# Deconvolution of cancer methylation patterns determines that altered methylation in cancer is dominated by a non-disease associated proliferation signal

**DOI:** 10.1101/2024.08.22.609153

**Authors:** H Lalchungnunga, H Atasoy, EC Schwalbe, CM Bacon, G Strathdee

## Abstract

All cancers are associated with massive reorganisation of cellular epigenetic patterns, including extensive changes in the genomic patterns of DNA methylation. However, the huge scale of these changes has made it very challenging to identify key DNA methylation changes responsible for driving cancer development. Here, we present a novel approach to address this problem called methylation mapping. Through comparison of multiple types of B-lymphocyte derived malignancies and normal cell populations, this approach can define the origins of methylation changes as proliferation-driven, differentiation-driven and disease-driven (including both cancer-specific changes and cancer absent changes). Each of these categories of methylation change were found to occur at genomic regions that vary in sequence context, chromatin structure and associated transcription factors, implying underlying mechanistic differences behind the acquisition of methylation at each category. This analysis determined that only a very small fraction (about 3%) of DNA methylation changes in B-cell cancers are disease related, with the overwhelming majority (97%) being driven by normal biological processes, predominantly cell proliferation. Furthermore, the low level of true disease-specific changes can potentially simplify identification of functionally relevant DNA methylation changes, allowing identification of previously unappreciated candidate drivers of cancer development, as illustrated here by the identification and functional confirmation of *SLC22A15* as a novel tumour suppressor candidate in acute lymphoblastic leukaemia. Overall, this approach should lead to a clearer understanding of the role of altered DNA methylation in cancer development, facilitate the identification of DNA methylation targeted genes with genuine functional roles in cancer development and thus identify novel therapeutic targets.

## Introduction

Cancer development is a multistep process involving the accumulation of genetic and epigenetic changes, including dramatic genome-wide alterations in the pattern of DNA methylation [1]. This includes global reductions in the level of 5-methylcytosine across the cancer genome versus normal cells, in association with the acquisition of methylation (hypermethylation) at CpG island sequences that typically have low methylation [2]. Hypermethylation of CpG islands has long been known to be important in tumour development and several well established tumour suppressor genes (TSG), such as *BRCA1* and *MLH1*, are known to be primarily inactivated by promoter hypermethylation [3]. However, array and sequencing based approaches that allow genome-wide assessment have identified that thousands of gene-associated CpG islands can become aberrantly methylated in a single tumour, although likely only a small number of these have a direct functional role [1]. In addition, while for genetic changes, a high mutation frequency can be taken as a proxy for functional relevance, many of the identified methylation changes were found to be present in essentially all cases of a particular tumour type and so a high frequency of occurrence of specific methylation changes does not provide evidence of a functional role in cancer development [4]. Thus, there is no clear mechanism for identifying the small number of methylation changes that are potentially important in driving tumour development from the overwhelming majority of passenger changes.

As an initial step to overcome this limitation we recently reported a novel bioinformatic approach that utilised genome wide DNA methylation patterns to identify synthetic lethal-like (referred to as subtype specific vulnerability genes) genes that are specific for individual molecular subtypes within a certain cancer type, initially in acute lymphoblastic leukaemia and medulloblastoma [5]. As part of this study, we also demonstrated a very strong overlap between altered methylation seen when comparing acute lymphoblastic leukaemia (ALL) cells to normal progenitors [6] with differences seen between normal memory B-cells and normal progenitors. Indeed, using a set of over 7,000 CpG sites originally identified as altered in acute lymphoblastic leukaemia (ALL), this analysis found that methylation change in ALL was highly predictive of changes in other B-cell malignancies and was highly predictive of mirroring methylation change in healthy memory B cells. Furthermore, the extent of methylation change in normal memory B-cells was essentially indistinguishable from more indolent B-cell malignancies such as chronic lymphocytic leukaemia (CLL) [5].

This demonstrates that many of the methylation changes seen in B-cell malignancies are not disease-specific, are independent of differentiation status, and are also present in specific populations of normal B-cells, which are long-lived and highly proliferative [7], like cancer cells. In addition, a number of other studies of B-cell malignancies have found similar results, with normal memory B-cells showing methylation changes that are highly overlapping with cancer derived from B-cells [8–10]. These results suggest that proliferation, independent of cellular transformation, may be the primary driver of altered methylation.

The similarity in the overall patterns of disrupted methylation across all malignancies [4] suggests that the driving forces underlying altered DNA methylation in different cancer types are likely to be highly similar. Most tissues lack populations of normal cells that, like memory B-cells, are long-lived and highly proliferative, that could be used for comparison to cancer cells. However, similarities observed between ageing-related methylation and cancer-related methylation in multiple tissue types [11] and similarities to altered methylation in cells reaching replicative senescence [12] is consistent with the hypothesis that proliferation may be the primary driver of methylation changes.

These observations not only contribute to our understanding of the basis of altered DNA methylation in cancer, but also suggest a potential approach to overcome the difficulty in identifying methylation changes in cancer cells that are functionally important in disease development. By screening out methylation changes that are known to also occur in normal cells following proliferation, true cancer-associated methylation changes could be identified. Taking advantage of the naturally occurring long-lived, highly proliferative memory B-cells, this study aims to “map” the derivation of all methylation changes occurring in B-cell malignancies and to classify them into four specific types: 1) Proliferation driven methylation changes, 2) Differentiation driven methylation changes, 3) True cancer specific changes (i.e. changes occurring in malignant B-cells but absent from normal memory B-cells) and 4) cancer absent methylation changes (i.e. changes occurring in memory B-cells but absent from malignant B-cells, suggesting direct selection against these methylation changes in the cancer cells). We hypothesised that the final two categories will include only a small fraction of the total methylation changes and that these methylation changes will have a much greater likelihood of being functionally relevant for disease development.

## Methods

### Data used for bioinformatic analysis

All datasets used in this study are from publicly available sources. The following methylation datasets were obtained from the Gene Expression Omnibus or the European Genome-Phenome Archive: B-cell progenitor cells GSE45459 [13] Naïve B-cells, memory non-switched cells (MEM_NCS) and memory switched cells (MEM_CS) EGAD00010000254 [14], for B-cell malignancies: ALL GSE49031 and GSE69229 [6, 15] CLL EGAD00010000254 [14], mantle cell lymphoma (MCL) EGAD00010001012 [16], diffuse large B-cell lymphoma (DLBCL) GSE37362 and TCGA-DLBC [17, 18], primary central nervous system lymphoma (PCNSL) GSE92676 [19]. Methylation data was processed as previously described [15].

Transcriptomic data sets used were as follows: Progenitor cell GSE45460 [13] Naïve B-cell, MEM_NCS, MEM_CS GSE24759 [20], ALL (GSE13159) [21], CLL EGAD00010000252 [22], MCL GSE93291 [23], DLBC GSE11318 [24], PCNSL GSE34771 [25].

### Bioinformatic and statistical analysis

All bioinformatic analyses were undertaken using R v3.4.0 (https://www.r-project.org/foundation). Differentially methylated regions (DMR), identified using DMRcate [26], were selected on the basis of an average beta value difference across the full DMR exceeding 0.2 (or 0.1, when stated), the minimum number of CpG sites to define a DMR was two and significance cutoff was p<0.0001. For the initial methylation mapping, DMR data sets were generated by comparing 1) B-cell malignancies (all five B-cell malignancies combined as a single dataset) vs B-cell progenitors and 2) Normal memory B-cells vs B-cell progenitors. The generated DMR datasets were then compared to allow characterisation of the identified DMRs using the following criteria (also illustrated in Figure 1):

**Figure 1.**
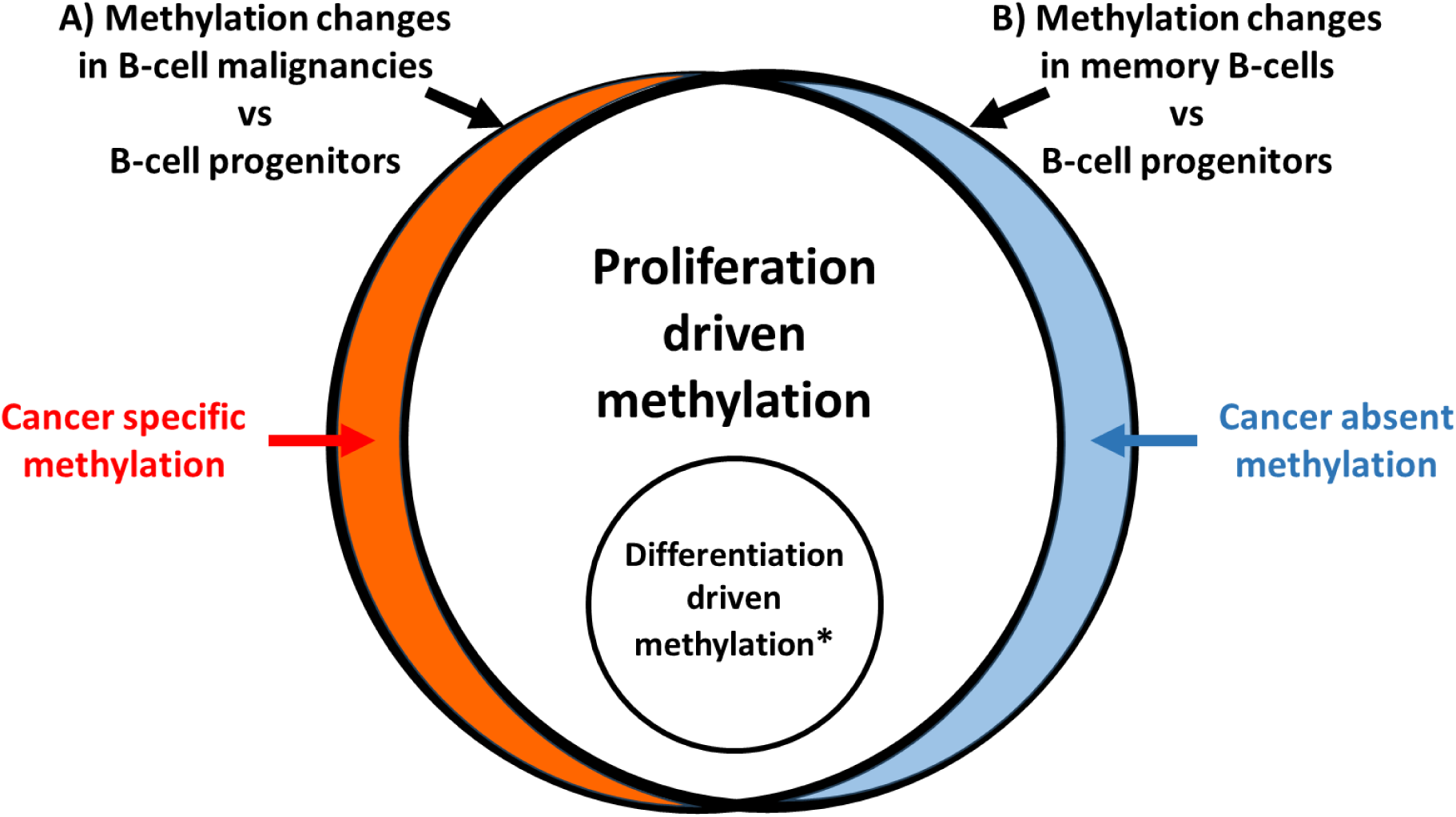
Identification of altered DNA methylation associated with cancer development. Initially methylation differences are identified separately from two comparison A) All B-cell malignancies (ALL, CLL, PCNSL, MCL, DLBCL) vs B-cell progenitors and B) Memory B-cells versus B-cell progenitors. DMRs that are identified in all B-cell malignancies and also in memory B-cells (in comparison to B-cell progenitors) are regarded as “proliferation driven). *Differentiation driven methylation changes are also present in both sets, but only shared with mature B-cell malignancies (CLL, PCNSL, MCL, DLBCL) and specifically absent from ALL. Cancer specific methylation changes represent a small minority of the total changes and include both “cancer specific methylation” (present in all B-cell malignancies but absent in memory B-cells) and” cancer absent methylation” (present in memory B-cells but absent in all B-cell malignancies).

For cancer specific methylation changes, criteria used to identify changes specific for all transformed B-cell population were: 1) Methylation change in all five B-cell malignancies combined vs B-cell progenitors >0.2 beta value. 2) Methylation change seen in each individual B-cell malignancy vs B-cell progenitors >0.1 beta value. 3) Methylation change seen in Memory B-cells vs progenitors is <0.1 beta value and at least four times smaller than change in each individual B-cell malignancy.

Cancer absent methylation changes met the following criteria: 1) Methylation changes seen in B-cell memory greater than >0.2 beta value as compared to B-cell progenitor. 2) Methylation change seen in B-cell memory cell is four times greater than methylation in all five B-cell malignancies combined. 3) Methylation change seen in all individual B-cell malignancies is <0.1 beta value.

For proliferation related methylation changes, multiple criteria were defined to ensure DNA methylation changes are due to B-cell proliferation: 1) Methylation changes in total proliferated B-cells (B-cell malignancies and B-cell memory cells) are greater than >0.2 beta value compared to B-cell progenitor. 2) Methylation change seen in all individual B-cell malignancies as well as in B-cell memory B-cells compared to B-cell progenitors is >0.1 beta value.

For methylation changes due to B-cell differentiation: 1) Methylation changes in differentiated B-cells (i.e., B-cell malignancies (with the exception of ALL) and B-cell memory cells) vs B-cell progenitors >0.2 beta value. 2) Methylation change seen in an individual differentiated B-cells (B-cell malignancies and B-cell memory cells), with the exception of ALL, vs B-cell progenitors >0.1 beta value. 3) Methylation change in ALL vs B-cell progenitors <0.1 beta value.

For individual disease specific methylation changes, criteria used to identify changes were: 1) Methylation change in the specific disease vs B-cell progenitors >0.2 beta value. 2) Methylation change seen in all other B-cell malignancies and normal memory B-cells vs B-cell progenitors <0.1 beta value. To increase stringency, a third criteria 3) Methylation change in the specific disease vs B-cell progenitors at least four times greater than methylation change vs B-cell progenitors for any of the other diseases or memory B-cells, was also included where indicated.

Identification of disease specific dependency genes and disease specific tumour suppressor genes was performed using our previously developed bioinformatic approach [5] with the exception that the five different B-cell malignancies were used as the “subtypes” for the analysis.

The SeSAMe package [27] was used to identify enriched genomic features within the filtered DMRs. For each comparison, SeSAMe was used to perform Fisher’s Exact tests for the enrichment of defined genomic feature sets, comparing the local genomic features defined by CpGs within a set of DMRs against a universe of genomic features associated with all probes within the Illumina Human Methylation 450k array; two-sided testing was performed throughout. These genomic feature sets were CpG island context (i.e. location with respect to proximal CpG islands), chromHMM (chromatin state), HMconsensus (histone modification), and TFBSconsensus (transcription factor binding sites (TFBS). Enrichments were visualised using SeSAMe.

Statistical assessments of differences in cell growth and apoptosis were carried out using t-tests, assuming equal variances, with p-values less than 0.05 deemed to be statistically significant.

### Cell culture

All leukaemia and lymphoma cell lines were cultured in RPMI 1640 media with L-glutamine and sodium bicarbonate (Sigma-Aldrich, UK). 293T cells were cultured in Dulbecco’s Modified Eagle’s Medium with 4500mg/l glucose, L-glutamine supplemented with foetal calf serum (Gibco, UK). Mycoplasma infection of cell lines was regularly checked for by using Mycoalert® Detection Kit (Lonza, Basel, Switzerland). Cells grown in an incubator at 37⁰C and 5% CO2.

### RNA extraction and cDNA synthesis

Total RNA was extracted using a GeneJET RNA Purification Kit (Thermo Fisher Scientific, Cat No: K0731) according to the manufacturer’s protocol. The extracted RNA was stored at -20oC (for use in the short term) or at -80oC if long-term storage was required.

The purified RNA was quantified using the Nanodrop ND-1000 spectrophotometer (Nanodrop, Delaware, USA). About 2μg of RNA was used for cDNA synthesis using the High-Capacity cDNA Reverse Transcription Kit (Applied Biosystems, UK Cat No: 4368814) according to the manufacturer’s protocol. Samples with no reverse transcriptase (RT) were also included in the reaction to act as a control in subsequent PCR reactions.

### Quantitative RT-PCR

All qPCR reactions were carried outed using Platinum SYBR Green qPCR SuperMix-UDG with ROX (Invitrogen, UK) following to the manufacturer’s protocol. Total volume per reaction was the 10μl. This includes 5μl Syber Green Master mix, 2μl CDNA, 0.8μl primers mix (Eurofins Genomics, Luxembourg) at 300ng/μl and 2.2 μl water to top up 10μl. Each sample was performed in triplicate in a 384-well plate ((Thermo Fisher Scientific)) and sealed with MicroAmp optical adhesive film (Applied Biosystem). A water control, and no RT controls were included along with assay samples. The samples were run on a QuantStudio 7 Flex Real-Time PCR system and analysed using QuantStudio Real-Time PCR software v1.2 (both Applied Biosystems). *GAPDH* was used as reference.

### Apoptosis assays

Apoptosis was assessed by two methods. Firstly, using the APC Annexin V (APC-labelled) and PI staining kit (BioLegend Cat No: 640932). This was performed according to the manufacturer’s instructions to stain 100,000 cells for each cell line/condition and performing flow cytometry using the FACS Fortessa X-20. Apoptosis was assessed at multiple time points post-transduction, as indicated in the text. FCS Express 7 software was used to analyse the results. Further confirmation of apoptosis was performed using the Caspase-Glo 3/7 Assay (Promega UK, Cat No:8091) system. Caspase 3/7 activity was measured at days 3, 4 and 5 post-transduction in 96-well plates, according to the manufacturer protocols. Blank wells, with only reagent and media without cells, were used as blank controls and untreated (parental) and mock-treated cells were used as negative controls. All experiments were performed in triplicate. Luminescent readings were taken using the FLOUstar Omega plate reader (BMG LABTECH).

### Assessment of cell growth

Staining with eBioscience™ Cell Proliferation Dye eFluor™ 450 (Invitrogen) was used to asses cell proliferation in transduced leukaemia cell lines. At day3 post-transduction, 250,000 cells were washed twice with pre-warmed PBS, re-suspended in 500μl PBS and then stained with cell proliferation dye according to the manufacturer’s instructions. Samples were then analysed at multiple subsequent time points (as indicated in text) until day 17 post-transduction, after which fluorescent levels dropped too low to see detectable differences after additional rounds of cell division. FCS Express 7 software was used to analyse results.

### Lentiviral transduction

The pSINE-SIEW vector (gift from Dr Paul Sinclair) was used for lentiviral transductions. This vector allows for high efficiency expression of cloned sequences, in addition to eGFP (expressed from the same transcript, but as a separate protein via an internal ribosome entry site). The THEM4 donor clone was purchased from GeneCopoeia (GC-11844), TTC12, MAP9 and SLC22A15 donor clones were purchased from as Genescript. Subcloning into pSIN-SIEW-GTW plasmid was performed using the gateway technologies (Gateway™ LR Clonase™ II Enzyme Mix, Invitrogen) as per manufacturer’s protocol. Generation of lentiviral particles for each clone was performed by co-transfection of the appropriate lentiviral expression vectors, along with the pMD2.G and pCMVΔR8.91 packing vectors, into the 293T packing cell line, using the EndoFectin Lenti transfection reagent (GeneCopoeia).

Transduction with lentiviral particles was performed using 2.5 x 10^5^ target cells/well in a 12-well plate at in 1ml fresh RPMI1640 media, containing 5% calf foetal serum then 500µl lentiviral-containing media was added to each well in a total volume of 1.5ml. This was incubated at 37°C and 5% CO2 for 16-20 hours. Untreated (parental) and mock-treated cells were used as negative controls. After incubation, at post-transduction day1, cells were washed twice with PBS and 1.5ml fresh RPMI 1640 media added. This was then incubated in the standard growth conditions and transduced cells were analysed by flow cytometry at multiple time points post-transduction, as indicated in the text.

### Western Blotting

Proteins were extracted using sonication after treating with SDS-lysis buffer. Pierce BCA protein Assay Kit (Thermo Fisher Scientific) was used to estimate protein concentration. 15-30µg of proteins were loaded onto gels (Mini-protein precast genes, Bio-Rad). After electrophoresis, proteins were then transferred onto a onto Polyvinyl difluoride transfer membranes (Immuno-Blot PDVF Membrane, Biorad, UK). Membranes were blocked in 5% dried skimmed milk with 0.1% Tween-20 (TBST) for 1 hr at room temperature, after which they were probed with primary antibodies overnight at 4°C. Primary antibodies used: anit-THEM4 (1:1000, Novus Bio (NBP2-94437)), anti-SLC22A15 (1:1000, Proteintech (20626-1-AP)) and anti-B-actin (1:10000, Abcam (ab49900)). Following incubation with the corresponding horseradish peroxidase-conjugated secondary antibodies, immunoreactive bands were visualized by enhanced chemiluminescence detection (Pierce ECL Western Blotting Substrate, Thermo Fisher Scientific Cat No: 32106) and ChemiDocTM MP imaging System (Bio-Rad).

## Results

### Methylation changes in B-cell malignancies are dominated by a proliferation signature

To map the derivation of methylation differences in B-cell malignancies, we identified differentially methylated regions (DMRs) identified between multiple types of B-cell derived malignancies and normal B-cell populations. This included malignancies arrested at the progenitor stage of differentiation (ALL) and at mature stages of B-cell differentiation (CLL, DLBCL, MCL and PCNSL) (Figure 1) and normal healthy B-cells at an early stage (B-cell progenitors) and a late stage (memory B-cells) of differentiation. B-cell progenitors are used as the “baseline” as, in addition to being normal/non-cancerous, they will have a relatively limited proliferation history (i.e. the total number of historical rounds of proliferation will be low for these cells). Inclusion of the memory B-cell population allowed identification of methylation changes that occur following proliferation of B-cells, independent of cellular transformation. Thus, methylation differences shared in the comparison of all B-cell malignancies to B-cell progenitors and memory B-cells to B-cell progenitors must result from processes independent of transformation (as they are present in normal memory B-cells) and independent of differentiation status (as present in ALL as well as differentiated normal and malignant cell types) (Figure 1). This analysis also allows identification of methylation changes specific for differentiation, which will be absent in normal and cancer cells derived from undifferentiated populations (i.e. normal B-cell progenitors and ALL) but present in differentiated normal and cancer cells (i.e. Memory B-cells, CLL, DLBCL, MCL and PCNSL) (Figure 1). The specific criteria used for identification of all DMR groups is detailed in Supplementary Figure 1.

Identification of DMRs associated with the normal cell processes of proliferation and differentiation then enables the identification of two categories of DNA methylation changes that are specifically associated with the development of B-cell malignancies. As shown in figure 1, we refer to these as “cancer specific”, to describe changes that occur in all B-cell malignancies (relative to the B-cell progenitor baseline) but do not occur in normal memory B-cells and a second category of differential methylation which we refer to as “cancer absent”, that is regions of differential methylation seen in memory B-cells (relative to the B-cell progenitor baseline), that are not replicated in any of the B-cell malignancies.

Initially, the analysis was performed by combining all types of B-cell malignancy together to allow identification of methylation changes that were specific for development of malignancy in B-cells as a whole (as opposed to being specific for a single type of B-cell malignancy). As shown in Figure 2, the overall altered methylation landscape of B-cell malignancies was highly similar to altered methylation seen in normal memory B-cells (r=0.94, p<0.0001) and was dominated by a proliferation signal (i.e. methylation changes that were present across all individual types of B-cell malignancy and also in memory B-cells), with 87.9% of DMRs mapping to this category (Table 1). A further 9% of DMRs were found to map to the differentiation category (i.e. alterations preserved across all mature B-cell malignancies (but not ALL) and in memory B-cells). By contrast, only 3% (174/5692) of DMRs mapped to the two categories in which methylation differences are specifically associated with malignancy (cancer specific, 2.7% and cancer absent, 0.3%) (Table 1).

**Figure 2.**
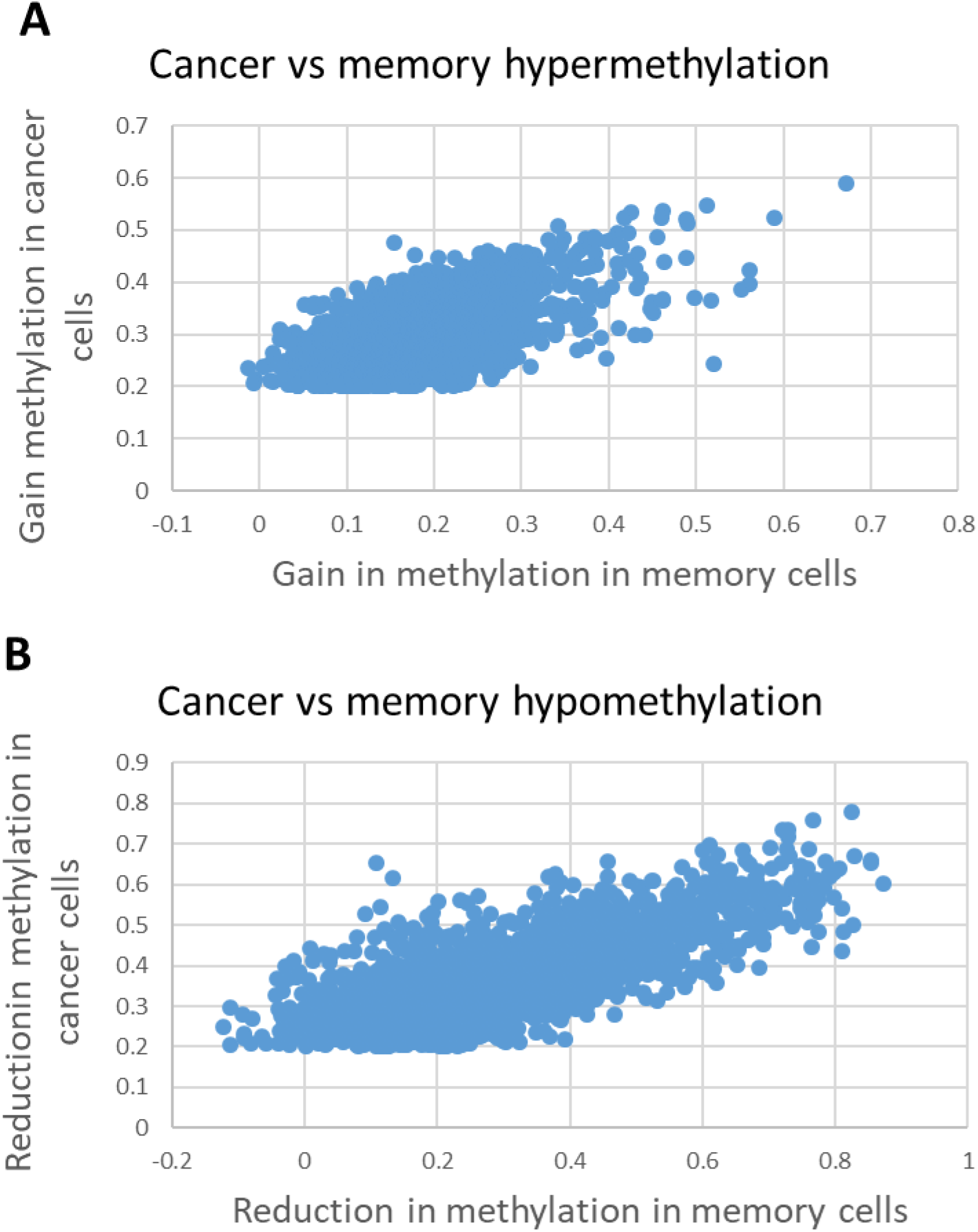
Methylation changes in normal memory cells closely mirror B-cell malignancies. Methylation differences (DMRs) were identified by comparing all B-cell malignancies combined vs B-cell progenitors. The level of methylation changes in the B-cell malignancies in DMRs that were hypermethylated **(A)** or hypomethylated **(B)** were then compared to methylation differences at the same DMRs in the comparison of normal memory B-cells vs B-cell progenitors. Overall, changes in the DMRs identified in B-cell malignancies were found to be largely replicated in normal memory B-cells (Pearson correlation coefficient r=0.94).

**Table 1.** Derivation of DMRs in B-cell malignancies.

| Table 1 - Derivation of DMRs in B-cell malignancies |  |  |  |  |
| --- | --- | --- | --- | --- |
| DMR group | Total DMRs<sup>a</sup> |  | Fraction of total DMRs (%) |  |
|  | 0.2 cut-off | 0.1 cut-off | 0.2 cut-off | 0.1 cut-off |
| Cancer specific | 156 | 224 | 2.7 | 1.7 |
| Cancer absent | 18 | 56 | 0.3 | 0.4 |
| Differentiation | 514 | 4091 | 9.0 | 31.5 |
| Proliferation | 5004 | 8628 | 87.9 | 66.4 |

<sup>a</sup>Total DMRs from a comparison of B-cell progenitors vs all B-cell malignancies combined

The analysis performed above focuses on DMRs with relatively large differences in DNA methylation (using a minimum beta value difference across the DMR of 0.2, approximately equivalent to a difference of 20% methylation). To examine the possibility that the low level of cancer specific changes was related to a focus on comparatively large differences in methylation, we repeated the full analysis for all datasets adjusting the cut-off for a significant DMR to a beta value difference of 0.1 (i.e. about 10% difference in methylation). As expected, this resulted in a large increase in the total number of DMRs identified when comparing B-cell malignancies to normal B-cell progenitors (from 5692 to 12,999, Table 1). Similarly, the total number of potential cancer specific DMRs increased (from 174 to 280). However, this represented a fall in the fraction of DMRs that mapped to the two cancer specific categories (from 3% to 2.1%). Methylation changes associated with normal cell processes still dominate the pattern, although a higher fraction of differentiation related changes was seen with this reduced cut-off (Table 1). These results demonstrate that, at least in B-cell derived cancers, the number of methylation changes that are specifically associated with disease development is far smaller than previously believed. This represents only a very small fraction of the differences identified by comparing cancer cells to normal cells which have not been through extensive proliferation. Even matching differentiation status would only remove a modest fraction of the non-disease related methylation differences (Table 1).

### Characterisation of DMRs mapping to proliferation, differentiation and cancer-specific categories

Assessment of the characteristics of the DMRs mapping to different categories suggests differences within the types of sequences present in each group. Proliferation associated DMRs exhibited significantly higher numbers of CpG sites (average 7.3 CpGs/DMR vs 3.5-4.4 for other groups, p<0.0001, Table 2) and were more likely to be closely proximal to transcriptional start sites (34% of DMRs were <1000bp from the nearest start site versus 19-28% for other groups, p<0.0001). Furthermore, there were pronounced differences in the fraction of hypermethylated versus hypomethylated DMRs in the different groups. Proliferation associated DMRs were relatively evenly divided, with 53.4% of DMRs hypermethylated, while the differentiation associated DMRs were heavily skewed towards hypomethylated DMRs, with only 5.1% hypermethylated (p<0.0001 vs proliferation DMRs). Cancer specific DMRs lay between proliferation and differentiation with 19.9% DMRs hypermethylated (p<0.0001 vs both). Cancer absent DMRs were predominantly hypermethylated (88.9%, P<0.01 vs any other group), although the absolute number of DMRs (18) was small.

**Table 2.**
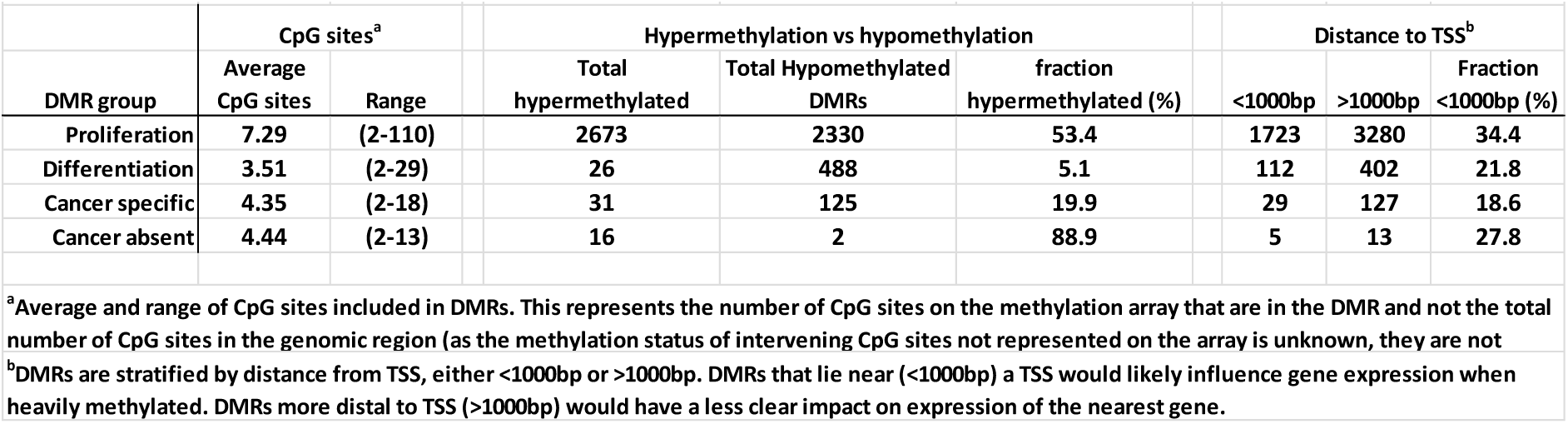
Characteristics of DMR groups.

### Identification of Zap70 as a functionally important target across all B-cell malignancies

A key aim of methylation mapping was that screening out of methylation changes that are driven by non-disease phenomena would allow identification of true cancer-related methylation changes. This should facilitate the identification of alterations in methylation that are of functional relevance in disease development. To this end, transcriptome datasets from each of the individual cancer types and normal cell populations were investigated to identify if any of the cancer specific or cancer absent changes (using the 0.2 beta value cut-off) exhibited significant negative correlations with expression from the nearest gene. Of the 174 cancer specific/absent DMRs identified, 10 exhibited a significant correlation with differential expression (Table 3). However, these loci exhibited features suggestive that this correlation may, in many cases, not be indicative of a direct role for differential methylation in controlling gene expression; the loci do not show an excess with the expected negative correlation (6/10 exhibit a positive correlation with gene expression), 6/10 are regions of low CpG density (only 2-3 CpG sites in the DMR) and 5/10 are highly distal (>40Kb) from the nearest TSS and only 2/10 are within the proximal promoter regions of a gene (<1kb from TSS, *SPATA6*, *ZAP70*) (Figure 3). *ZAP70* was the clearest candidate for a pan B-cell malignancy driver, and the only one of the two candidates with a beta value difference >0.2 (versus normal progenitors or memory cells) in all B-cell malignancies (Figure 3, Table 3). Consistent with this, the *ZAP70* gene has been clearly implicated in the development of chronic lymphocytic leukaemia [28]. However, the consistent differences in methylation and expression across all B-cell malignancies suggest that analysis of its wider role across many B-cell derived malignancies would be merited.

**Figure 3.**
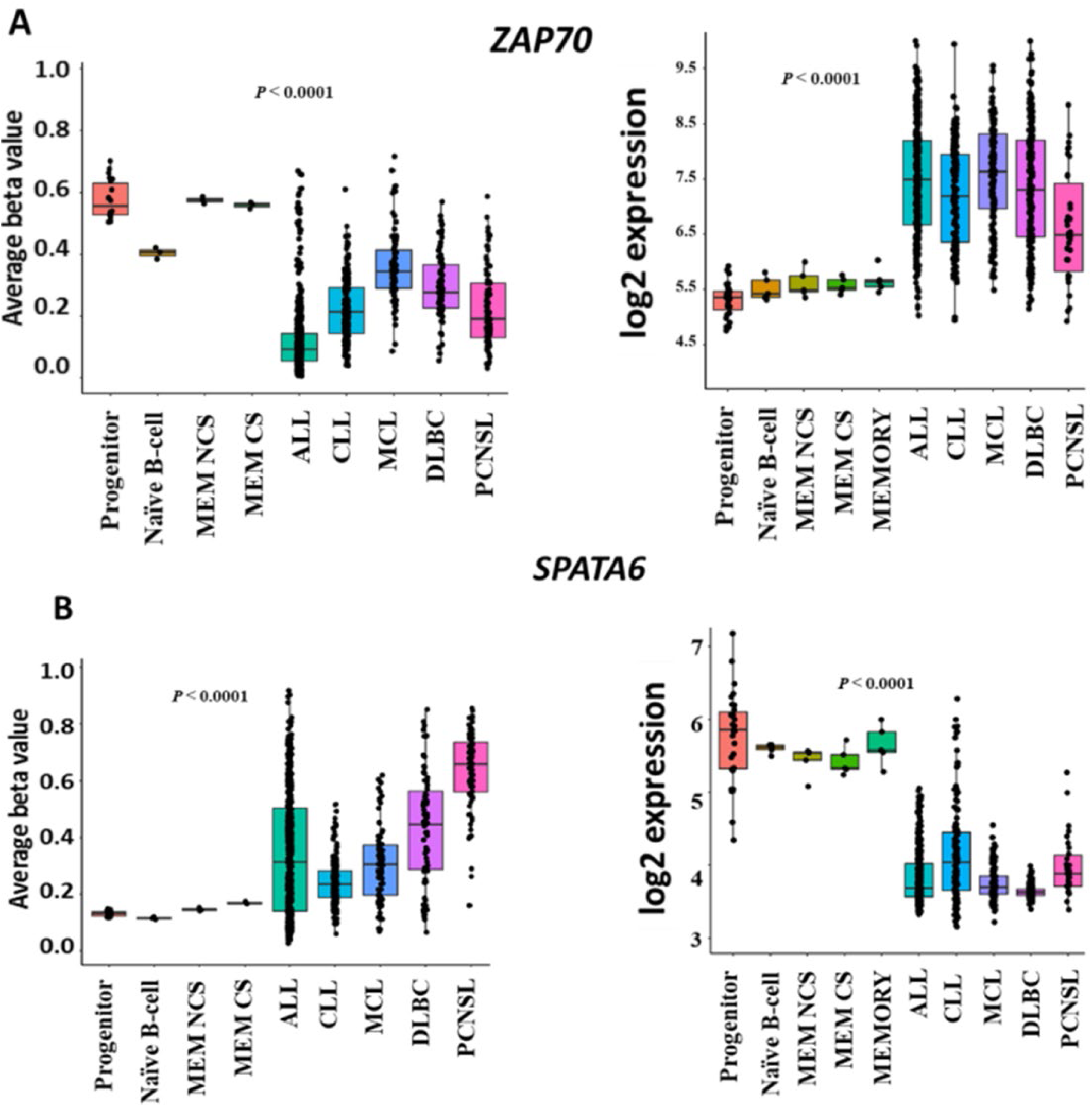
Loci exhibiting B-cell malignancy specific methylation and expression patterns. Methylation (left-hand graph) and expression (right-hand graph) patterns for the three loci identified with DMRs located in their proximal promoter regions and exhibiting a negative correlation with gene expression. A) ZAP70 and B) SPATA6. Methylation and expression is shown for normal cell types (B-cell progenitors (progenitor) and class-switched memory B-cells (MEM CS)) and for the five B-call malignancies used in this study.

**Table 3.**
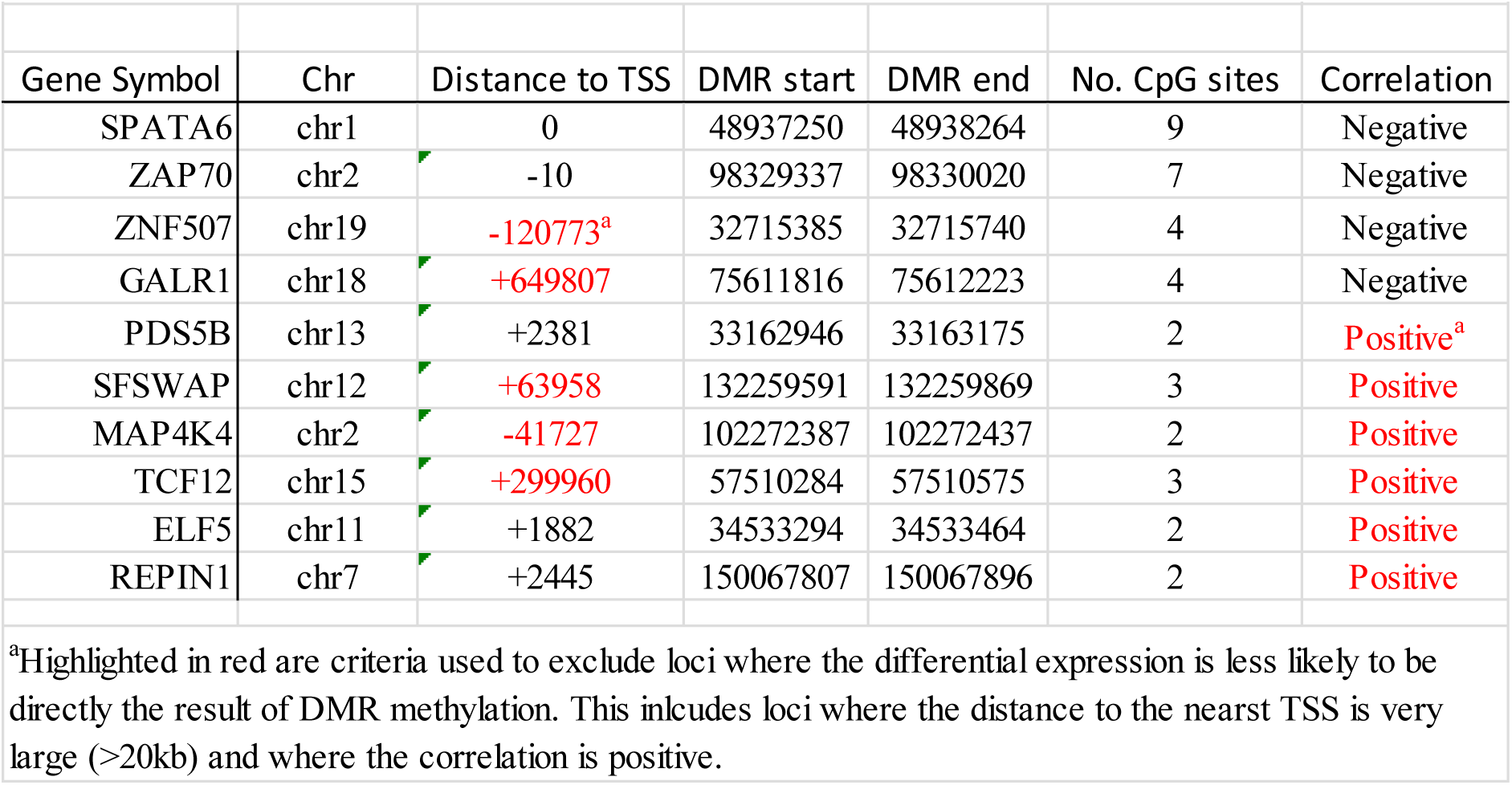
Cancer related DMRs with correaltion to gene expression.

### Methylation mapping applied to individual cancers

The above results suggest that despite the large number of methylation changes shared across B-cell malignancies, only a very small fraction of methylation changes are truly cancer-specific. However, it is possible that most key functional changes in DNA methylation may be specific for individual types of B-cell cancer. To assess this possibility, we repeated the analysis examining each individual B-cell malignancy individually to identify DMRs that were specific for individual disease types (Supplementary Figure 1E). Disease specific DMRs were defined as DMRs present in the specific disease vs B-cell progenitor comparison (with a beta value difference >0.2) but absent in any of the other groups (i.e. the other four B-cell malignancies and memory B-cells) versus B-cell progenitors. “Absence of change” was defined as <0.1 beta value change or a change in the opposite direction, so that minor differences in methylation in other diseases didn’t prevent the identification of disease specific DMRs. As can be seen from Table 4, this analysis also identifies a comparatively low fraction of methylation changes as being specific for individual disease, ranging from 0.1% to 5.4% across the five malignancies. Notably, three of the five malignancies exhibit only very small numbers of disease specific DMRs (6-17 DMRs or 0.1-0.15% of total DMRs for CLL, MCL and DLBCL (Table 4), indicating a near absence of disease specific methylation changes in these malignancies. In contrast the results for ALL and PCNSL suggest that disease specific methylation change may be for more relevant in these malignancies with 240 (4.8% of total DMRs) in ALL and 1104 (5.4% of total DMRs) in PCNSL. We have previously observed higher absolute levels of methylation change in PCNSL at loci that also acquire methylation changes in other B-cell malignancies (Schwalbe et al 2021). Thus, the high level of DMRs in PCNSL may primarily reflect a greater absolute size of methylation change at loci also changed in other diseases/memory B-cells, as opposed to true specificity. Consistent with this hypothesis, use of a more stringent definition for specificity of the methylation change (by defining “absence of change” in the other four cancer types as >4x lower than the change in the specific disease, as opposed to the original definition of anything <0.1) would retain only 28% of the “specific” DMRs in PCNSL (reducing the fraction of specific DMRs to 1.5% of total DMRs in PCNSL). Applying the same criteria to ALL would retain 78% of the DMRs, (reducing the fraction of specific DMRs to 3.4% of total DMRs in ALL).

**Table 4.** Disease specific DMRs.

| Table 4 - Disease specific DMRs |  |  |  |  |  |  |  |  |  |  |  |  |
| --- | --- | --- | --- | --- | --- | --- | --- | --- | --- | --- | --- | --- |
| Disease | Total DMRs | Disease related (0.2 cut-off)<sup>a</sup> |  |  |  |  | Disease related (0.1 cut-off)<sup>d</sup> |  |  | DMR characteristics (0.1 cut-off) |  |  |
|  |  | All disease related | Fraction of total DMRs<sup>b</sup> | Disease specific | Disease absent |  | All DMRs | All disease related | Fraction of total | average CpG sites | Fraction <1000bp to TSS | Fraction hypermethylated |
| ALL | 4988 | 240 | 4.80% | 240 | NA<sup>c</sup> |  | 14208 | 463 | 3.26 | 7.1 | 38.0 | 69.4 |
| CLL | 6326 | 6 | 0.10% | 2 | 4 |  | 18544 | 48 | 0.26 | 6.4 | 25.0 | 22.9 |
| MCL | 6659 | 10 | 0.15% | 4 | 6 |  | 21709 | 61 | 0.28 | 7.3 | 24.6 | 26.2 |
| DLBCL | 11701 | 17 | 0.15% | 7 | 10 |  | 27335 | 106 | 0.39 | 5.8 | 24.5 | 20.8 |
| PCNSL | 20583 | 1104 | 5.40% | 1093 | 11 |  | 34819 | 1612 | 4.63 | 6.4 | 33.3 | 3.4 |
| <sup>a</sup>DMRs (relative to B-cell progenitors) using a minimum beta value cut-off of 0.2 |  |  |  |  |  |  |  |  |  |  |  |  |
| <sup>b</sup>Fraction of the total DMRs that are disease related (including disease specific and disease absent) |  |  |  |  |  |  |  |  |  |  |  |  |
| <sup>c</sup>Disease absent DMRs cannot be identified in ALL as their methylation patterns would be identical to normal differentiation related DMRs |  |  |  |  |  |  |  |  |  |  |  |  |
| <sup>d</sup>DMRs (relative to B-cell progenitors) using a minimum beta value cut-off of 0.1 |  |  |  |  |  |  |  |  |  |  |  |  |

As with the analysis of all B-cell malignancies combined, reducing the stringency of defining DMRs by using a beta value difference from 0.1 (as opposed to the original 0.2) increases the total number of DMRs identified, but does not increase the fraction of those DMRs which are identified as disease specific (Table 4). Thus, analysis at the individual level also determined that only a very small fraction of DMRs identified in the individual diseases represent disease specific changes.

### Regions targeted in disease specific methylation show similarities to proliferation driven methylated regions

Characterisation of the DNA structure of the identified disease specific DMRs shows a number of disease specific differences, most notably that ALL shows a clearly higher fraction of hypermethylated regions compared with any of the other B-cell malignancies (p<0.0001 for all comparisons, Table 4) and also the highest fraction of DMRs located proximal to gene promoters (although differences from other diseases were not statistically significant, Table 4). In contrast, PCNSL exhibited a strong link with hypomethylation, with 96.6% of all PCNSL-specific DMRs being hypomethylated (p<0.0001 versus all other individual B-cell malignancies). For all five B-cell malignancies the disease specific DMRs exhibited relatively high CpG densities (average of 5.8-7.3 CpG/DMR) which were in a similar range to those seen for proliferation derived DMRs in the original analysis (average of 7.3 CpG/DMR) (Table 4).

To begin the process of investigating the potential molecular basis of the identified DMR groups, the SeSAMe package [27] was used to identify specific DNA/chromatin structural elements associated with the different DMR groups. The results of this analysis are outlined in Figure 4 (full details are in supplementary Figures 2-5). This approach assesses the different DMR groups across four domains: Sequence context, chromatin state, histone modifications and transcription factor binding sites. For this analysis, the larger DMR groups from the pan-b-cell cancer analysis (i.e. the proliferation and differentiation groups) were separated into hypermethylated and hypomethylated DMRs. Consistent with previous reports [29, 30], both proliferation and differentiation related hypermethylated DMRs are associated with CpG island and shore sequences, with polycomb complex regulated and bivalent promoter regions, and with regions linked to H3K9 and H3K27 trimethylation. In contrast, hypomethylated DMRs were linked to CpG sparse regions, with regions associated with quiescence and enhancer regions and with H3K36 trimethylation. Notably, the disease specific DMRs show patterns similar to the proliferation/differentiation hypermethylated pattern in terms of three of these domains and also show strong associations with associated transcription factor binding sites. This is true for ALL, in which the disease specific DMRs are predominantly hypermethylated, but is also true for CLL, MCL and DLBCL in which the DMRs are predominantly hypomethylated. Of note, in terms of histone modifications, all these groups of disease specific DMRs are specifically associated with regions linked to H3K27 trimethylation but not H3K9 trimethylation. The same pattern is also seen for differentiation associated hypermethylated DMRs, while proliferation DMRs are uniquely, and strongly, linked to H3K9 trimethylation. Disease specific DMRs identified in PCNSL, unlike the other diseases, exhibited patterns related to proliferation/differentiation hypomethylated region and is consistent with the extreme association within this group to hypomethylated DMRs. In contrast to the disease specific DMRs, the DMR groups identified as occurring across all B-call cancers (cancer absent and cancer specific) demonstrate few significant associations, although both groups demonstrated very strong links to enhancer regions.

**Figure 4.**
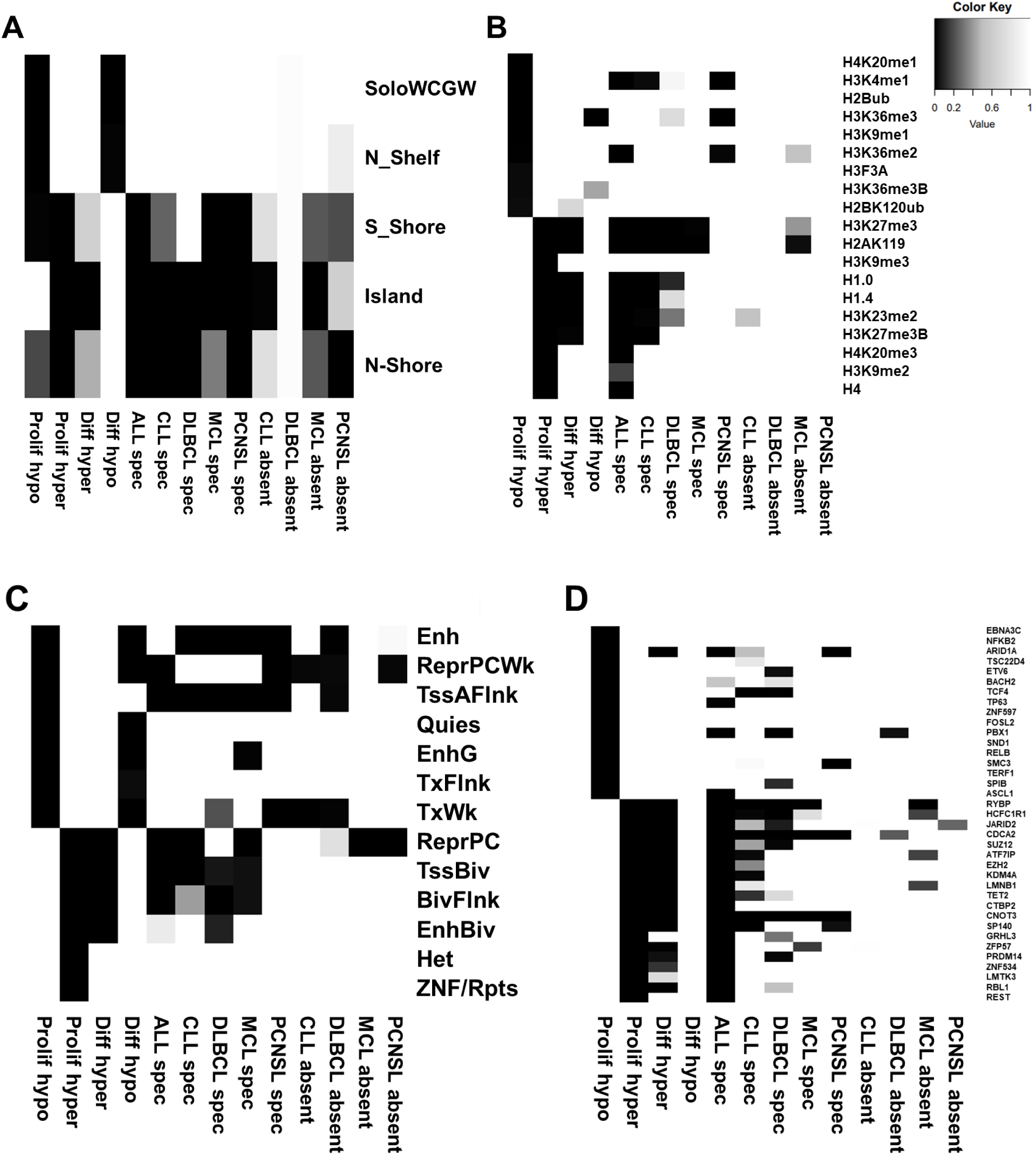
SeSaMe analysis of DMRs associated with normal and cancer specific methylation changes. SeSaMe analysis was carried out to compare the characteristics of DMR regions associated with normal and cancer specific processes. For all four domains up to 20 associations are shown for each group (with fewewer shown is there were less than 20 associations with significant (<0.05) FDR values. Associations are shown for **A)** Sequence context, **B)** Histone modifications, **C)** Chromatin States, **D)** Transcription factor binding sites. While the chromatin states (C) exhibit a mixed pattern, for the other three domains there is a pronounced overlap between the cancer specific associations in the B0cell malignancies and the normal methylation changes associated with hypermethylated regions.

**Figure 5.**
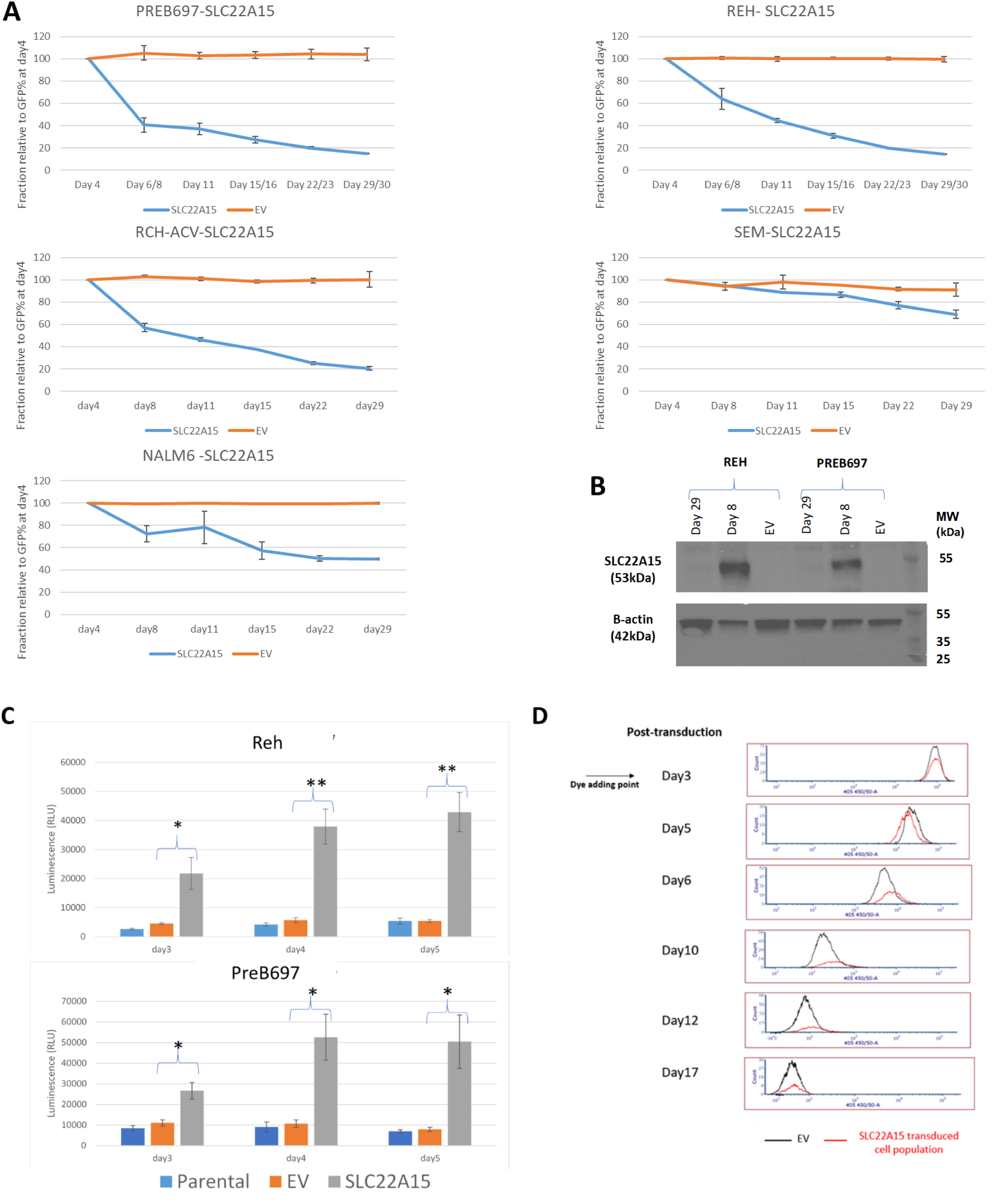
Selection against SLC22A15 expression across cell lines derived from multiple ALL genetic subtypes. A) Following transduction of ALL cell lines with a lentiviral vector expression SLC22A15 the SLC22A15 expressing cells were lost from the population in all cell lines. By contrast, a control empty vector (expressing only eGFP) remained stable throughout the period. **B)** SLC22A15 protein expression was assessed in PreB697 and Reh cells at day 8 and 29 post-transduction and in control empty vector transduced cells. Re-expression of SLC22A15 is clearly detectable at day 8, but is lost by day 29. **C)** Caspase activation was assessed post-transduction in the Reh and PreB697 cells. While transduction with the control vector (EV) had no detectable impact on caspase activation, a pronounced activation of caspase 3/7 was initially detectable at day 3 post-transduction with SLC22A15 expressing vector, increasing to a maximal level by day 4/5 in both cell lines. **D)** Proliferation of SLC22A15 or control EV transduced cells was assessed following staining with VPD450 (results are shown for the Reh cell line, although essentially identical results were also found in PreB697 cells). While a slight delay in proliferation appears to be present at days6-12, there is no evidence for a population of arrested cells following SLC22A15 re-expression.

The analysis of transcription factor binding sites may allow identification of factors that could play active roles in promoting methylation changes in the specific groups. Consistent with the association of most disease specific DMRs with the hypermethylated regions driven by normal processes of proliferation and differentiation, multiple transcription factor binding sites was found to be shared across all these groups, including a core set of five TFBS (CDCA2, CNOT3, HCFC1R1, RYBP, SP140) (Figure 4). Despite this similarity, associations with core PRC2 proteins (EZH2, SUZ12, JARID2) were amongst the strongest in the proliferation/differentiation DMR groups but absent or only very weakly associated in the disease specific DMR groups. TET2 binding, a factor strongly linked to myeloid malignancies, was found to associate specifically with proliferation DMRs and ALL specific DMRs (supplemental Figure 5X). Furthermore, most of the DMR groups are associated with a set of unique TF binding sites that may have disease/process specific functions (supplemental Figure 5X).

### Candidate gene identification in individual diseases

As for the cancer specific methylation changes, genome wide expression data for each of the five malignancies was assessed to identify genes linked to DMRs that showed reciprocal specific differences in gene expression (i.e. gene whose expression pattern show an equivalent, but opposite, specificity for the disease in which the DMR was identified). This analysis identified a total of 15 candidate genes across four of the five diseases (no candidates were identified for DLBCL). Analysis of ALL, which had the greatest fraction of disease specific methylation changes, also resulted in the identification of the largest number of candidate genes (Table 5).

**Table 5.** Identified candidate functional genes.

| Table 5 - Identified candidate functional genes |  |  |  |  |
| --- | --- | --- | --- | --- |
| Disease | Gene | Candidate type<sup>a</sup> | Methylation mapping<sup>b</sup> | Subtype Analysis<sup>c</sup> |
| ALL | *THEM4* | TSG | Yes | Yes |
| ALL | *MAP9* | TSG | Yes | Yes |
| ALL | *SLC22A15* | TSG | Yes | Yes |
| ALL | *TTC12* | TSG | Yes | Yes |
| ALL | *DTD1* | TSG | Yes | No |
| ALL | *NPY* | DSDG | Yes | Yes |
| ALL | *BMP2* | DSDG | Yes | Yes |
| ALL | *CTGF* | DSDG | No | Yes |
| ALL | *ZNF423* | DSDG | Yes | Yes |
| ALL | *SMIM3* | DSDG | Yes | No |
| ALL | *CMTM2* | DSDG | Yes | No |
| CLL | *SNX18* | DSDG | Yes | Yes |
| MCL | *SOX11* | DSDG | Yes | Yes |
| MCL | *PAPPA* | DSDG | No | Yes |
| MCL | *MFHAS1* | DSDG | No | Yes |
| PCNSL | *FSCN1* | TSG | Yes | Yes |
| PCNSL | *STMN4* | DSDG | Yes | No |
| <sup>a</sup>TSG; tumour suppressor candidate, identified as having disease specific hypermethylation and loss of expression. DSDG; disease specific dependency gene candidate, identified as having disease specific lack of acquired methylation and increased expression |  |  |  |  |
| <sup>b</sup>Also identified by the methylation mapping approach |  |  |  |  |
| <sup>c</sup>Based on our previously published approach (ref 5) |  |  |  |  |

### Similarity of candidates identified through adaption of an alternative bioinformatic approach for identification of subtype specific functionally relevant genes

The relatively small number of candidate genes identified in B-cell malignancies above could suggest that the methylation mapping approach lacks sensitivity. To investigate this possibility, we used an alternative approach for identification of functionally relevant targets for aberrant methylation and then assessed the extent of overlap between the two approaches. This second approach is based on our recently reported bioinformatic approach [5], which can be used to identify cancer subtype specific synthetic lethal and tumour suppressor genes. This approach takes advantage of the extremely high degree of overlap in methylation changes between molecular subtypes of the same cancer and was able to identify functionally confirmed synthetic lethal genes with a high specificity (Schwalbe et al 2021). The very high degree of overlap in DNA methylation changes between the different B-cell malignancies allows a similar approach to be used, utilising the different B-cell malignancies as the “subtypes” in the analysis. There are a number of key differences between this approach and the above described methylation mapping; 1) the original identification of DMRs is achieved by comparison of the disease under study versus other four diseases (and does not involve comparison to normal cell populations), 2) identified DMRs are further analysed to identify regions of biggest difference (consisting of two or more adjacent CpG sites) and the 3) cut-offs used to identify differential methylation in these regions of biggest difference are more stringent (beta value difference of 0.4, or 40% methylation difference). This approach identified a number of candidate functionally relevant genes across the five diseases, including five tumour suppressor candidates and eight synthetic lethal-like genes (which we refer to as disease specific dependency genes, as they are specific for a whole disease as opposed to a specific genetic/molecular change). As shown in Table 5, these exhibit a highly significant overlap with the candidates identified through methylation mapping (10/13 candidates identified by this approach (“Subtype Analysis”) were also identified through methylation mapping). Specific details of the methylation and expression of the identified candidates is shown in Supplementary Figure 6. While, due to the differences between the approaches, each approach identifies some additional unique candidates, the number of candidates identified remains low, further suggesting that the number of functionally relevant methylation changes in B-cell malignancies is low.

### Functional assessment of novel tumour suppressor candidates

Amongst the candidates that were identified by both approaches was a set of four candidate tumour suppressor gene candidates identified in ALL (*THEM4, MAP9, SLC22A15, TTC12*). As all four genes were identified by both approaches and have not been previously implicated in ALL, this set of candidates was chosen for assessment of functional importance to provide proof-of-principle that novel functional genes can be identified by methylation mapping. As the candidates were identified as relevant for ALL as a whole (i.e. for all/most ALL genetic subtypes) a panel of cell lines was chosen that included representatives of multiple different cytogenetic ALL subtypes. All cell lines were assessed for DNA methylation and gene expression of the candidate genes, which confirmed high levels of promoter associated methylation and lack of expression for all candidates. Re-expression of the candidates was achieved using transduction with a lentiviral expression construct, into which each of the genes were individually cloned, which also co-expresses eGFP from the same transcript to allow transduced cells to be identified by flow cytometry. Correct cloning and lack of genetic disruption of the candidates in the expression construct as confirmed by sequencing. One of the four candidates (*MAP9*) failed to exhibit any detectable expression in transduced populations and so its functional impact could not be determined.

Of the other three candidates, clear expression from the polycistronic transcript was detected four days post transduction (Figure 5, Supplementary Figure 7). For comparison, a corresponding “empty vector” control transduction (i.e. vector expressing only eGFP) was also carried out. For one of the three candidates, *TTC12*, expression levels (assessed by fraction of eGFP positive cells) remained essentially unchanged throughout the time course in all cell lines assessed, suggesting that *TTC12* is either not a tumour suppressor gene or that its function is not effectively modelled in cell lines. *THEM4* expression was rapidly lost in the PreB697 cell line (t(1;19) genetic subtype) but remained relatively constant in all other cell lines, including a second t(1;19) derived cell line (RCH-ACV). In contrast, following re-expression of *SLC22A15*, loss of eGFP/SLC22A15 positive cells was seen in all five ALL cell lines assessed (Figure 5). This was associated with a corresponding loss of SLC22A15 protein expression (Figure 5B). Furthermore, analysis of caspase activation (in Reh and PreB697 lines) demonstrated that re-expression of SLC22A15 was associated with rapid induction of caspases 3/7 (Figure 5C) in the absence of alteration in the underlying rate of proliferation (Figure 5D), suggesting that re-expression of SLC22A15 resulted in induction of cell death. Thus, analysis of the derivation of cancer-related methylation changes allows identification of a much smaller number of truly cancer specific changes and can enable the identification of novel functionally relevant genes (such as *SLC22A15*) that would have previously been challenging to identify from the huge number of methylation differences previously thought to be disease specific.

## Discussion

Following the development of genome wide methods for DNA methylation analysis it has been widely reported that essentially all cancer types exhibit extensive alterations in DNA methylation compared with equivalent normal cell types [1]. However, the results presented here suggest that only a very small proportion of these methylation changes can actually be regarded as truly disease-associated and instead, the predominant drivers of alterations in DNA methylation in cancer cells are normal cell processes, specifically proliferation and differentiation, with proliferation being the predominant driver of the most extensive alterations in DNA methylation. Thus for the B-cell malignancies assessed in this study, normal B Memory cells showed highly overlapping patterns of altered DNA methylation (compared to B-cell progenitors) with B-cell malignancies. Most of these changes were shared with all B-cell malignancies (proliferation-driven), whilst a smaller subset were only shared with other B-cell malignancies derived from differentiated B-cells (differentiation-driven). Although, this analysis was restricted to cancers derived from the B-lymphocyte lineage, the strong similarity in overall patterns of altered DNA methylation seen in all cancers [2] strongly suggests that the driving forces behind altered DNA methylation in cancer will be broadly similar.

An important aspect of the analysis performed here is that it is based on comparison of both cancer and normal cell populations to a “base state”, in this case B-cell progenitors. This allowed us to identify loci with related changes in DNA methylation even when absolute changes in DNA methylation were larger in some cancer types (such as ALL, DLBCL, PCNSL), likely a product of most B-cell cancers being more proliferative than the normal memory B-cells. Consistent with this, more indolent cancers (CLL and MCL) exhibited methylation changes at very similar levels to memory B-cells.

Two key aspects that lead to difficulty in identifying functionally relevant DNA methylation changes in cancer are the very large number of methylation changes and also the high frequency with which many of these DNA methylation changes occur [2]. This means that even determining that a particular gene is aberrantly methylated in almost all cases of a particular cancer type cannot be regarded as strongly suggestive of a functional role. The results presented here suggest that proliferation itself is the key driver of altered methylation and, as this is a feature of all cancers, the high frequency of invariantly aberrantly methylated genes is consistent with this hypothesis. If proliferation-driven methylated genes can be identified through analysis of normal cells before and after extensive proliferation and at different stages of differentiation, this approach would then allow identification of true disease related changes by subtracting out the overwhelming number of methylation changes related to proliferation or differentiation status. This approach allowed us to identify true disease related DNA methylation changes that occurred across all B-cell malignancies, as well as changes that were specific for individual B-cell malignancies.

The key aim of this approach is to identify DNA methylation changes that are truly disease specific and in this way enable the identification of alterations in DNA methylation that are important in driving disease development or progression. Initially we used this approach to look at DNA methylation changes occurring across all B-cell malignancies, to investigate the possibility of key drivers of transformation applicable to all malignant B-cell disease. Combined with corresponding analysis of gene expression this identified a single gene, *ZAP70*, as implicated across all B-cell cancers. ZAP70 has been extensively investigated for its role in CLL [28]. However, its role in other B-cell malignancies has received less attention [31] and the results presented here suggest that functional studies in all B-cell malignancies may be warranted. Expanding the analysis to identify DNA methylation events specific for individual B-cell malignancies allowed the identification of a larger number of candidate genes, including the identification of SLC22A15, a solute carrier with a specificity for zwitterions such as carnosine [32], as a novel tumour suppressor gene in ALL. Several recent studies have identified carnosine as a metabolite that shows anti-tumour activity against multiple different cancer types [33–35]. Thus, downregulation of carnosine uptake may be important in the functional role of SLC22A15 in ALL.

A feature of this approach is that it can theoretically be applied to identify DNA methylation changes associated specifically with any feature of cancer development or with any cancer type. Here we have concentrated on identification of DNA methylation changes associated with either all B-cell malignancies or associated with individual B-cell malignancies. However, previously we have used similar approaches to identify synthetic lethal candidates associated with haematological and solid tumours [5, 36] and also DNA methylation changes associated with chemotherapeutic drug resistance [37]. A similar approach could be used to investigate DNA methylation changes specific for essentially any feature of cancer development and progression by identifying DNA methylation differences between states and then subtracting out those known to be driven by proliferation/differentiation state. However, an additional challenge in expanding this approach to non-lymphocyte associated cancers would be the identification of normal cell populations that have undergone extensive proliferation, as for most tissue types, highly proliferative normal cells equivalent to memory B-cells, do no occur naturally. Thus, model systems that utilise age-related methylation or *in vitro* or *in viv*o models to induce proliferation in normal cells may be required.

The results presented here suggest at least three separate driving forces (proliferation, differentiation, disease development) underlie the DNA methylation changes seen in B-cell malignancies and likely other cancer types. Analysis of the chromatin features and transcription factor binding sites associated with these different groups found multiple differences in the sequences, locations relative to transcribed regions and associated chromatin structures, further emphasising the differences between these groups. This may provide a starting point to investigate differences in the underlying mechanisms responsible for these different groups of methylation change. Such studies could enable the identification of mechanisms that are predominantly or exclusively involved in the disease associated DNA methylation changes and thus potentially allow for strategies aimed at specifically reversing or preventing changes directly involved in cancer development without having more widespread effects on normal cellular epigenetic mechanisms.

Overall, the results presented here contribute to a much clearer understanding of the basis of the extensive altered DNA methylation patterns which characterise cancer development and open up new opportunities for utilising true disease associated methylation changes to identify novel therapeutic targets.

---

**Supplementary Figure 1.**
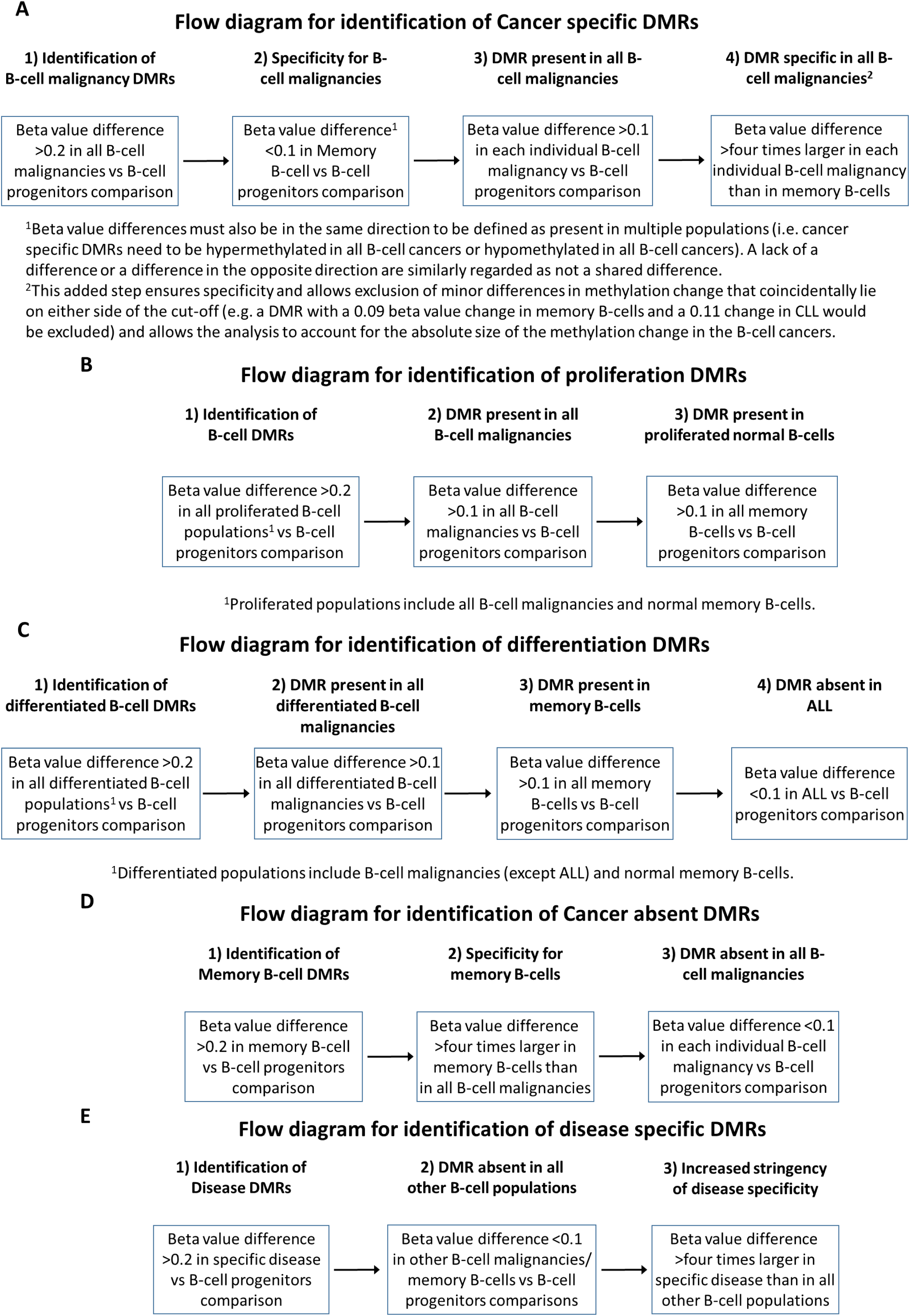

**Supplementary Figure 2.**
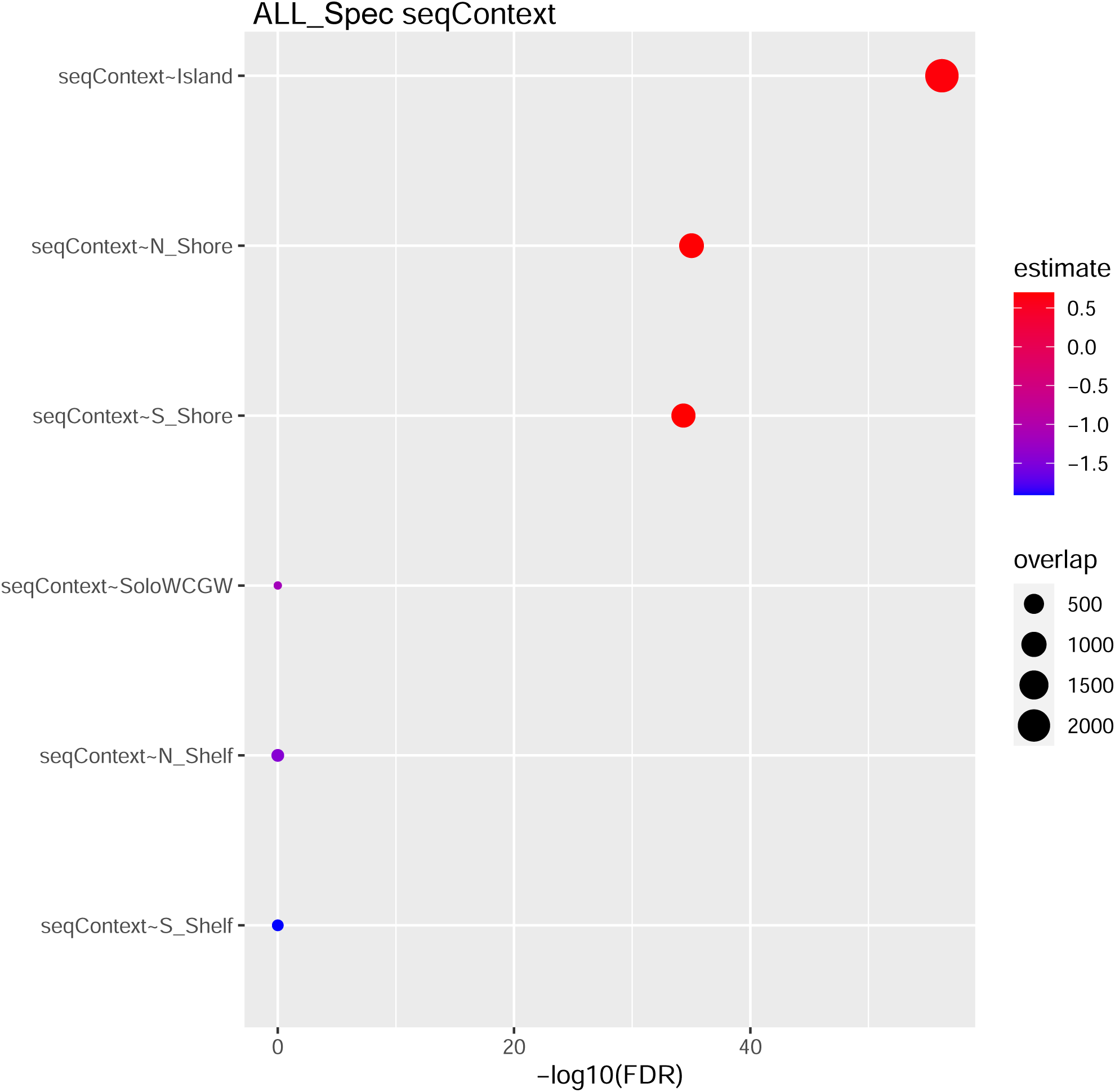
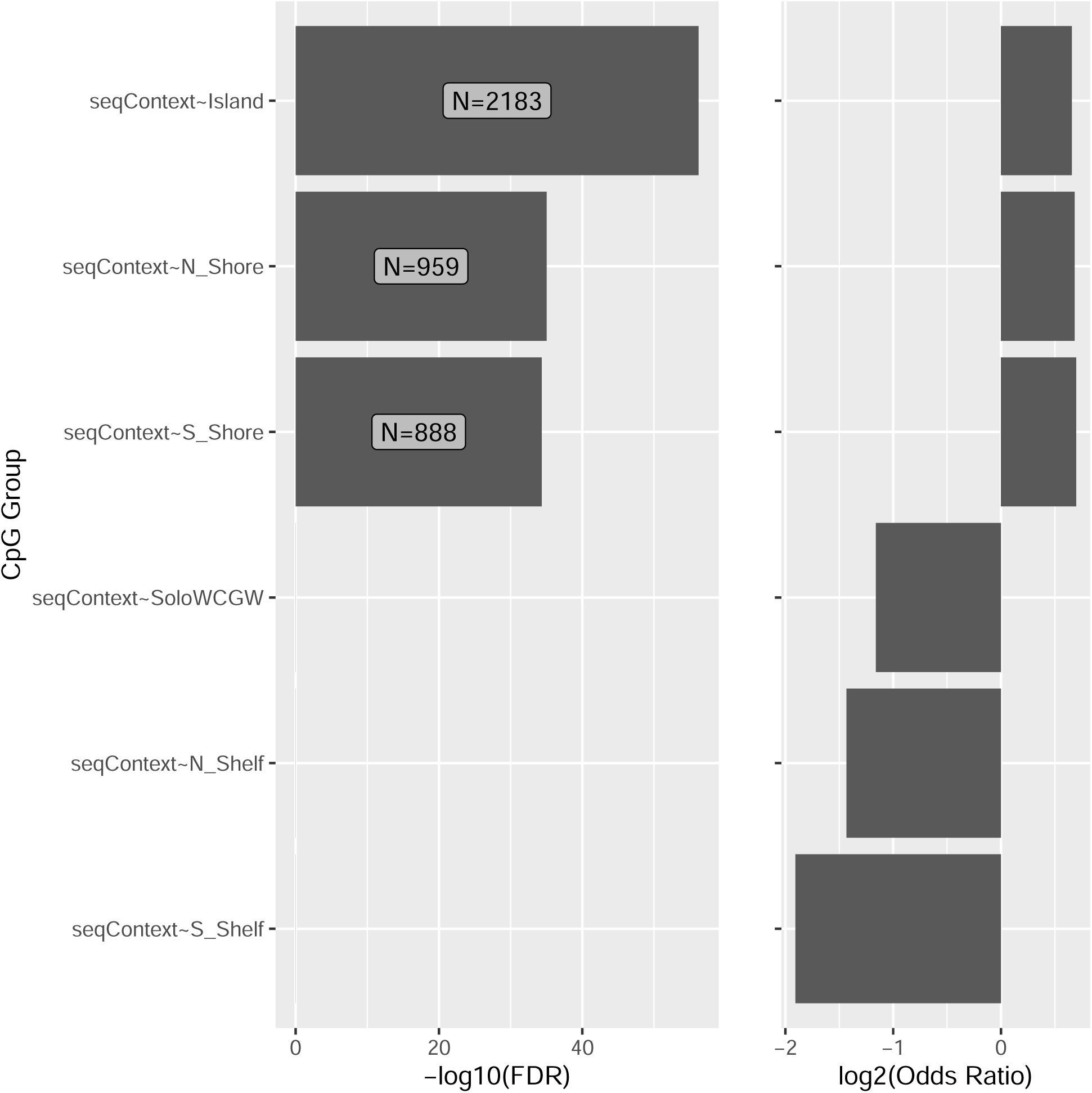
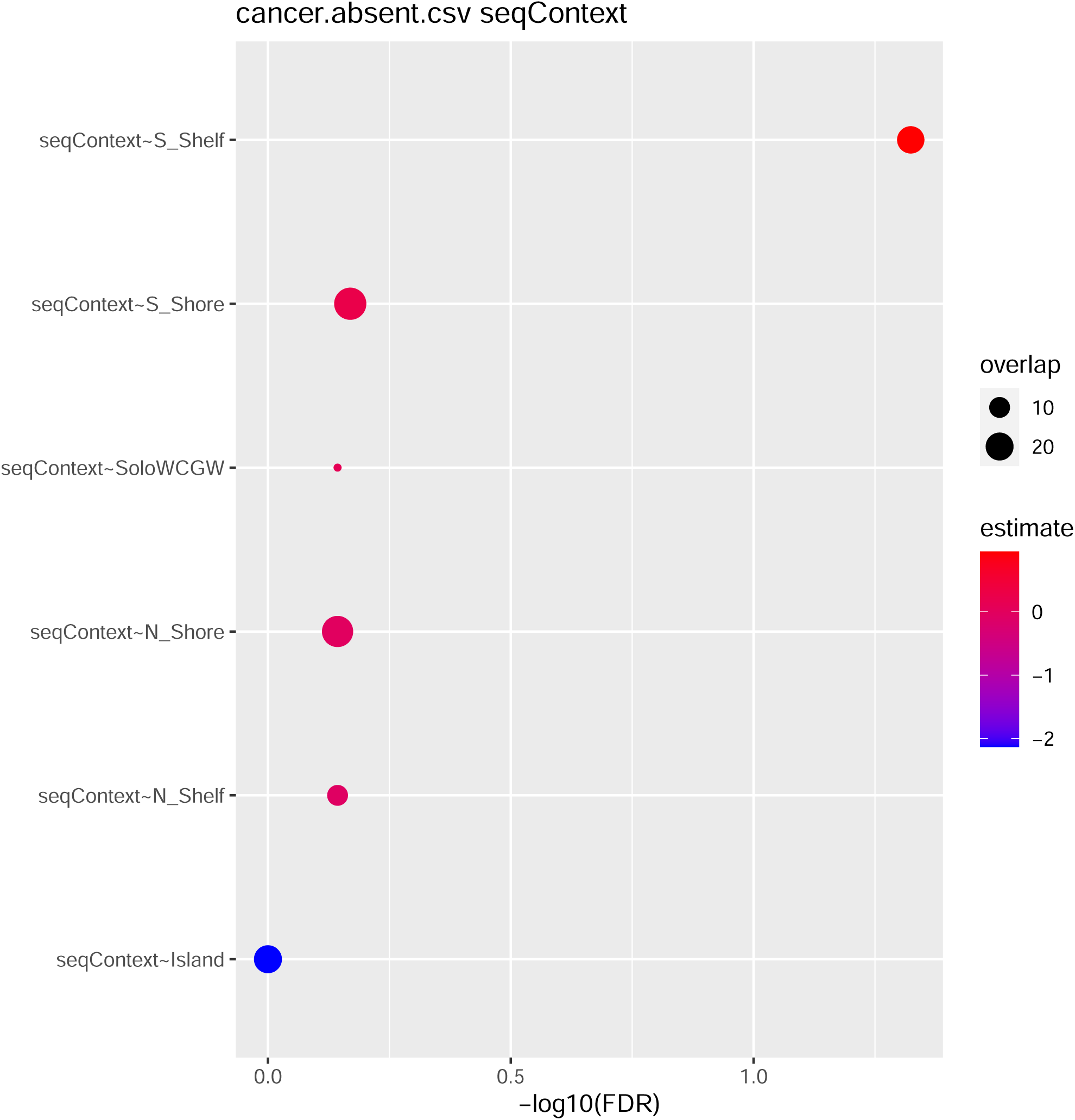
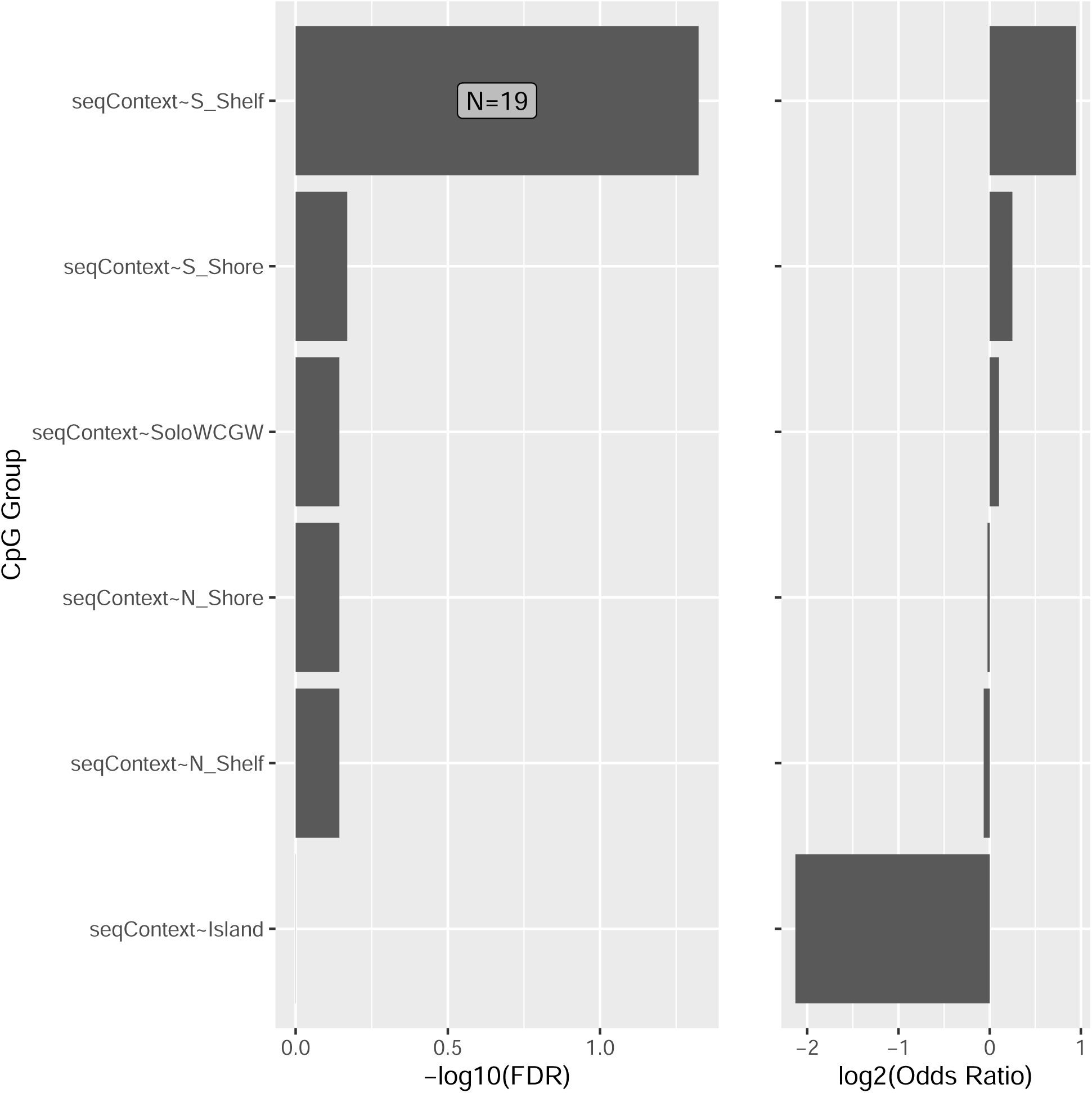
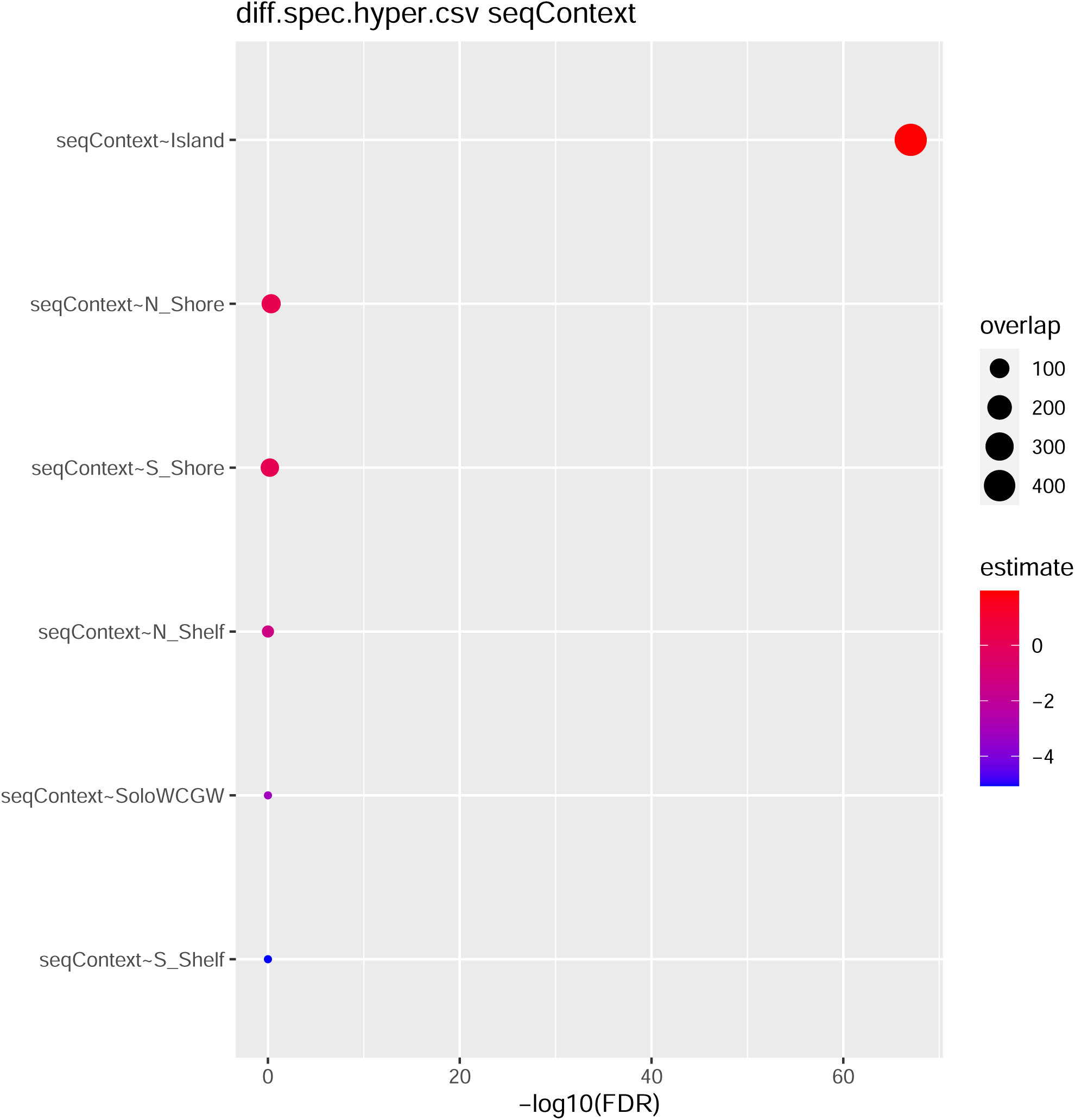
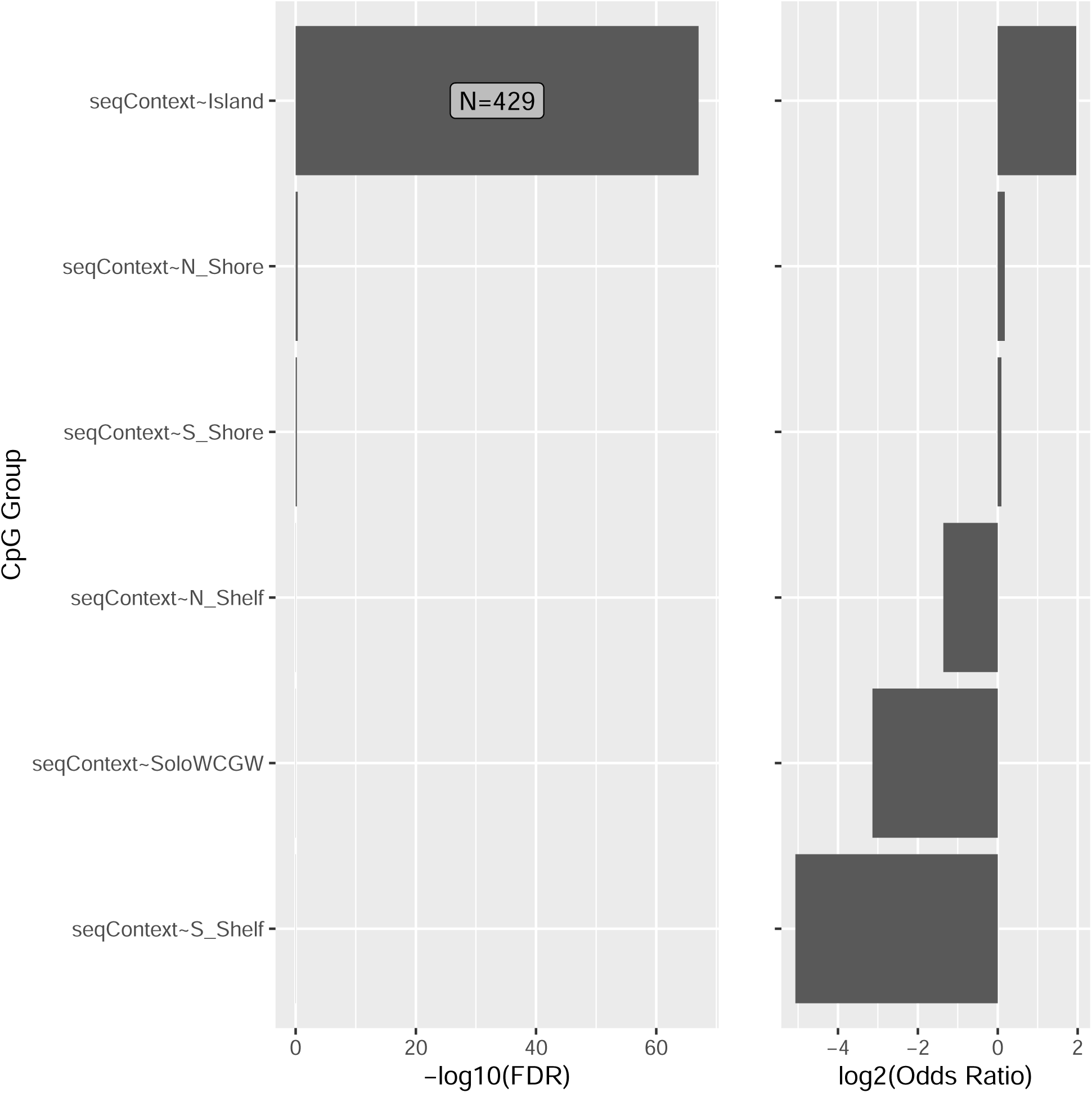
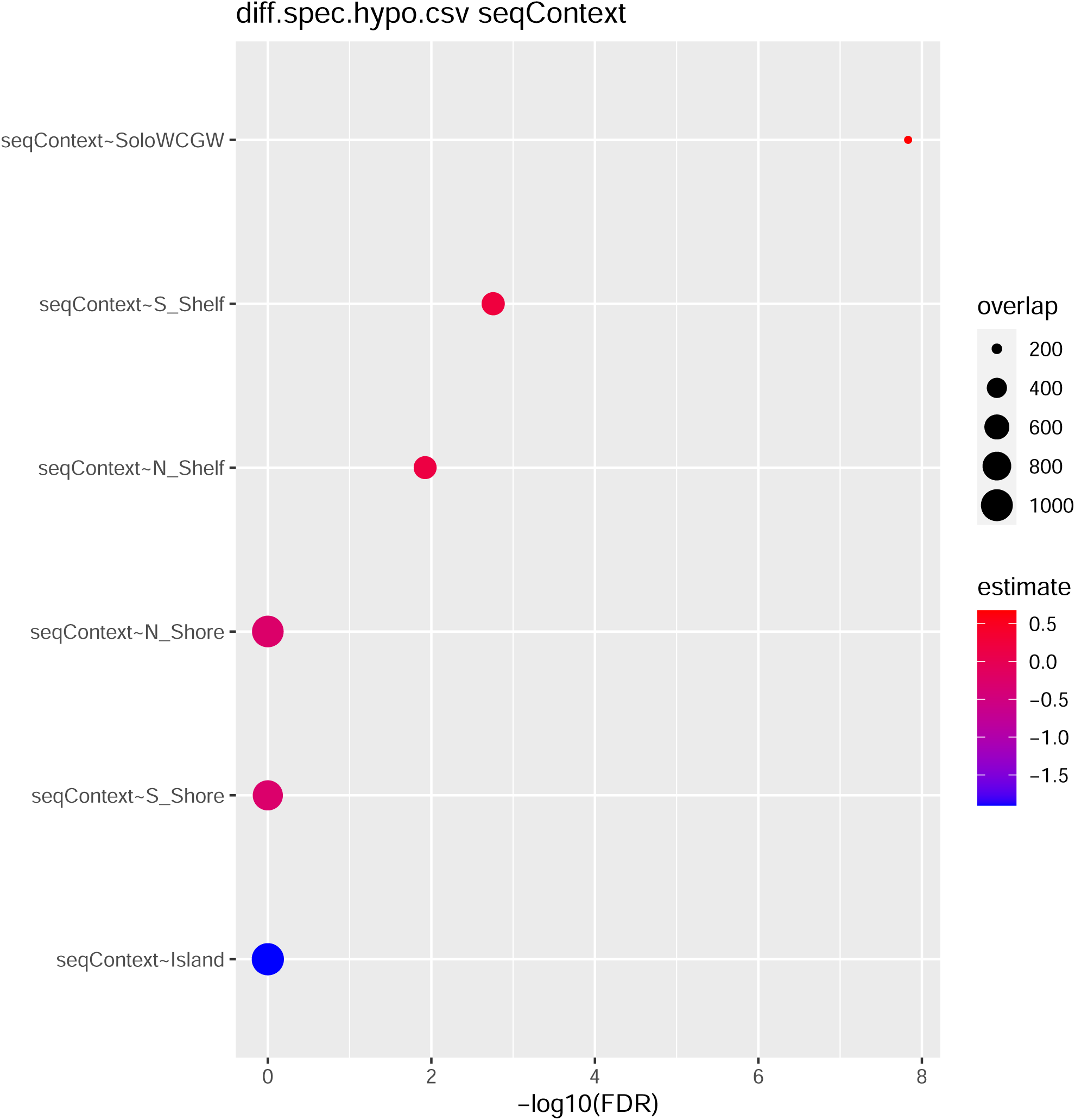
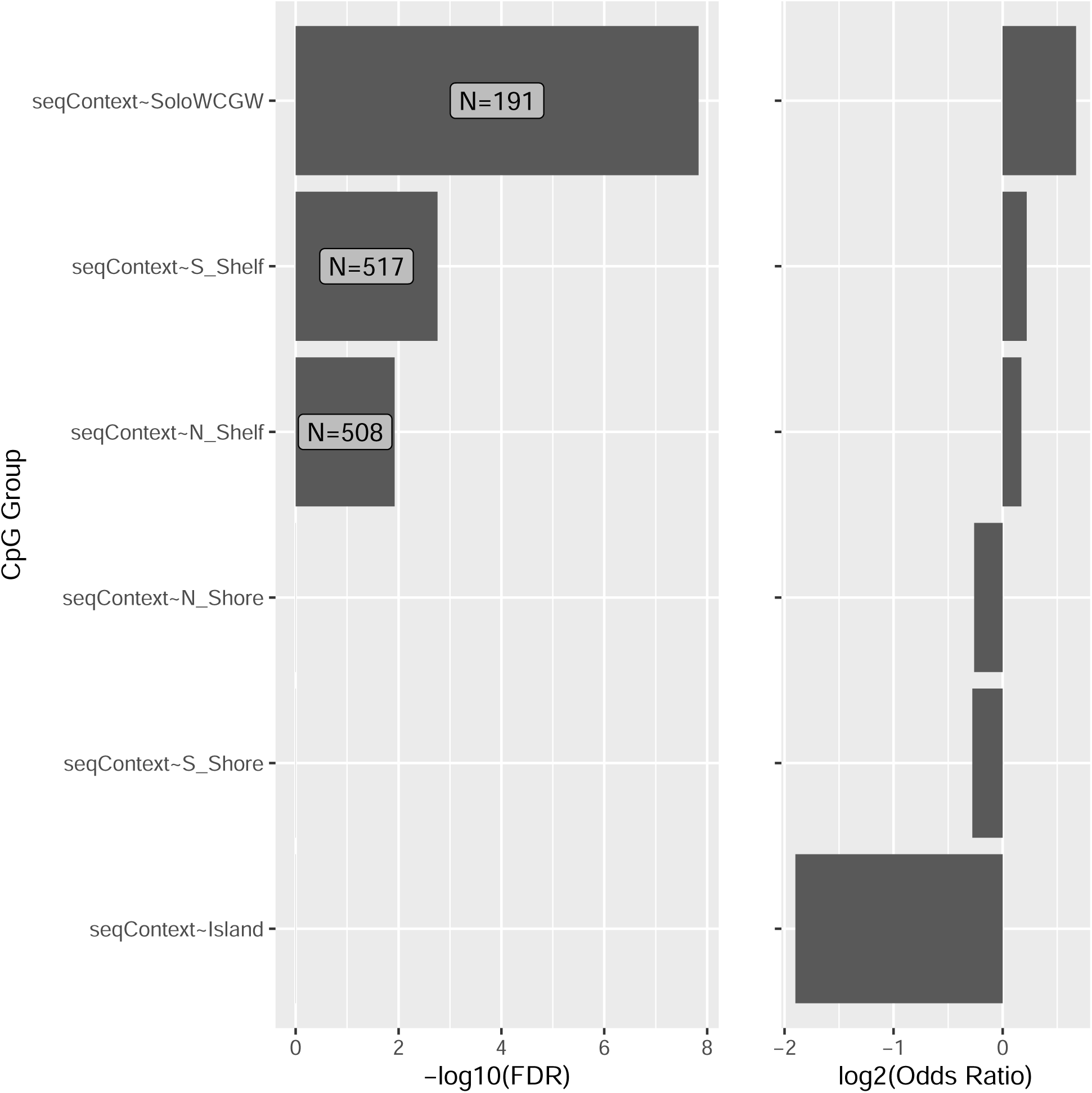
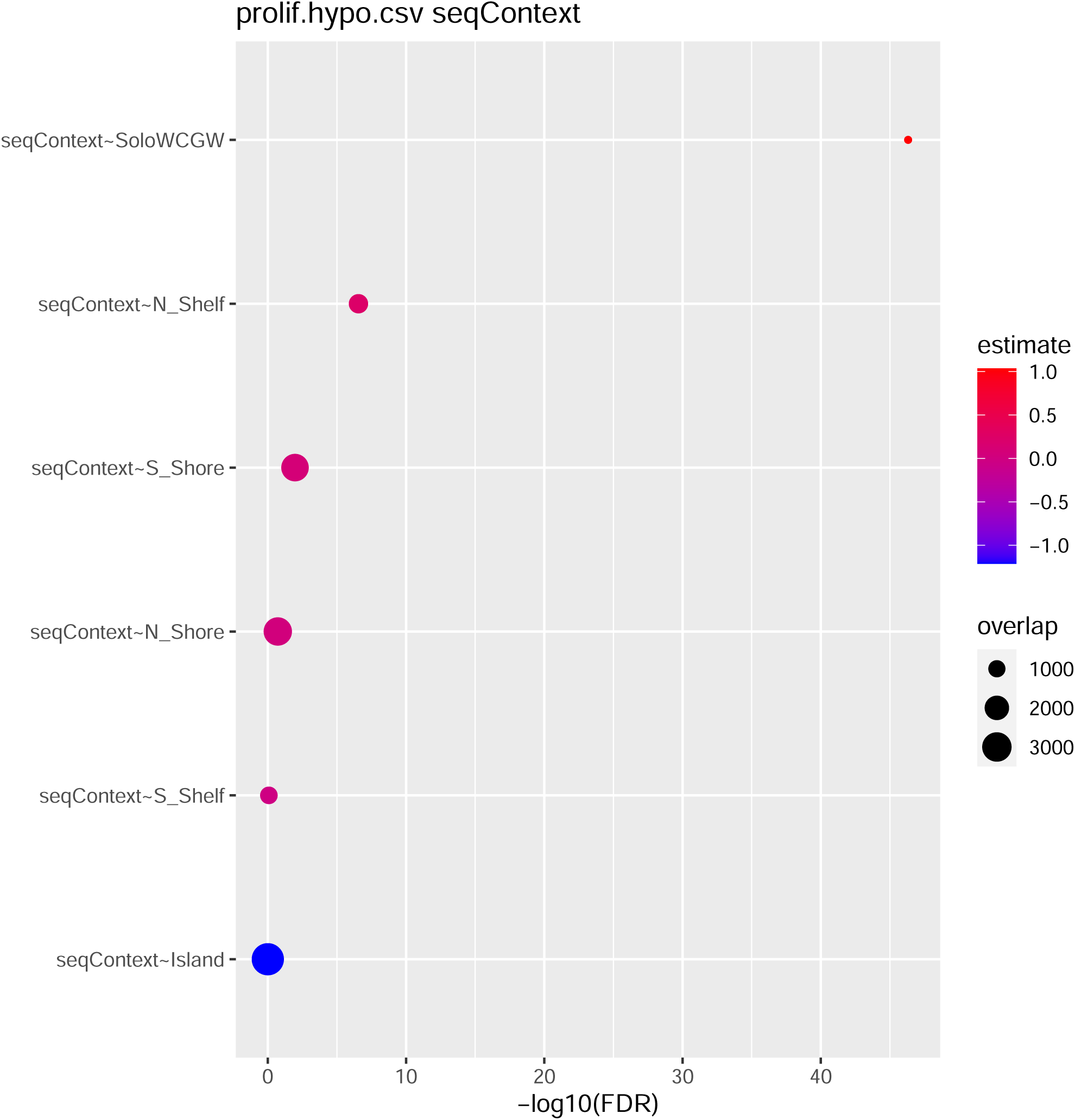
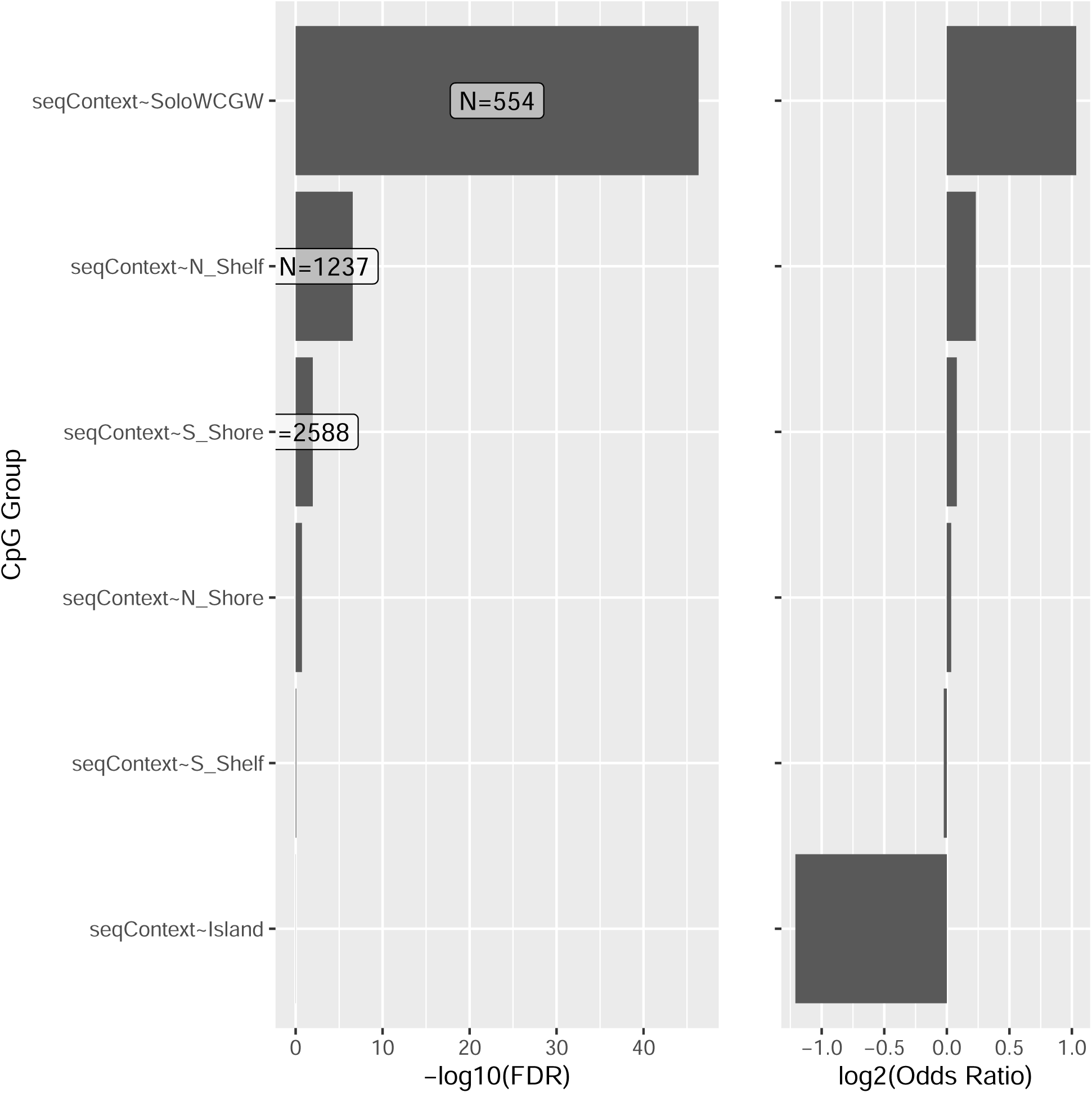
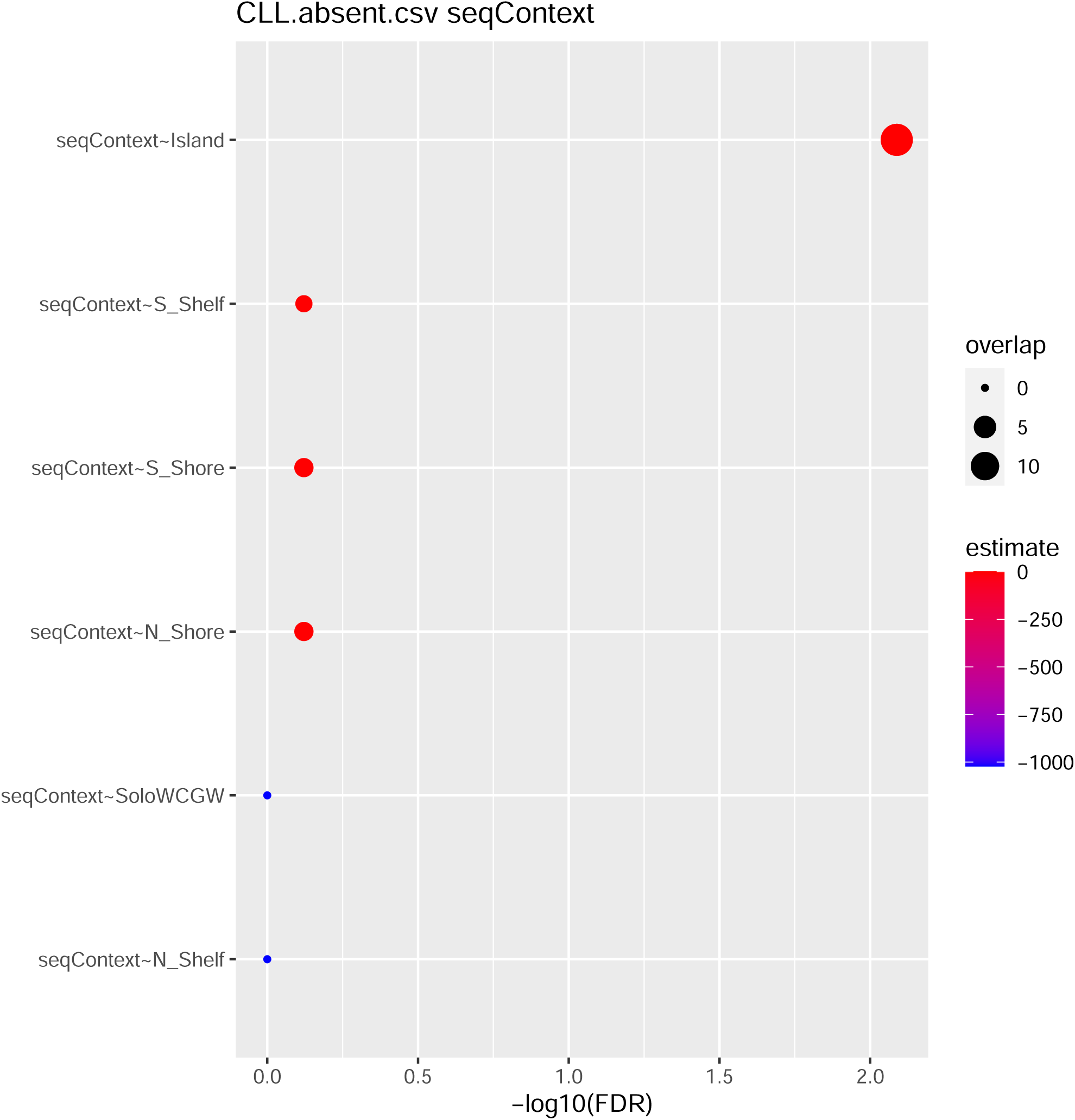
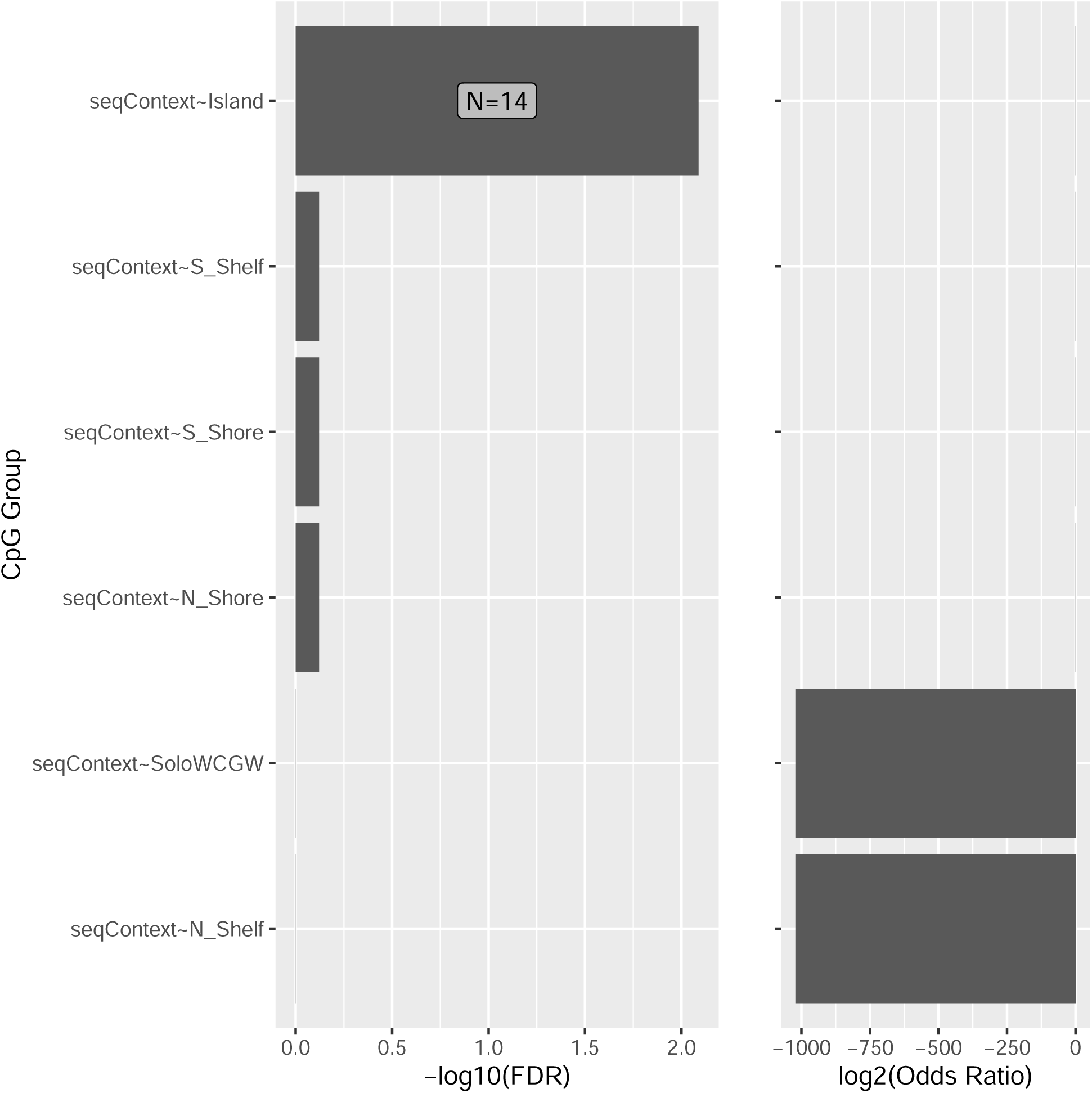
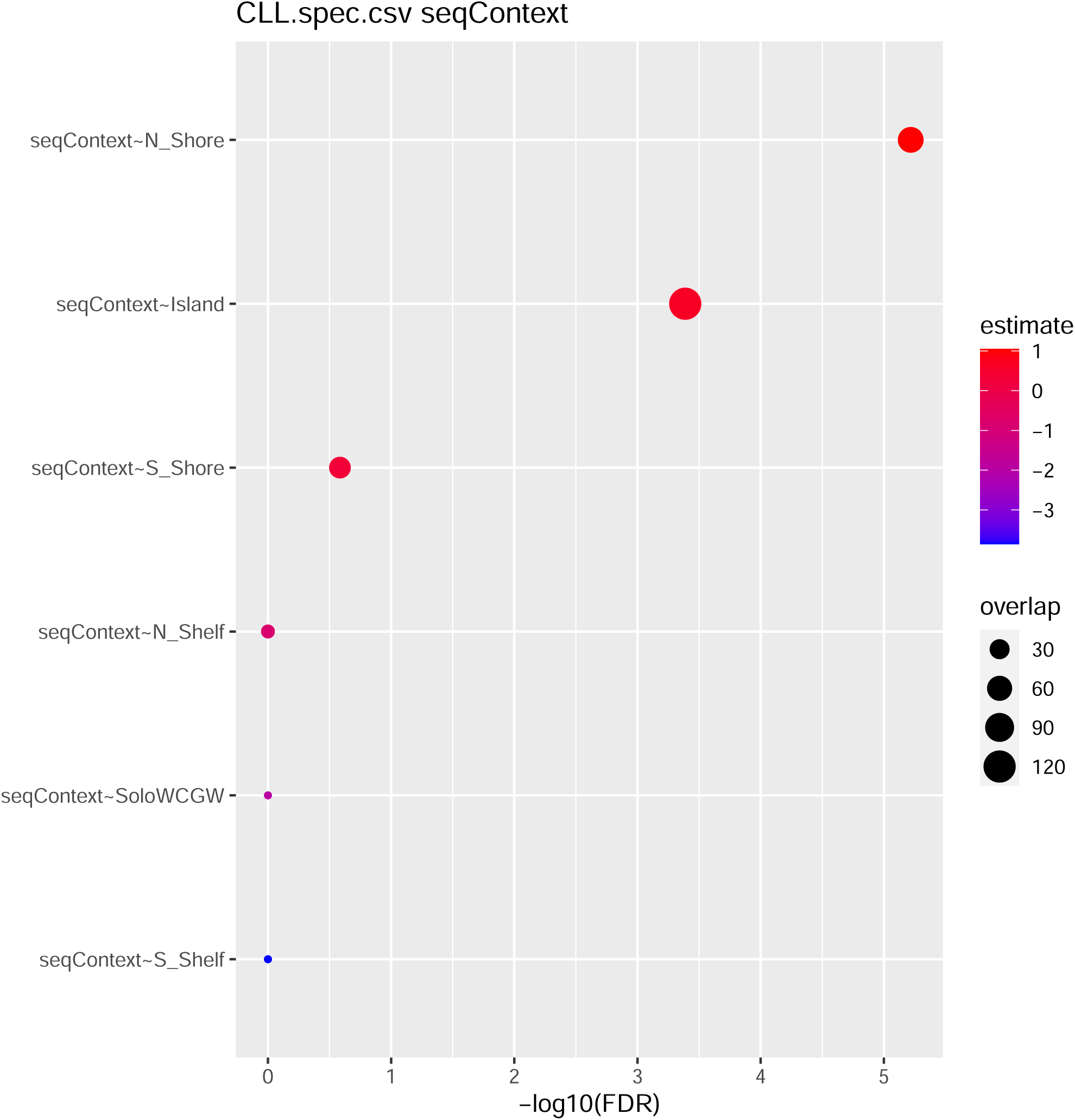
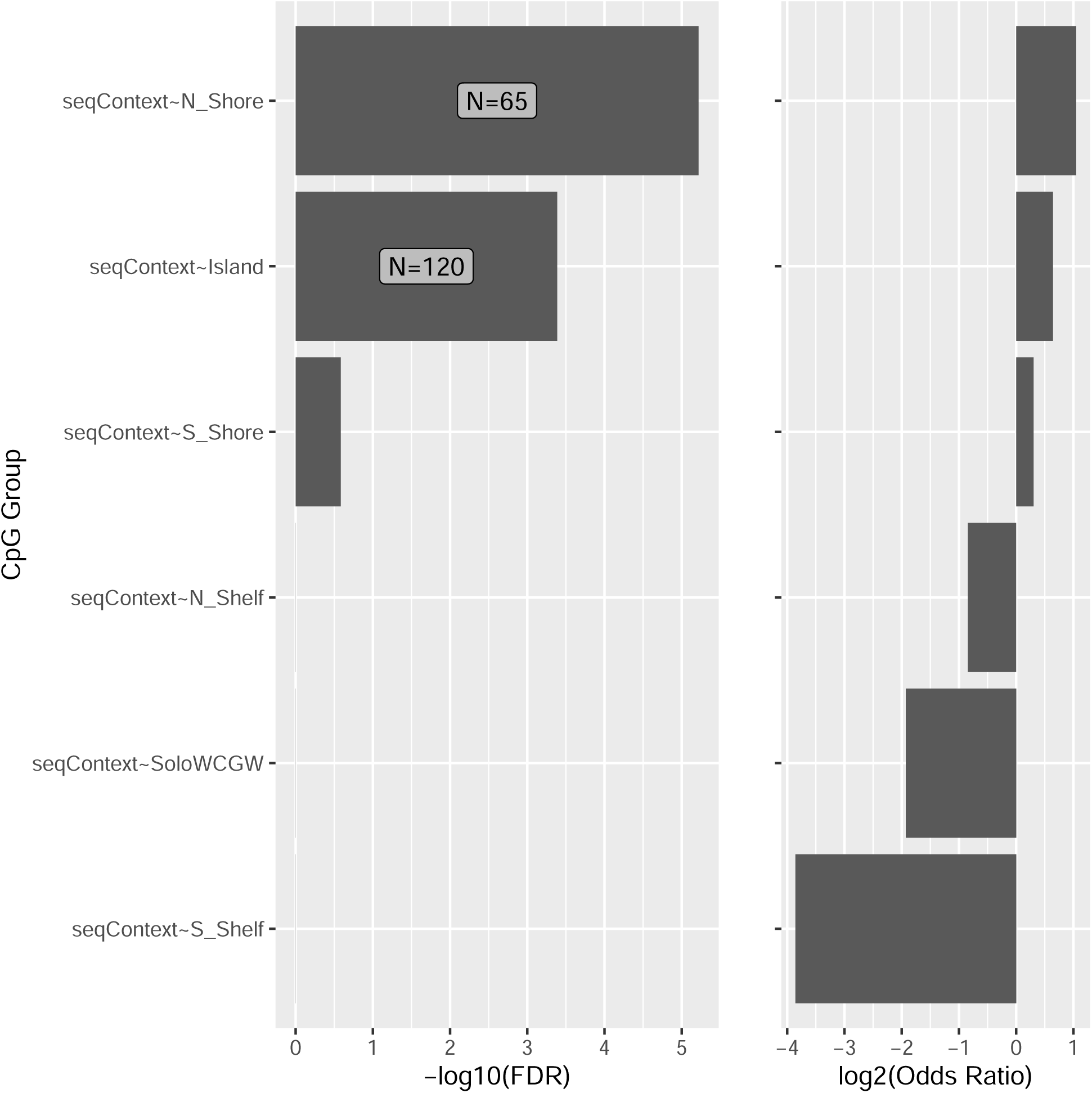
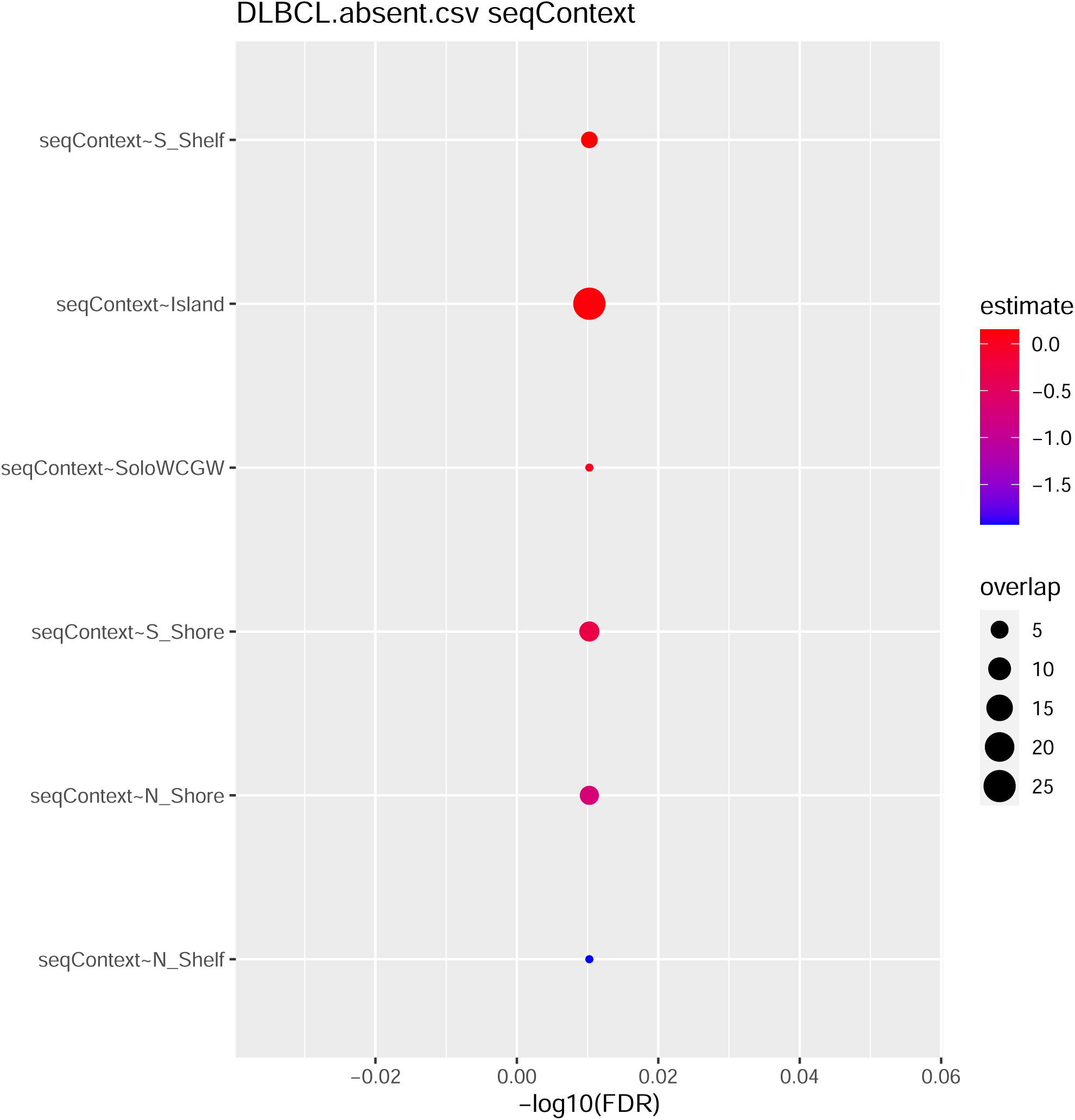
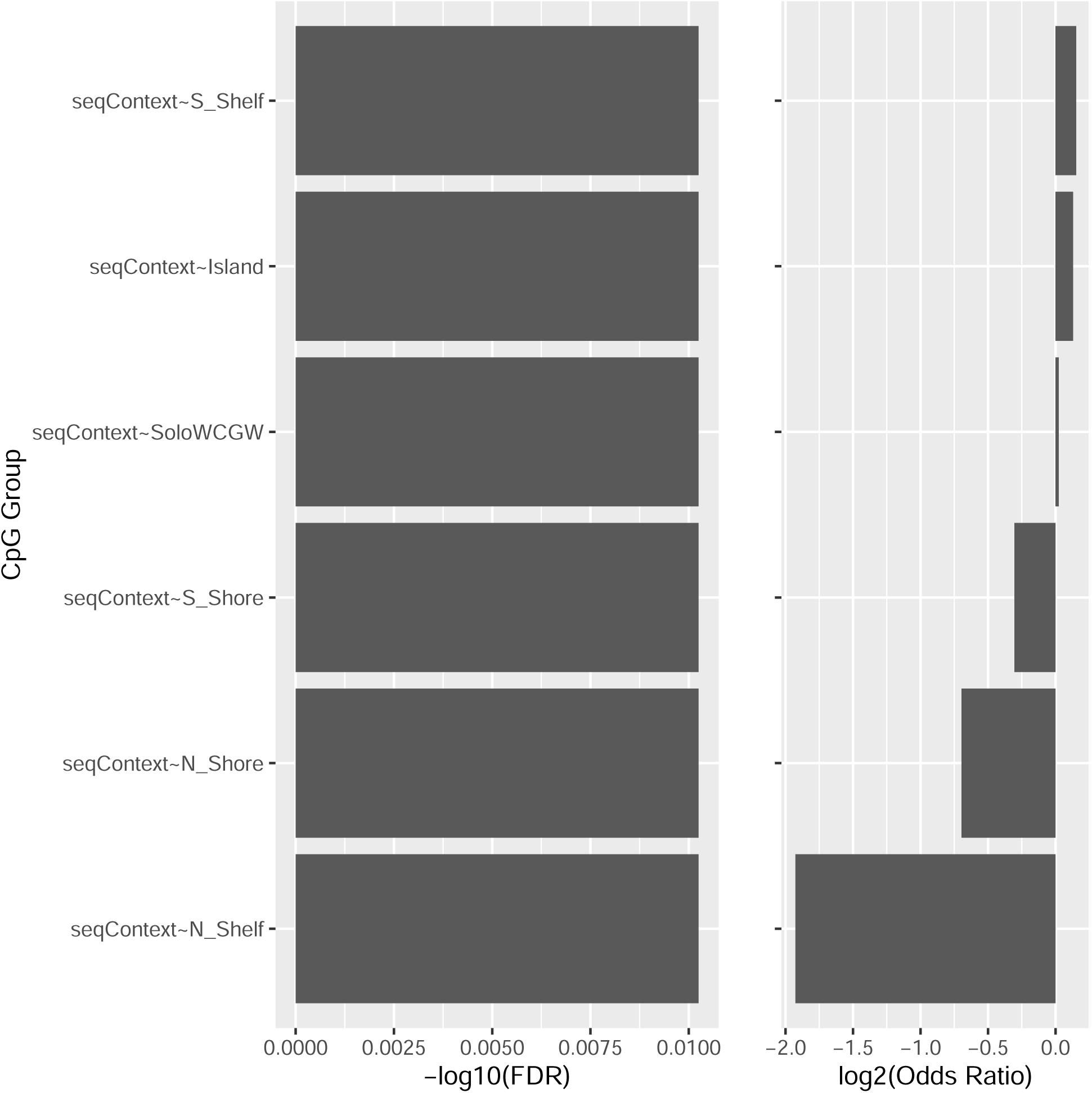
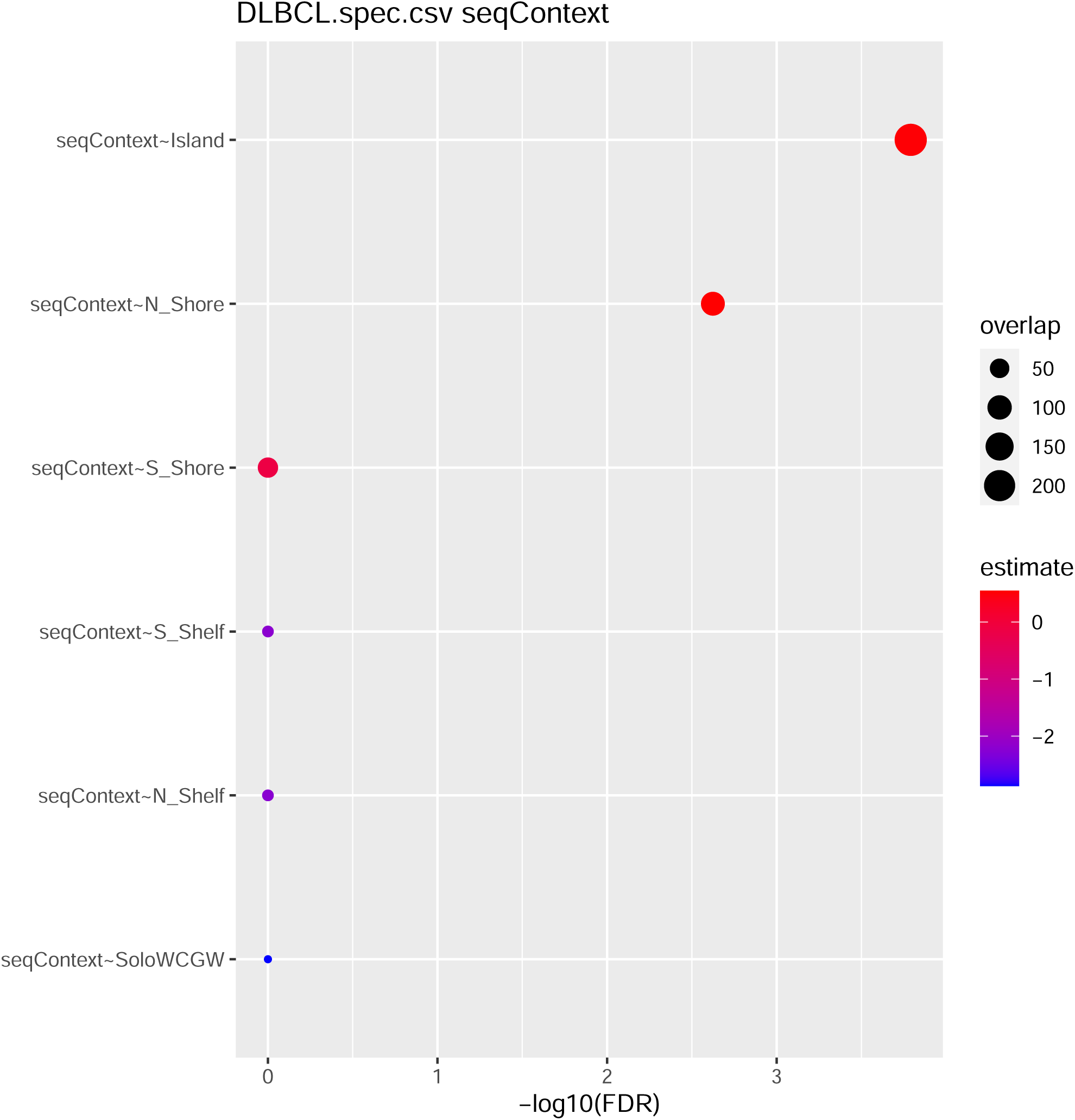
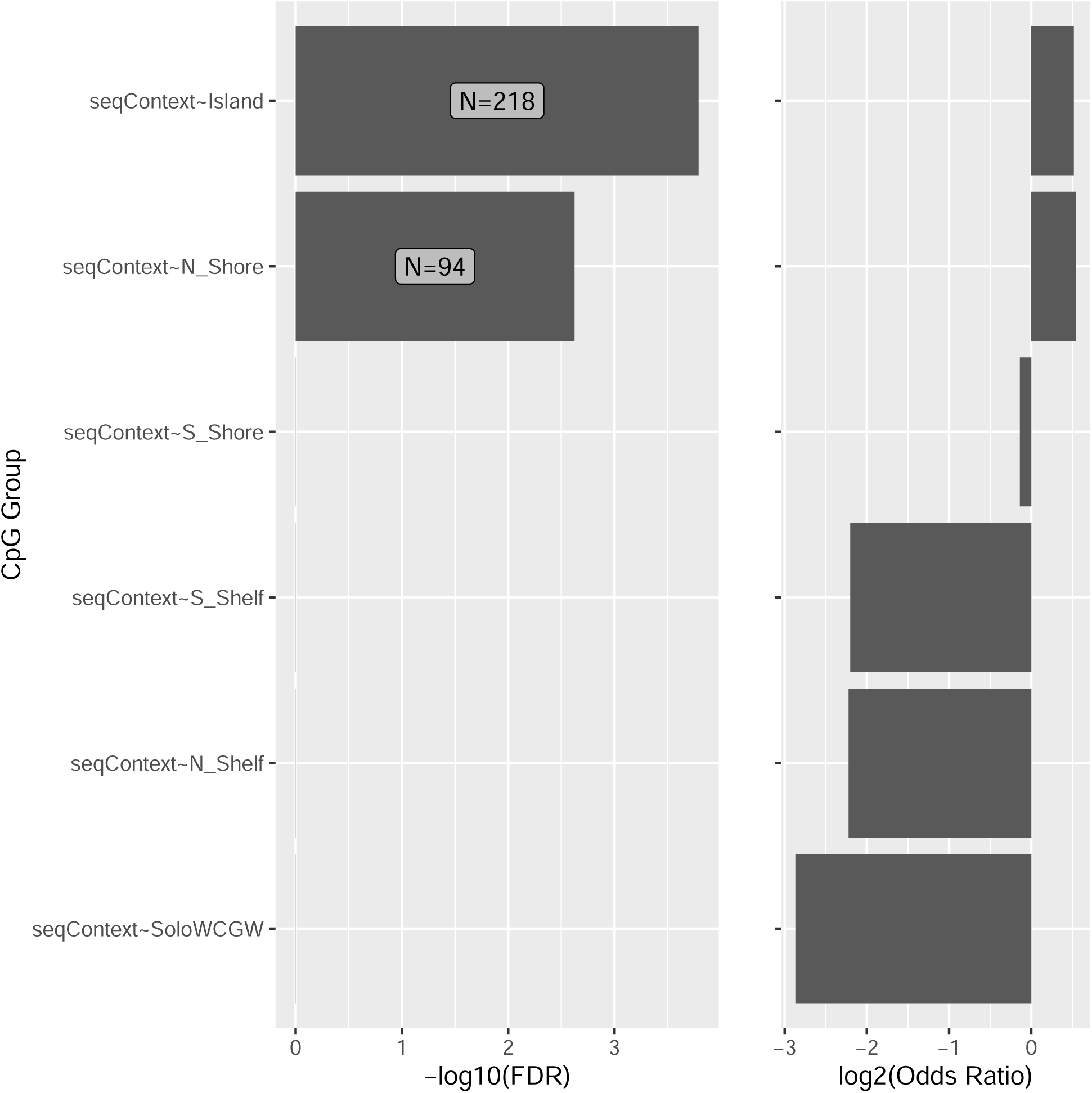
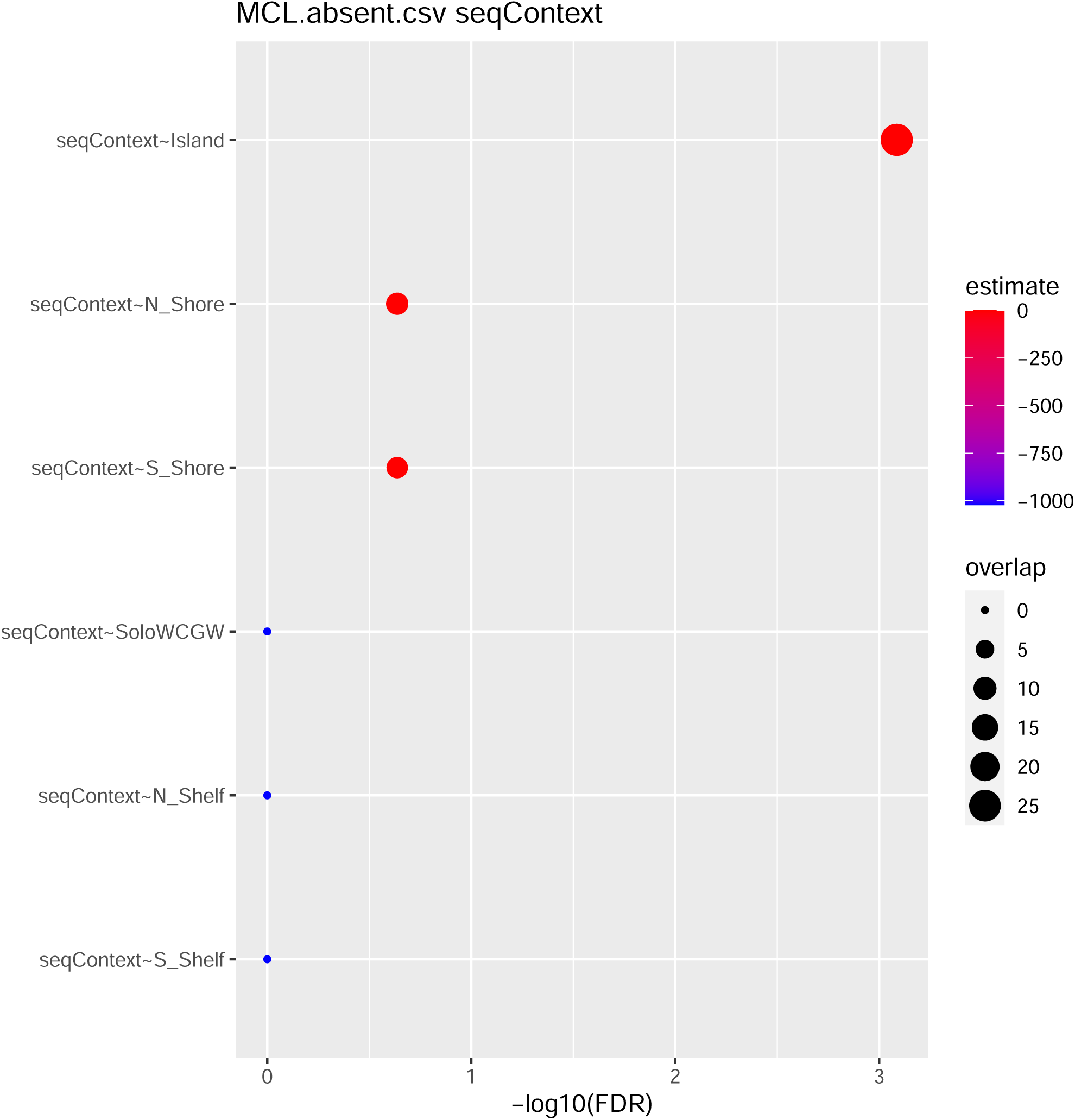
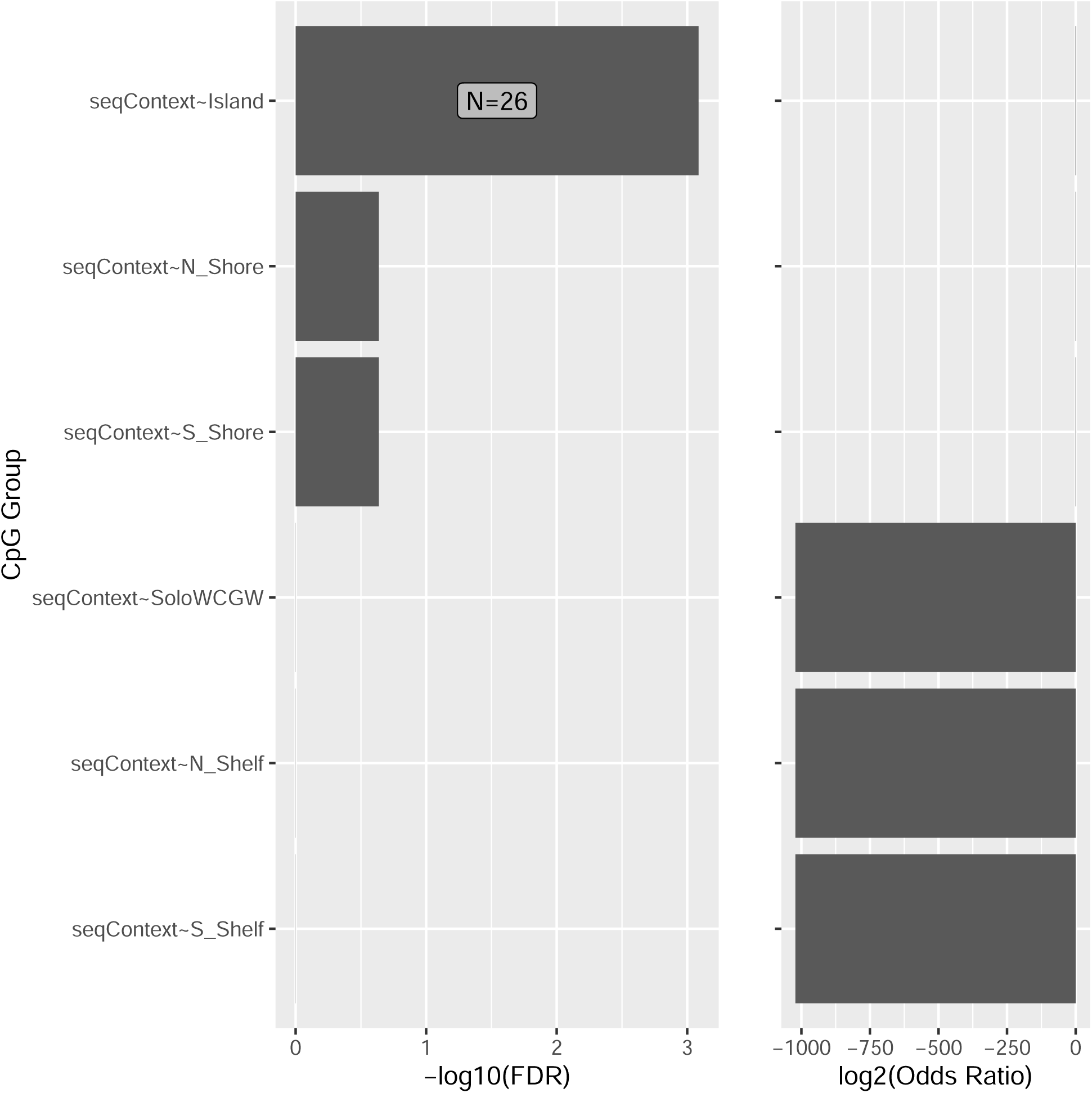
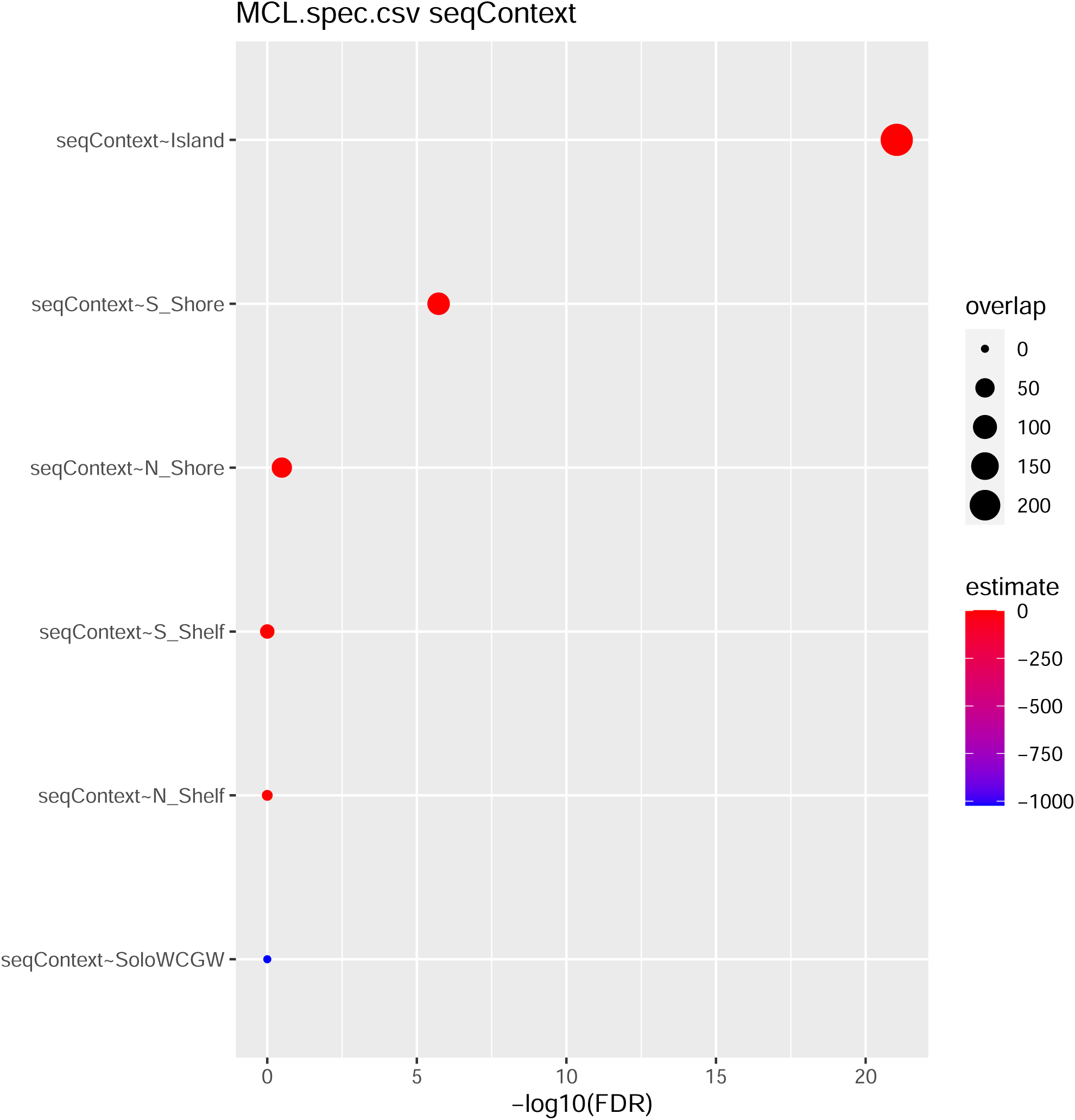
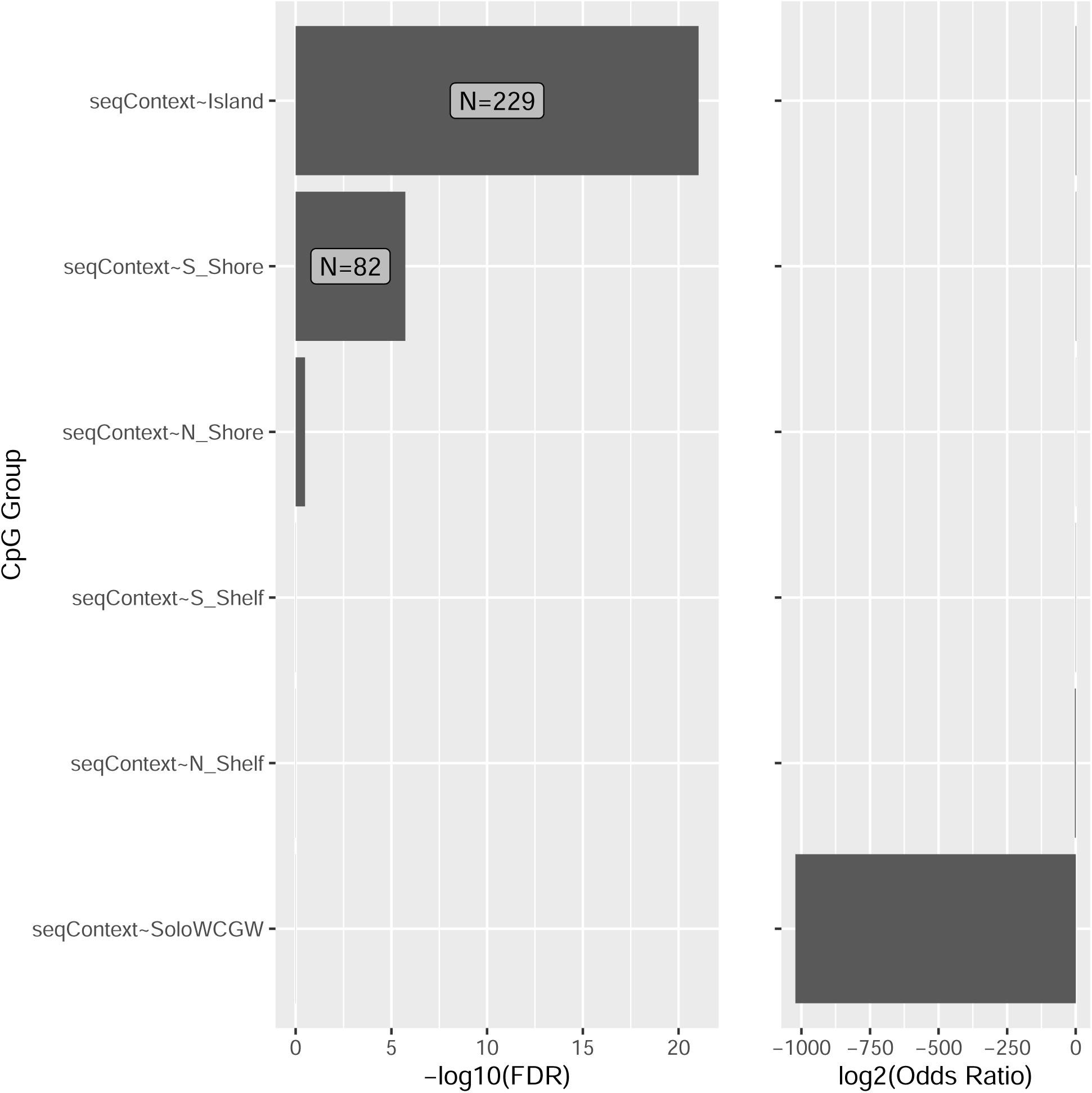
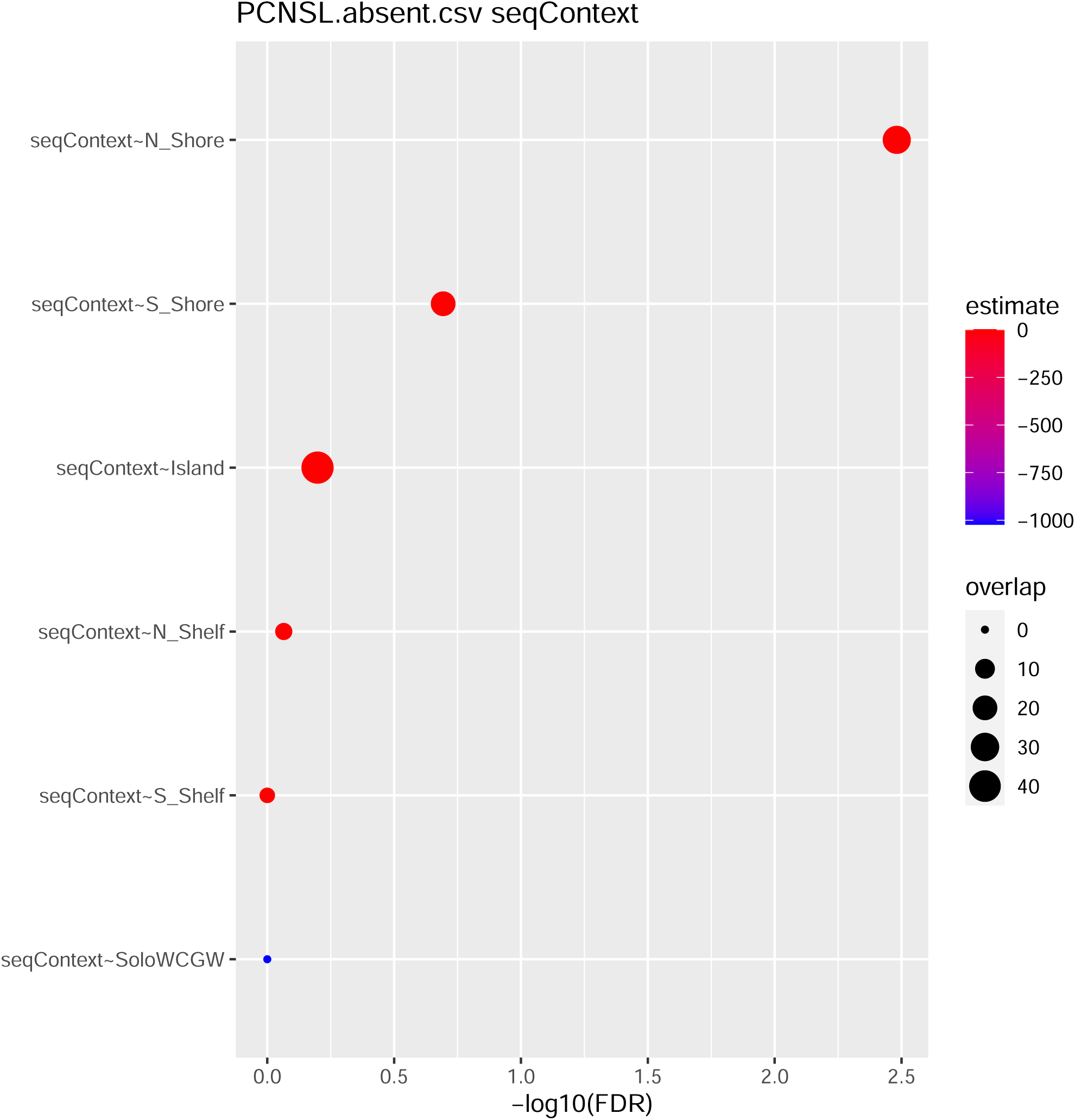
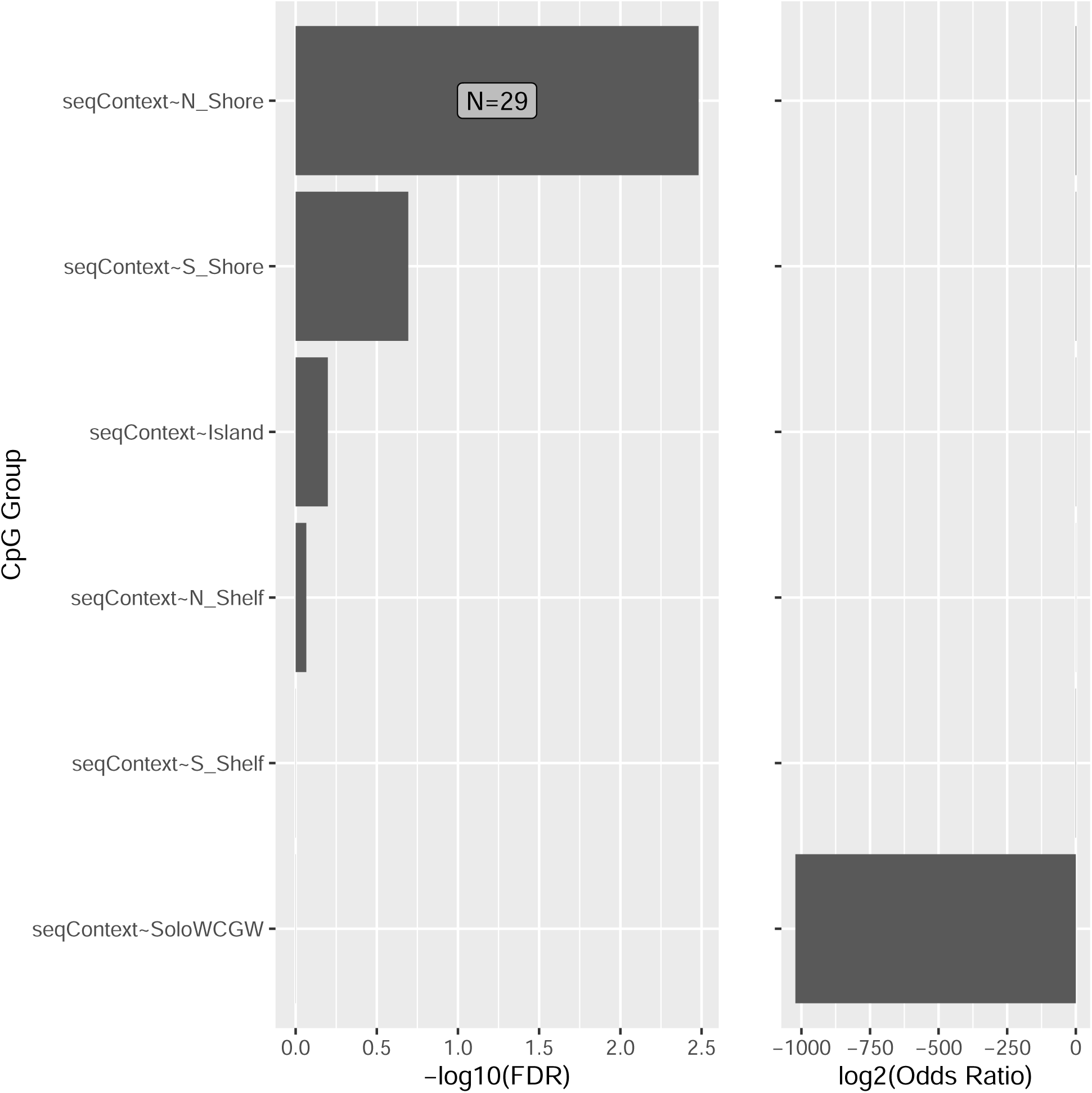
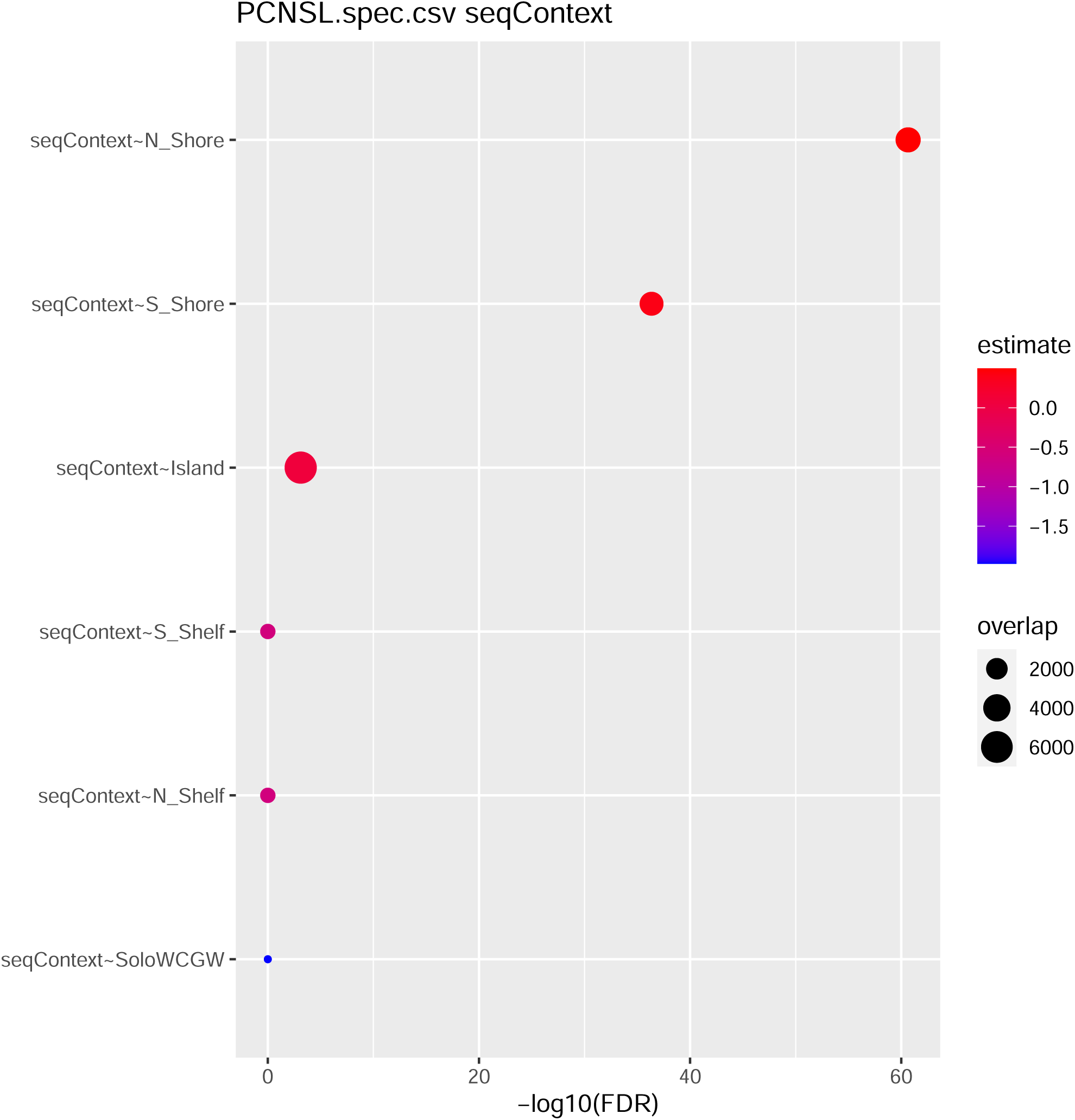
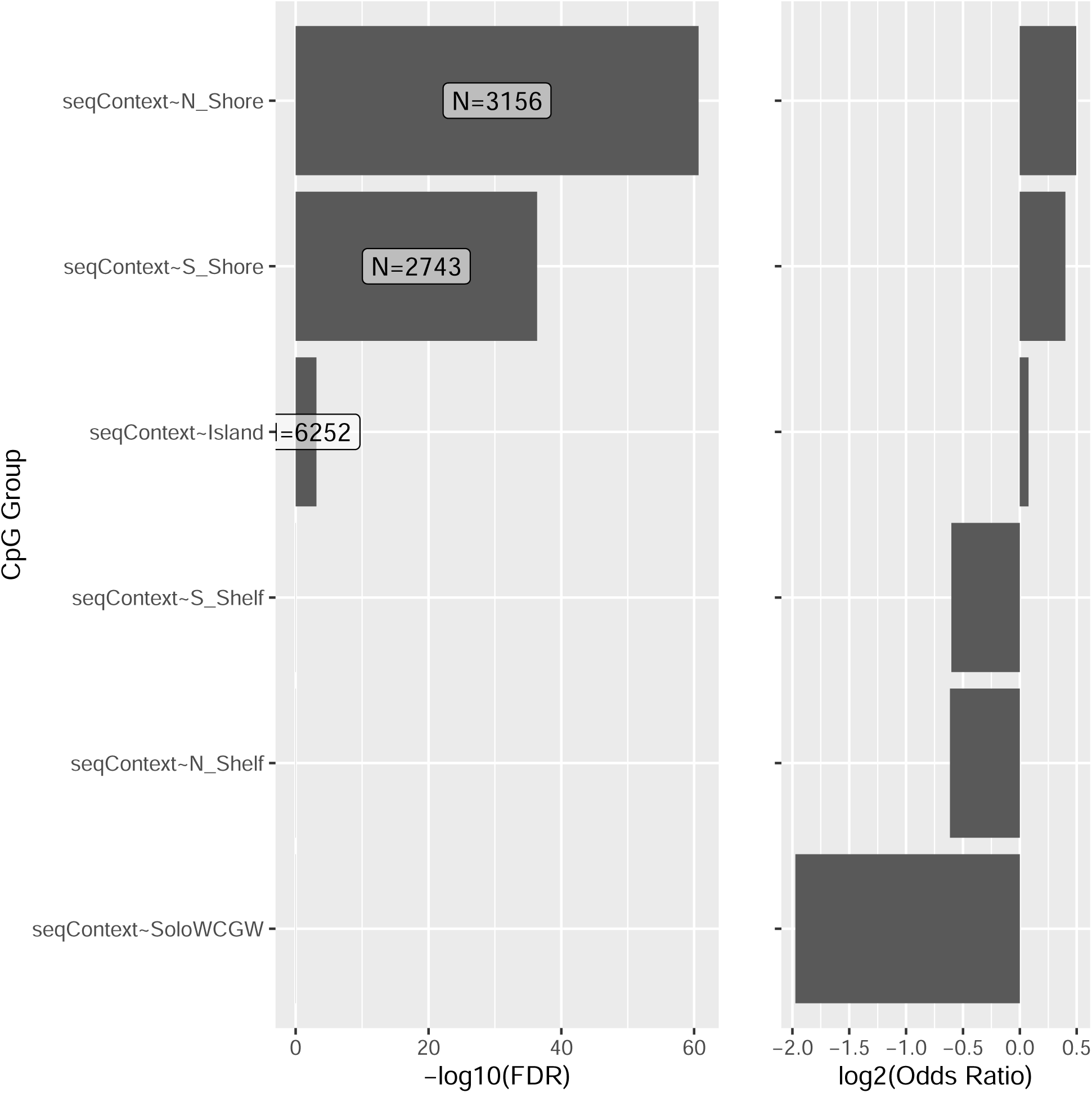
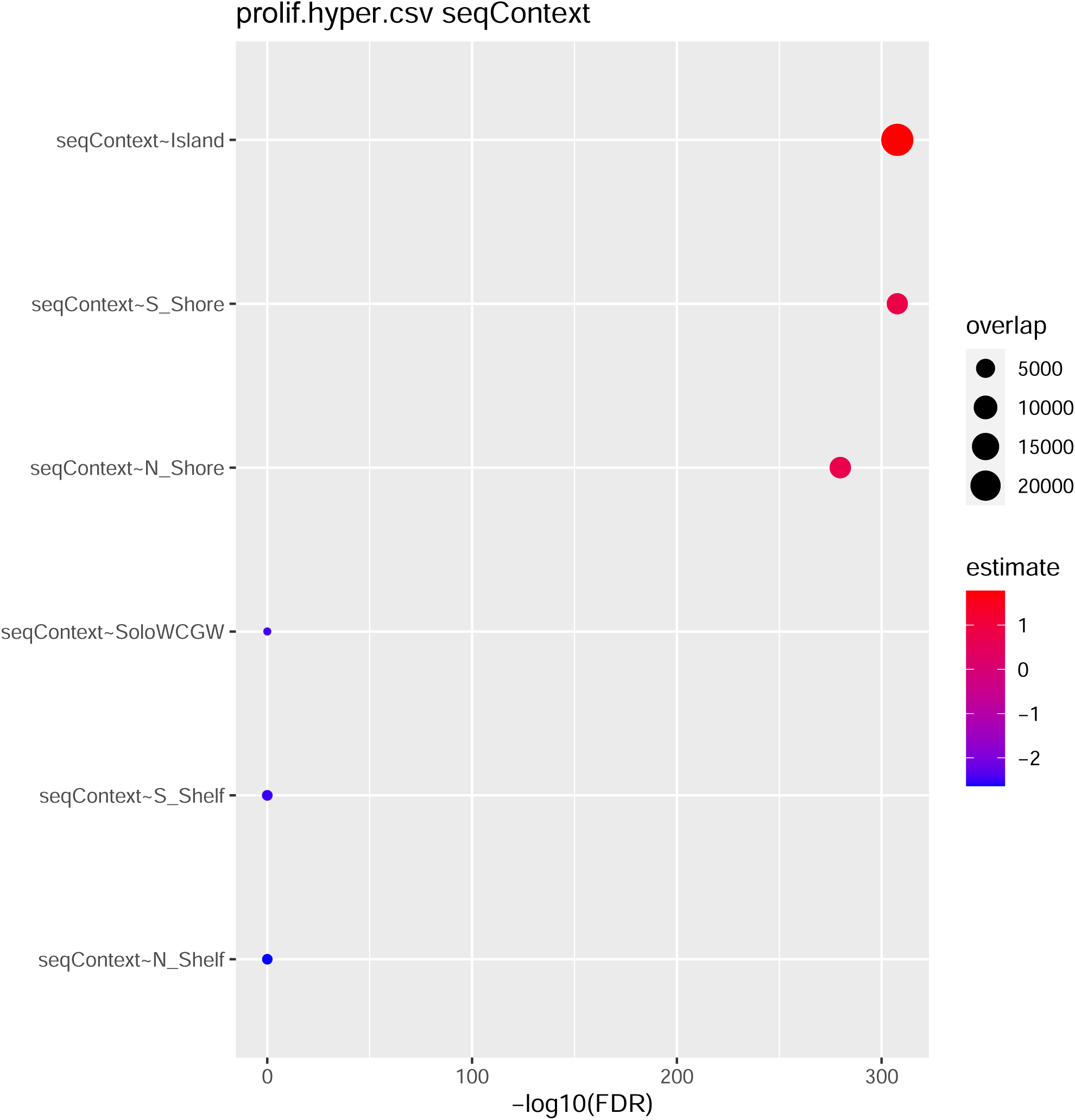
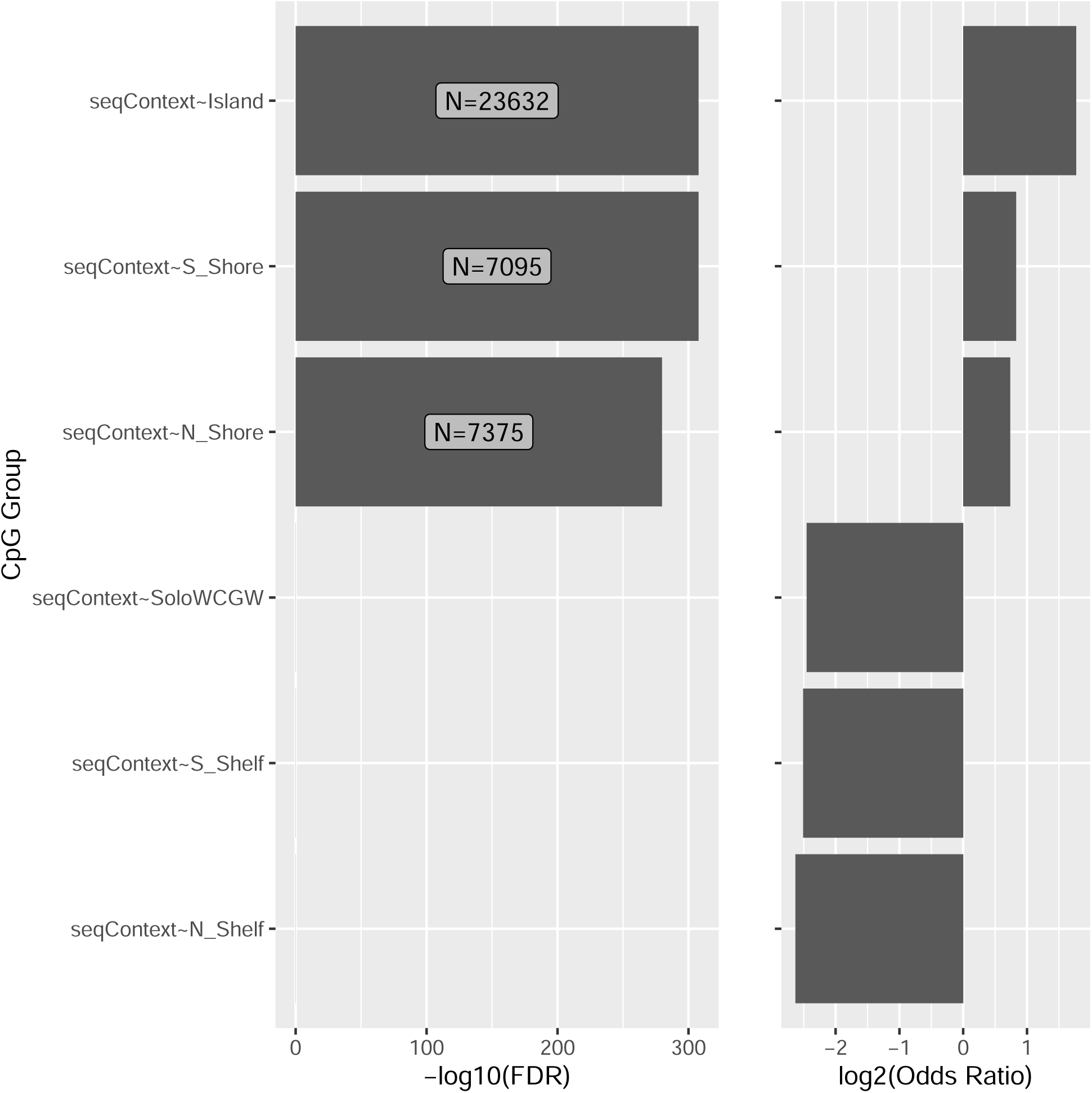

**Supplementary Figure 3.**
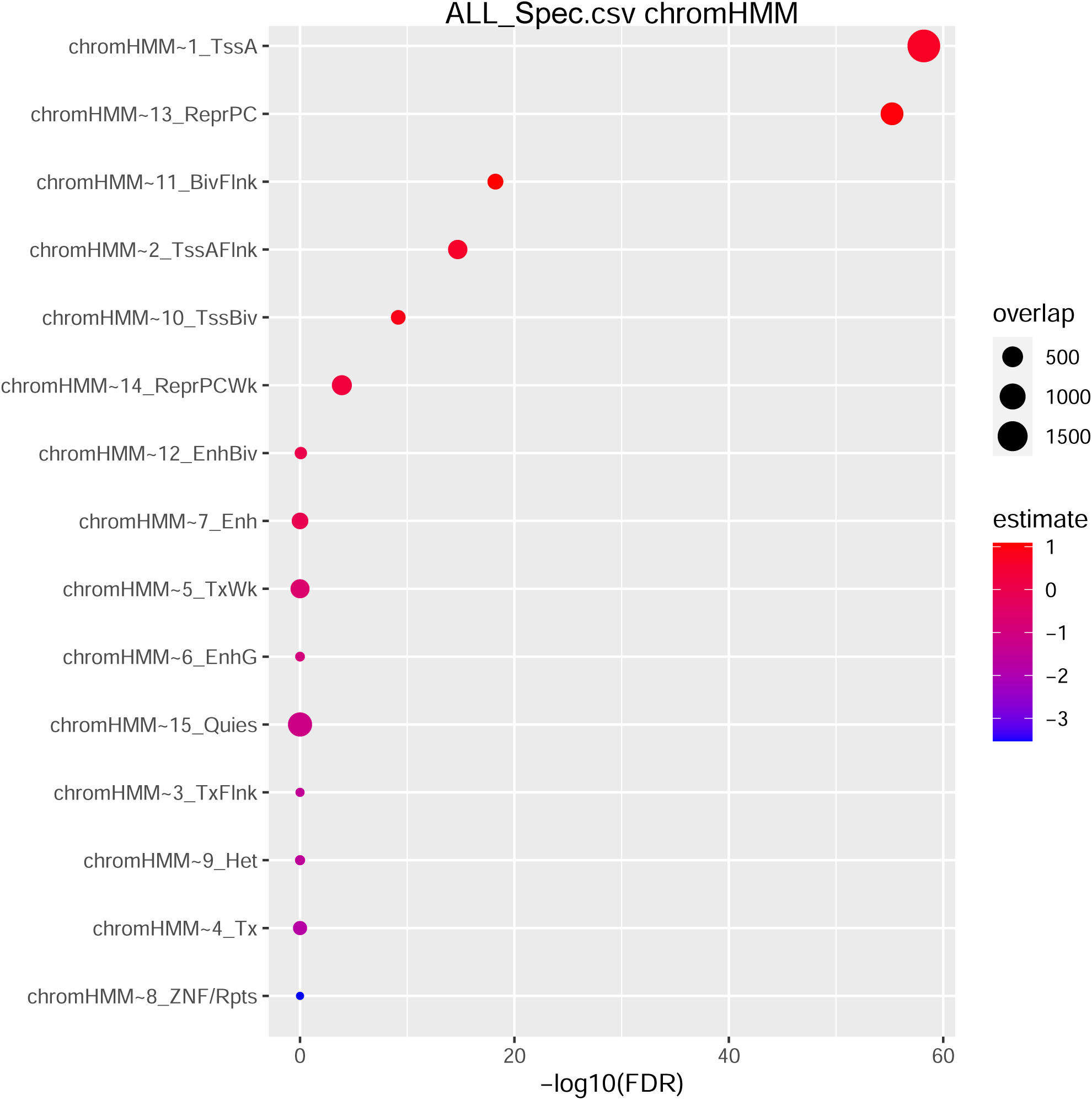
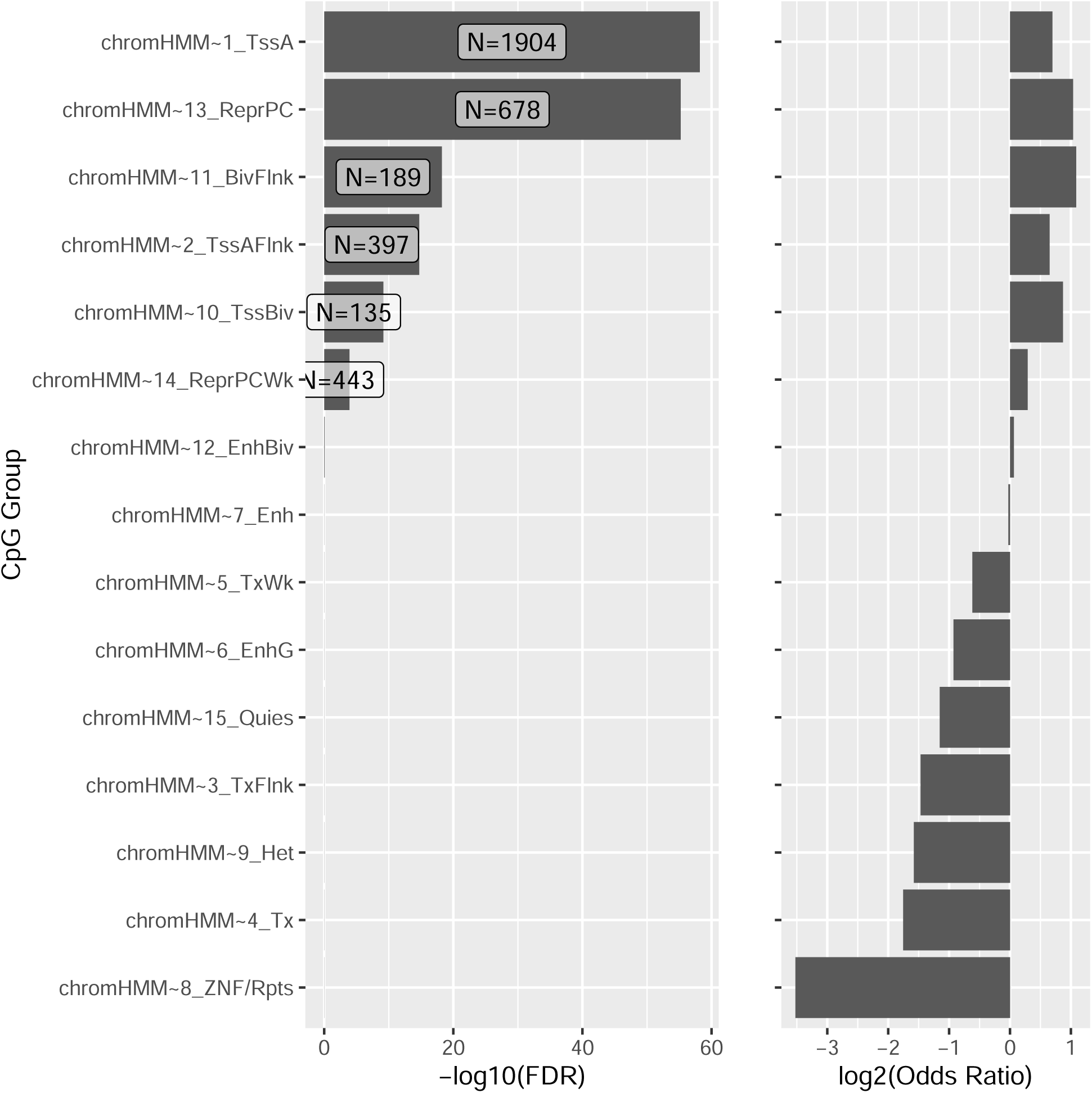
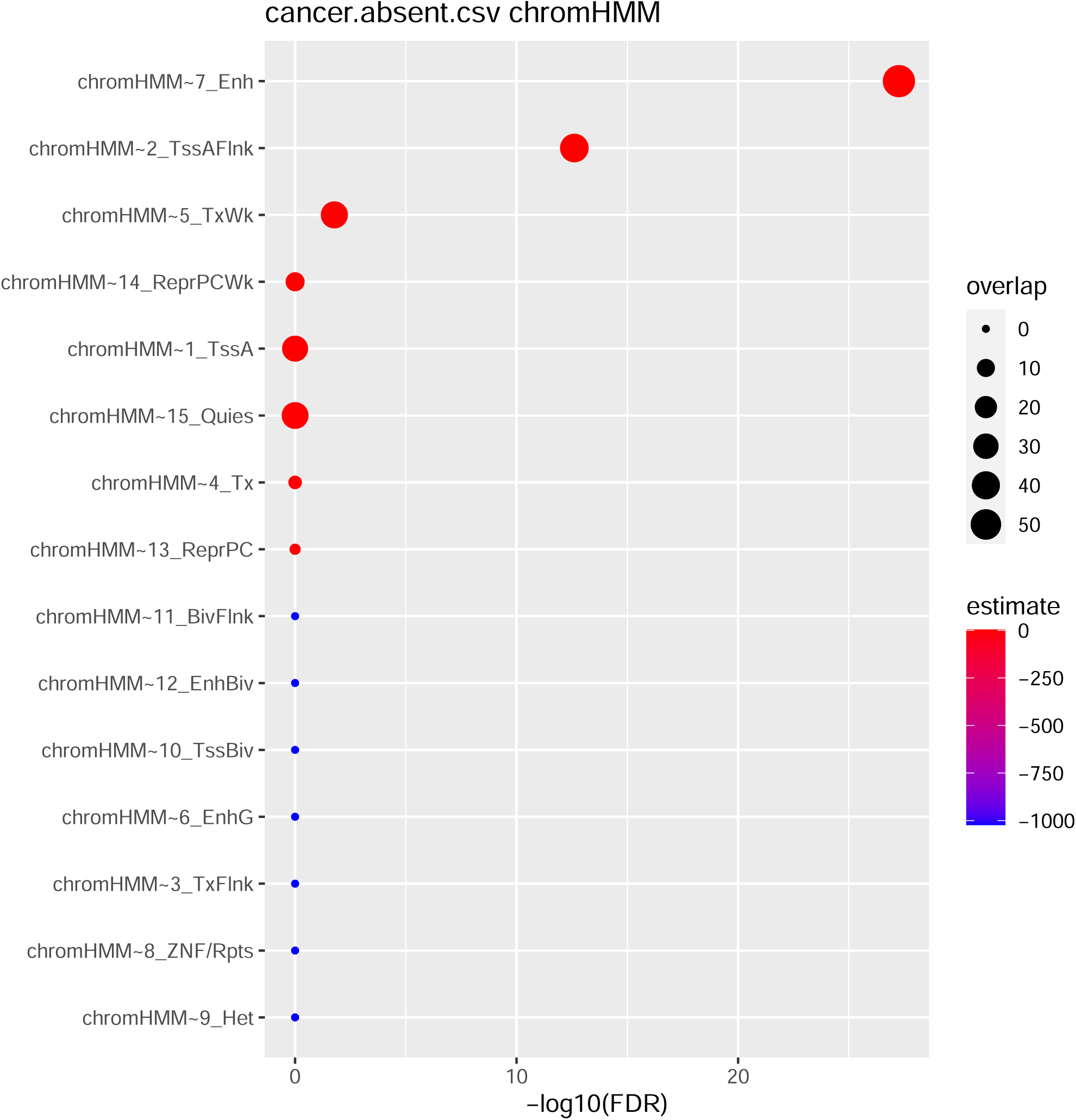
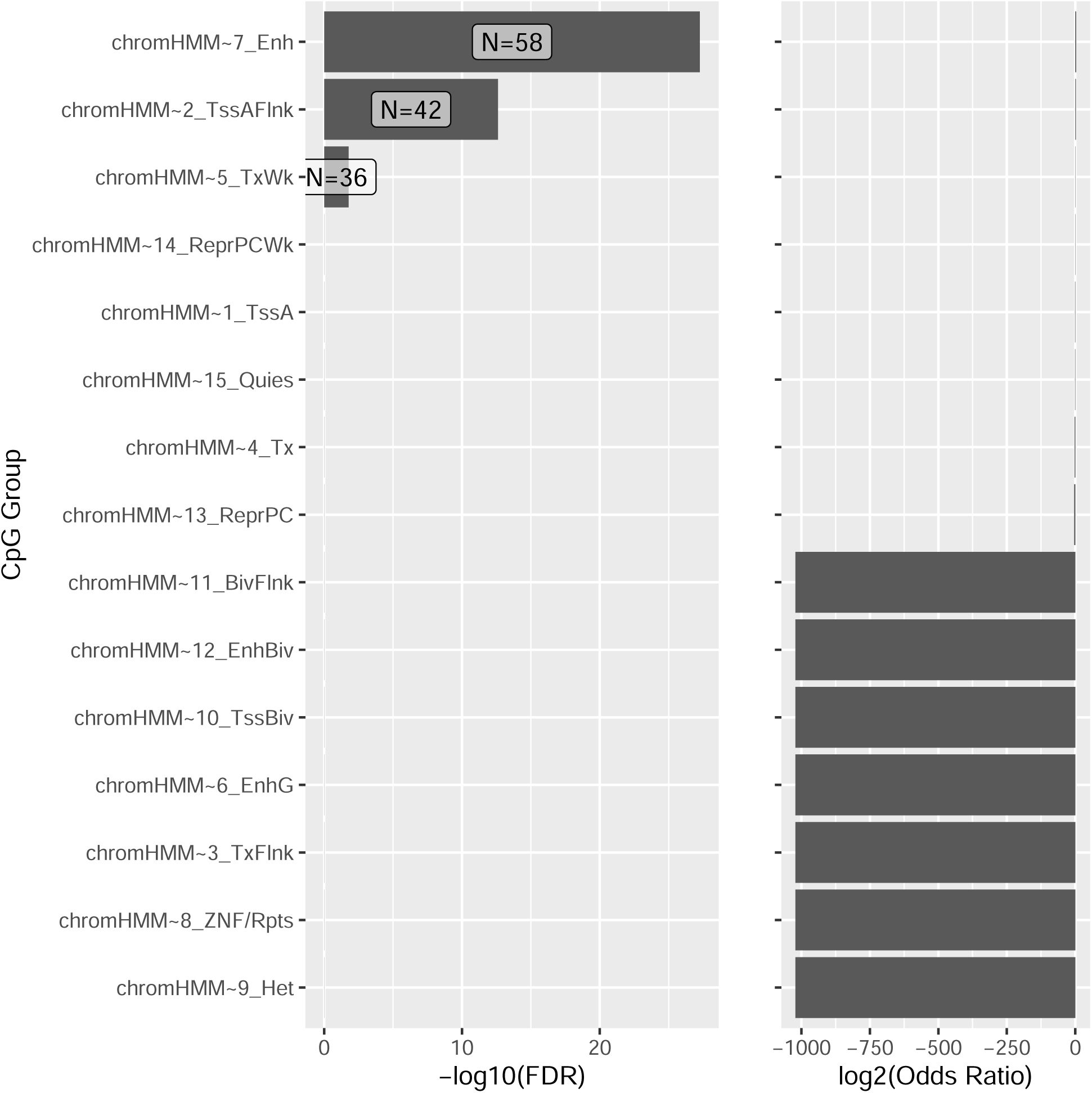
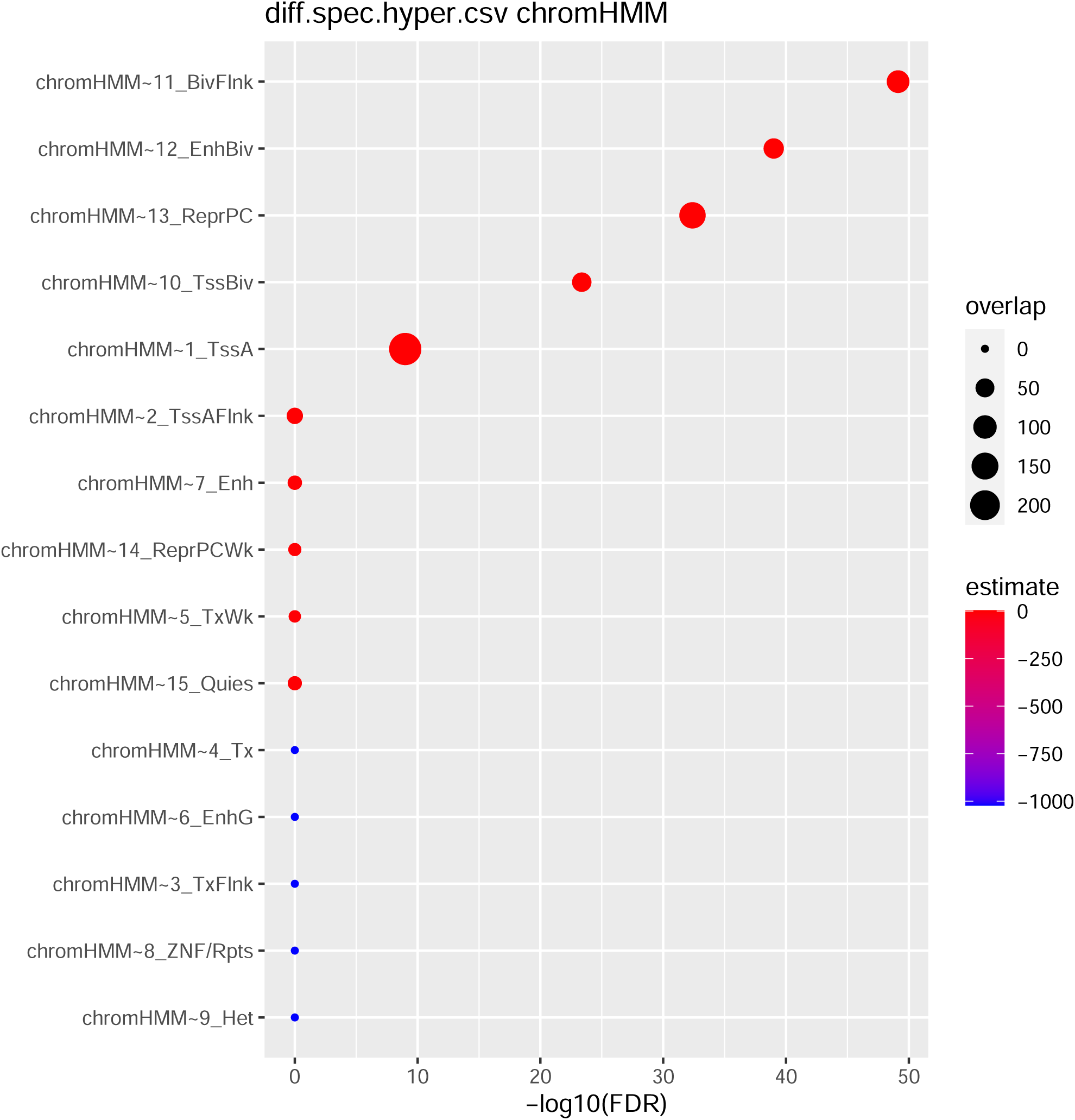
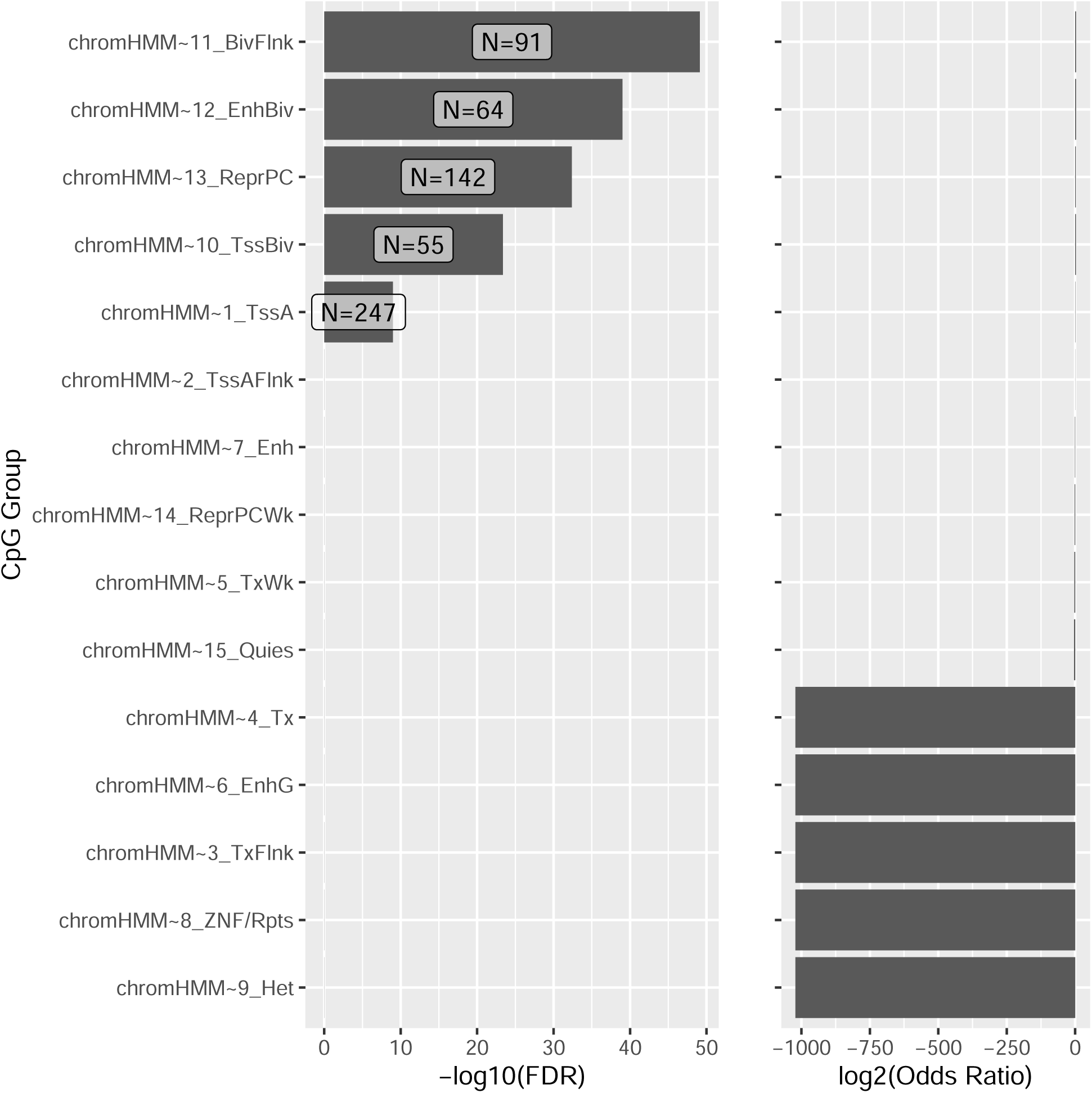
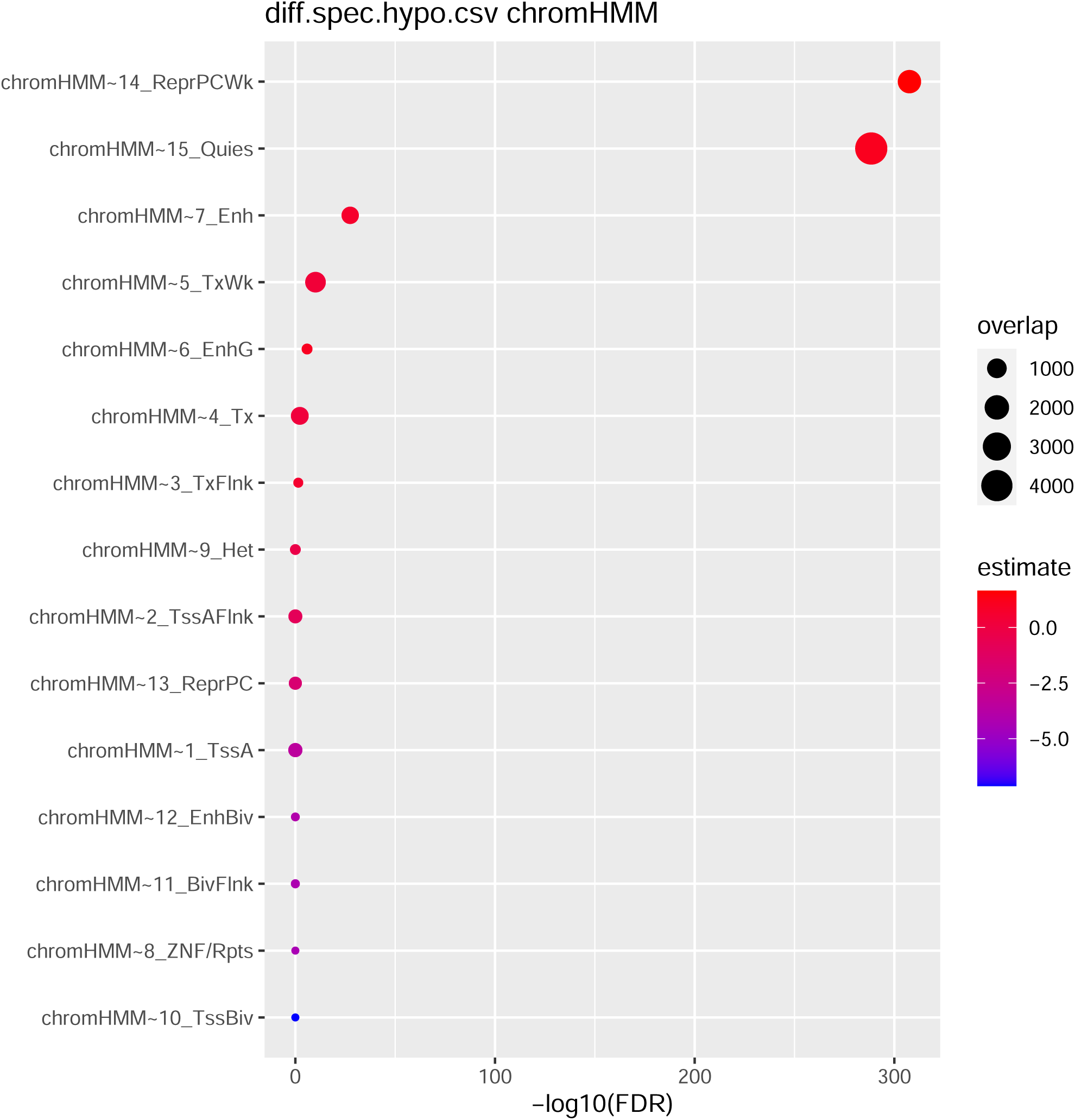
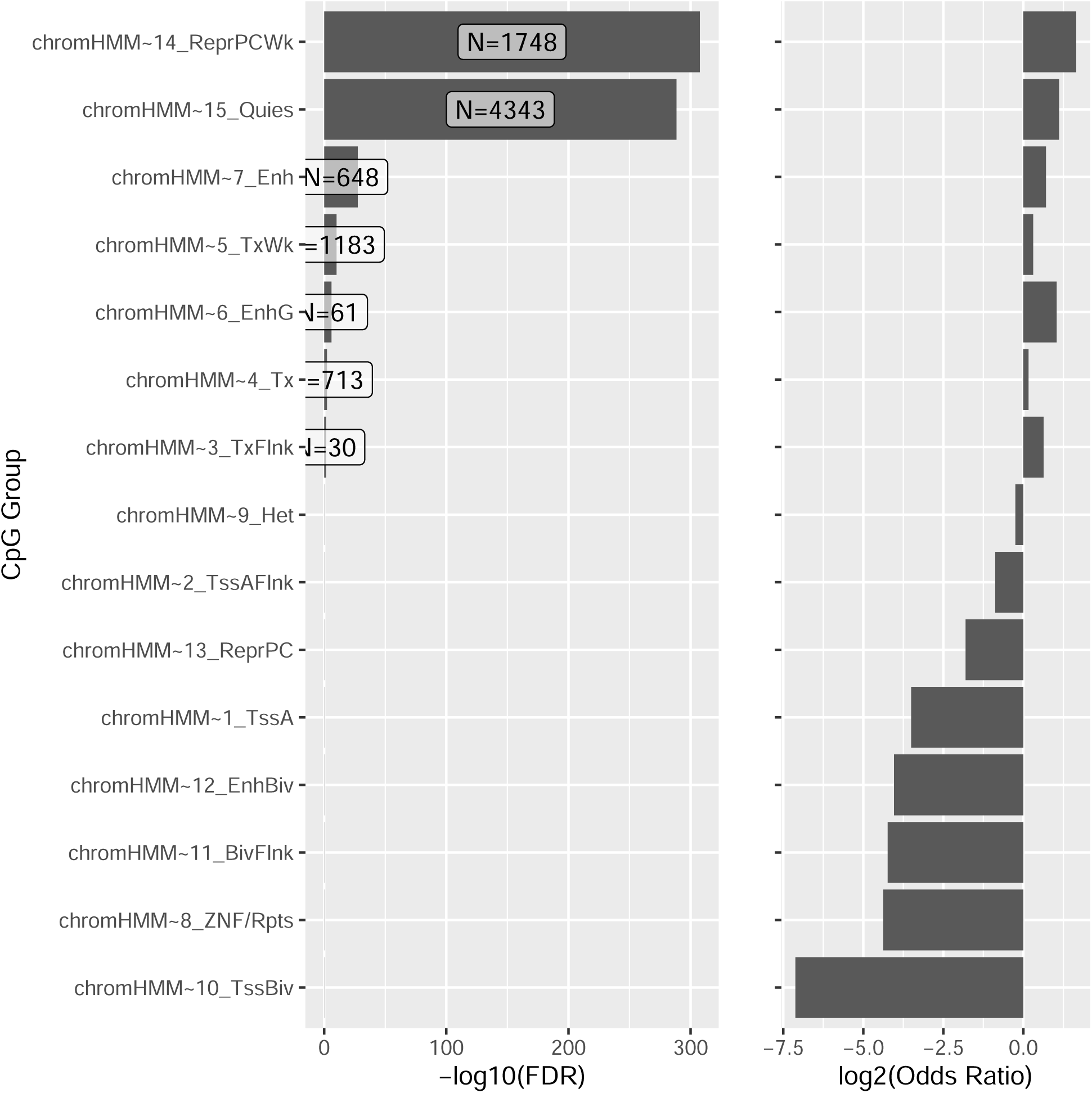
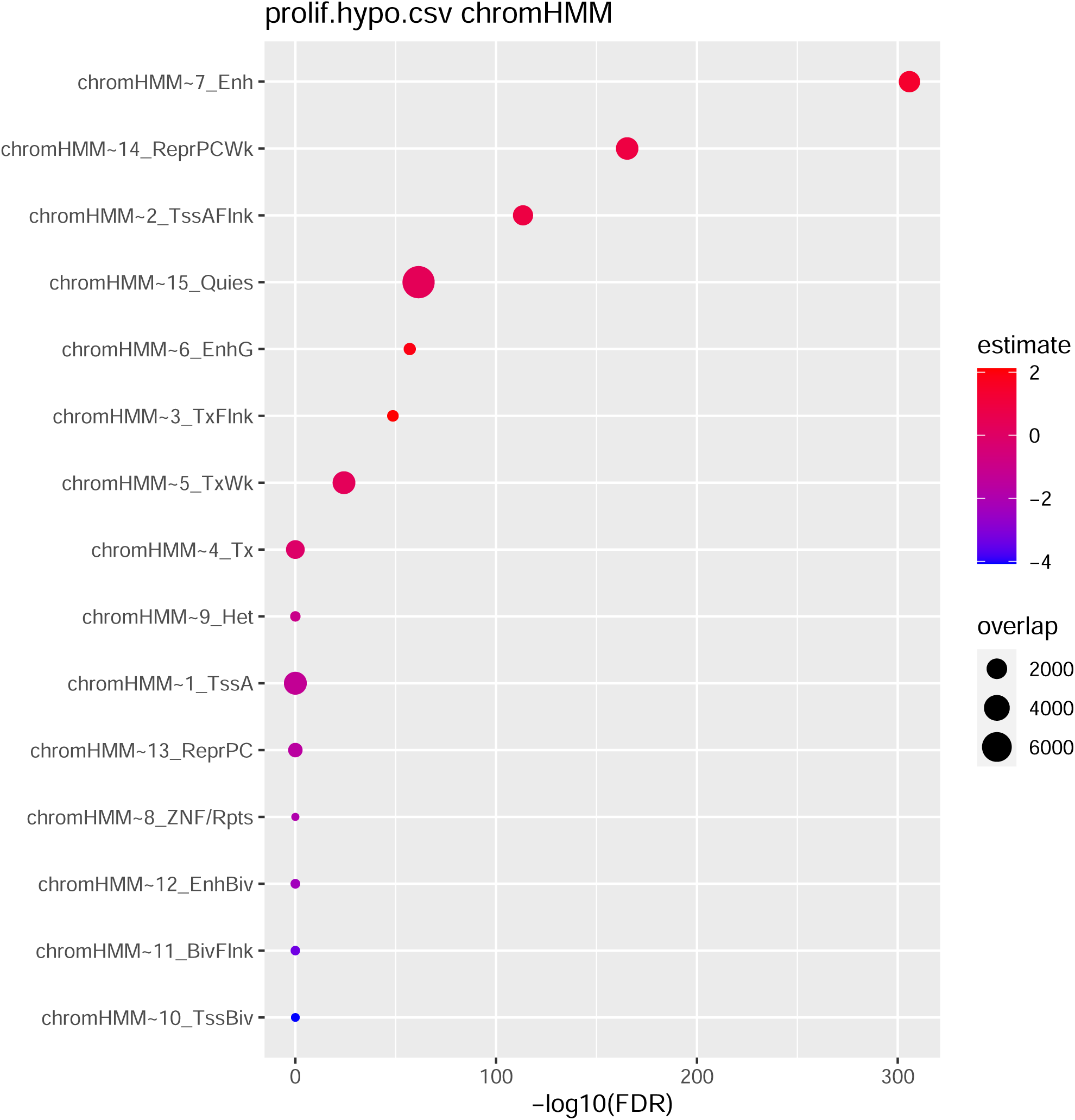
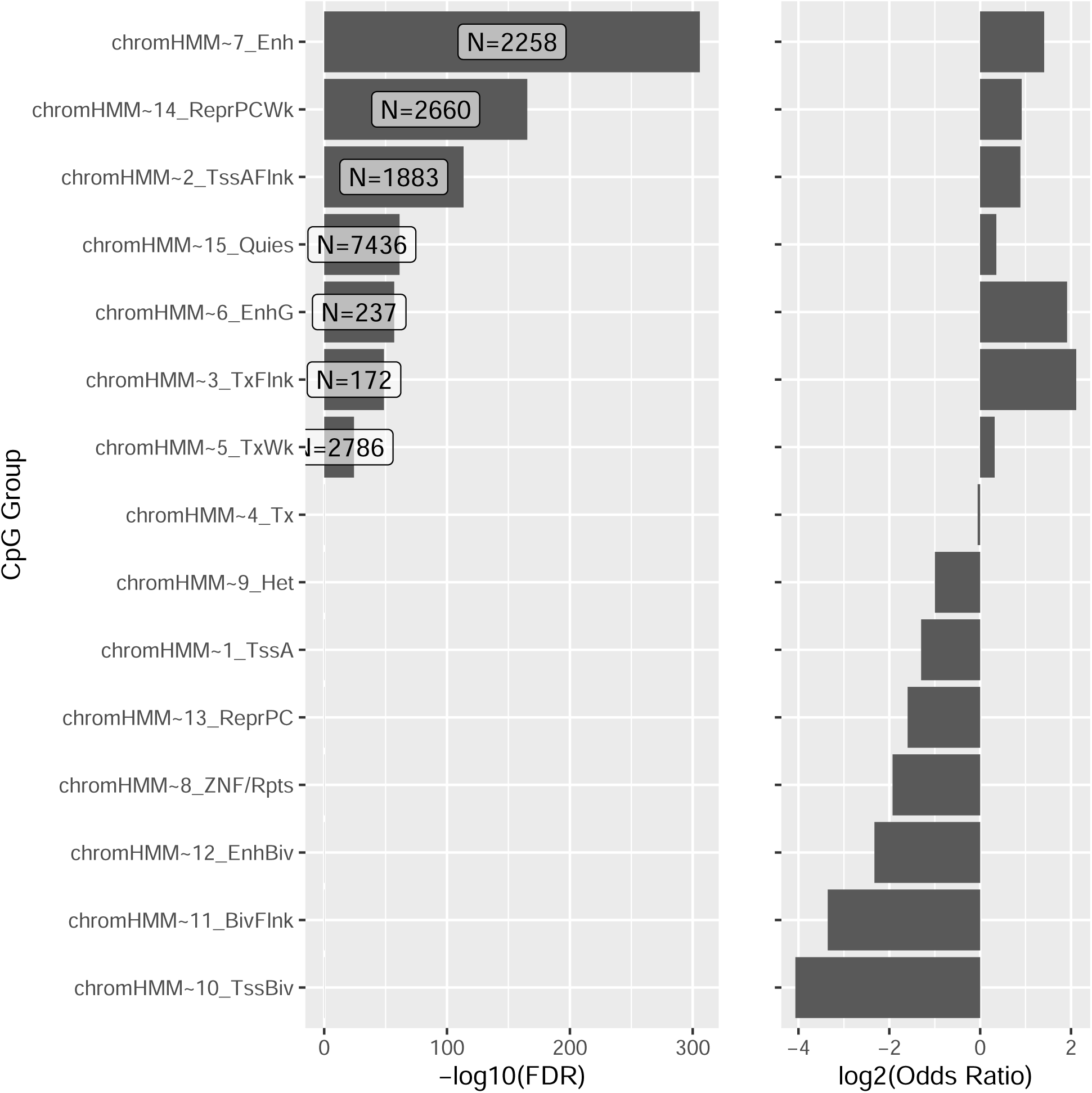
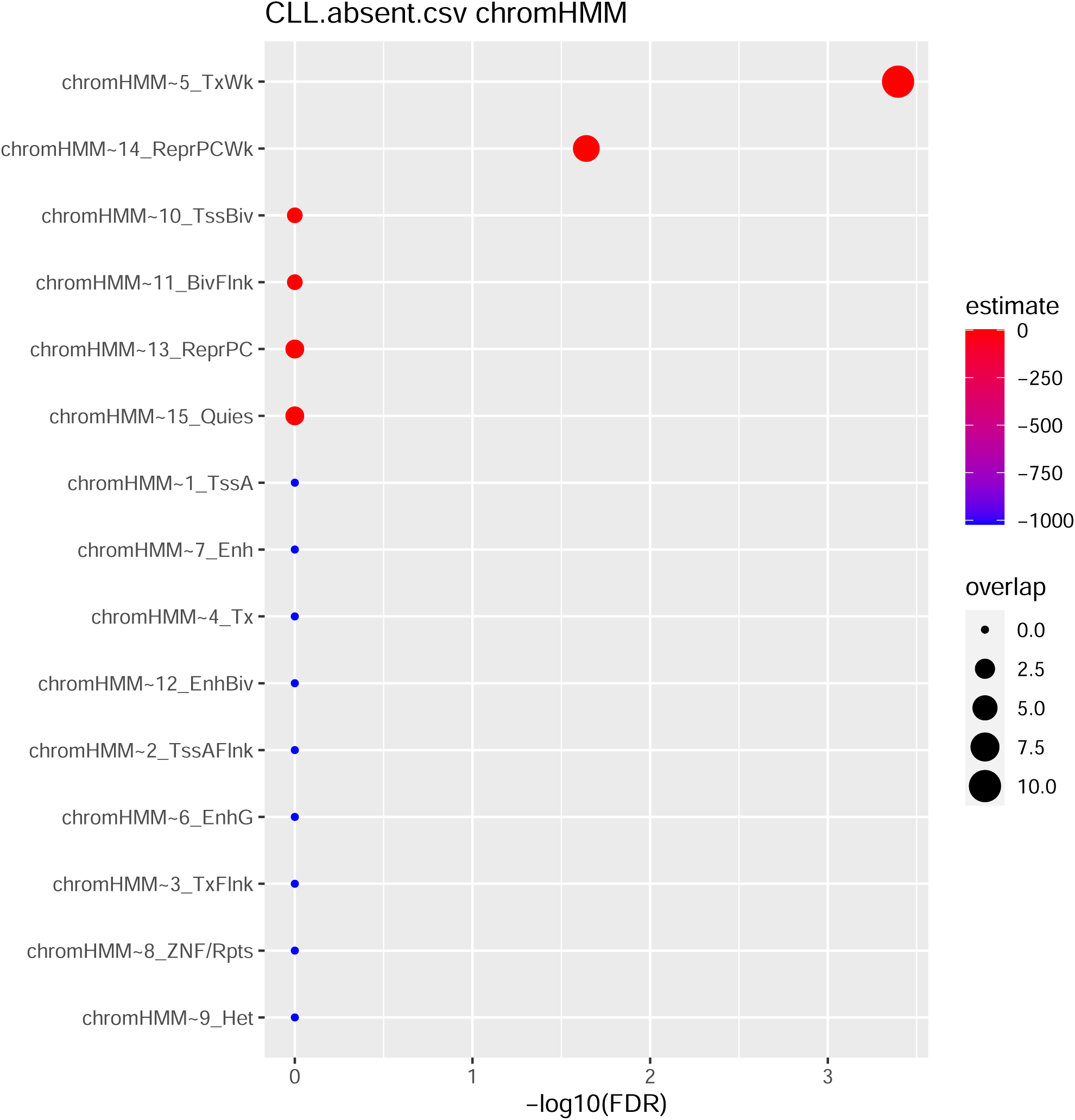
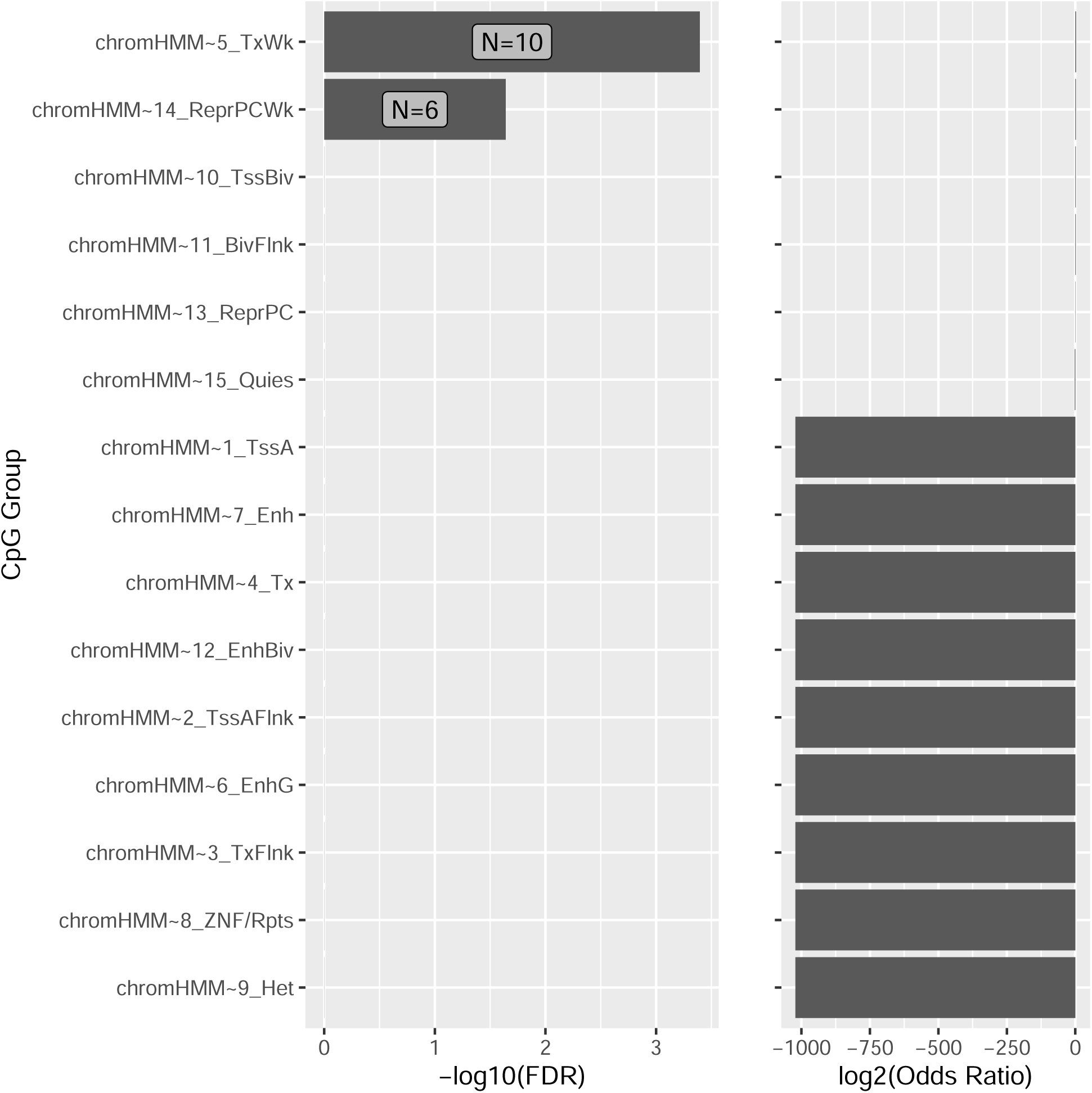
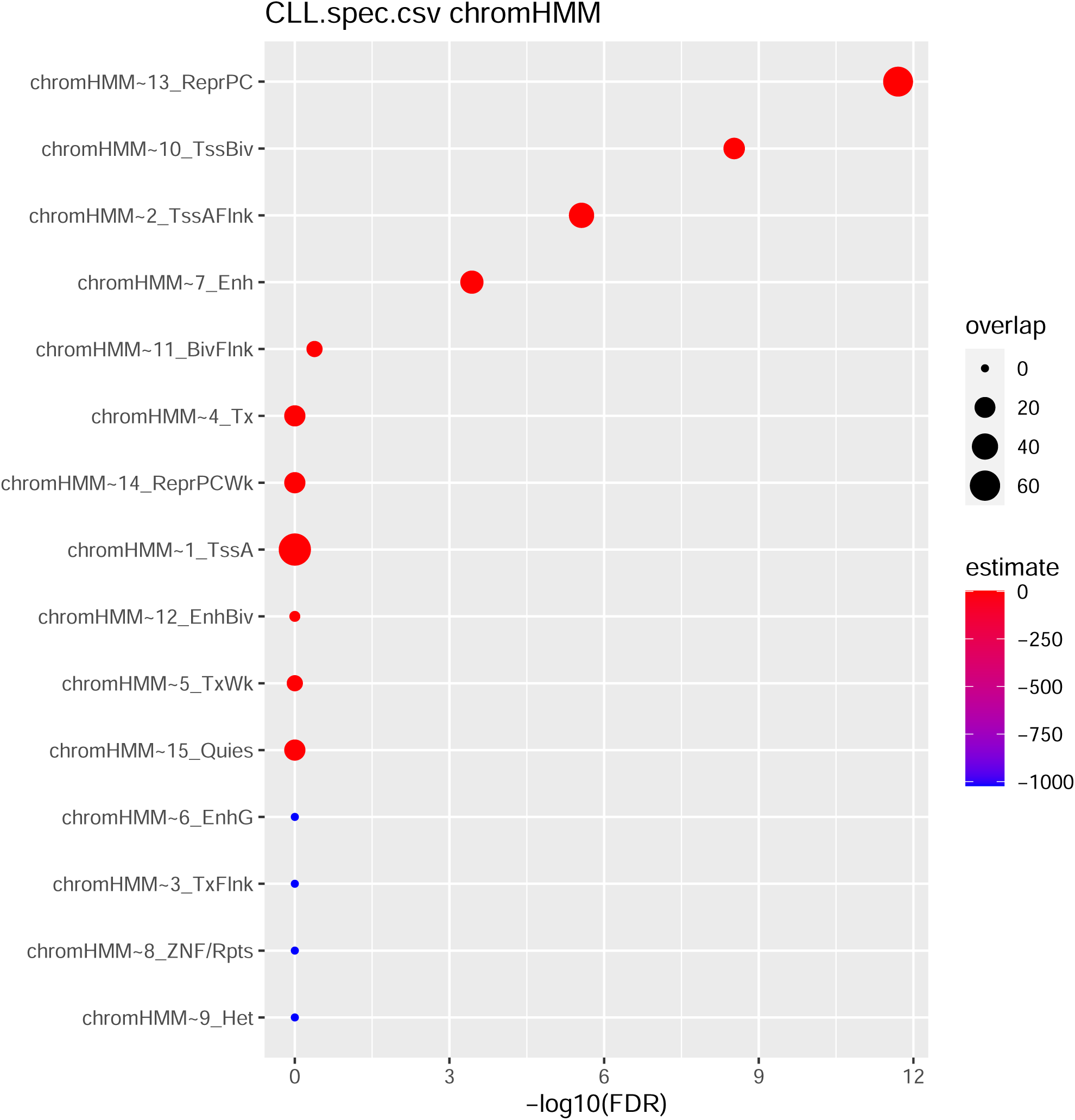
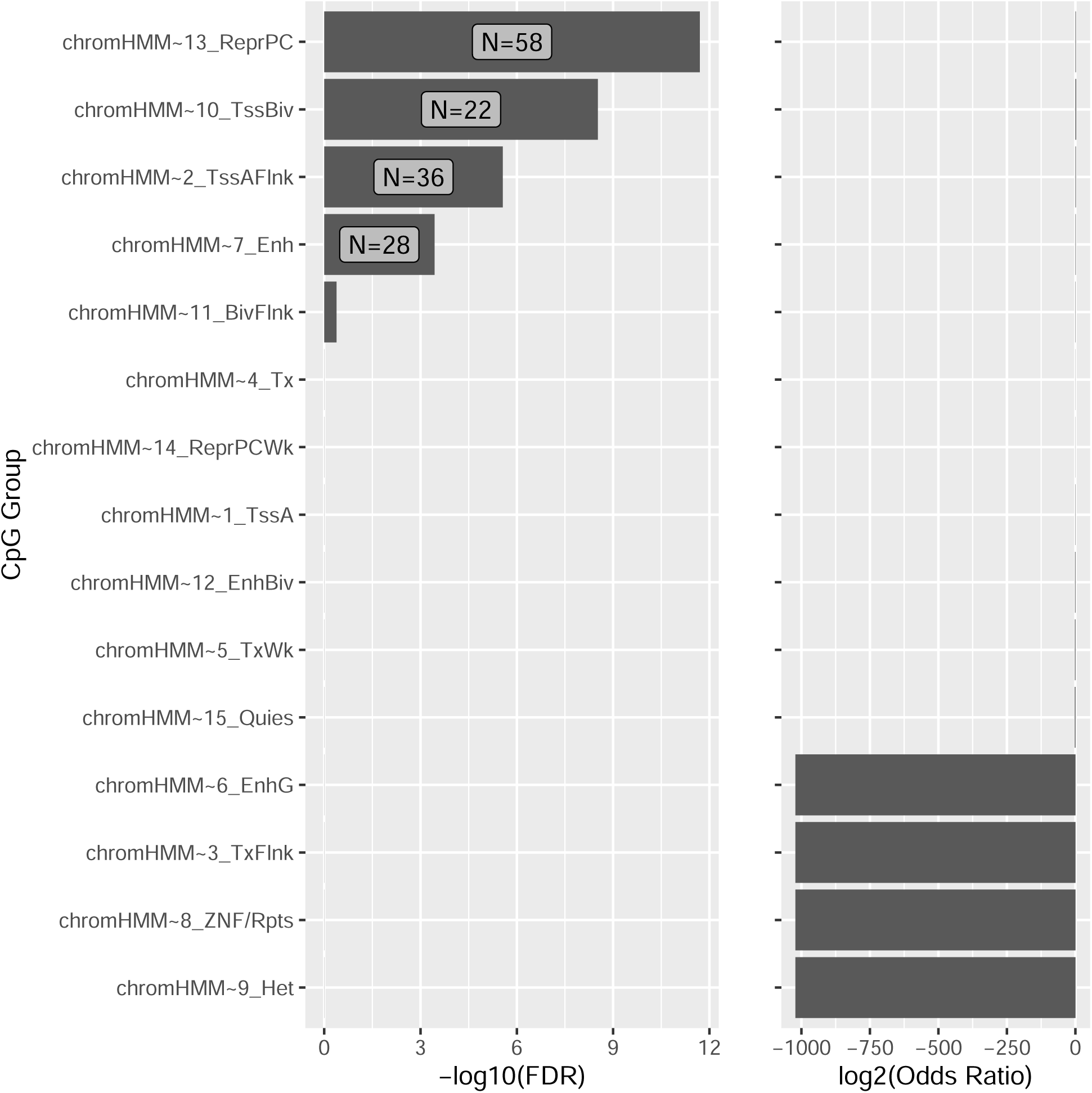
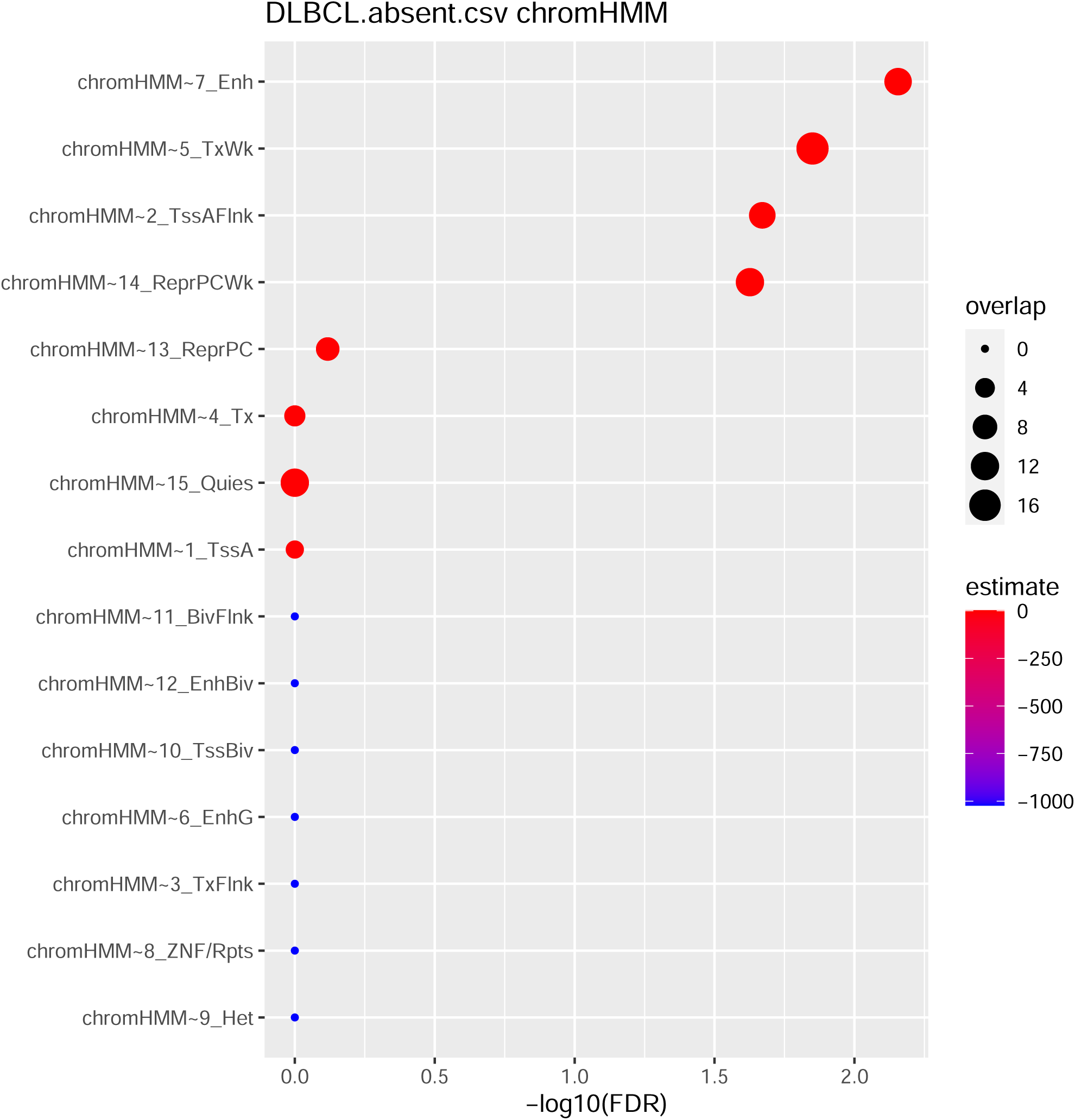
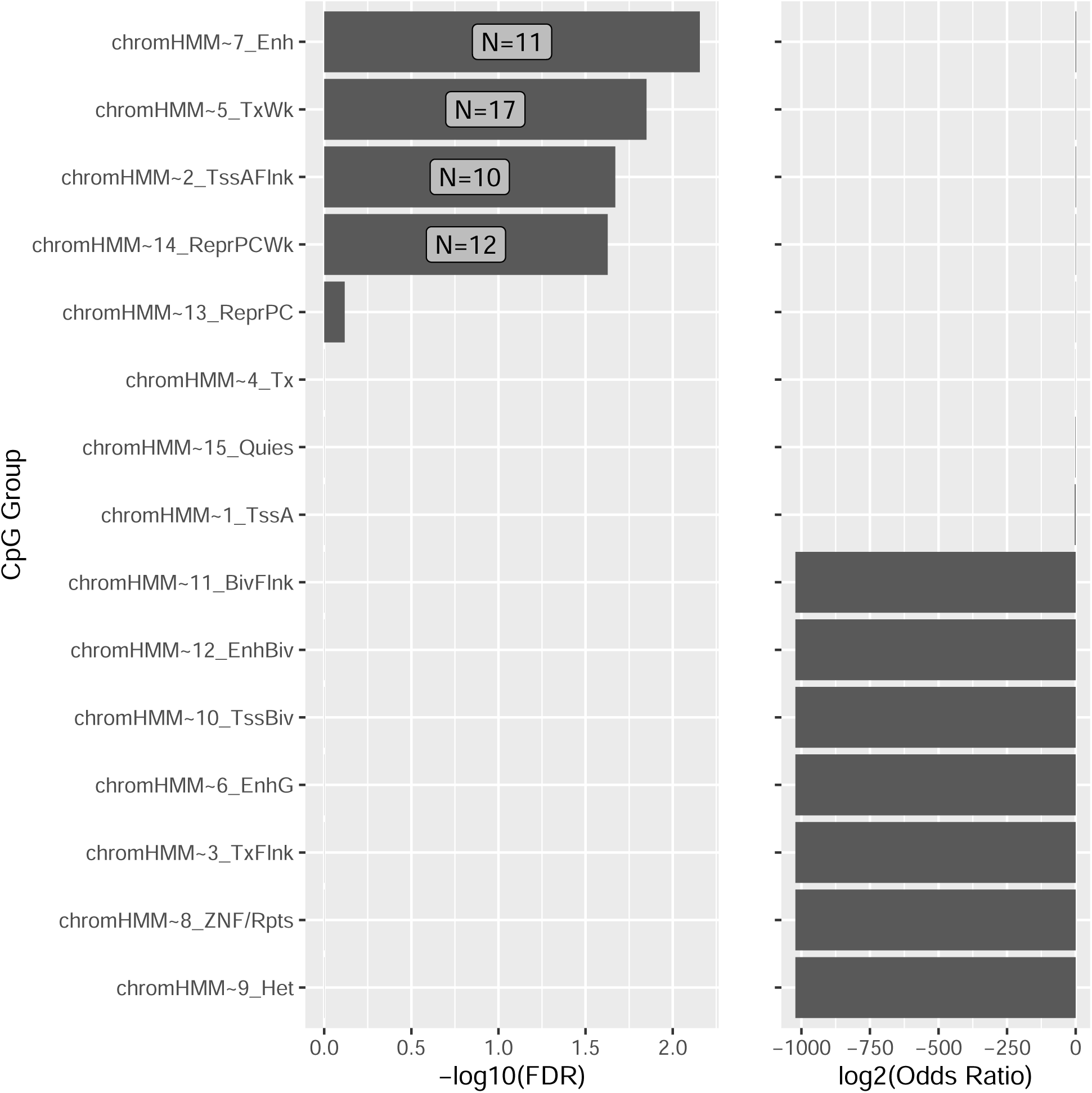
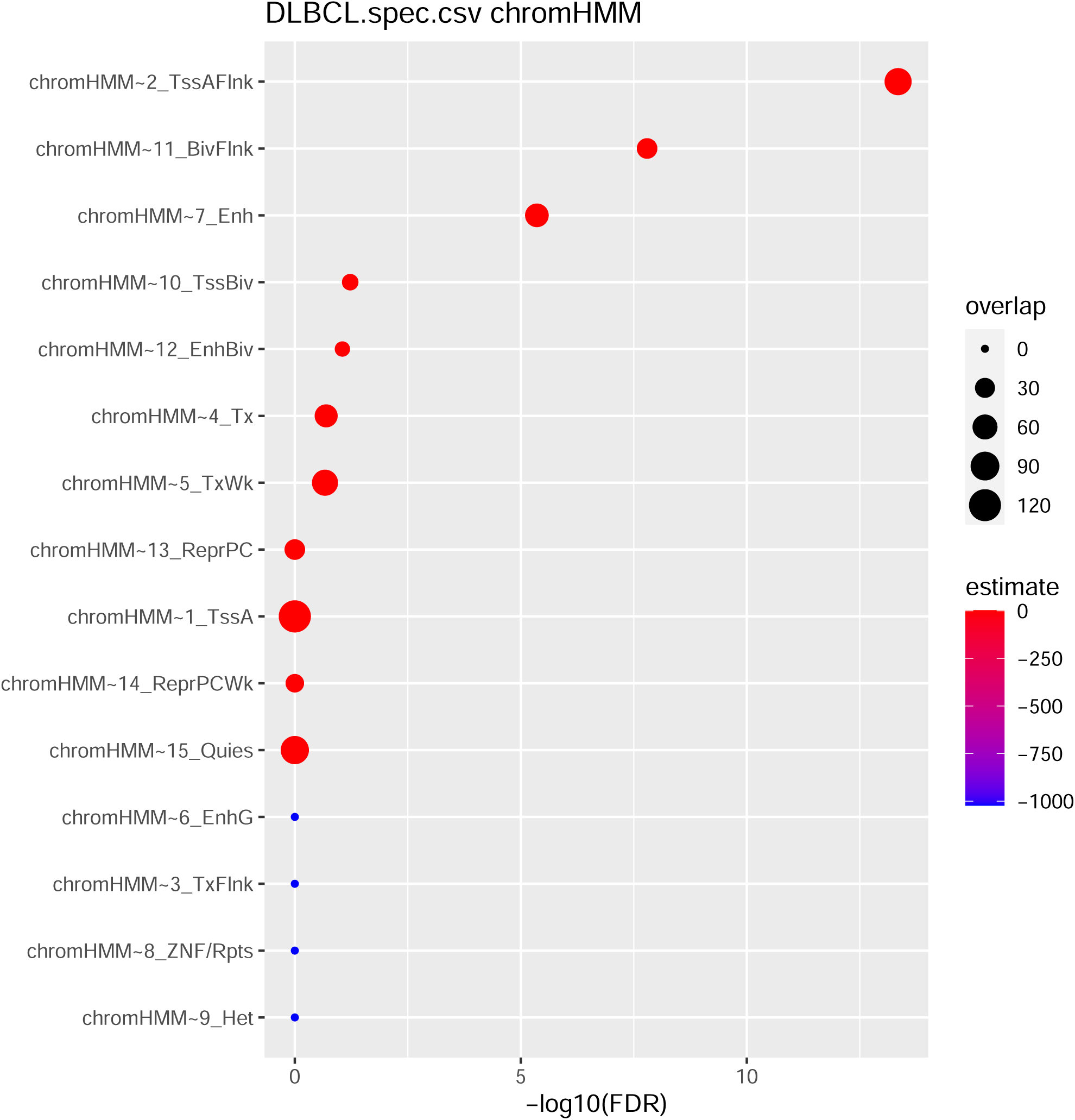
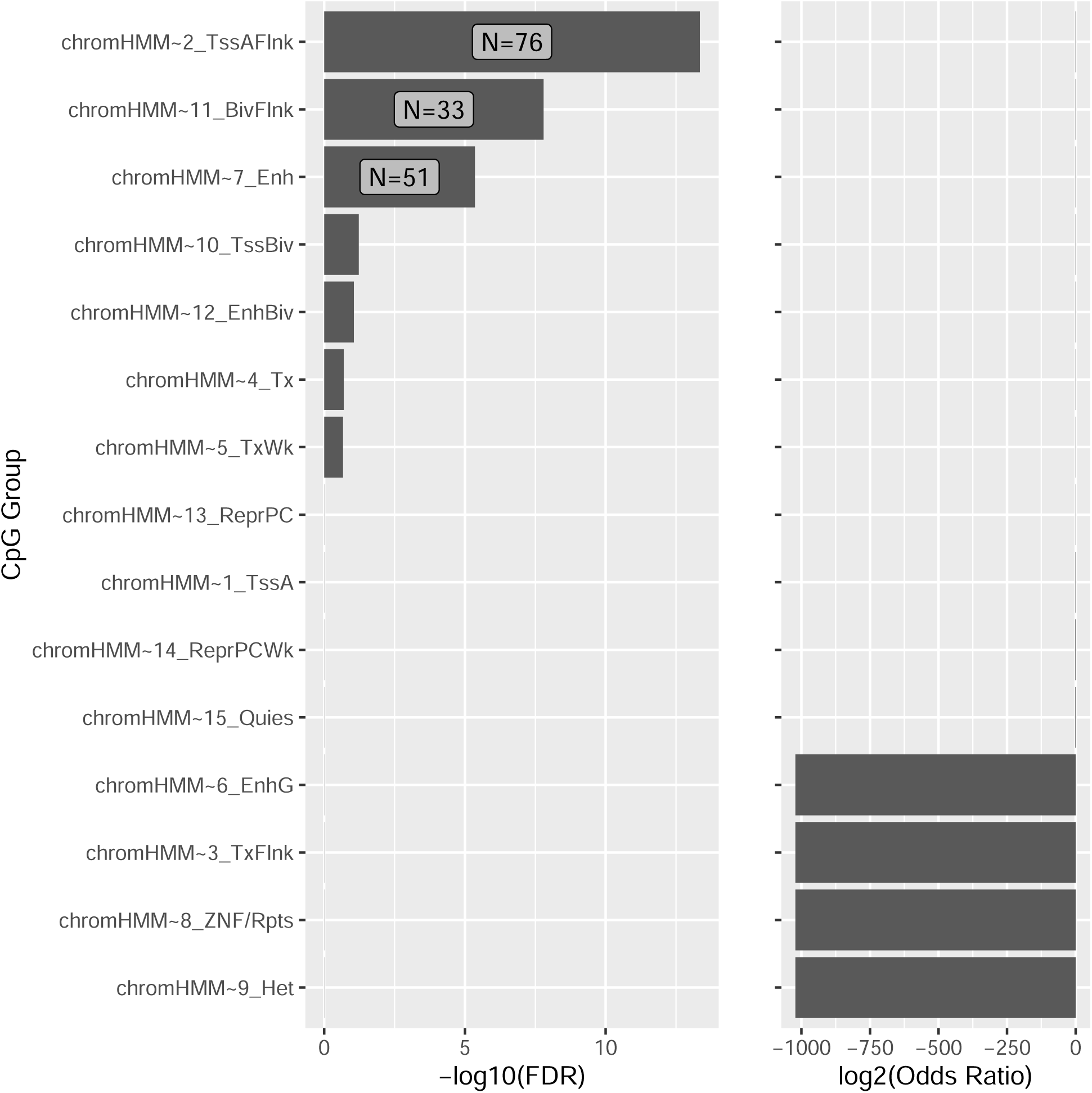
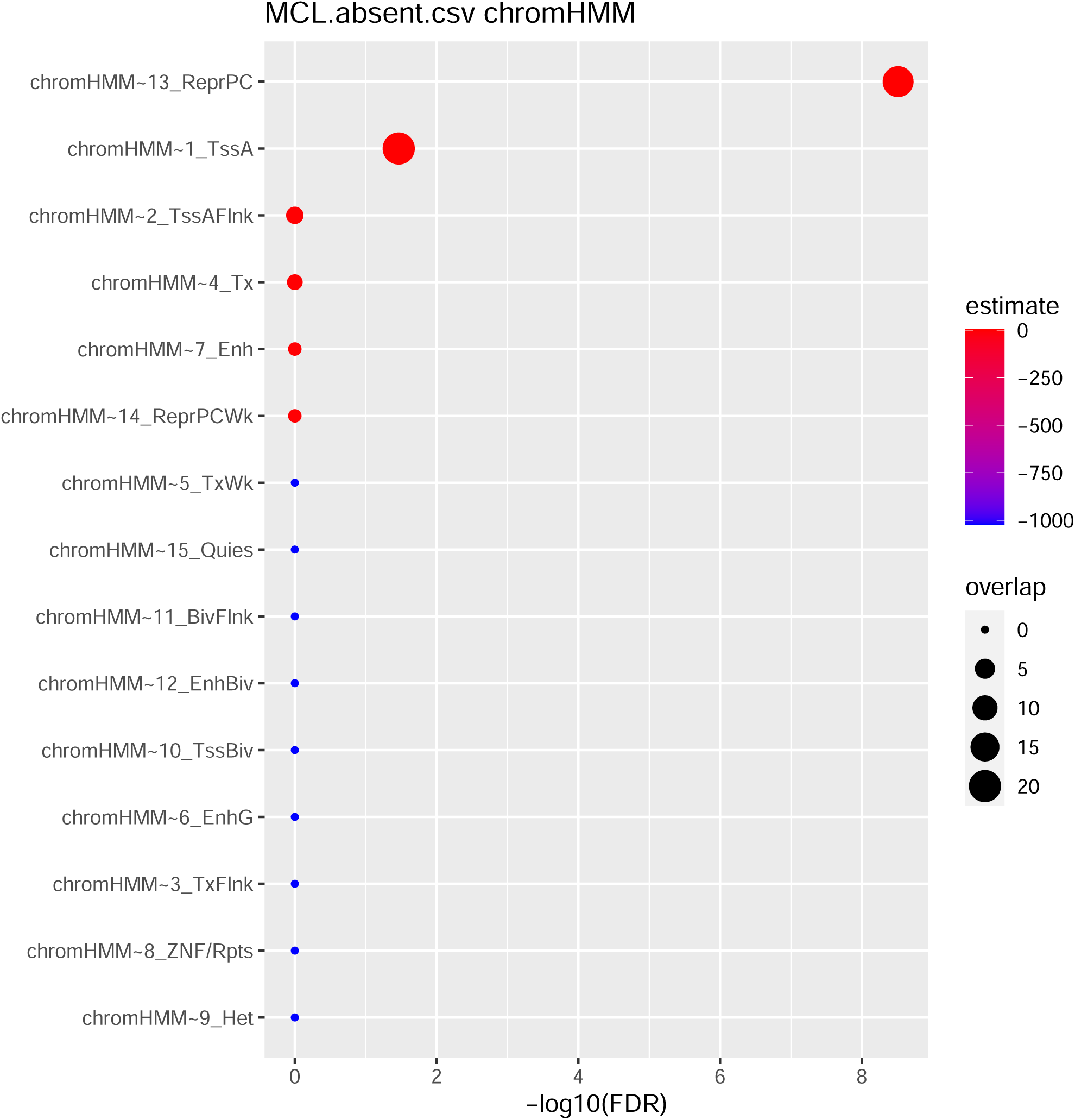
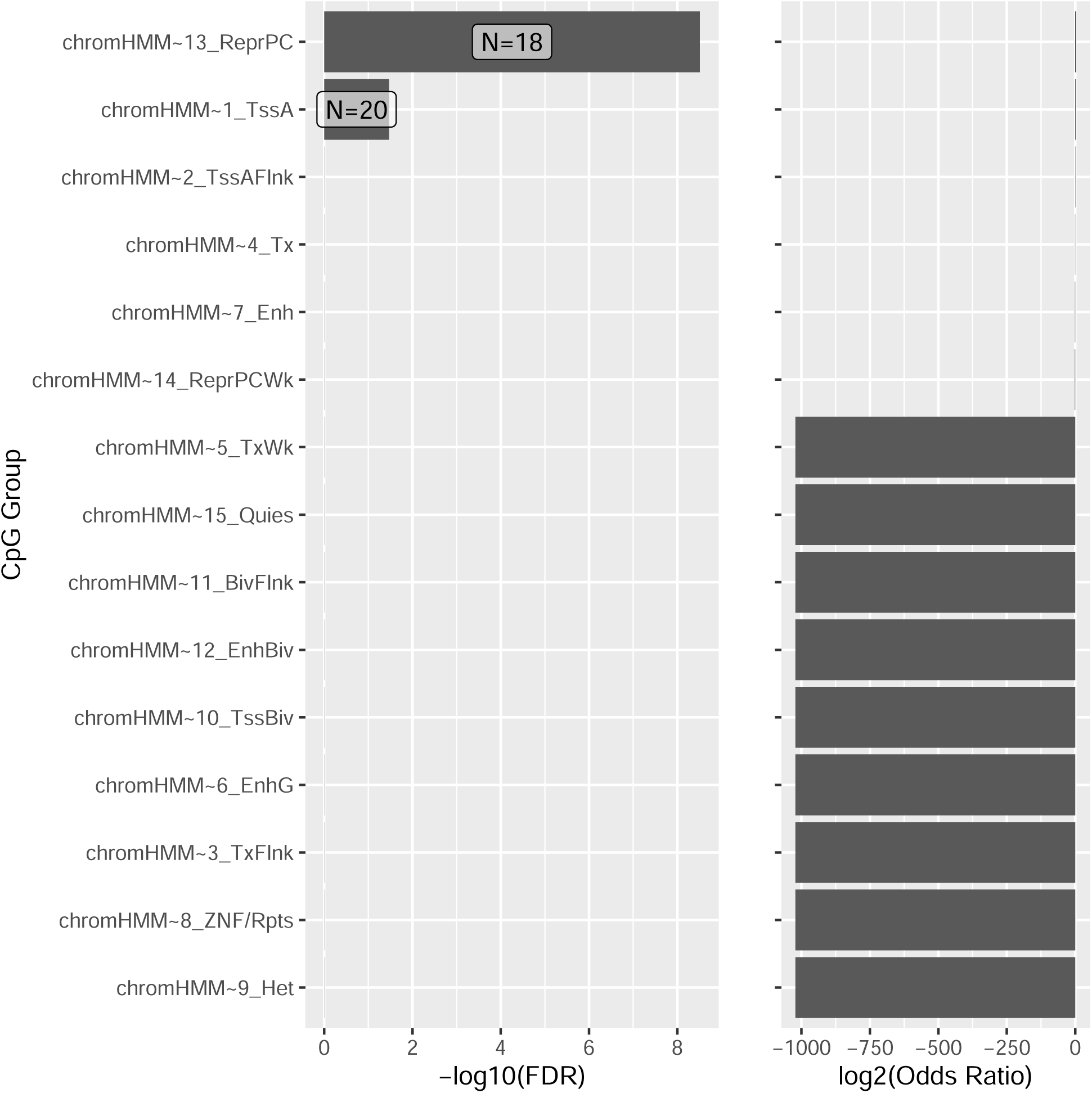
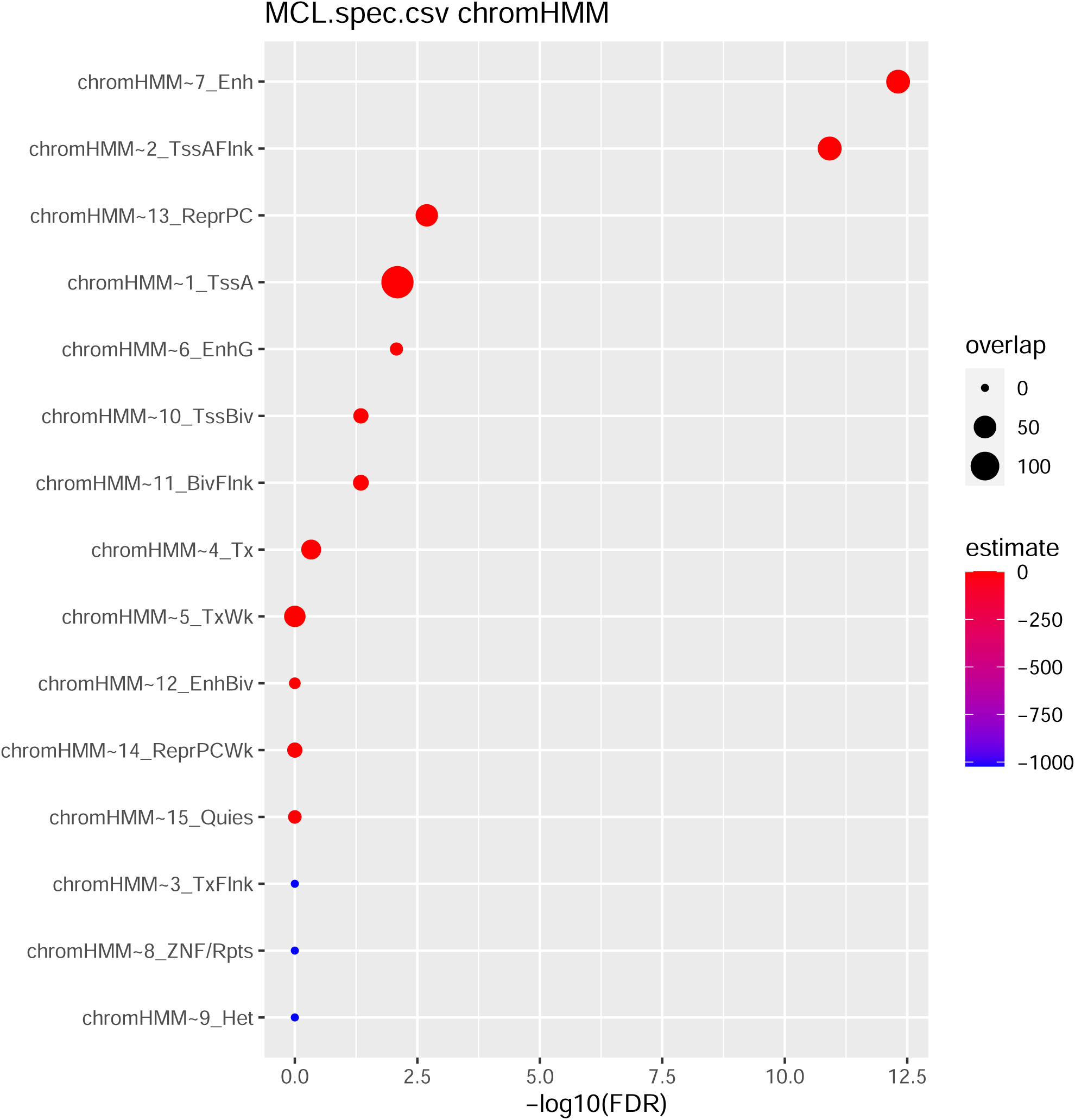
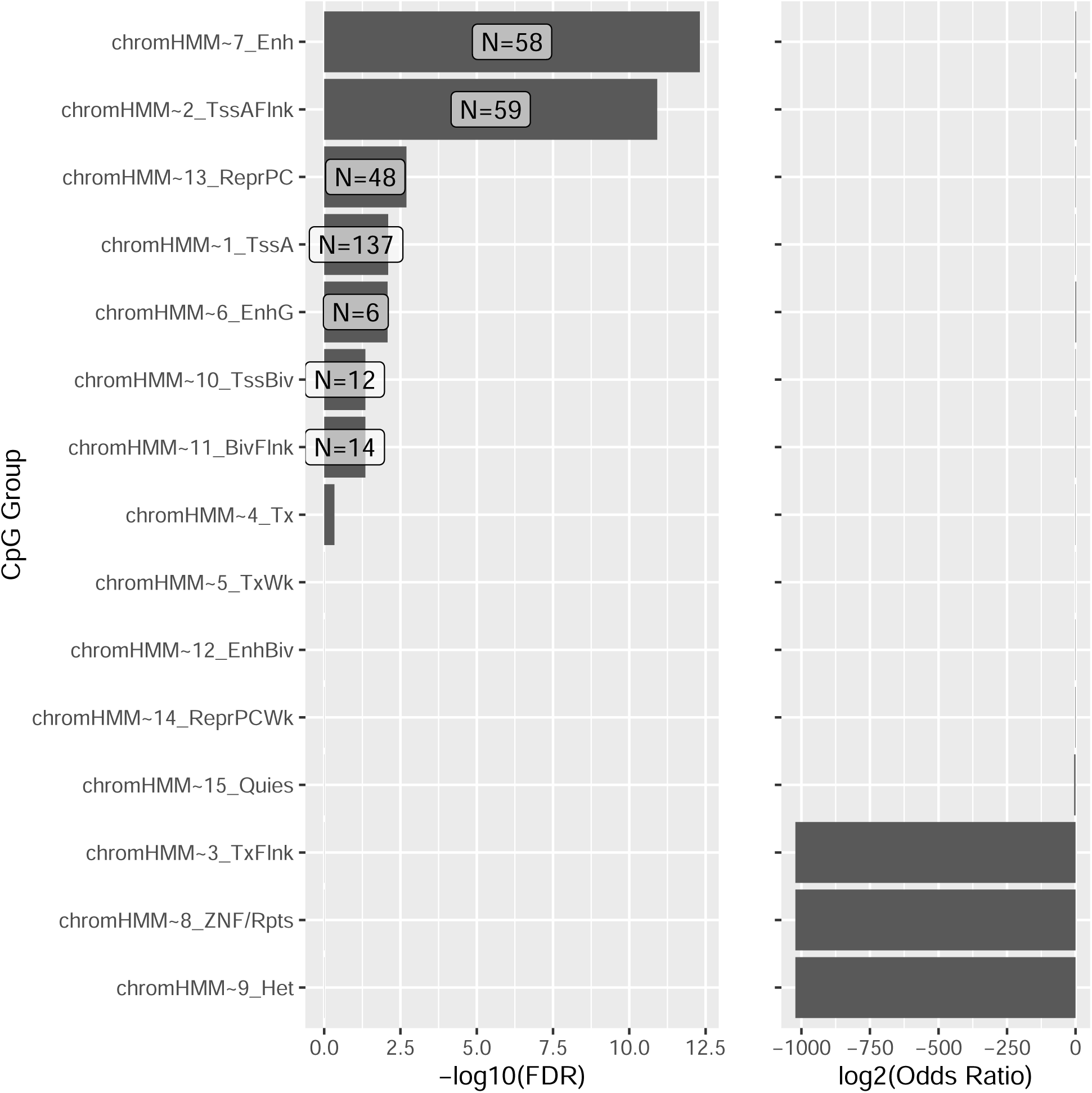
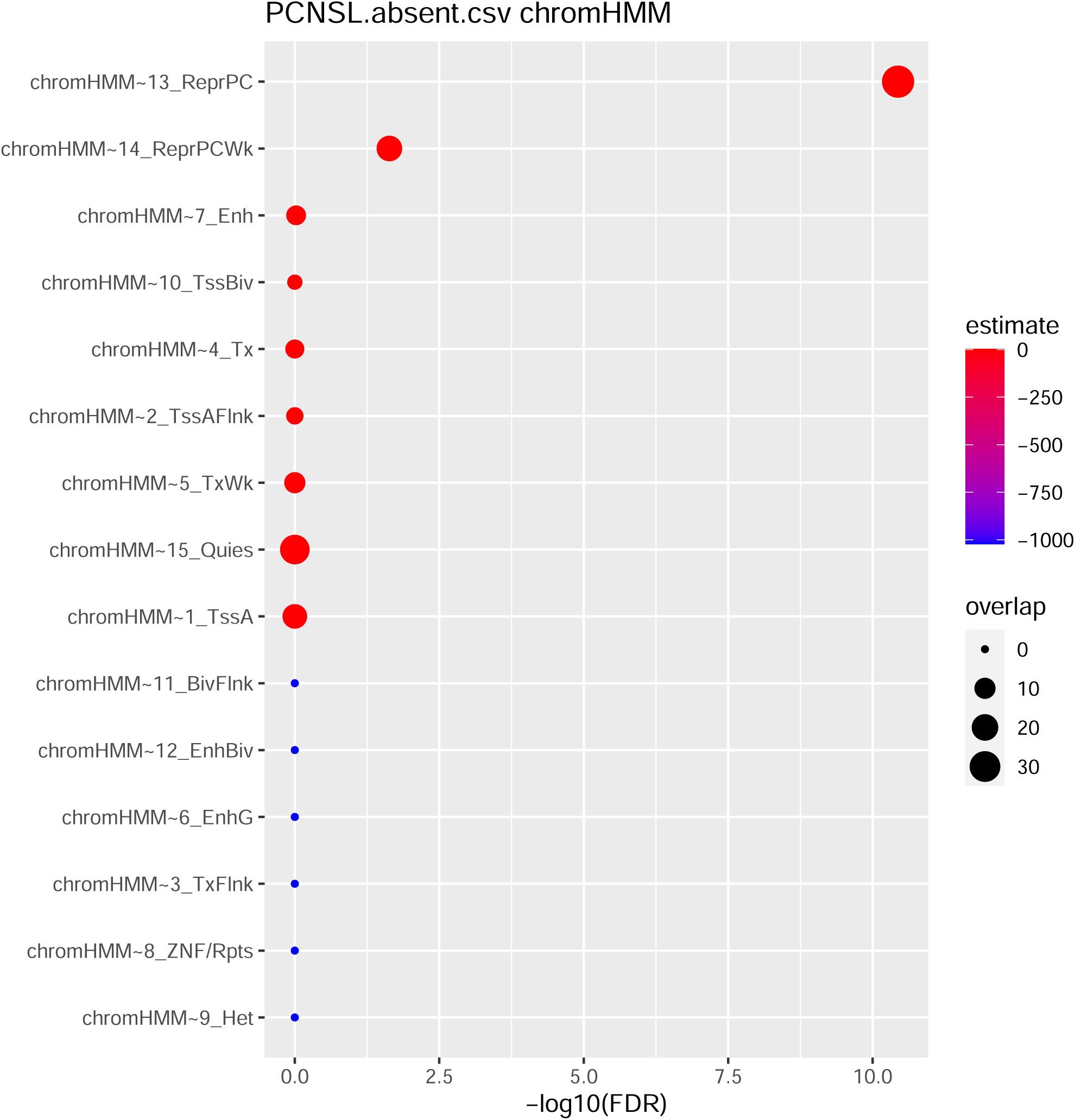
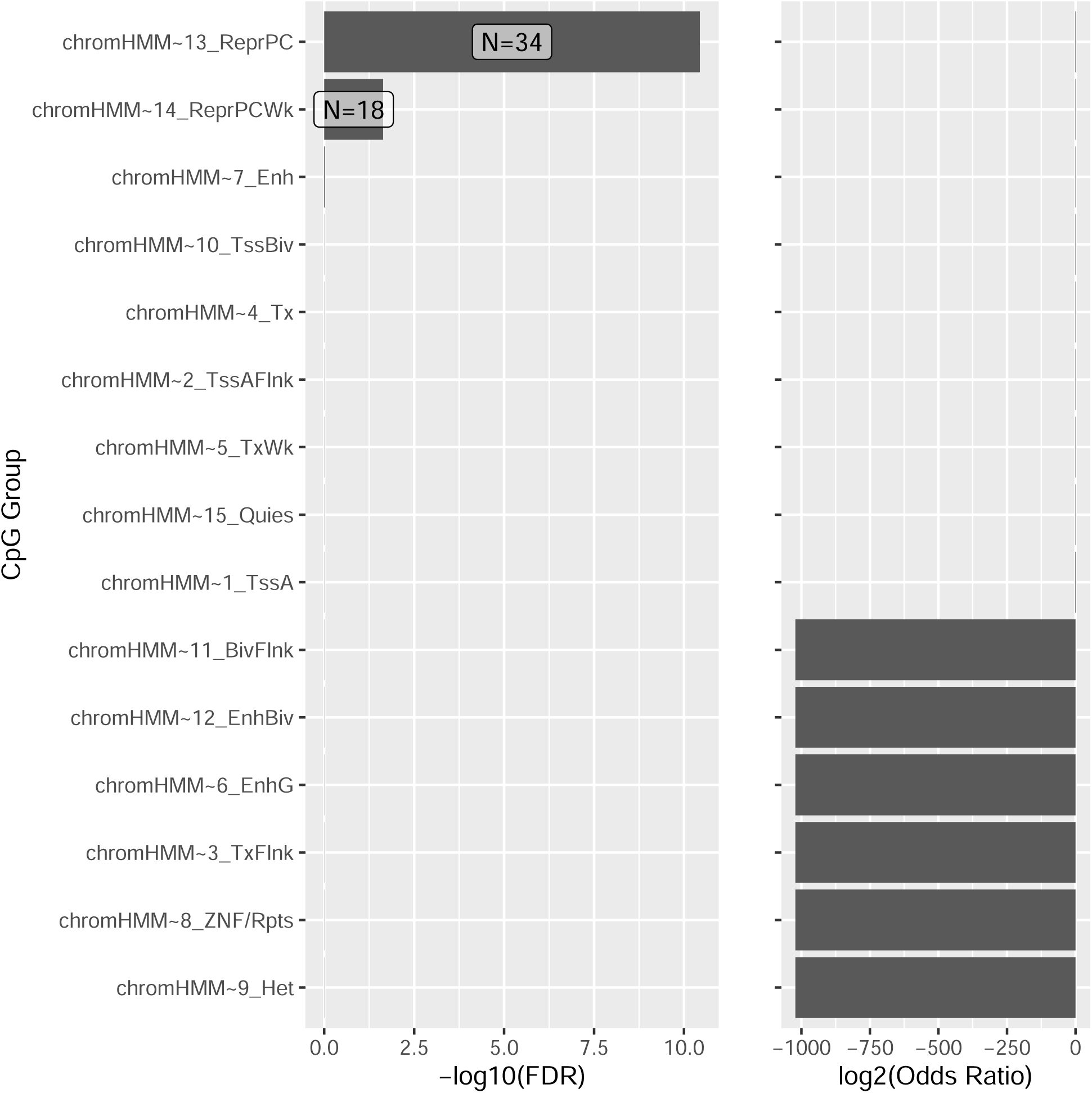
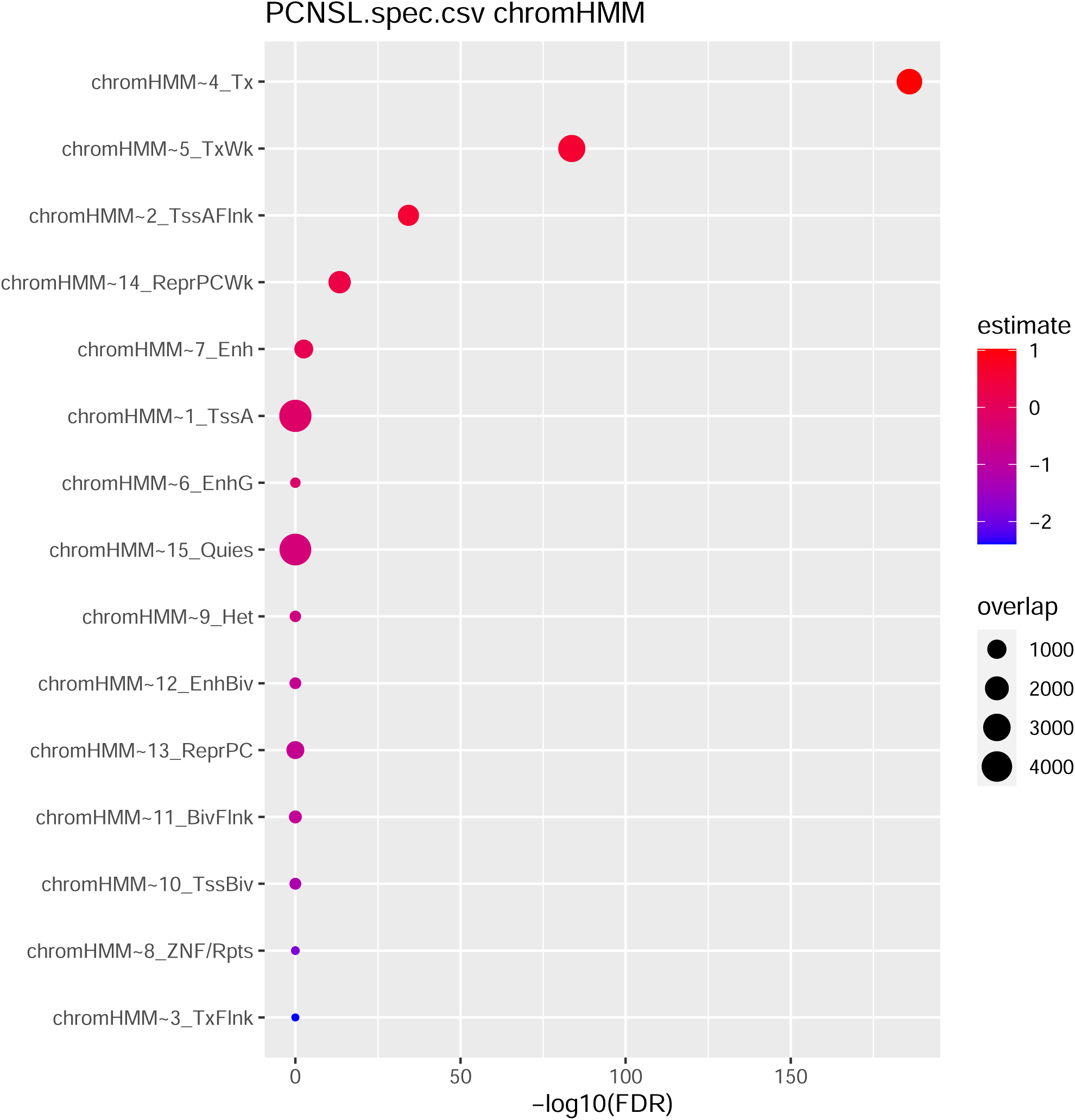
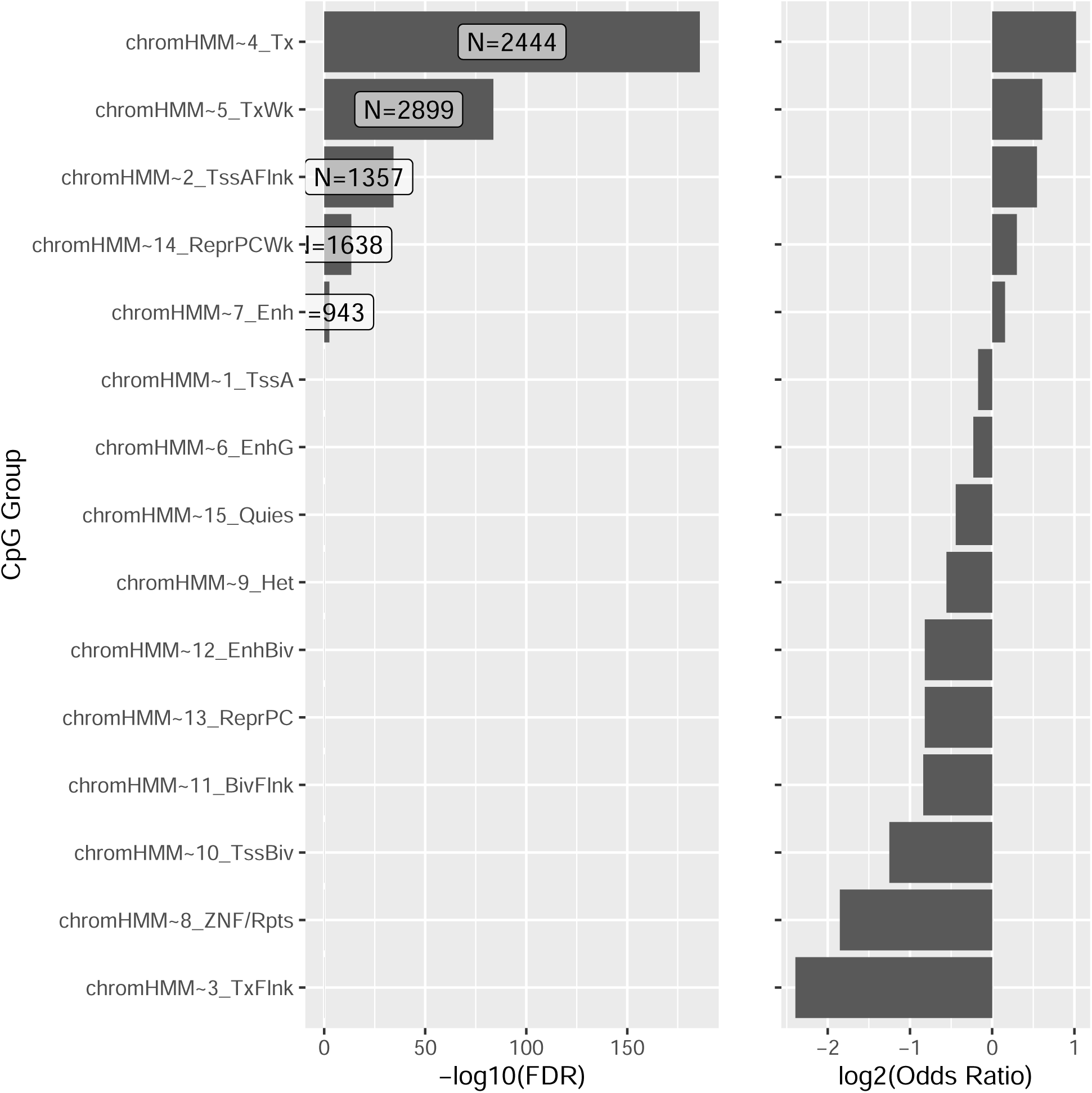
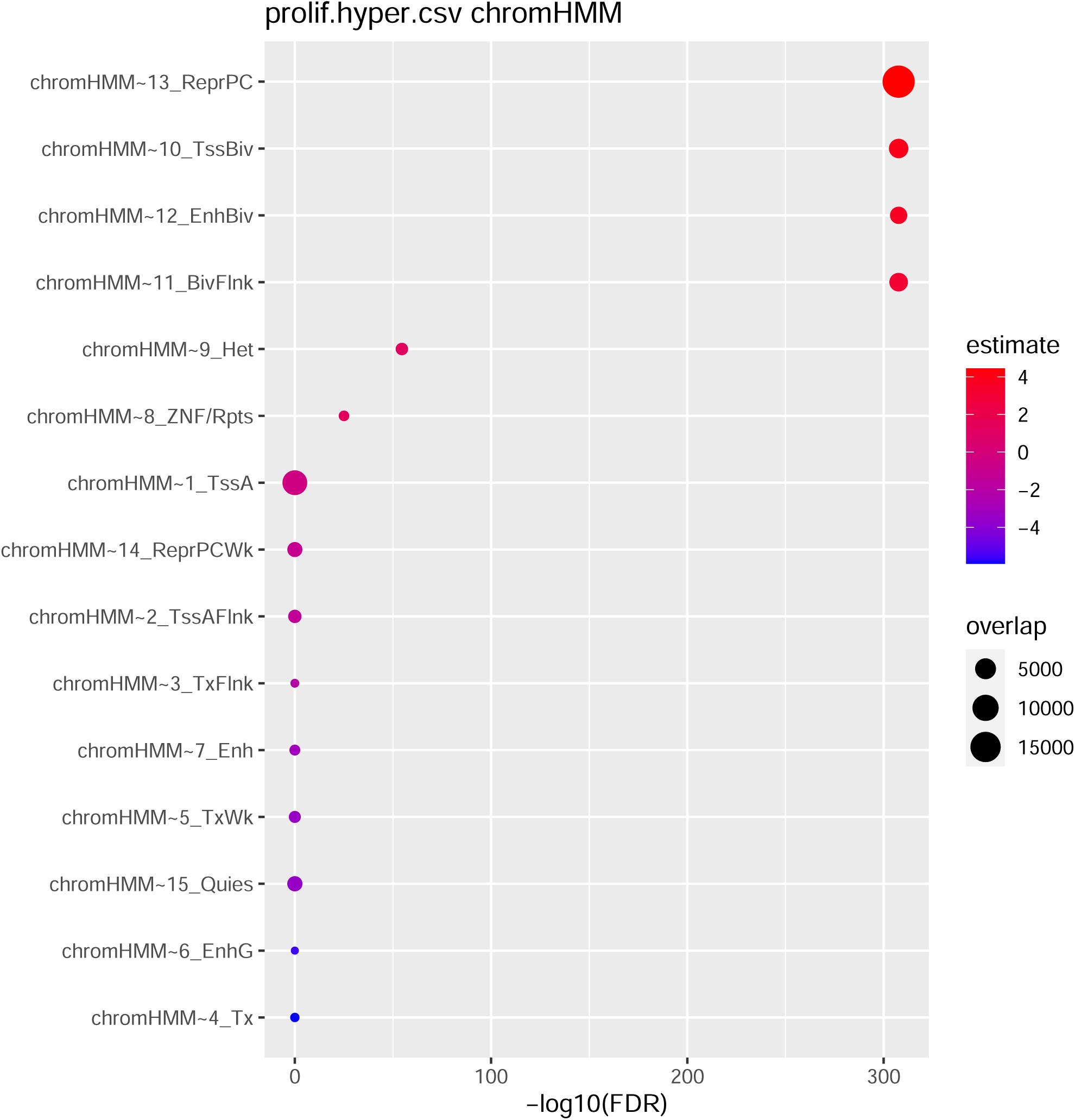
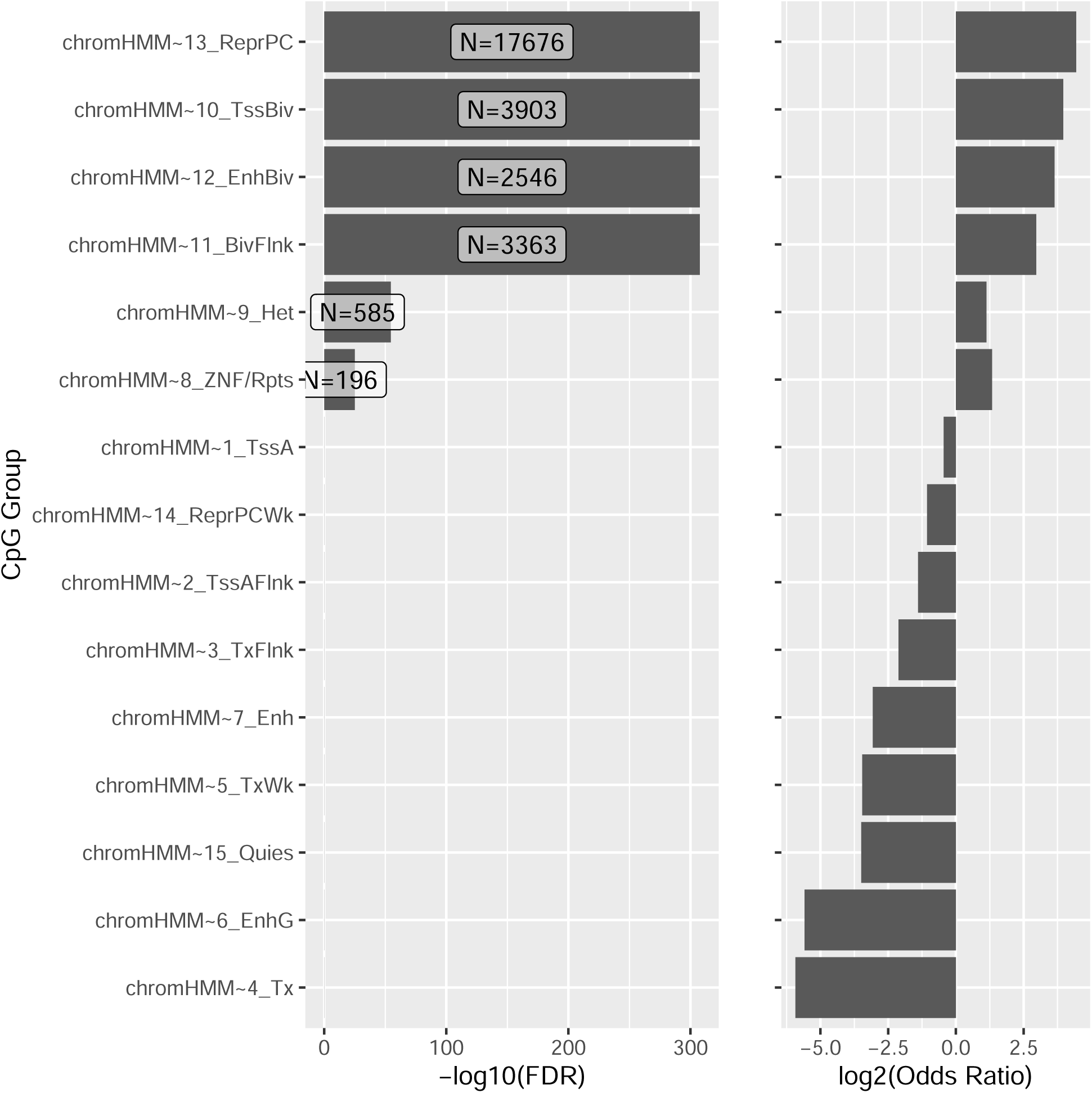

**Supplementary Figure 4.**
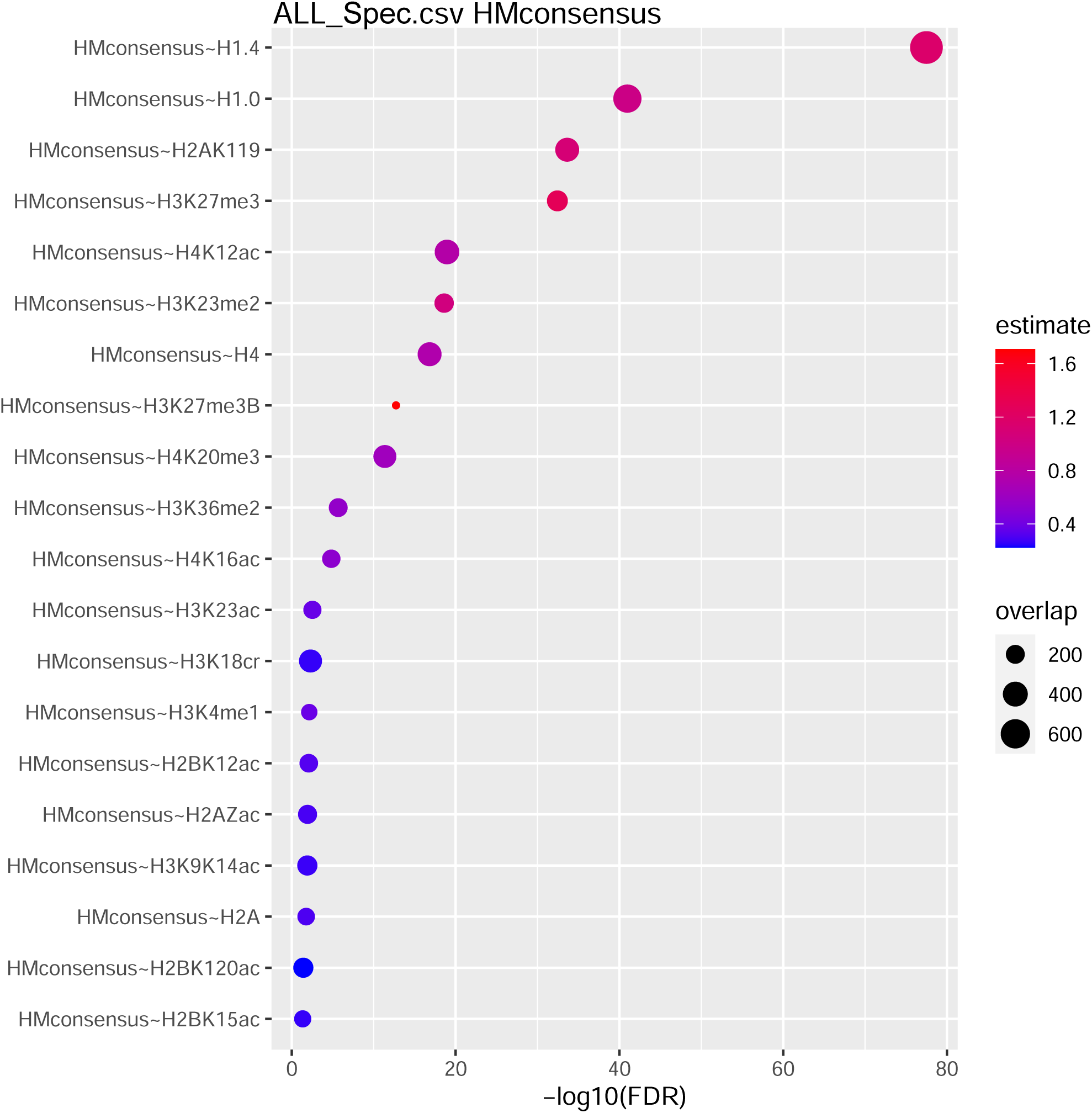
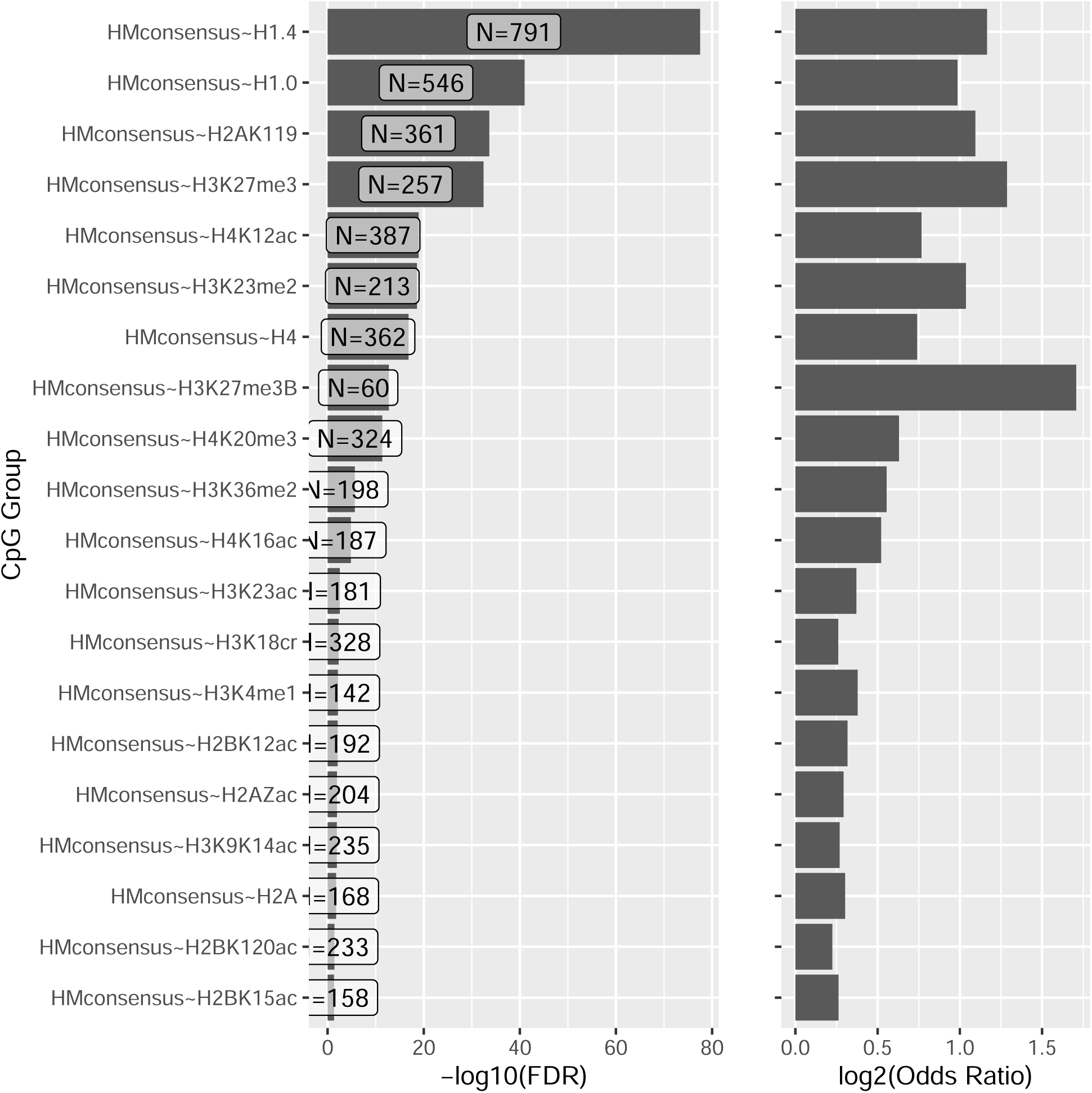
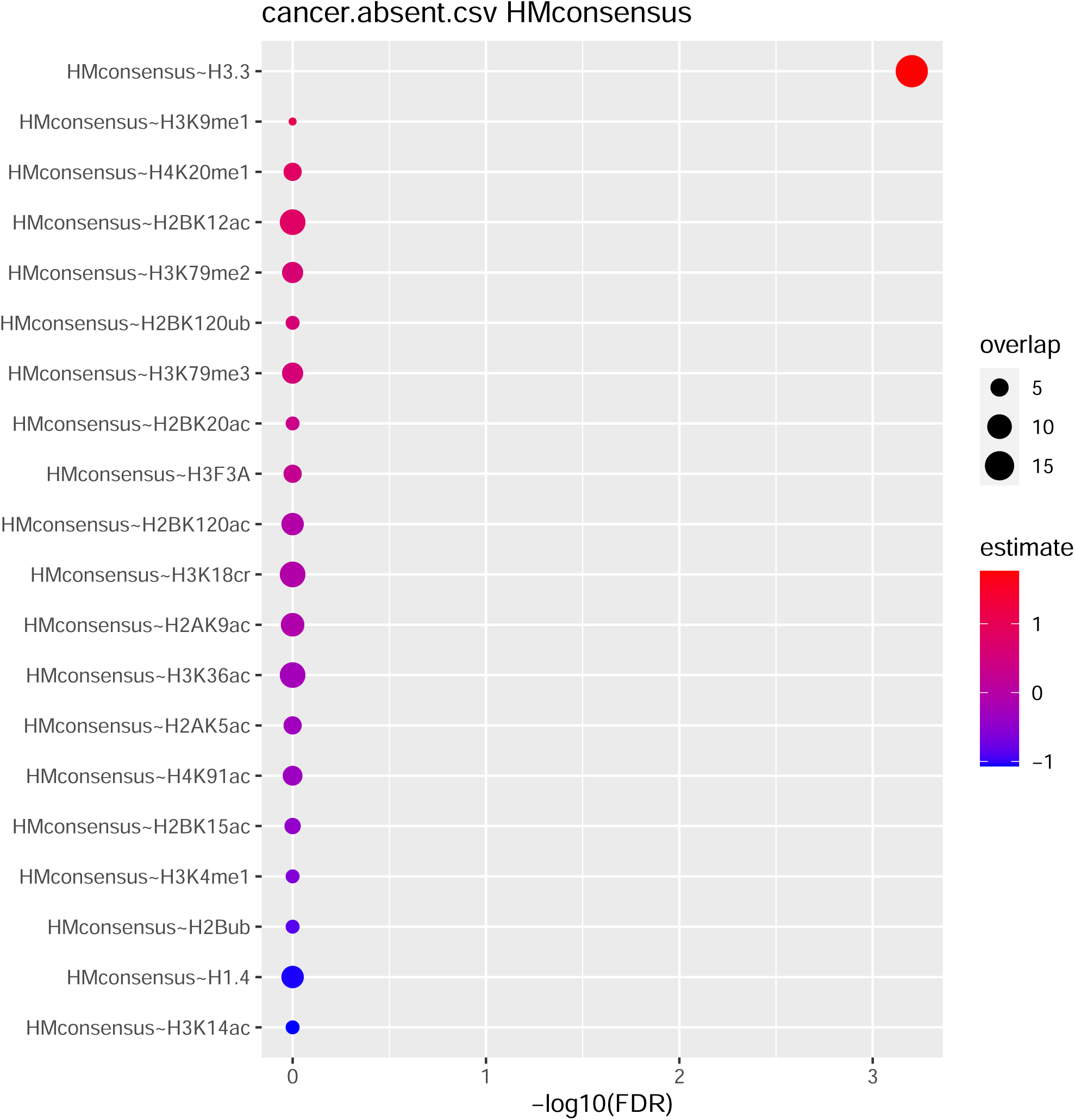
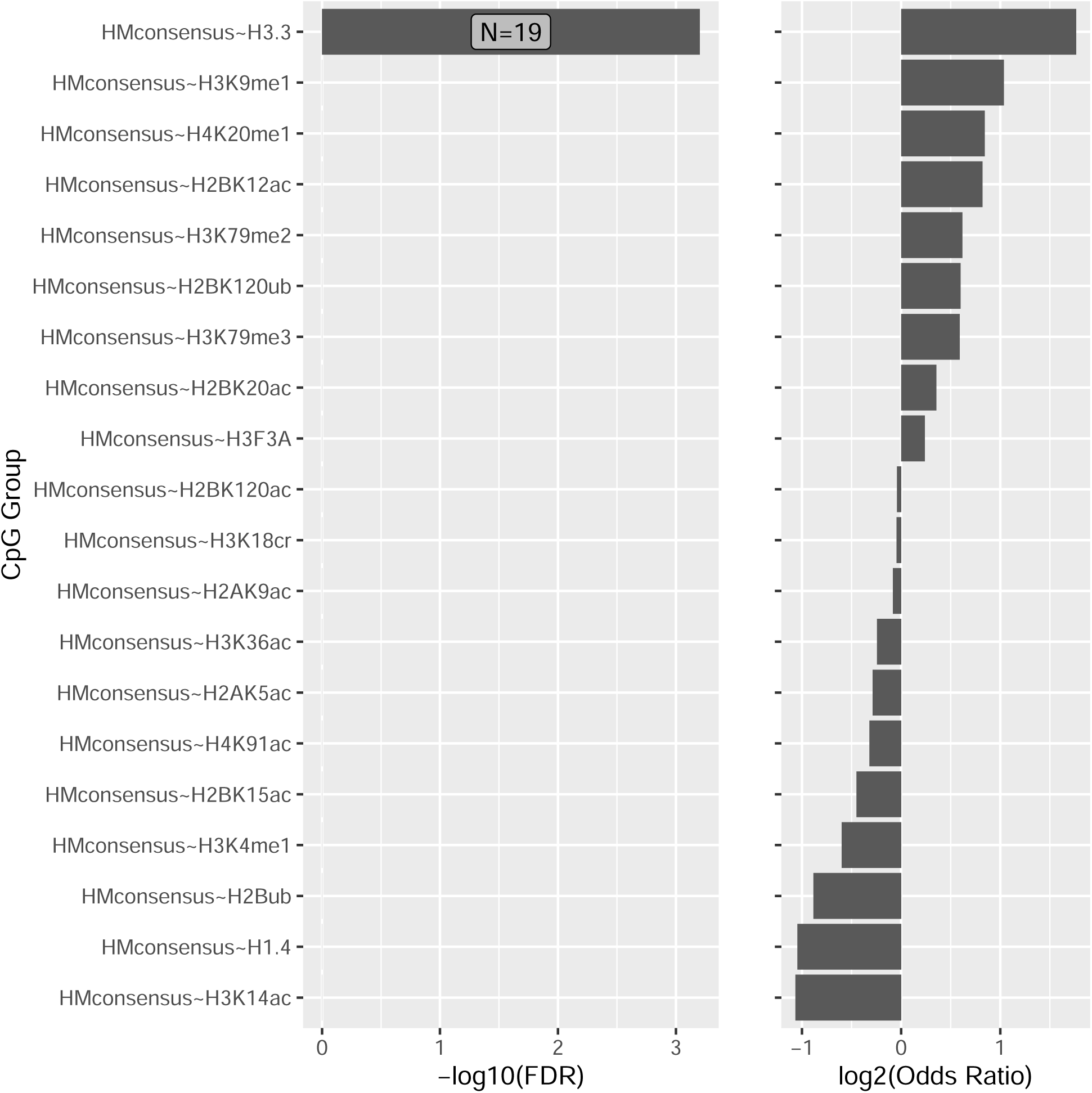
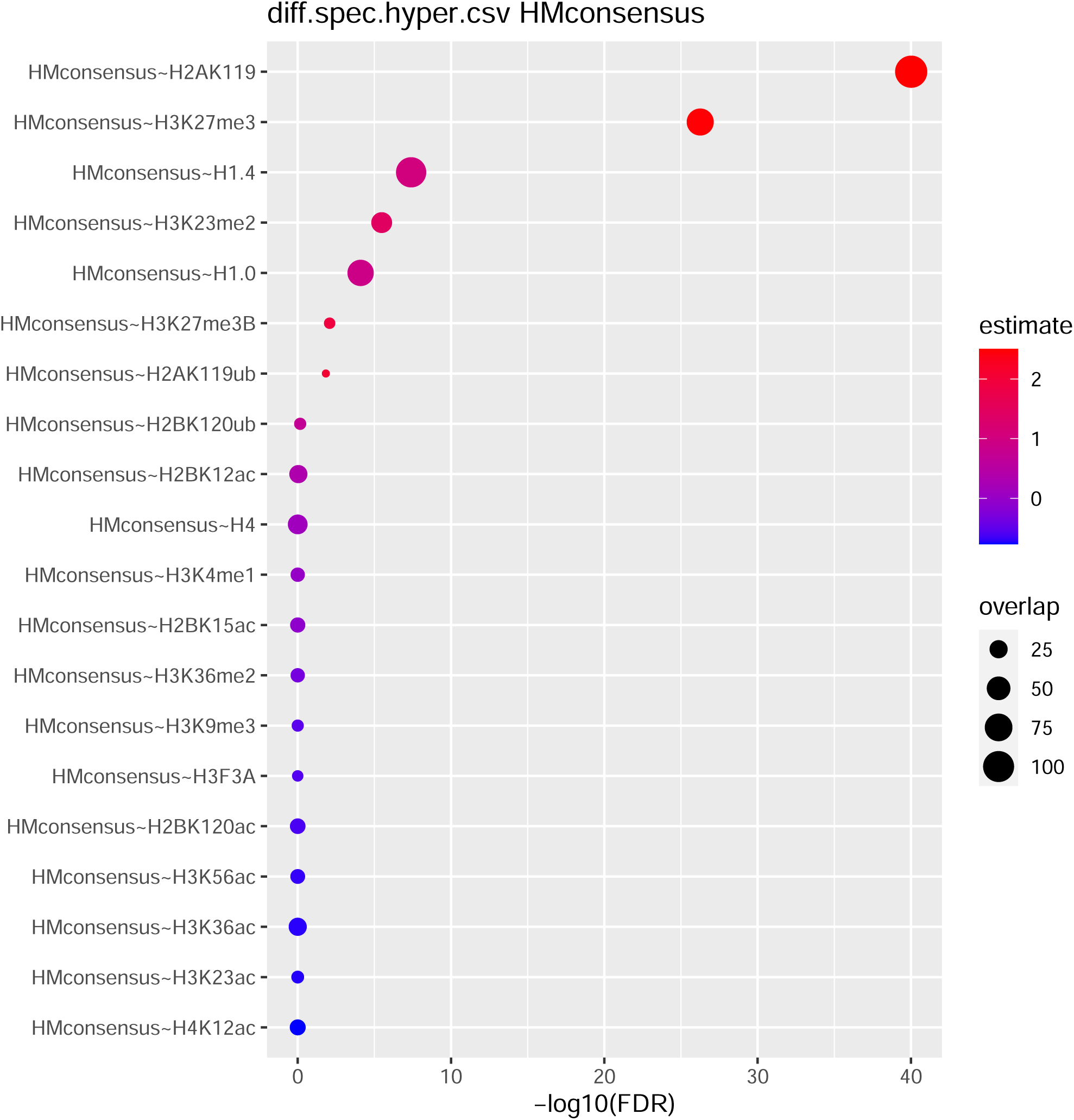
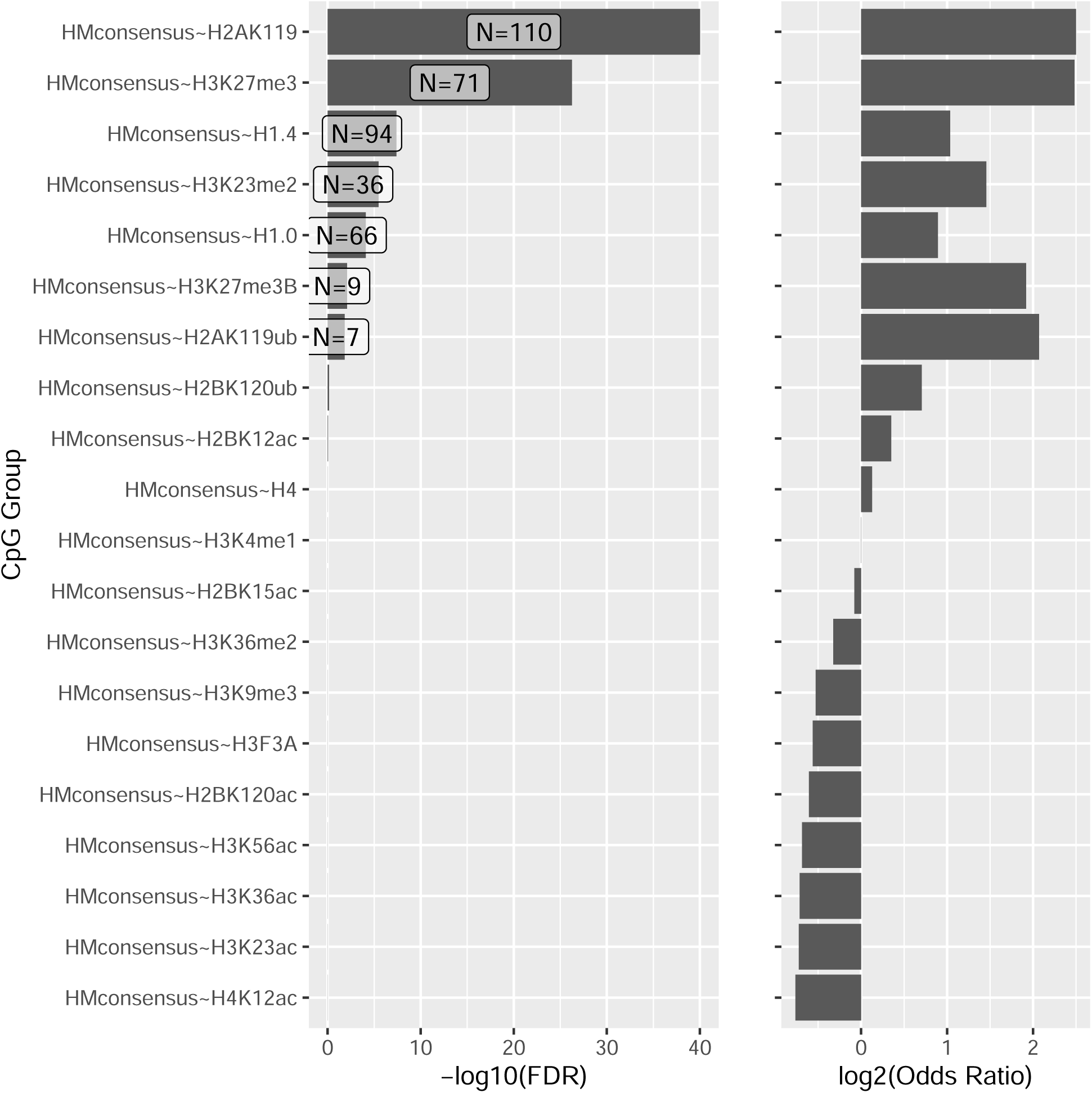
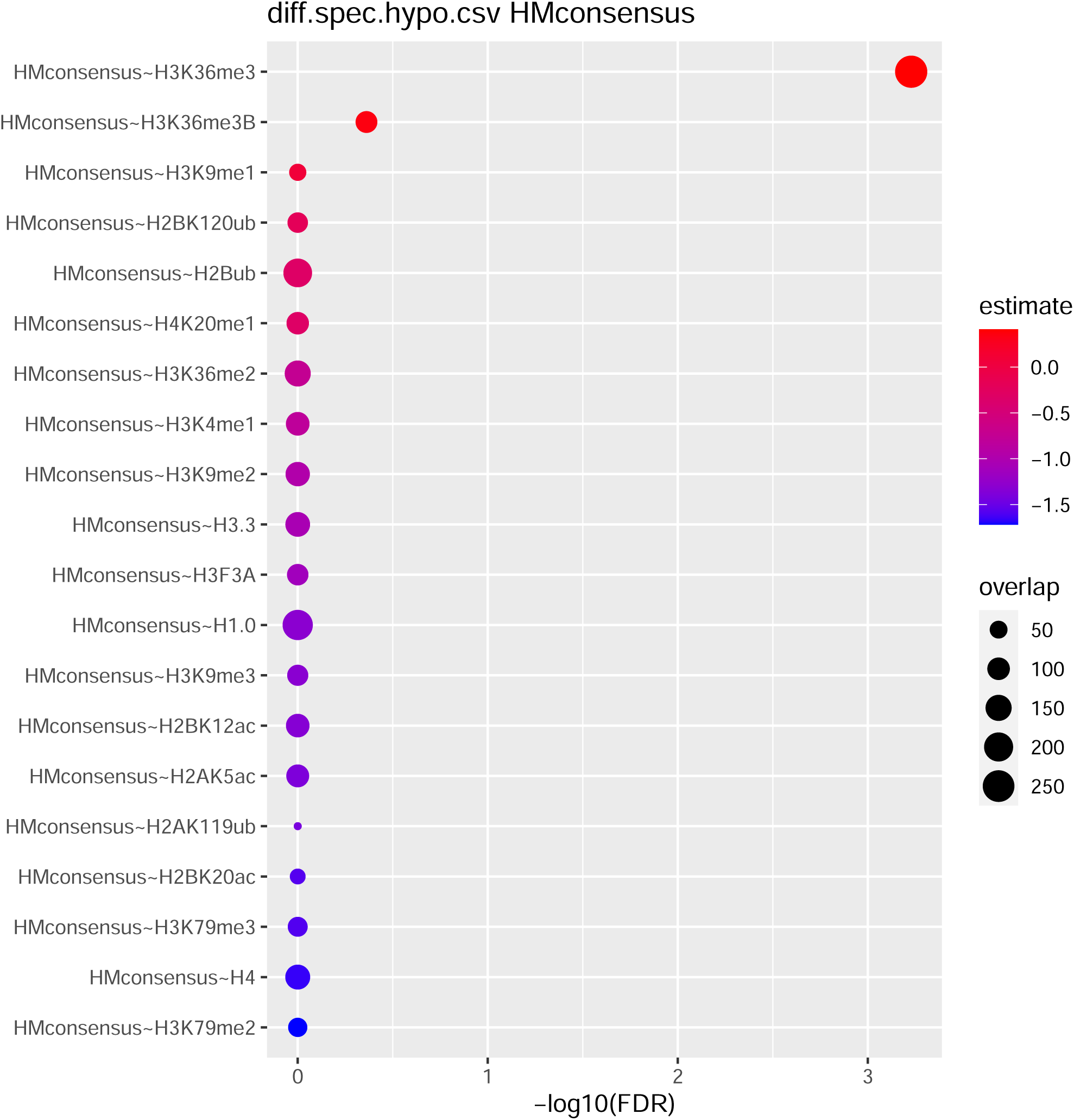
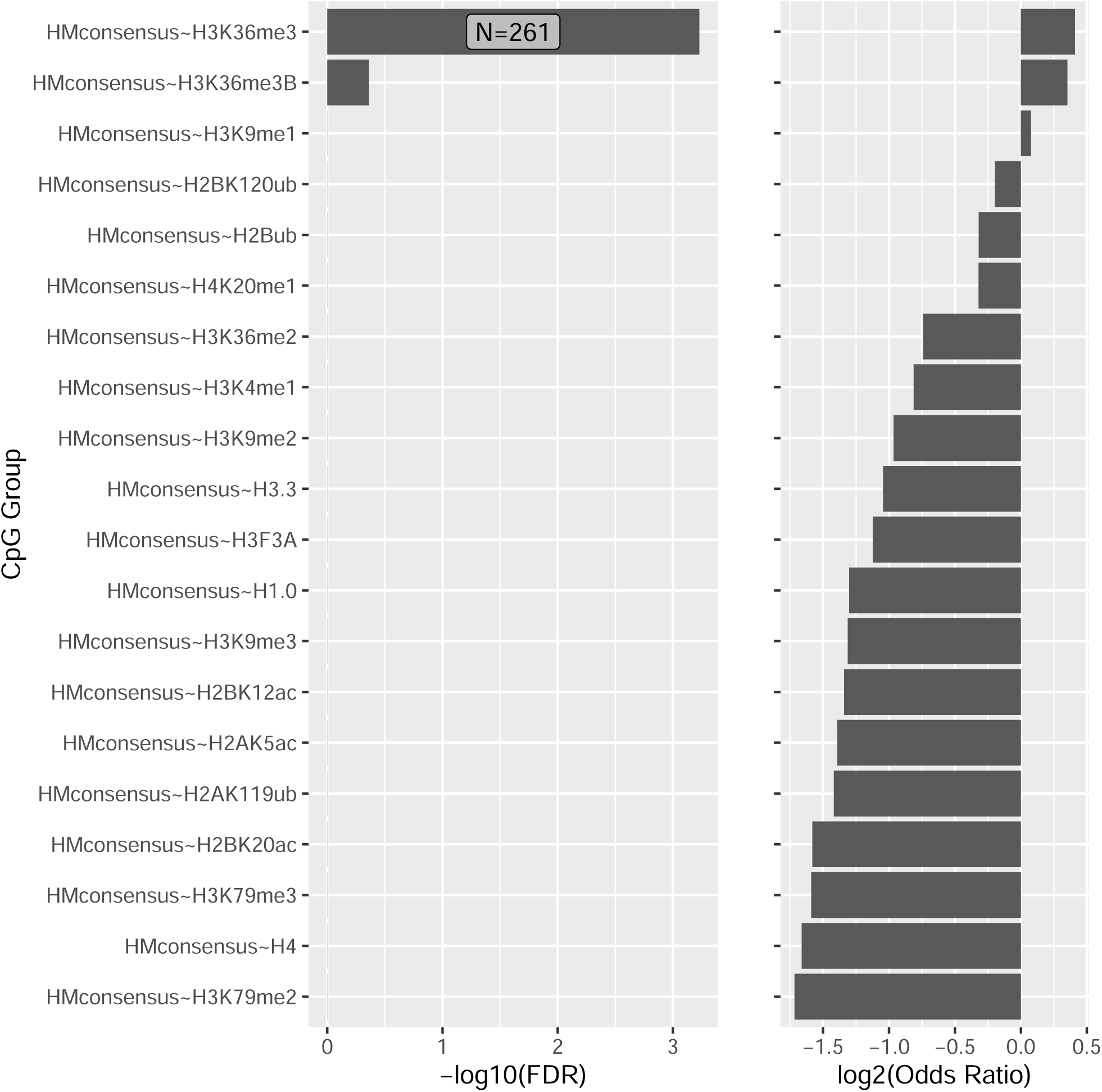
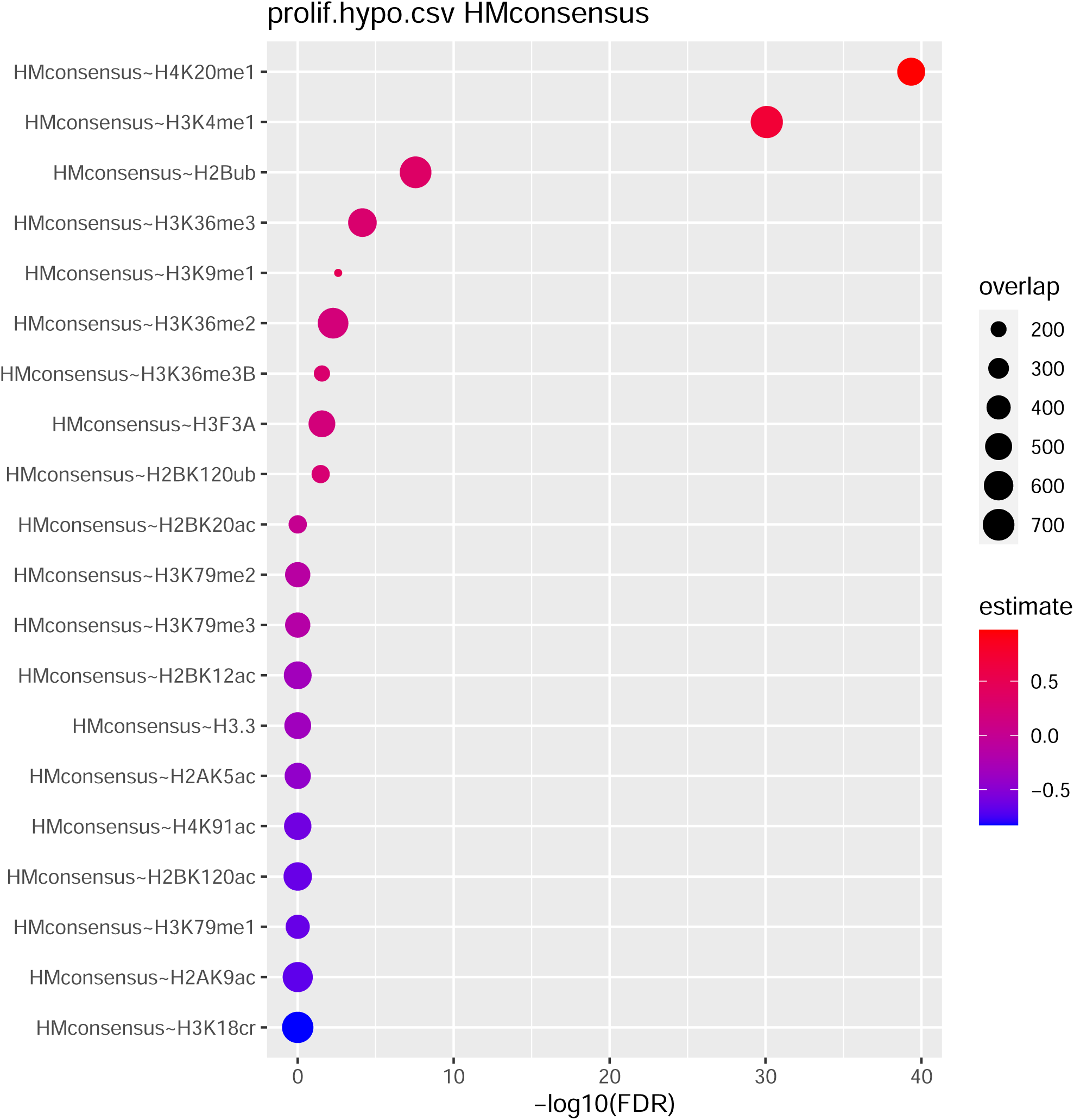
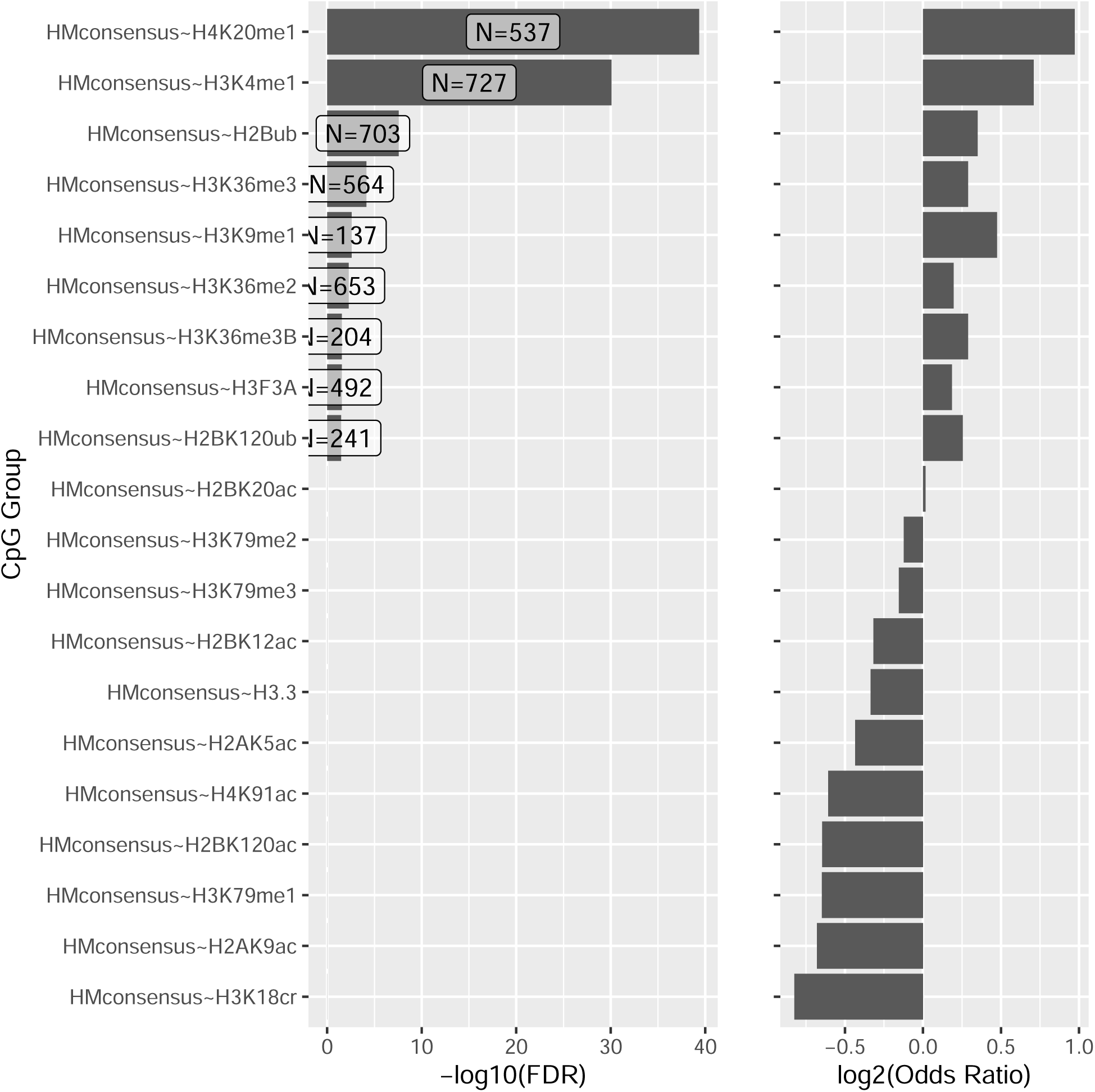
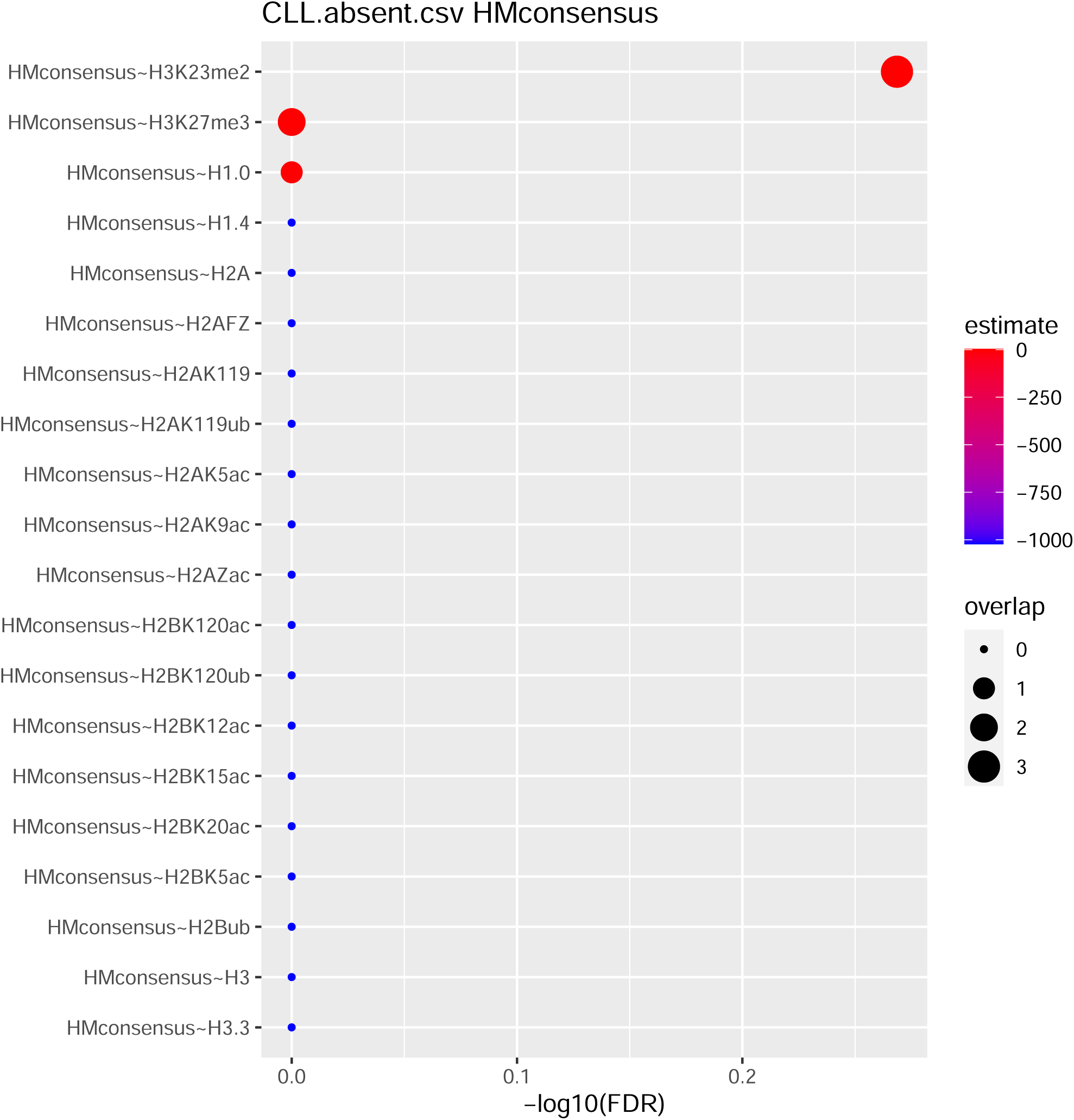
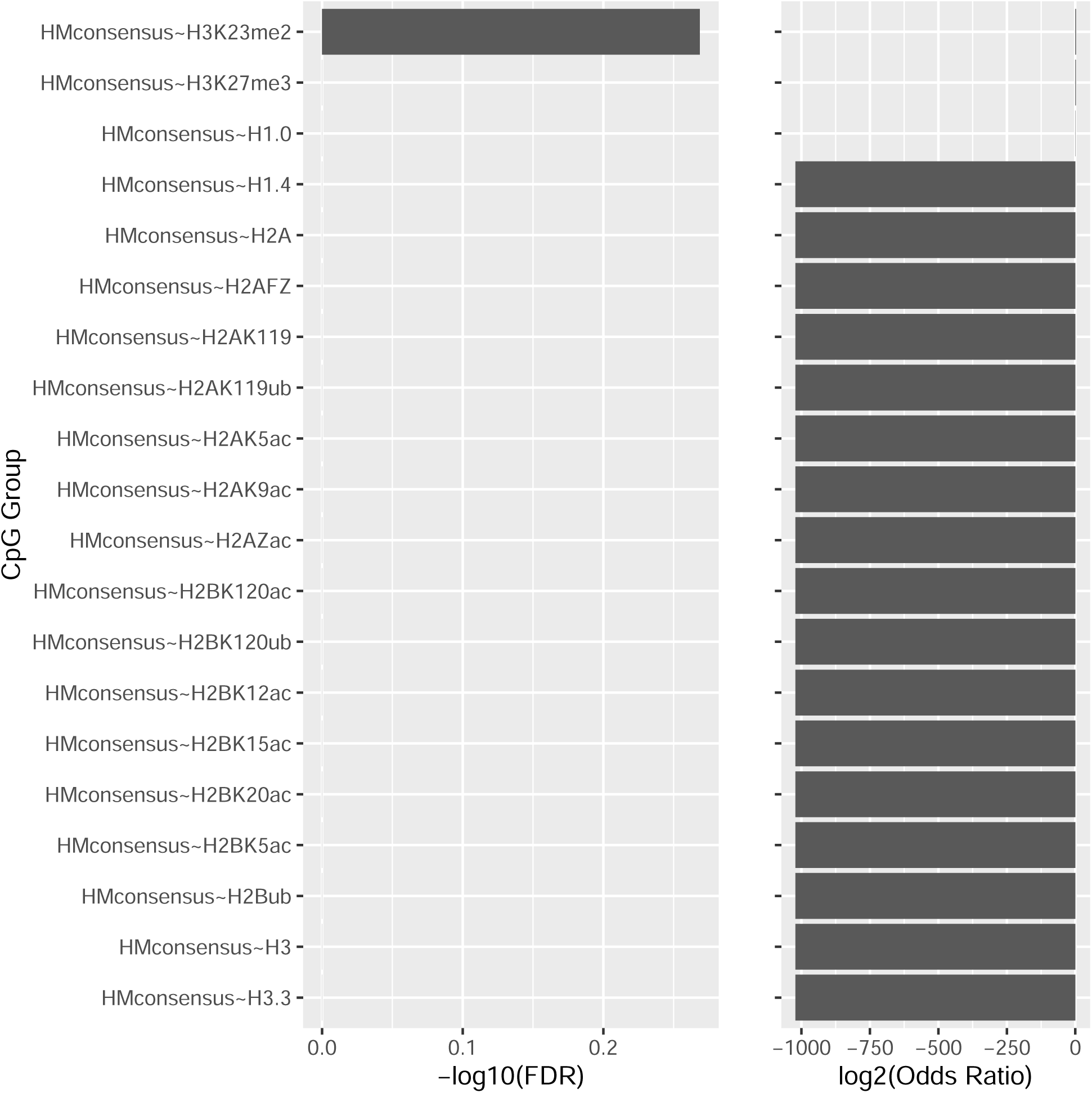
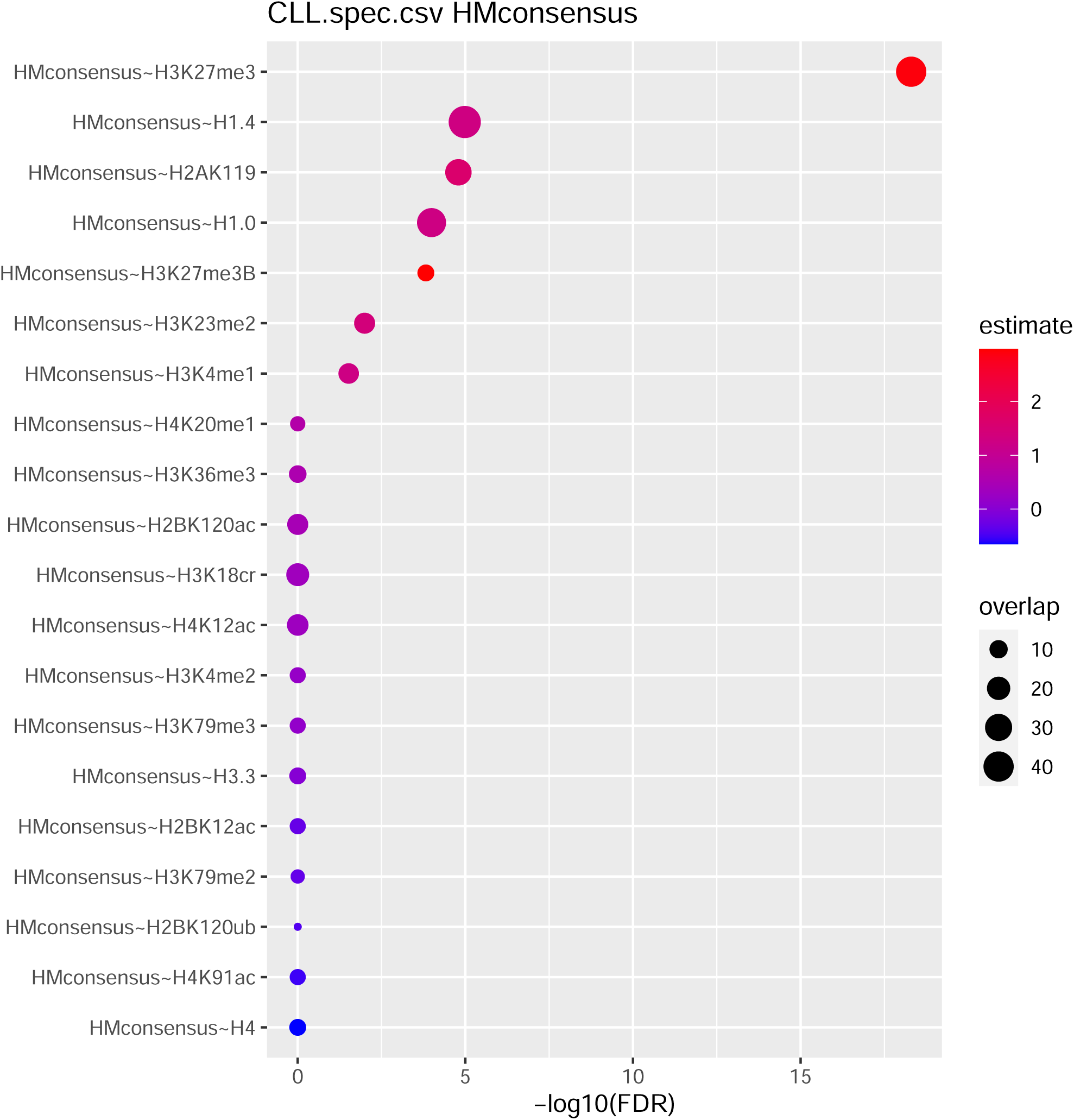
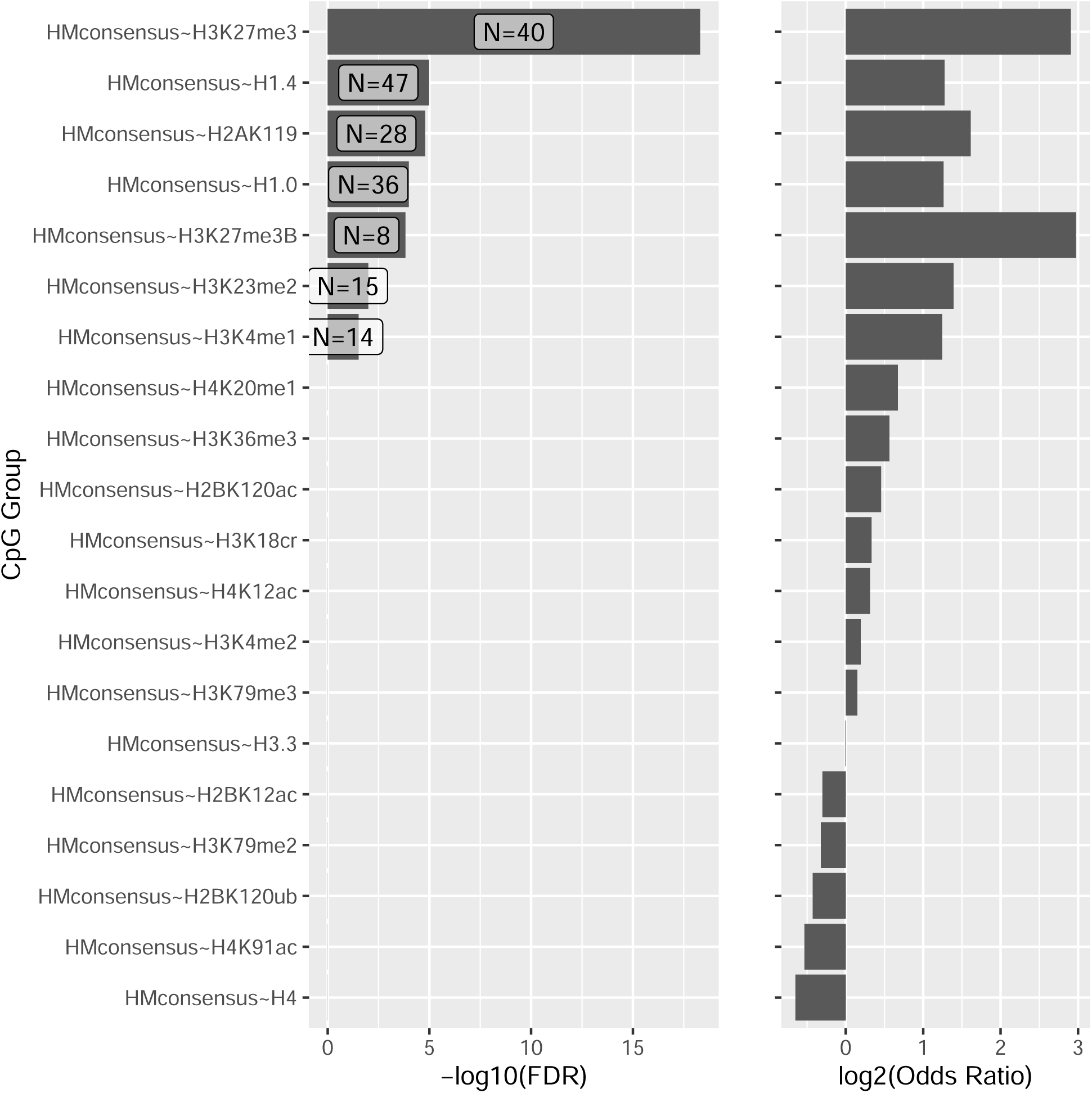
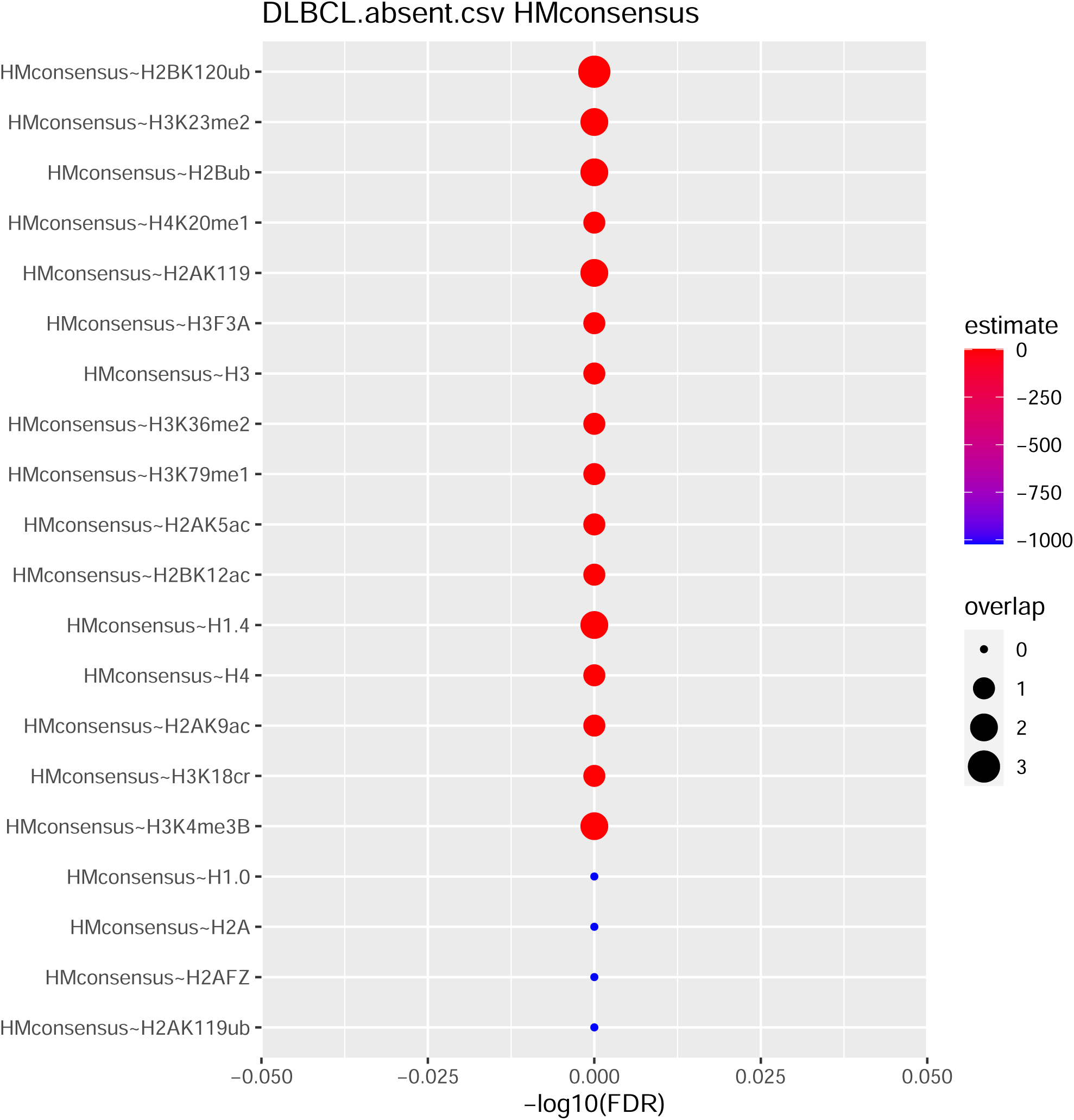
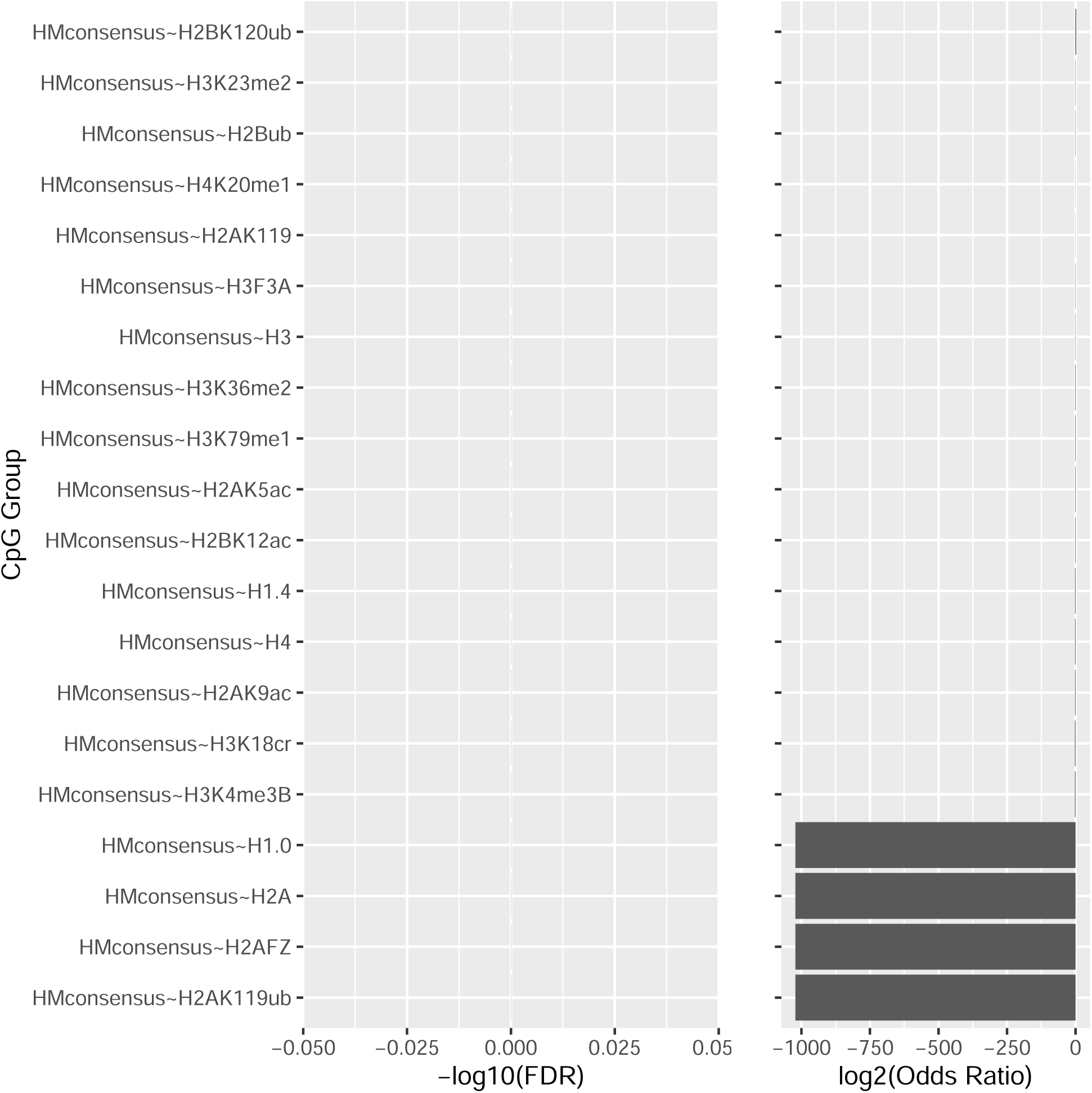
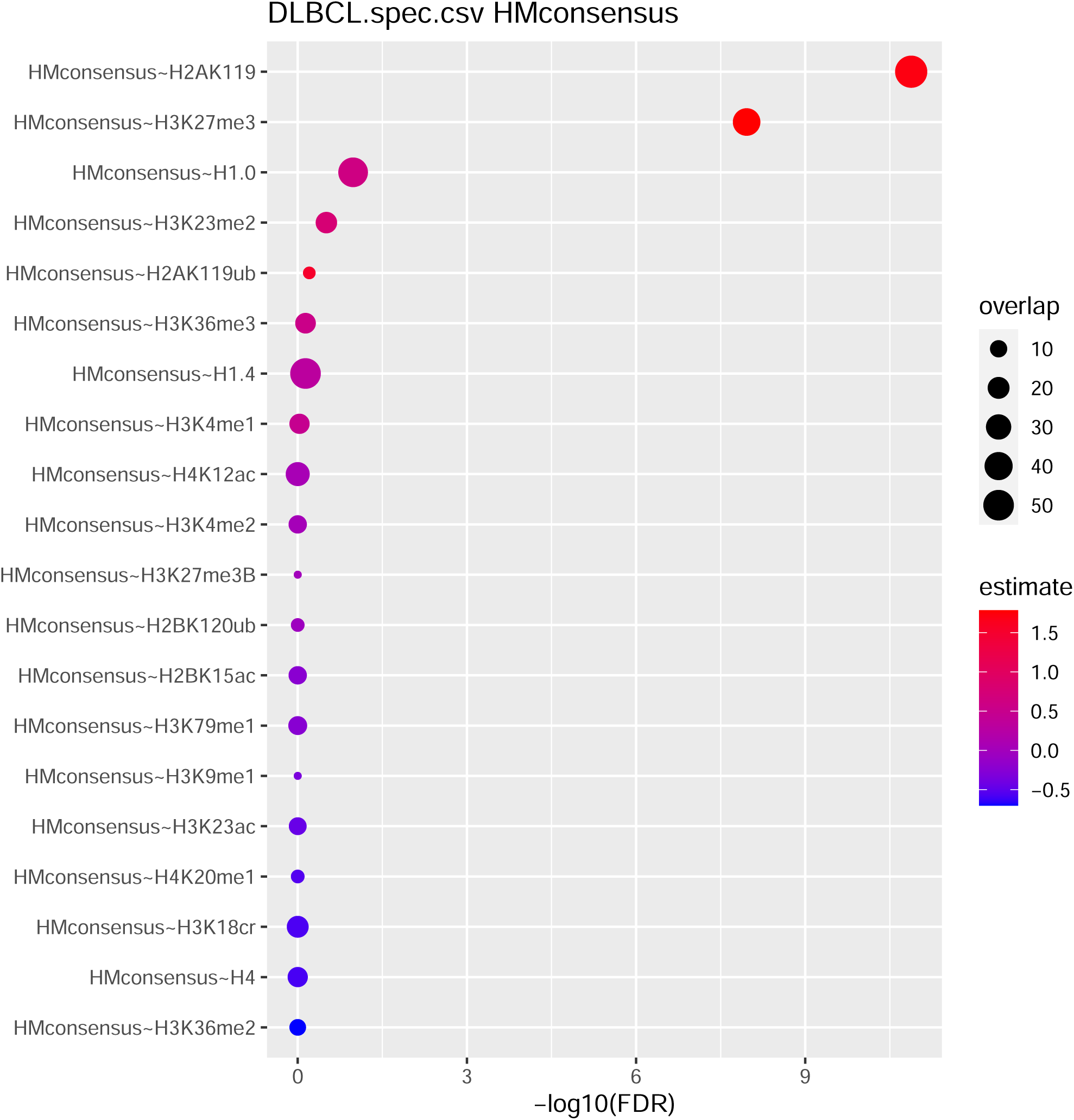
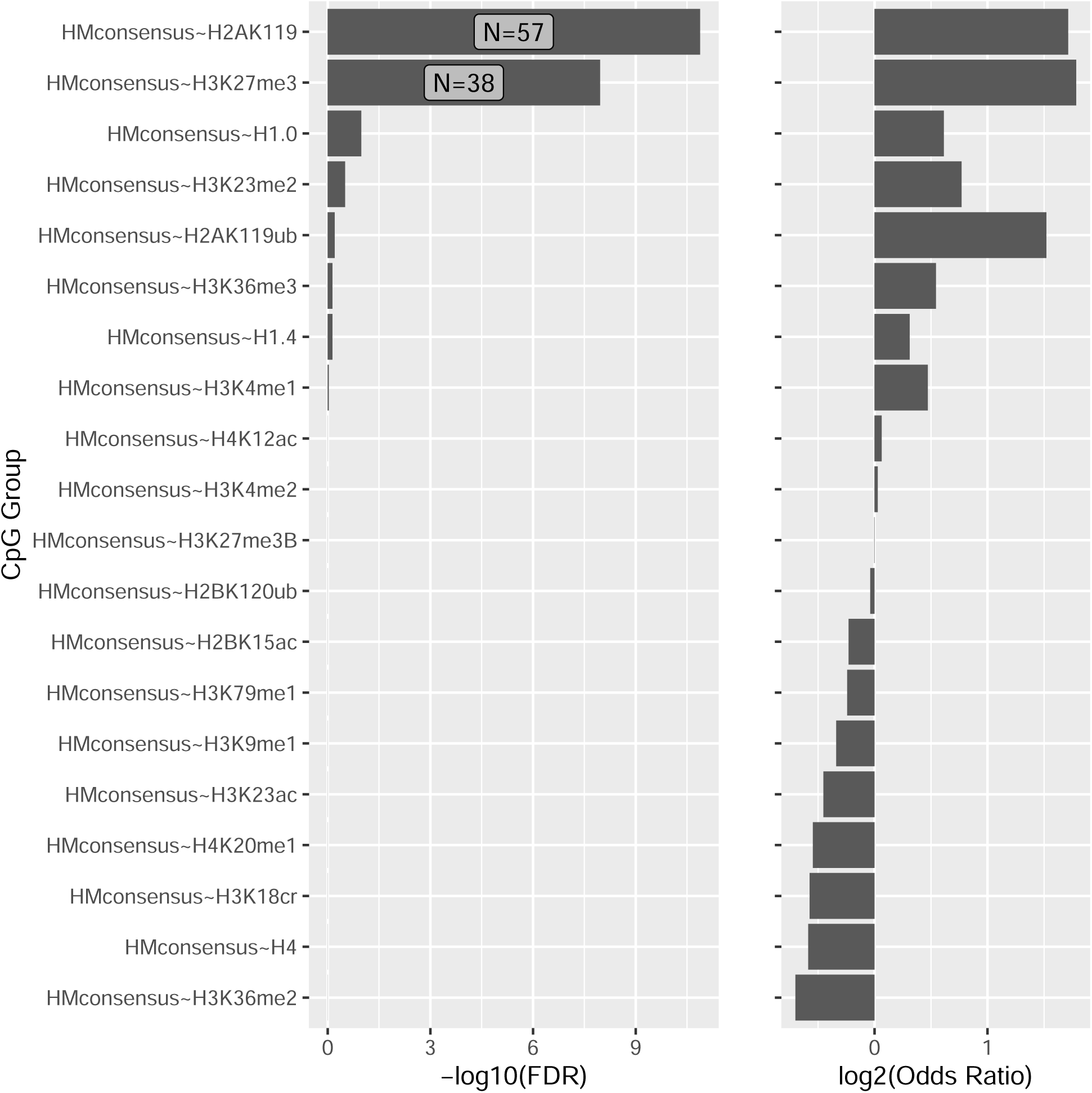
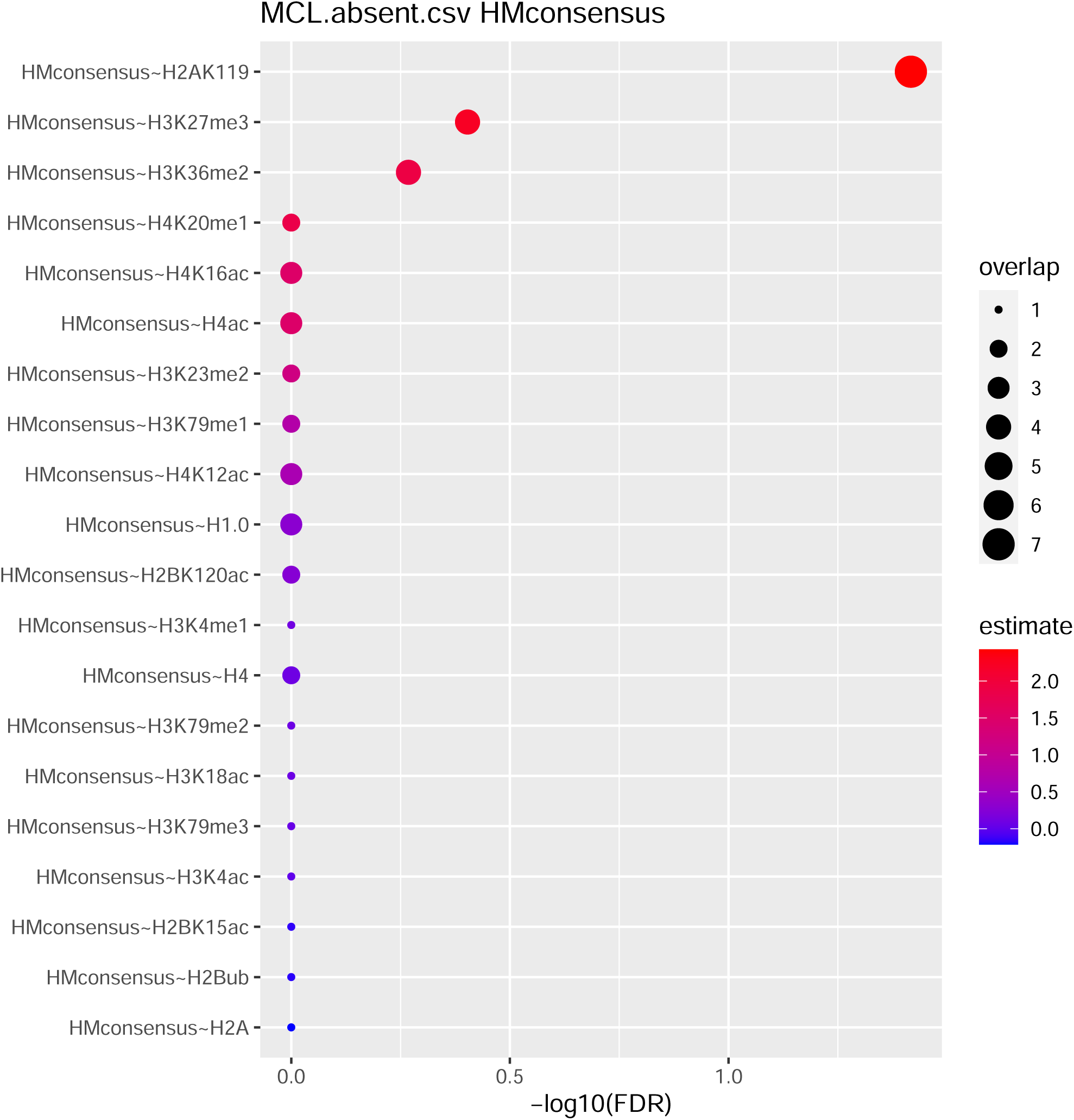
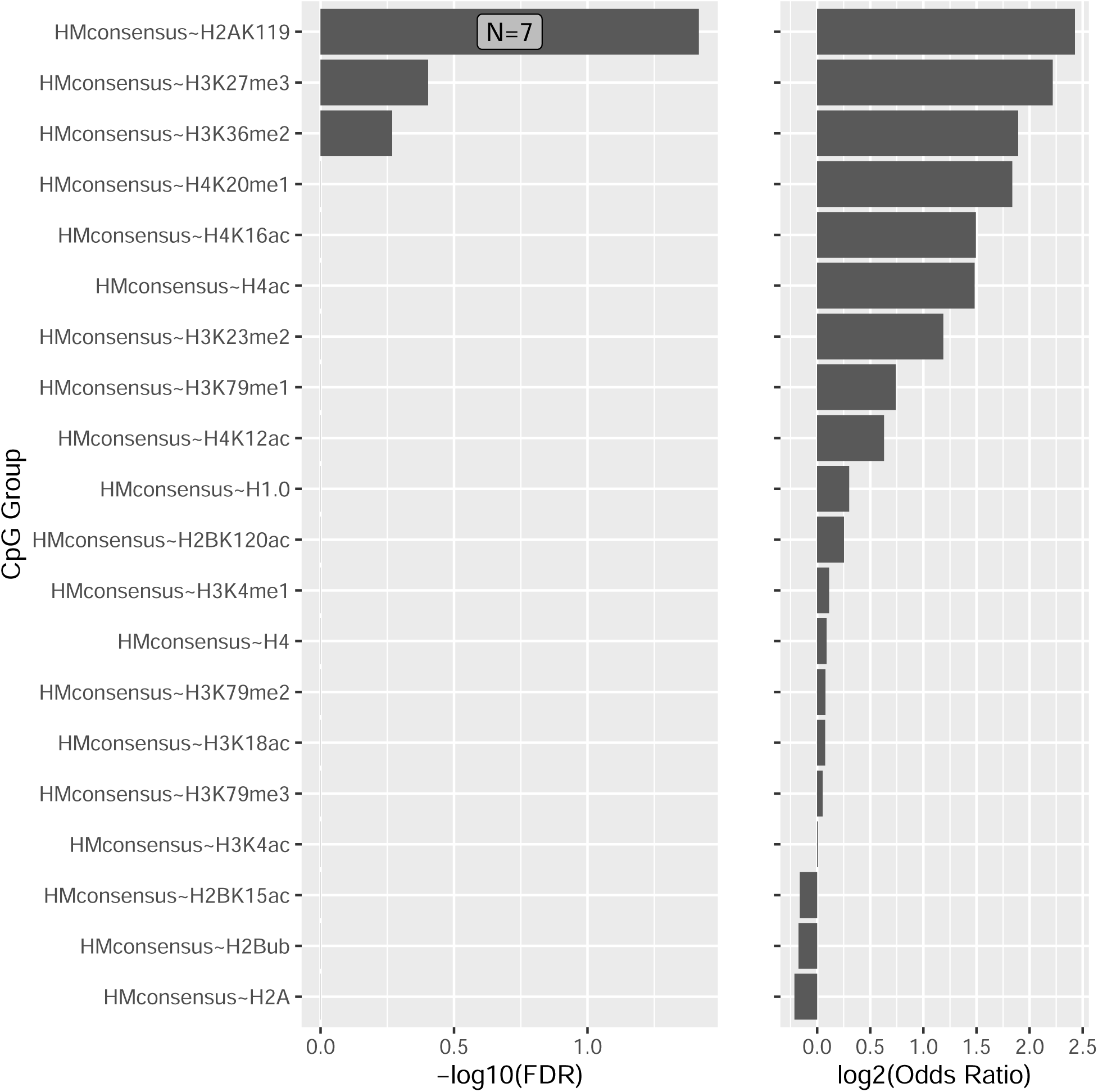
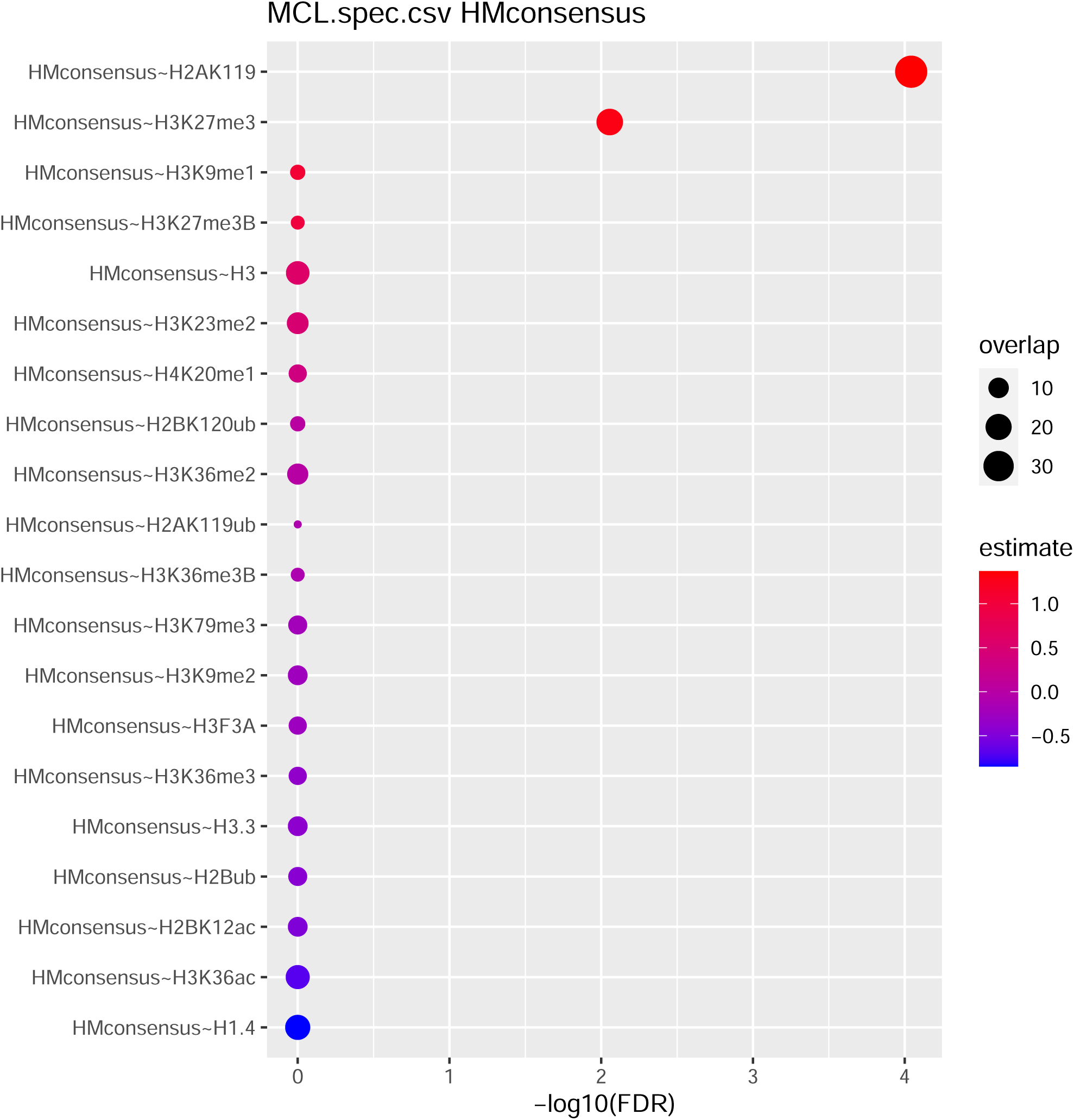
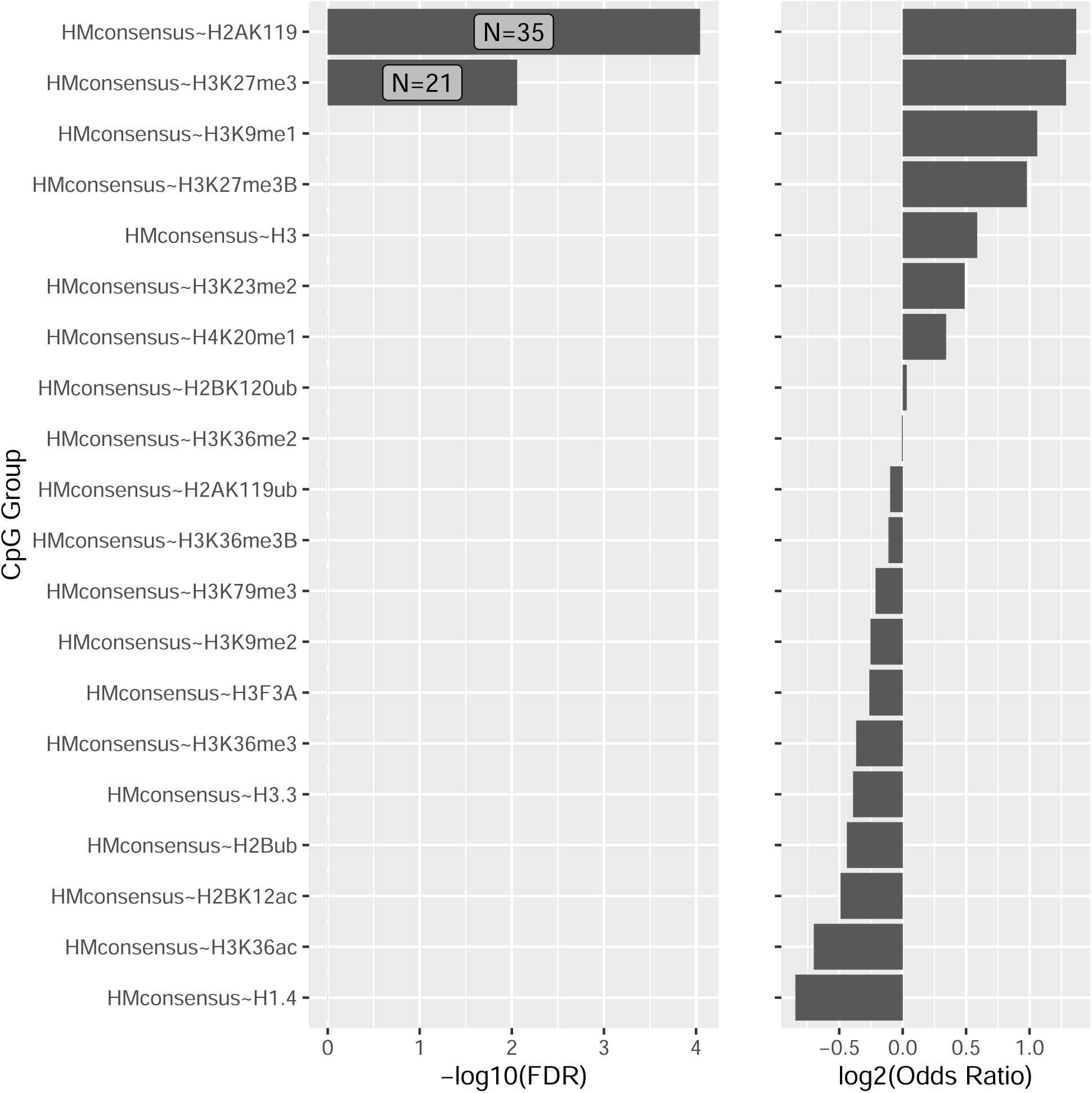
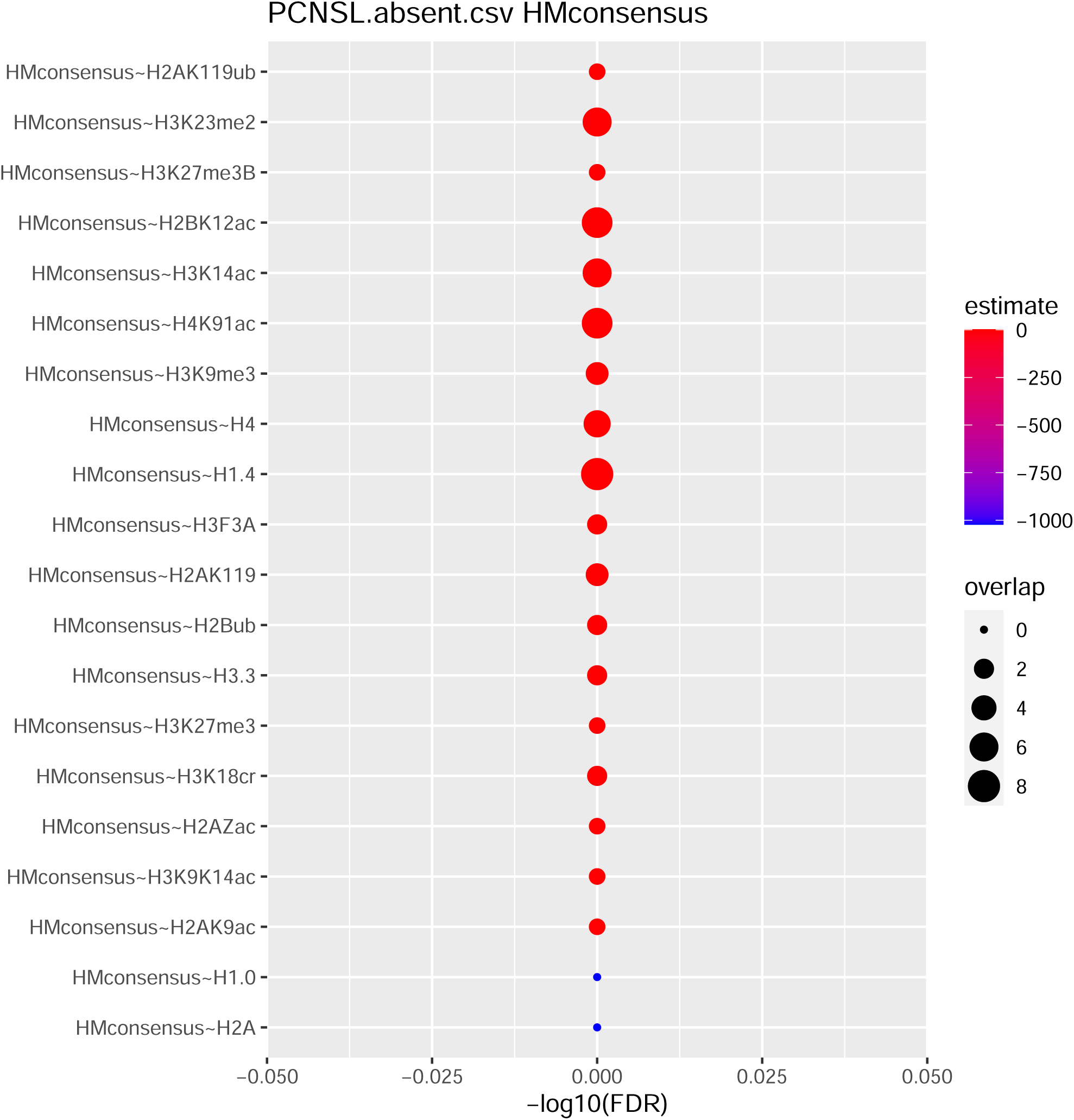
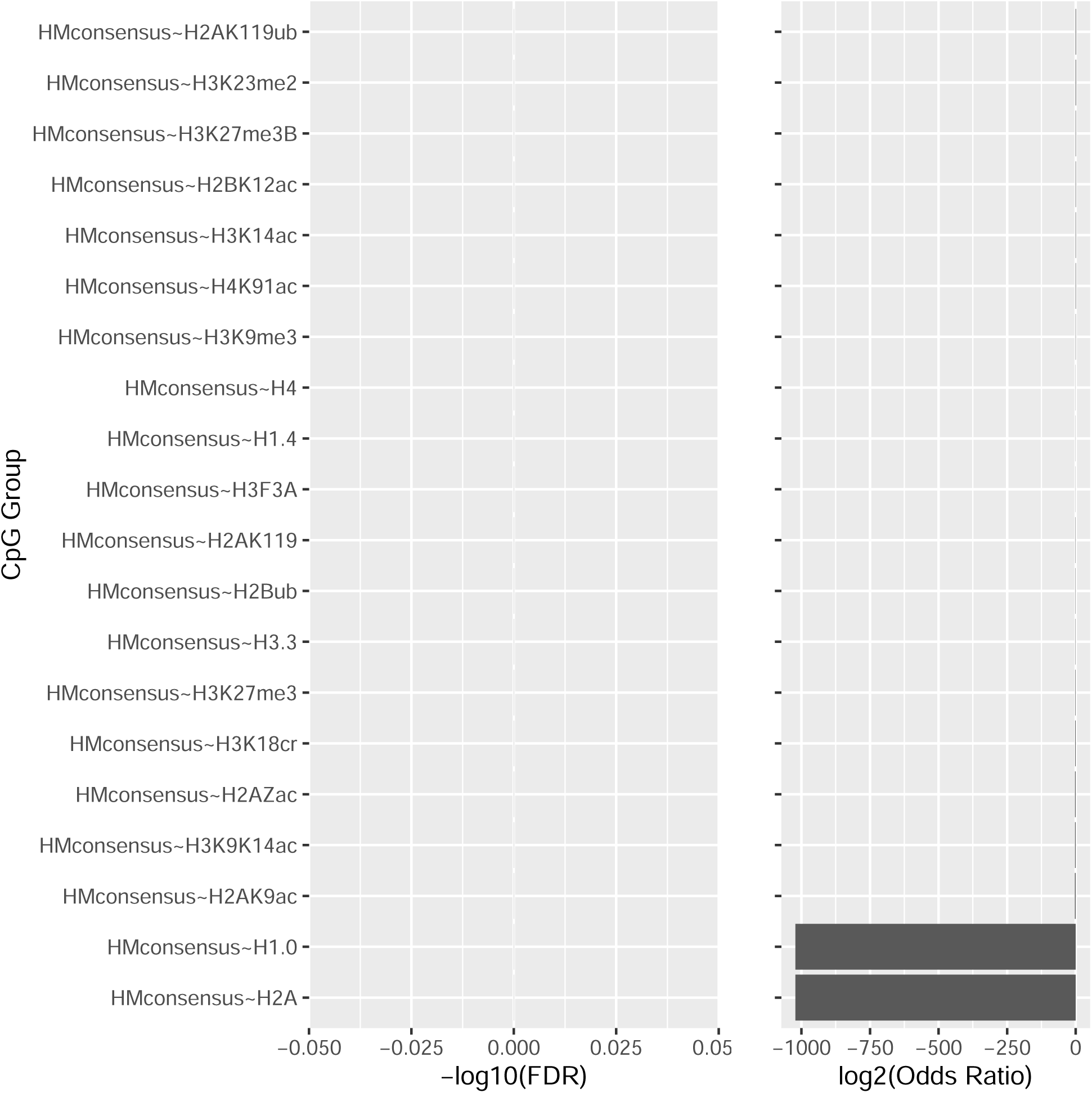
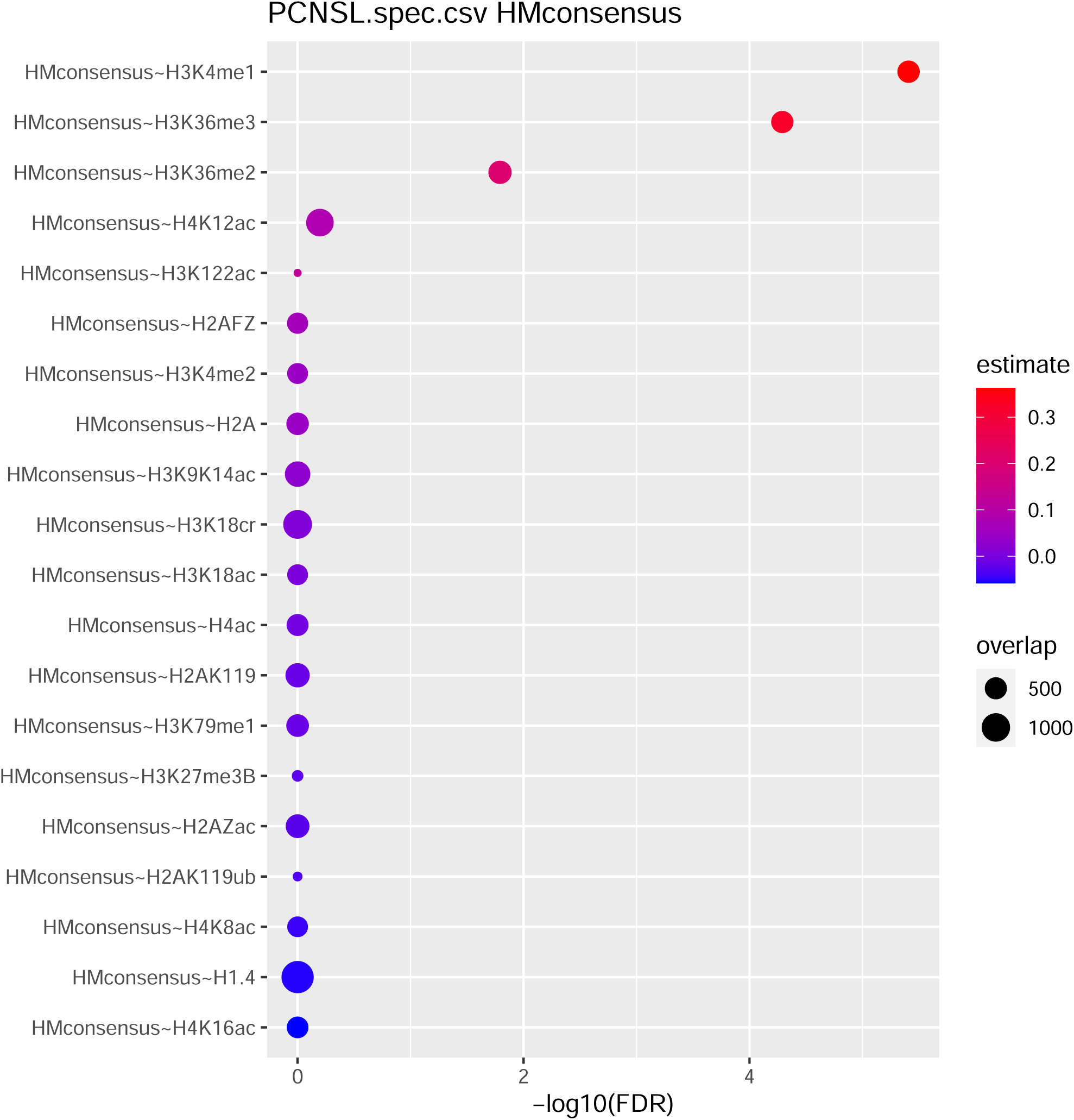
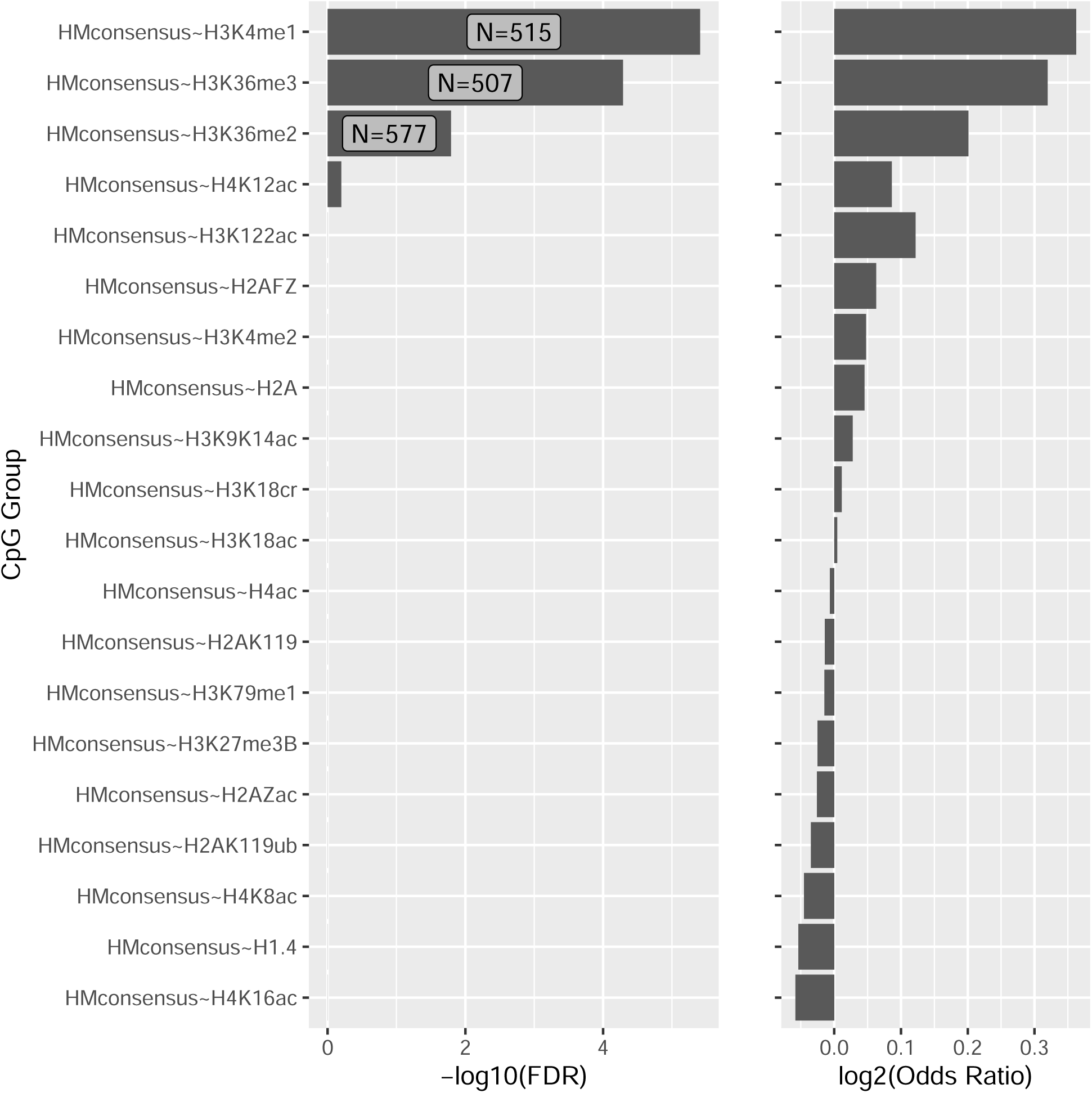
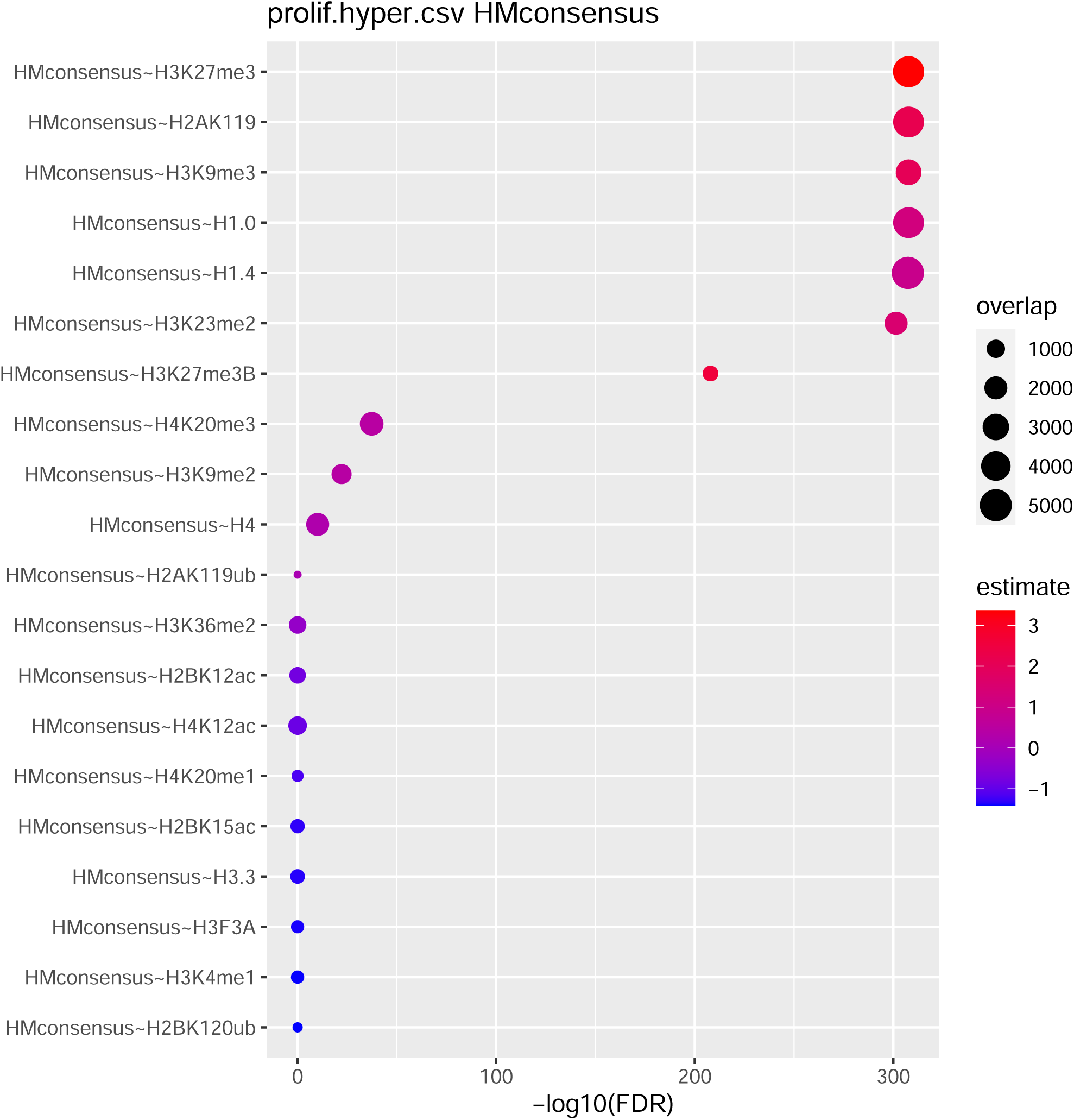
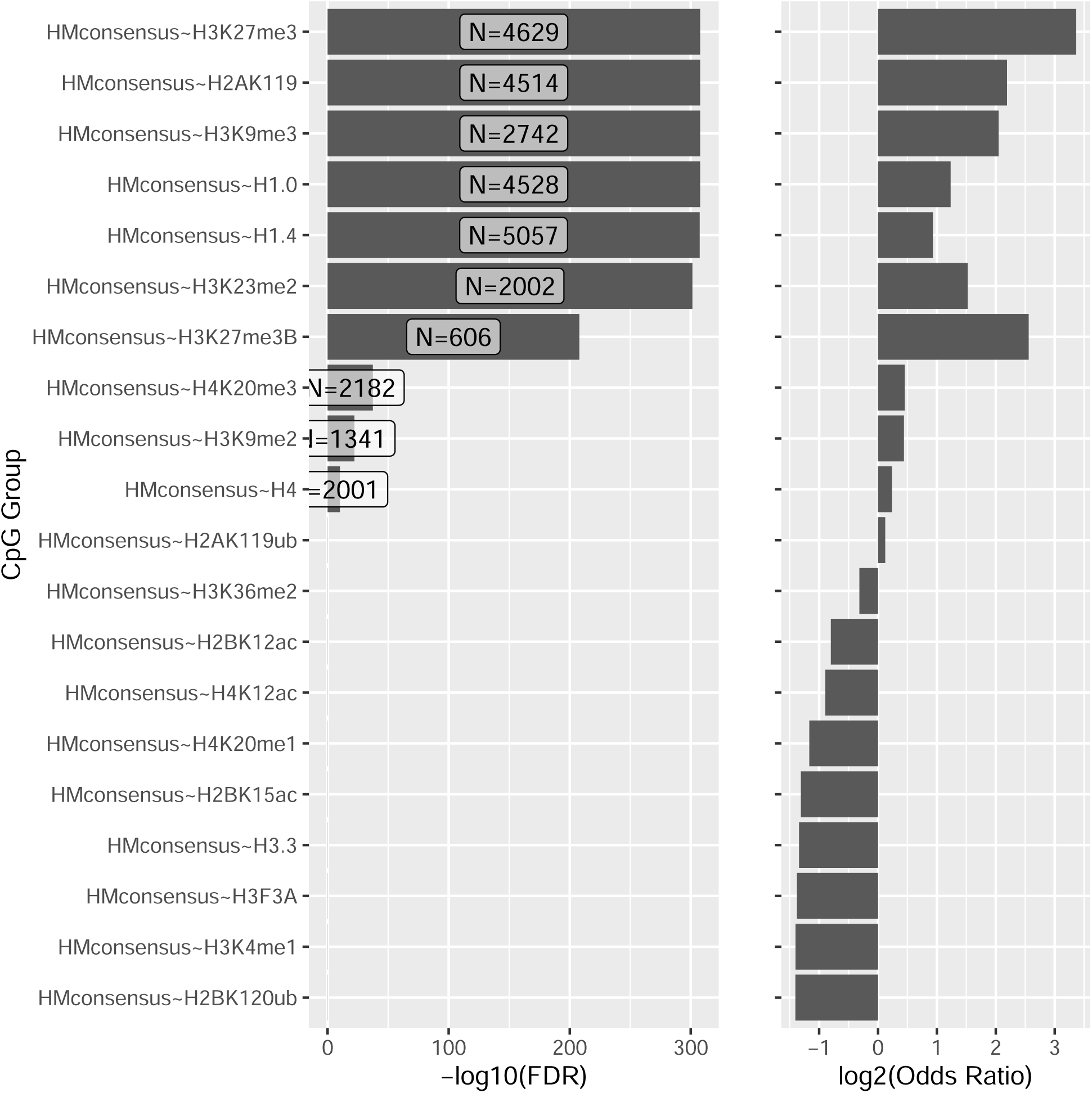

**Supplementary Figure 5.**
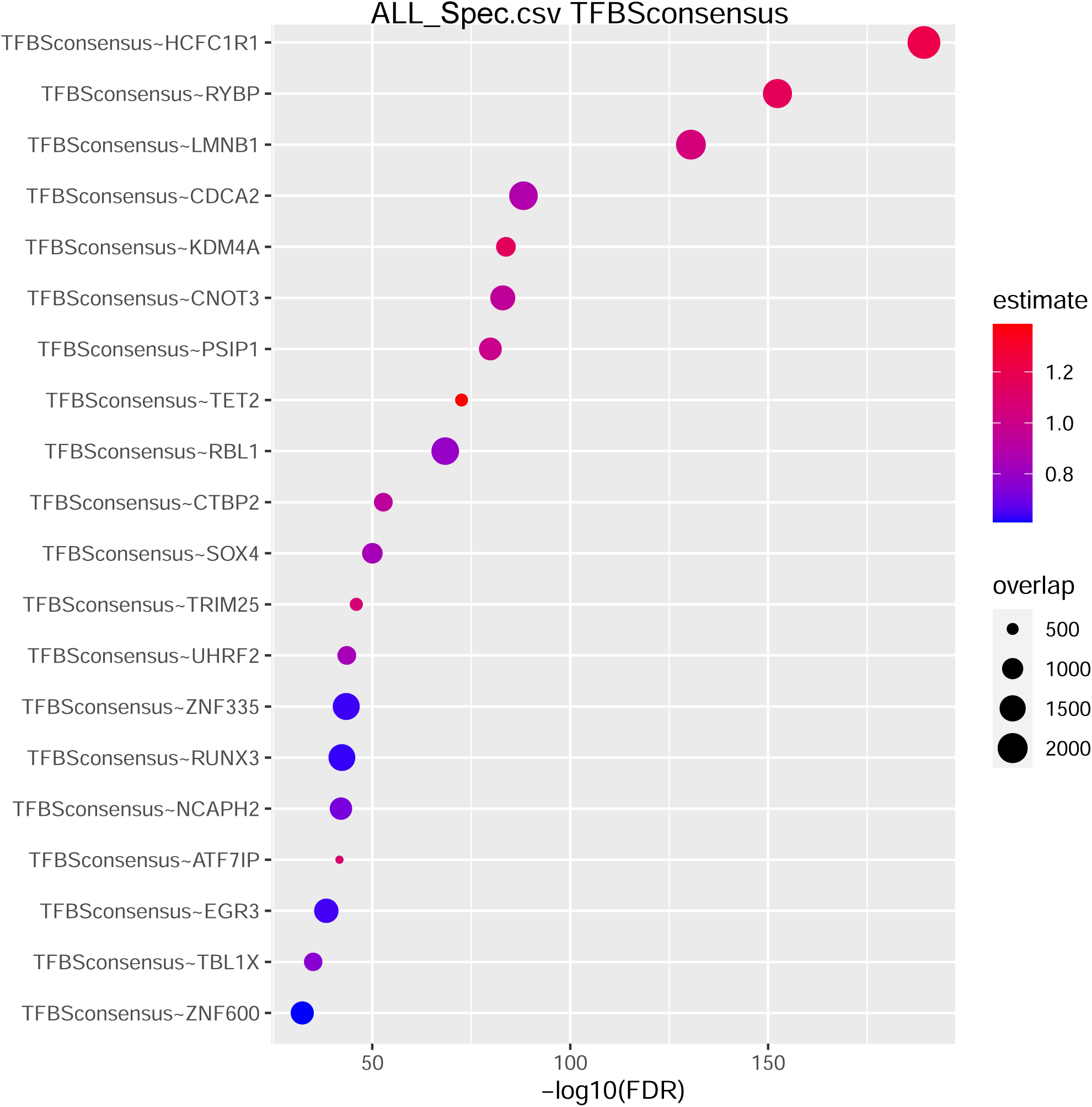
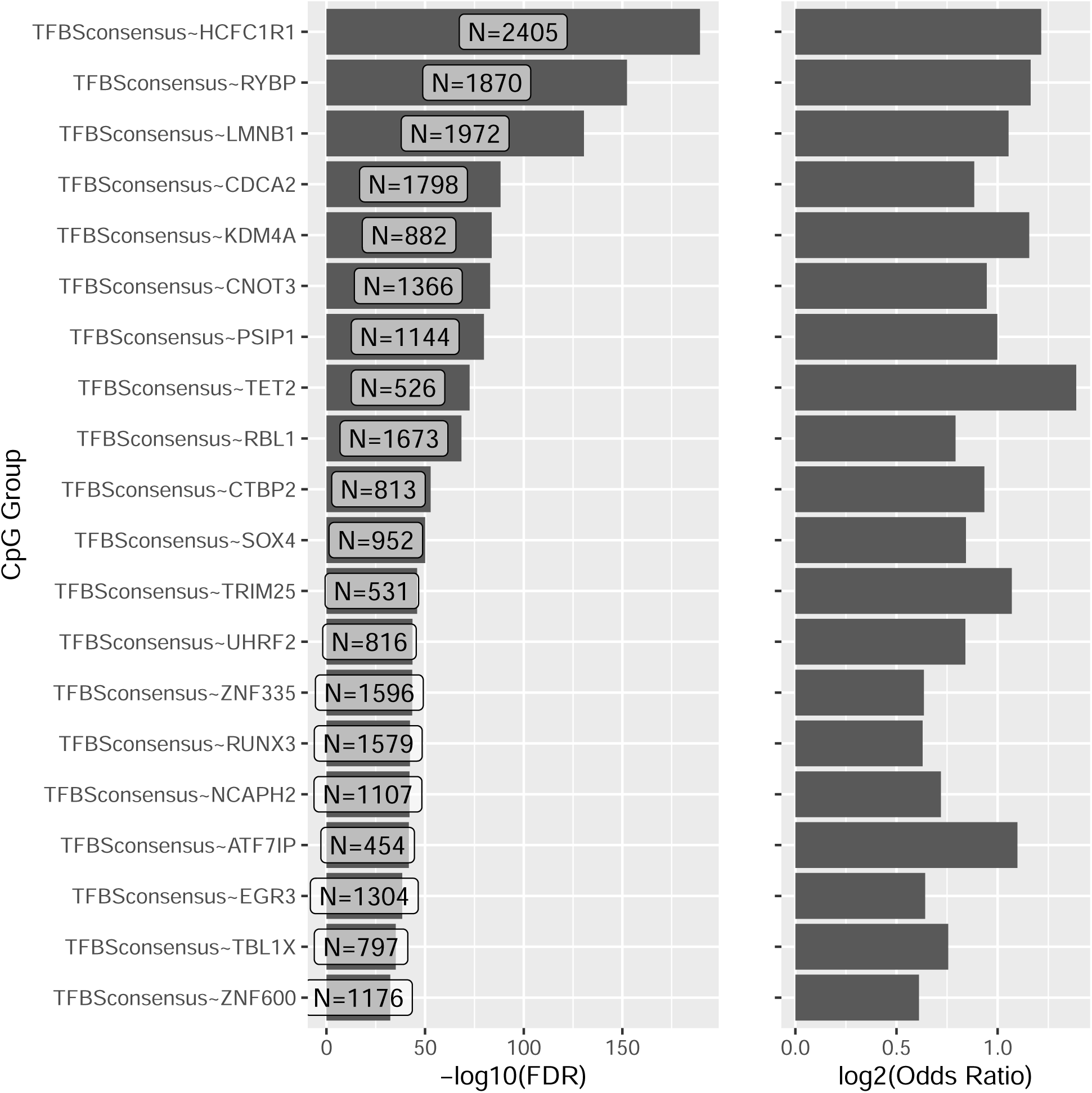
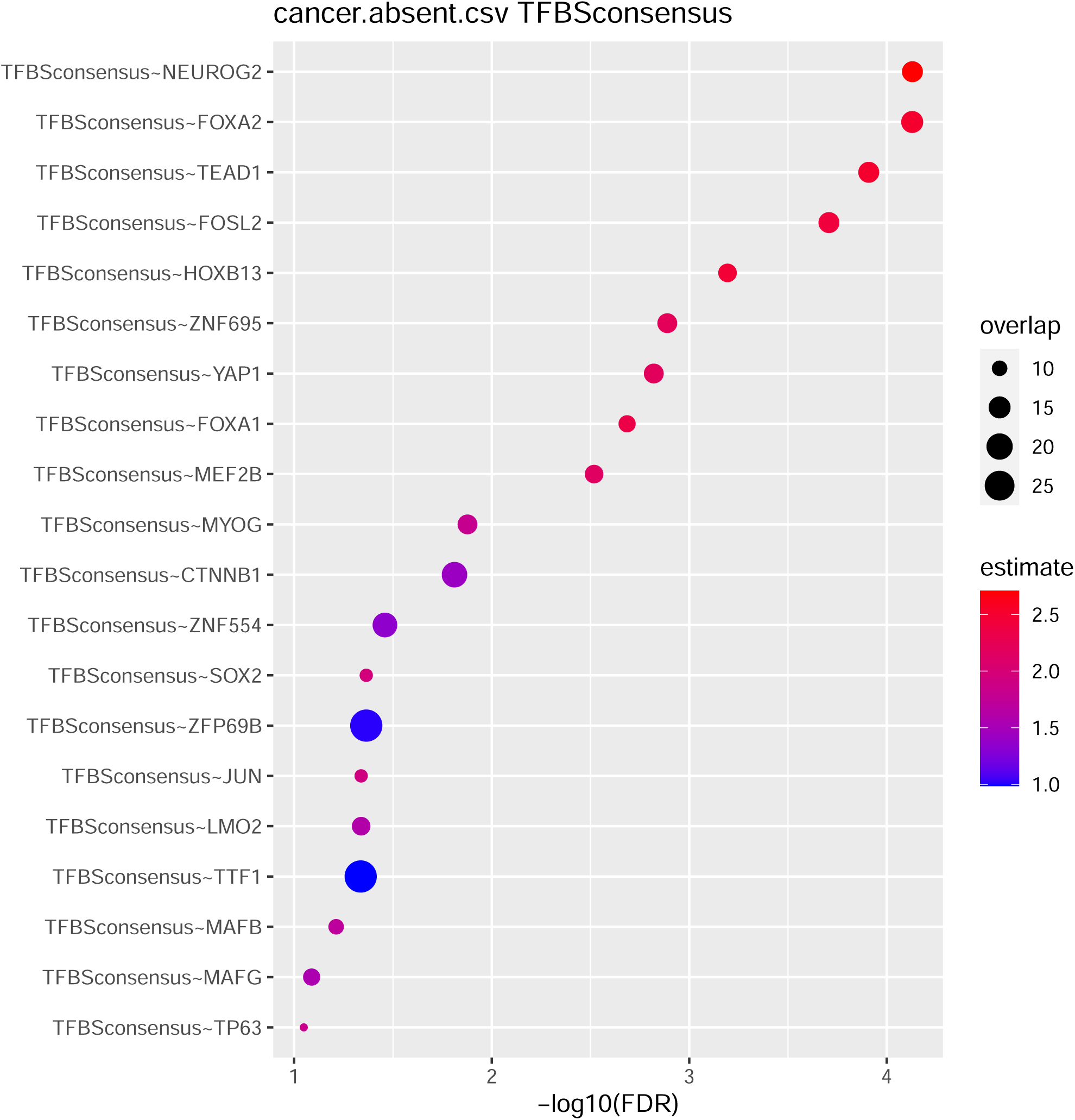
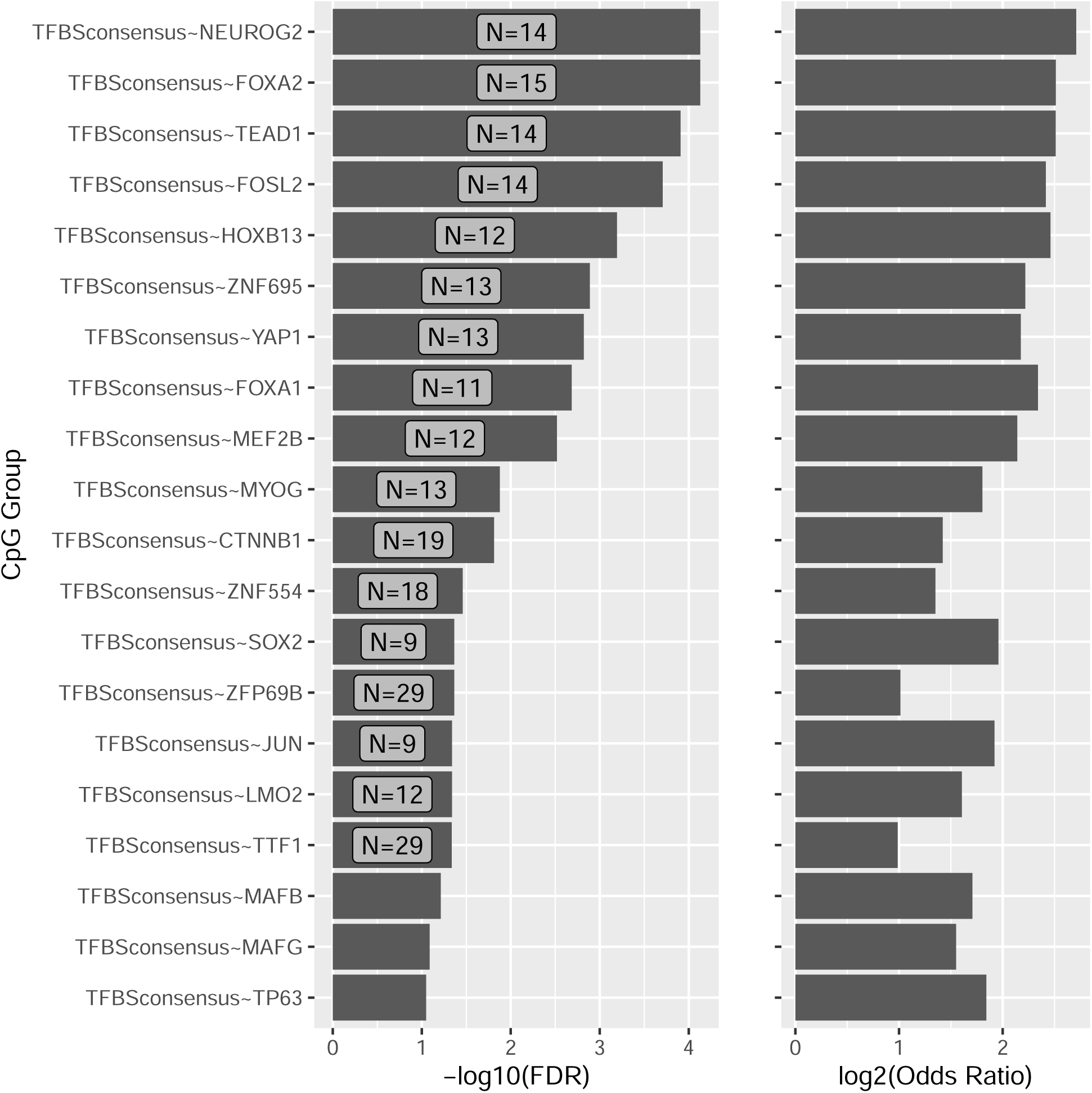
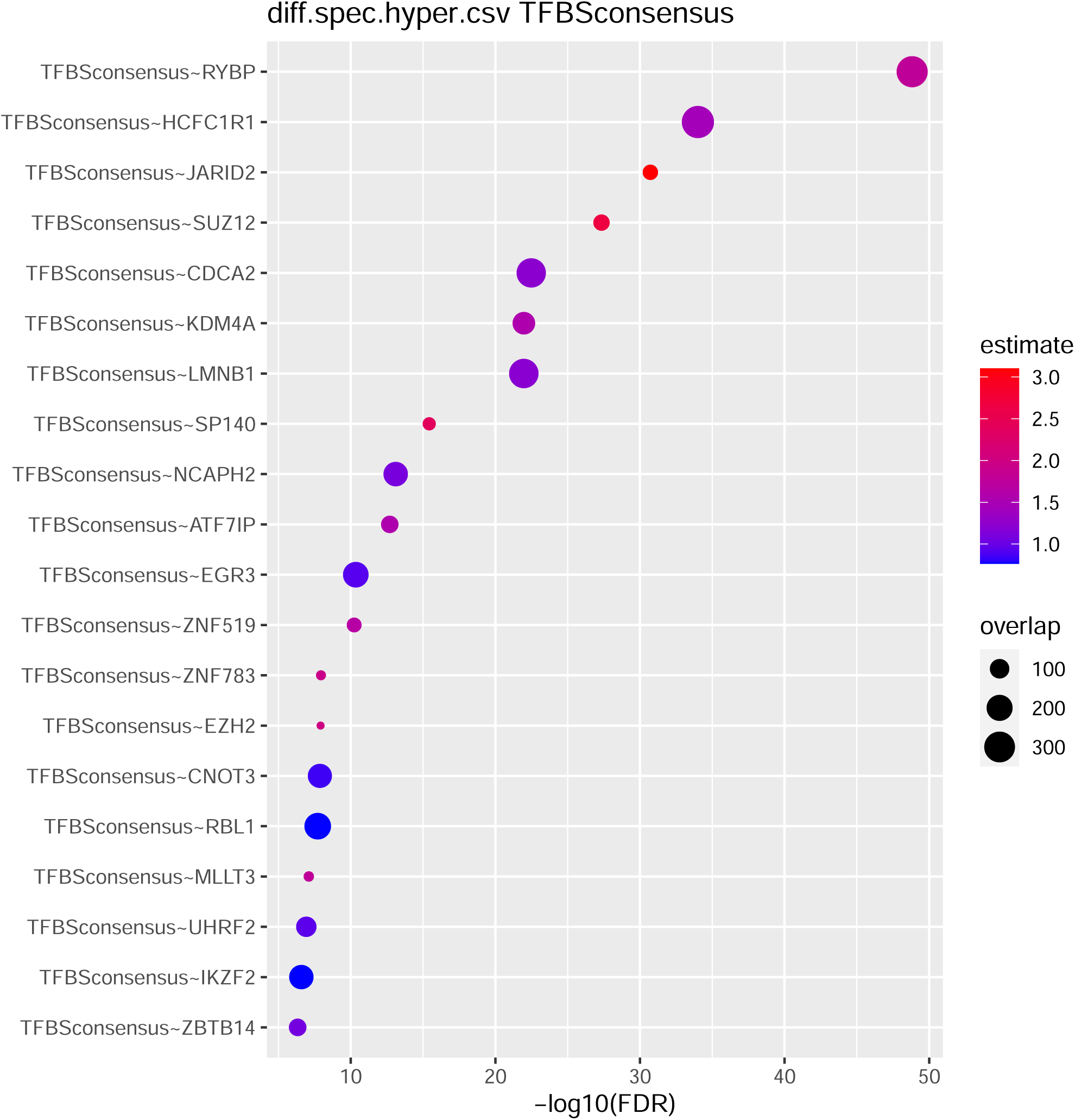
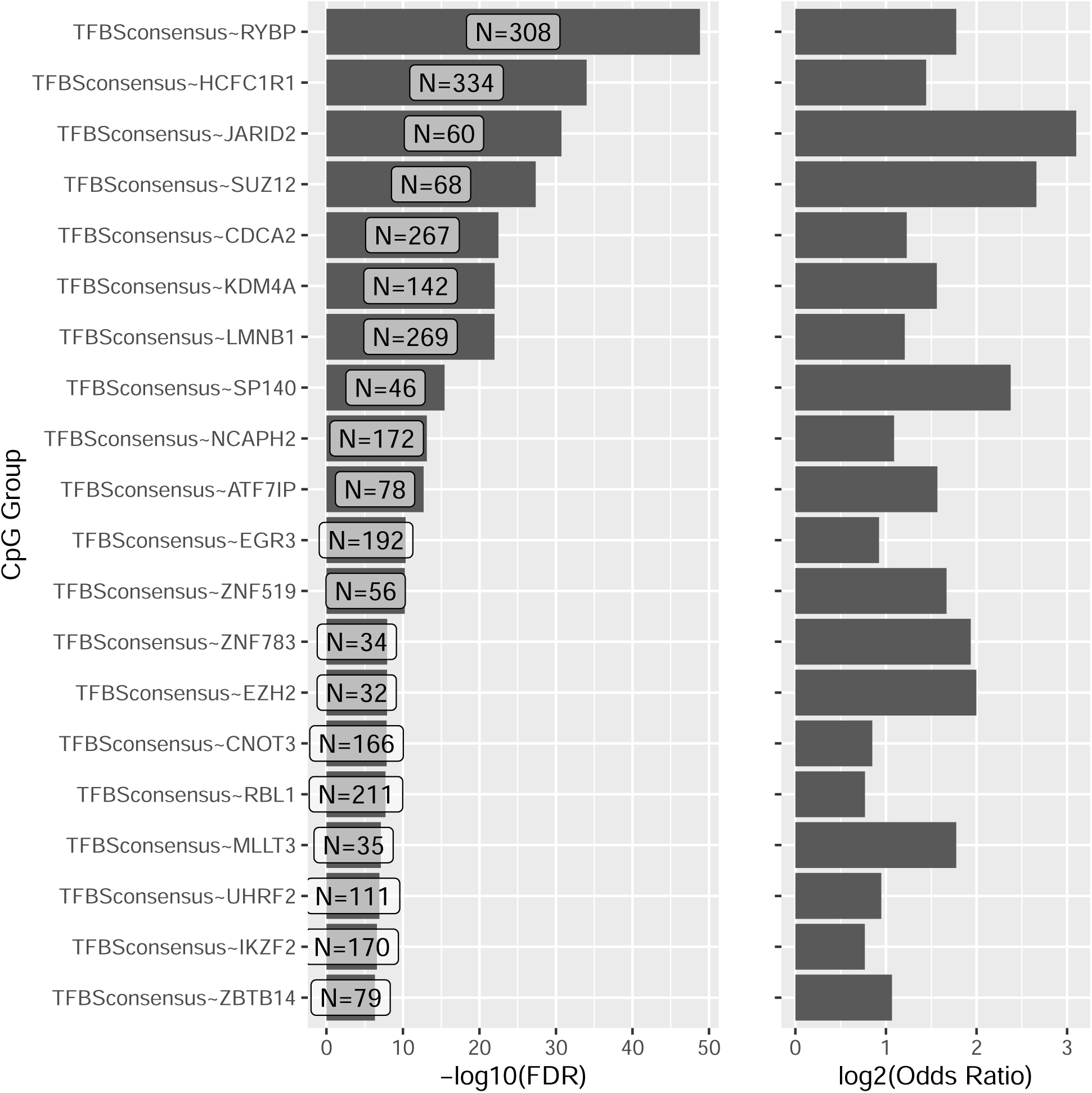
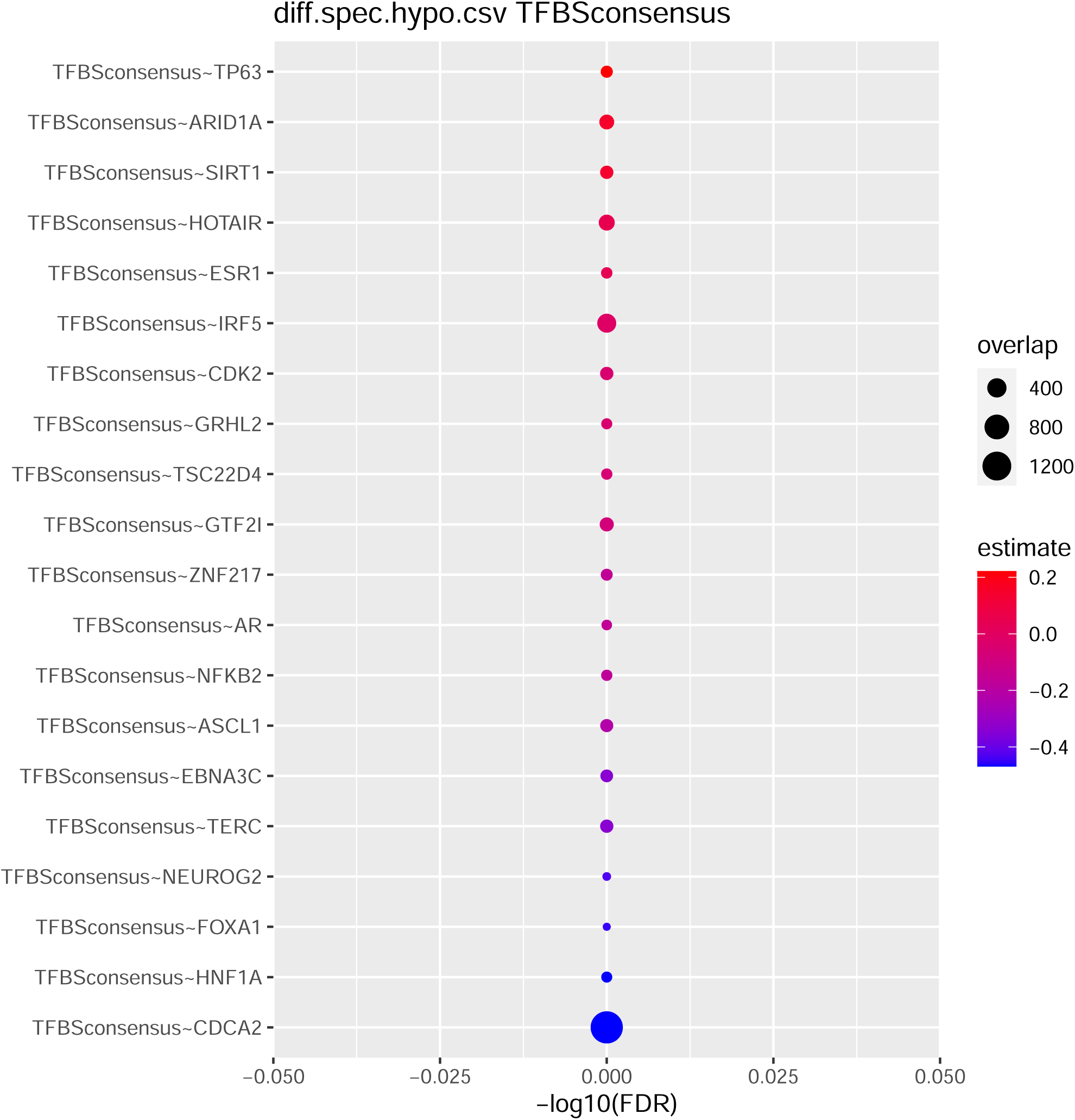
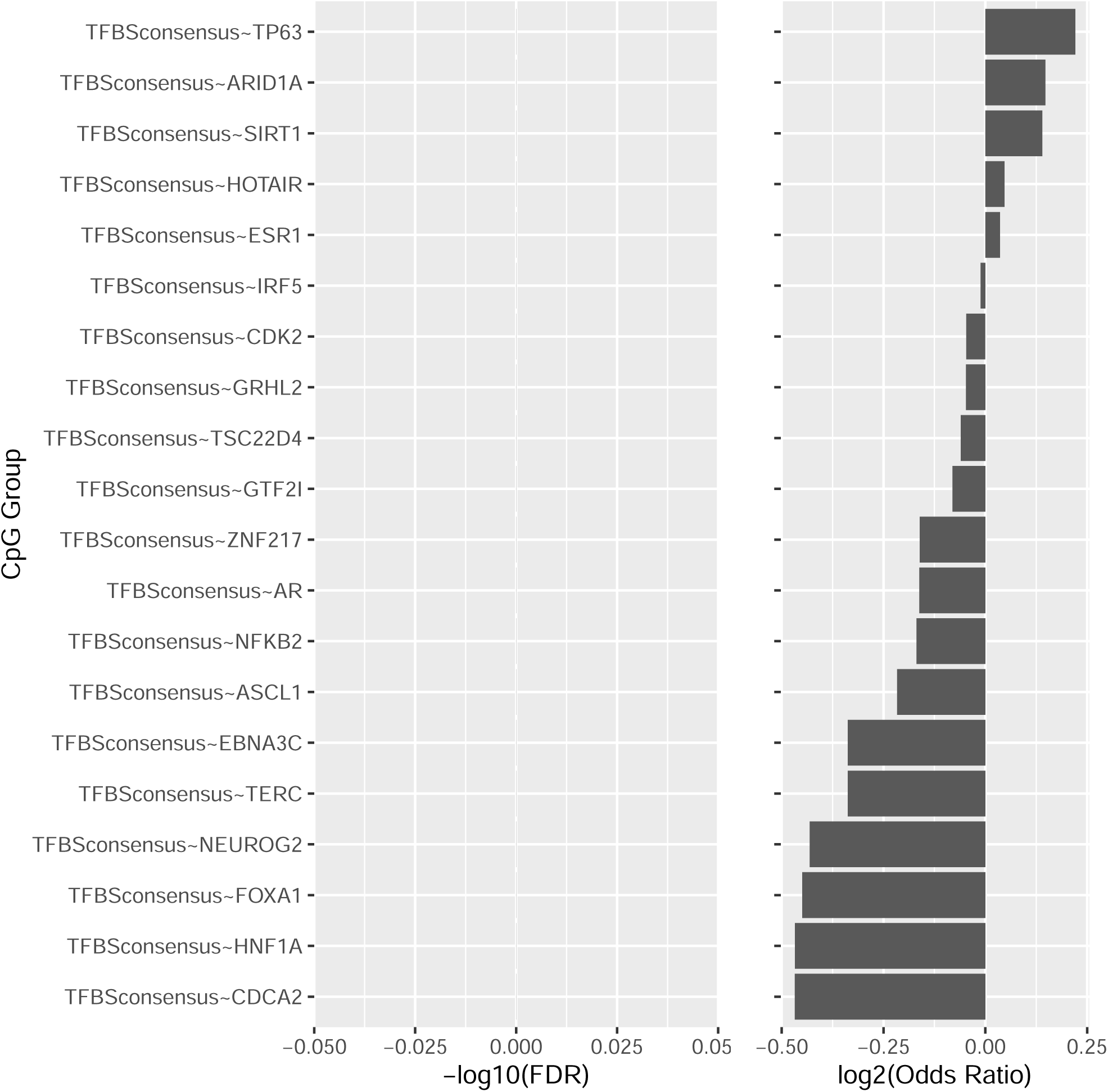
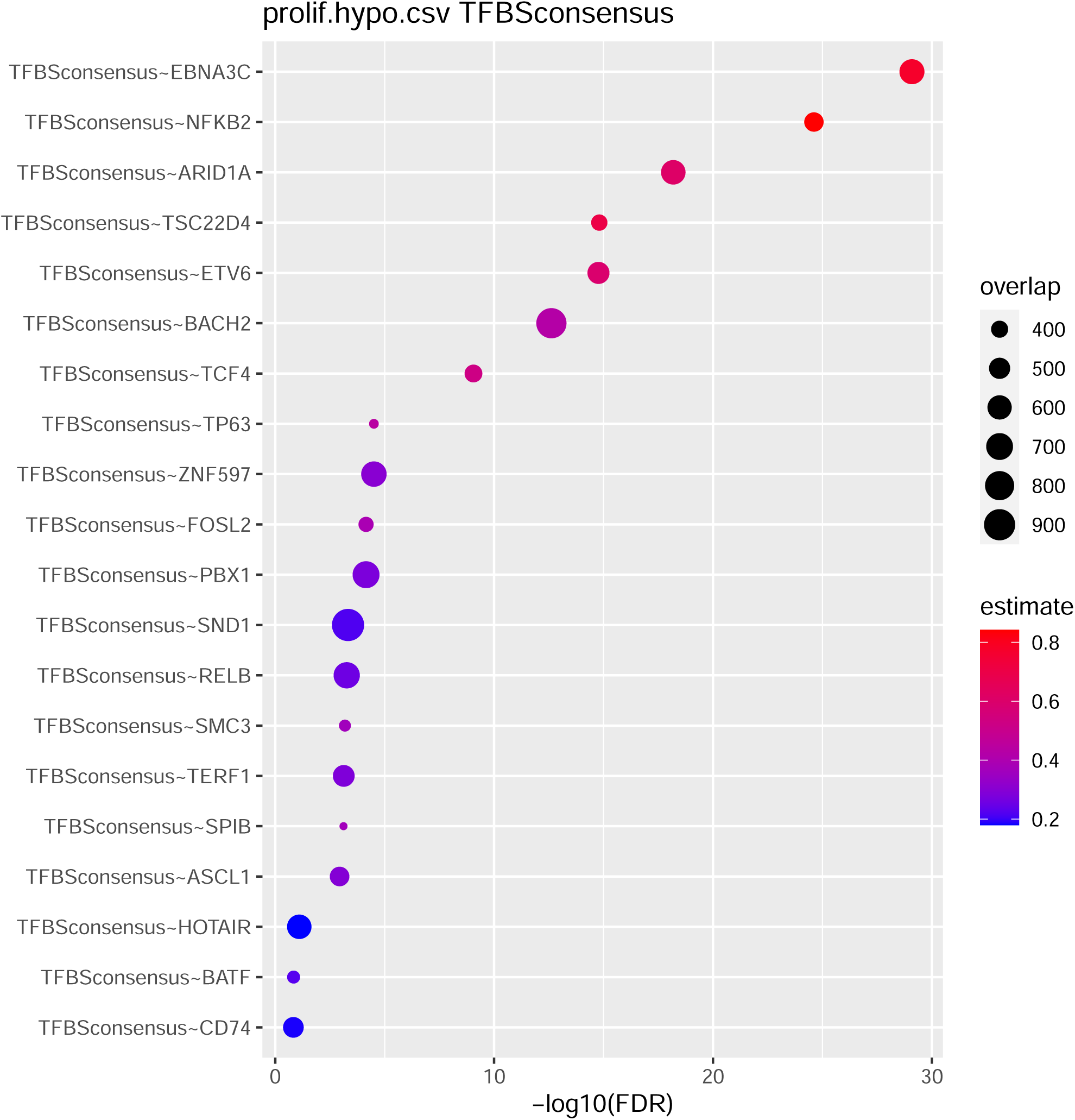
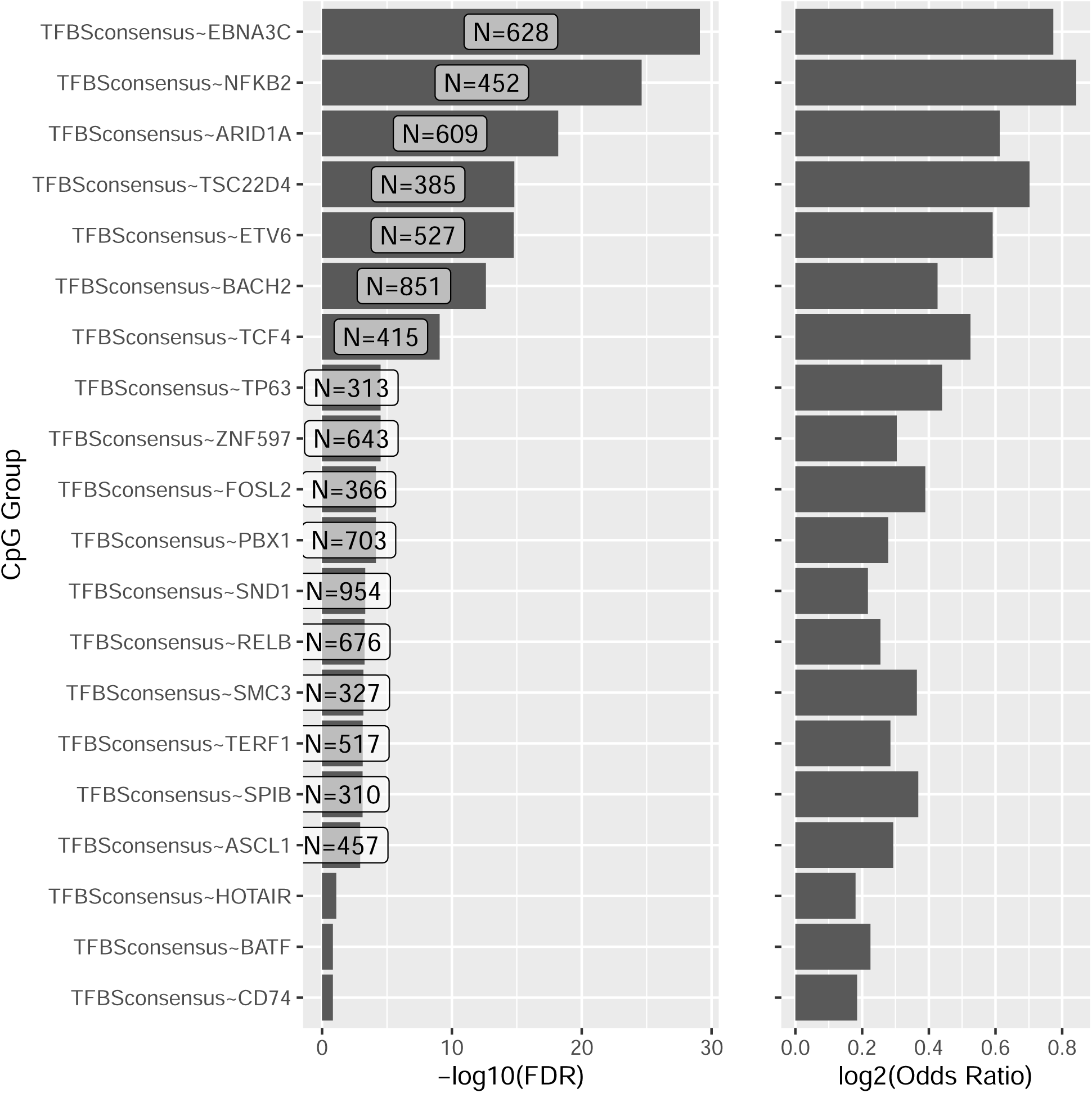
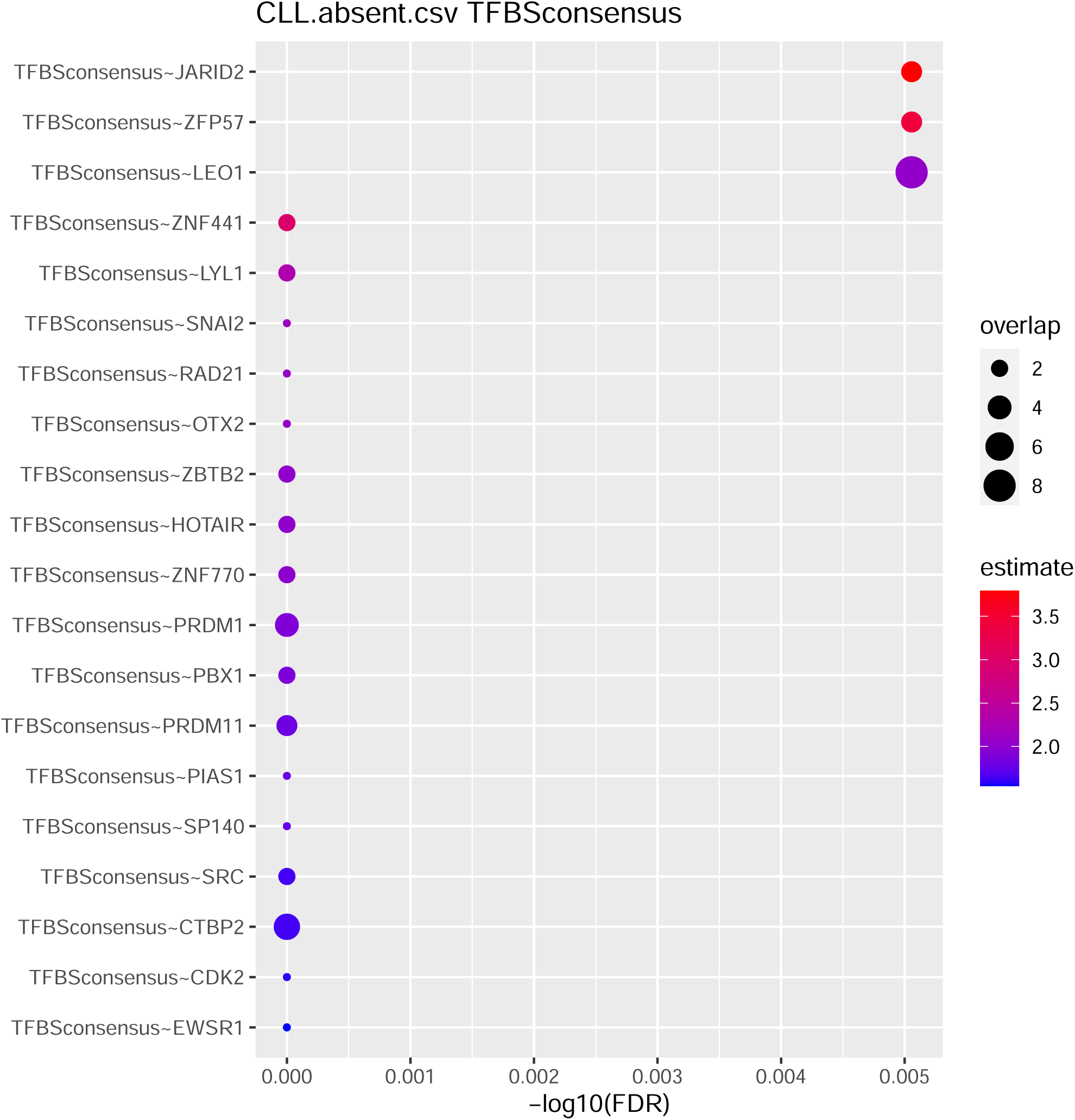
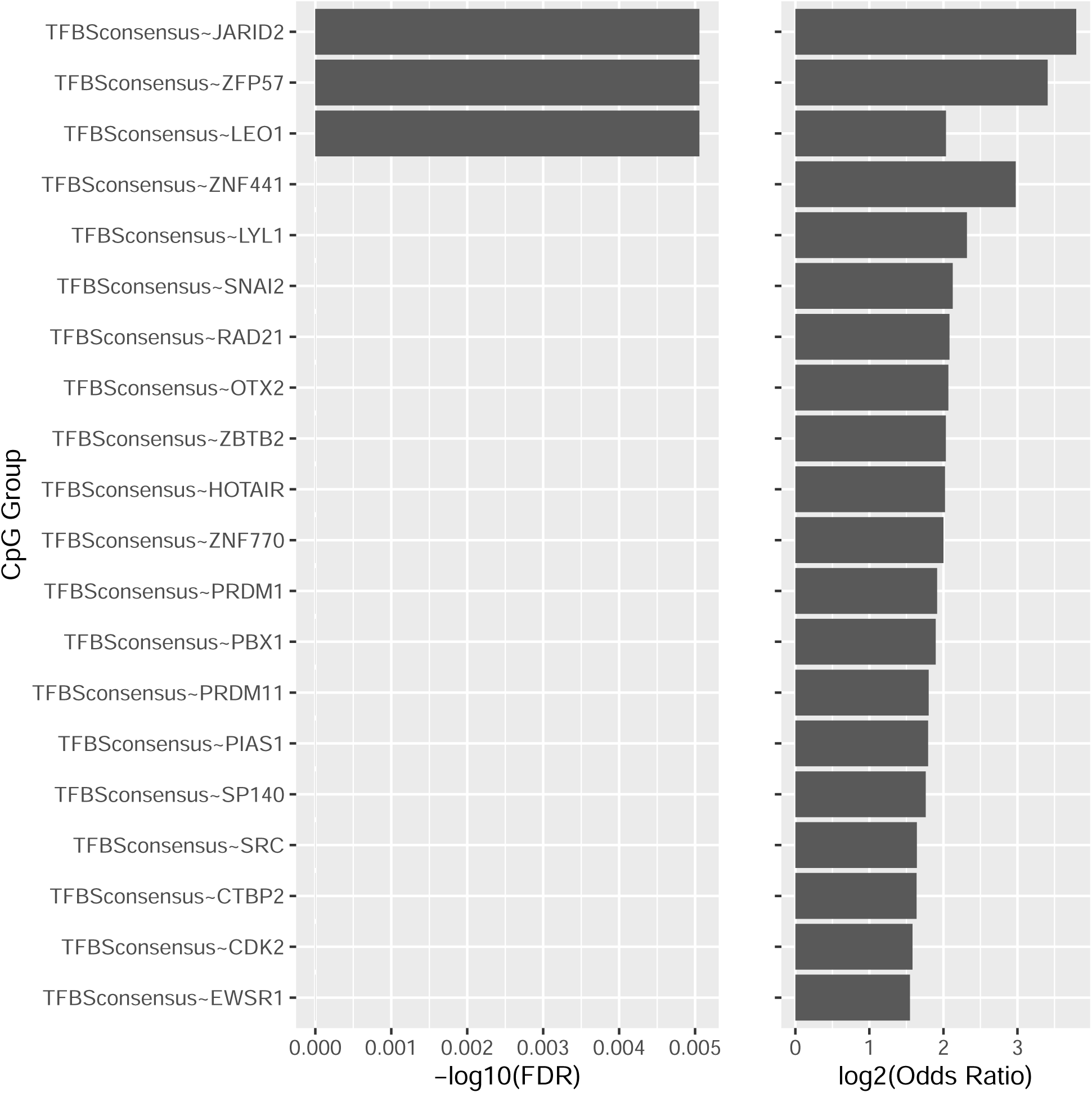
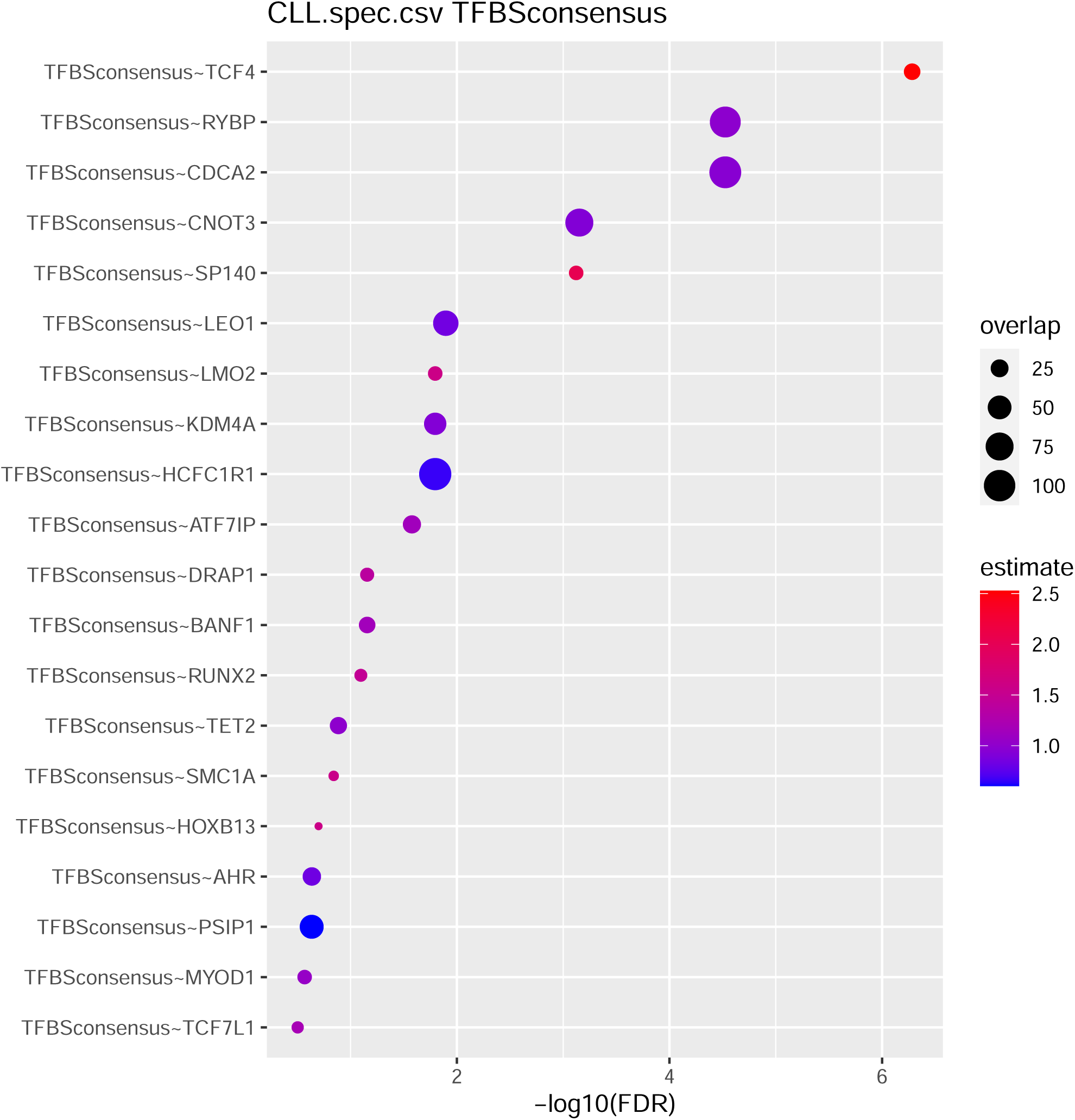
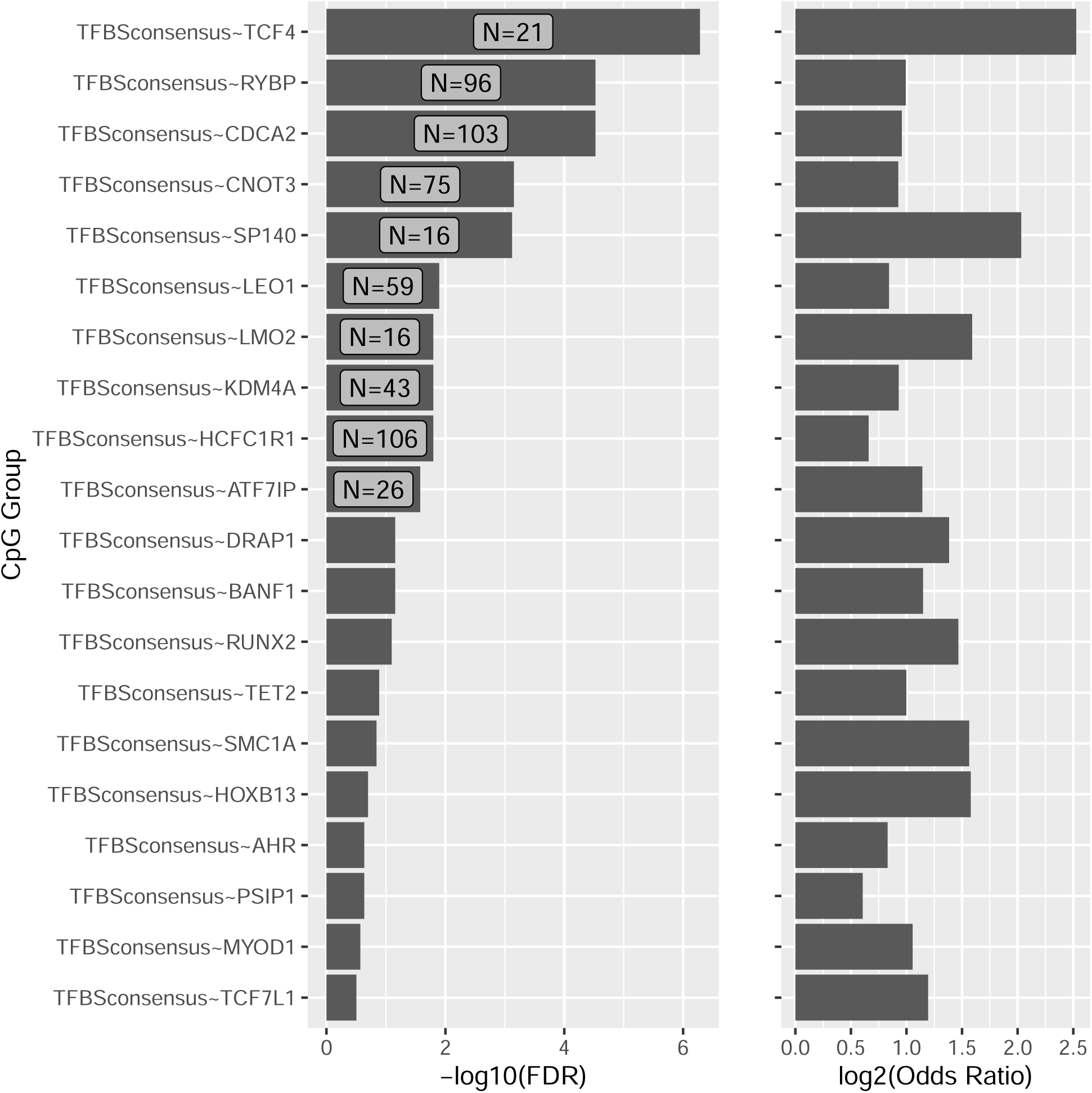
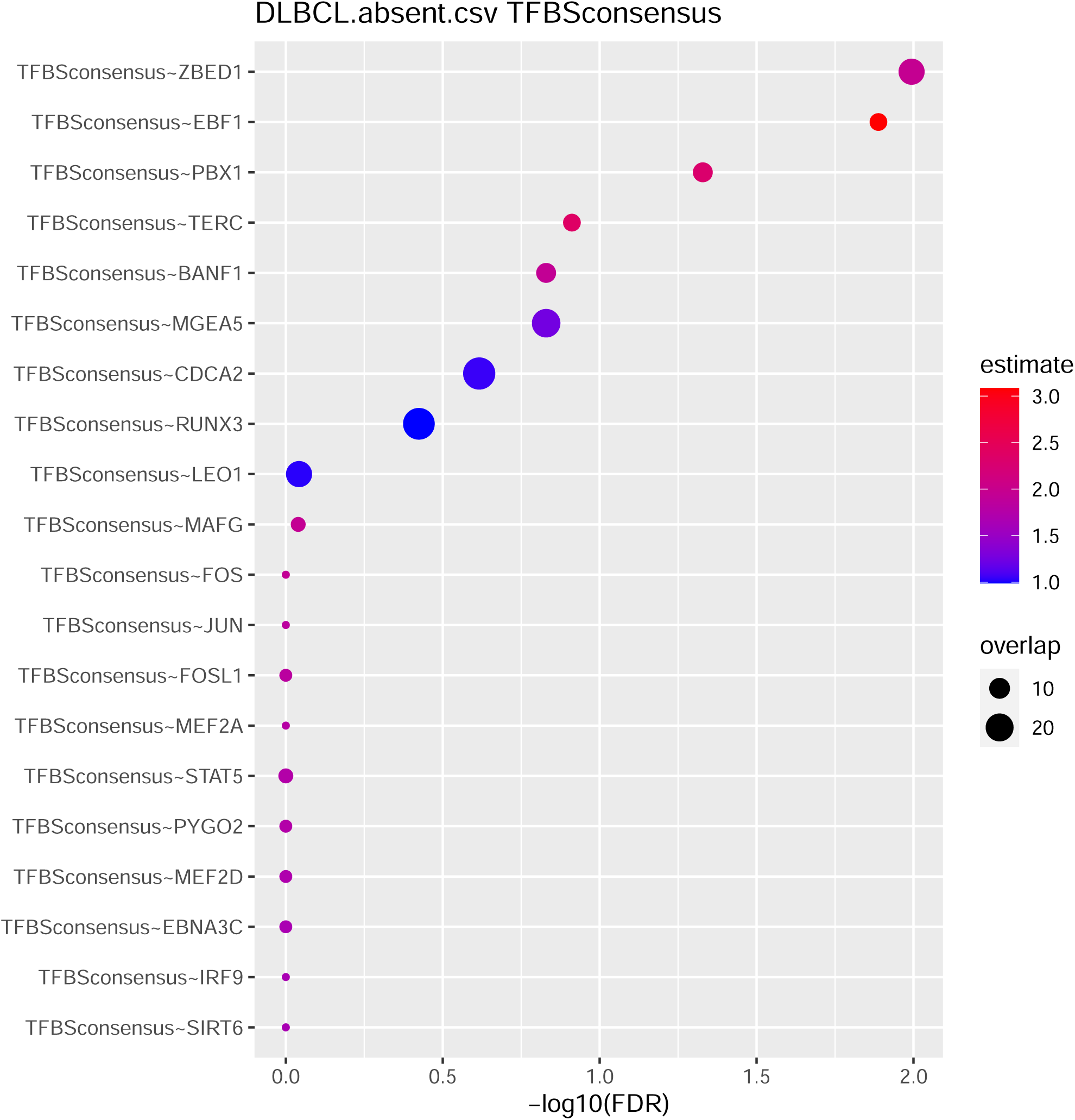
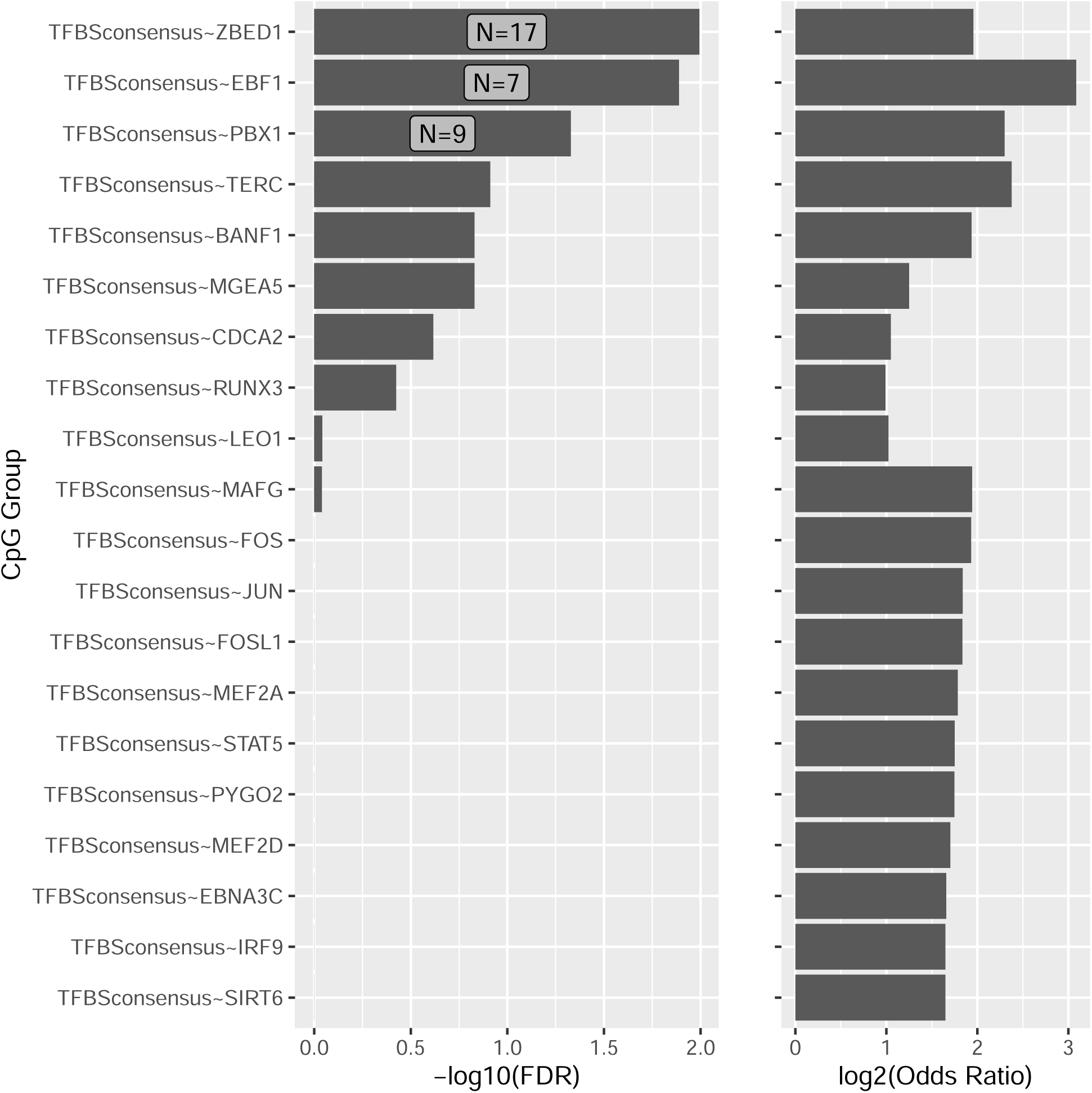
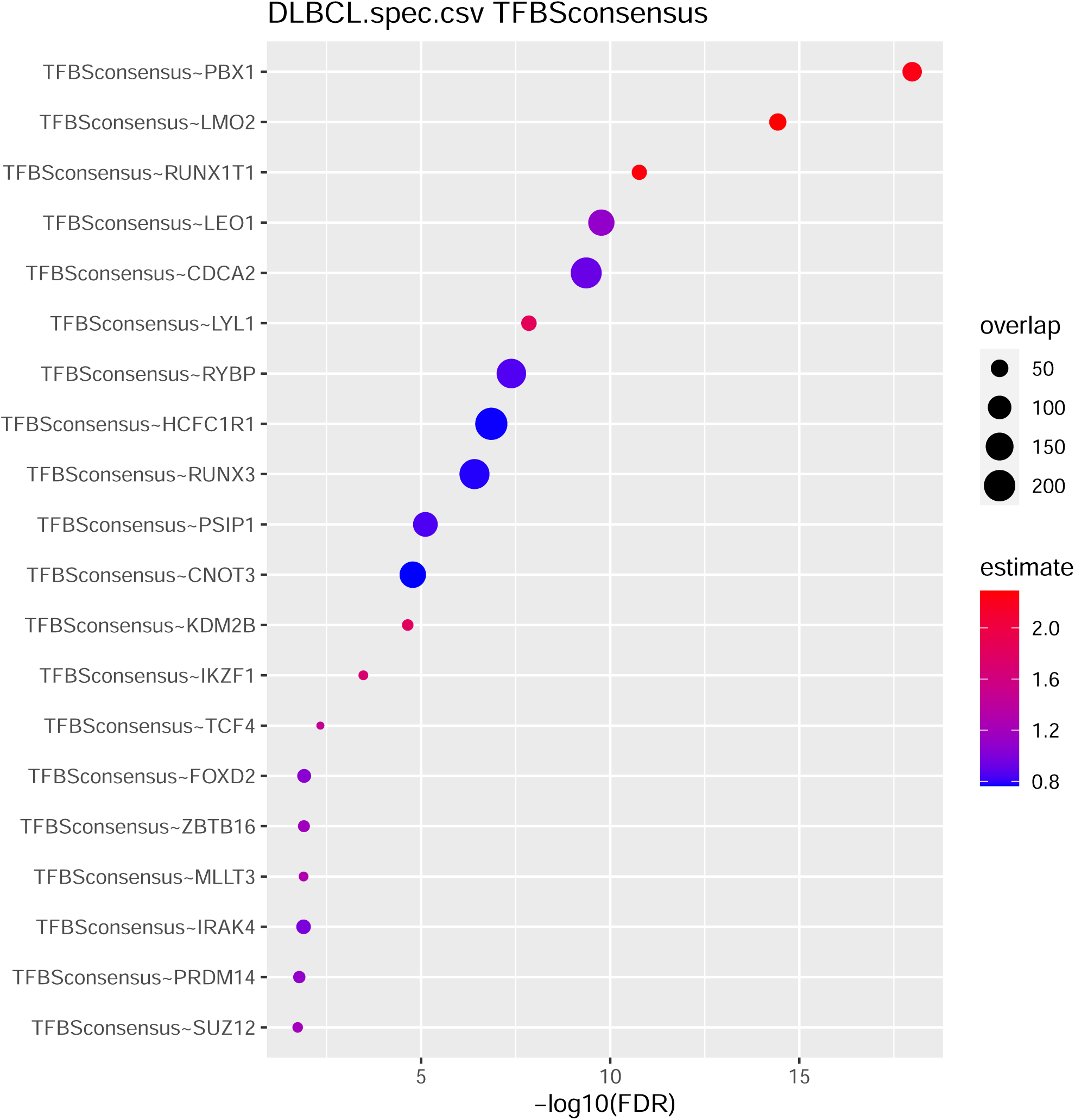
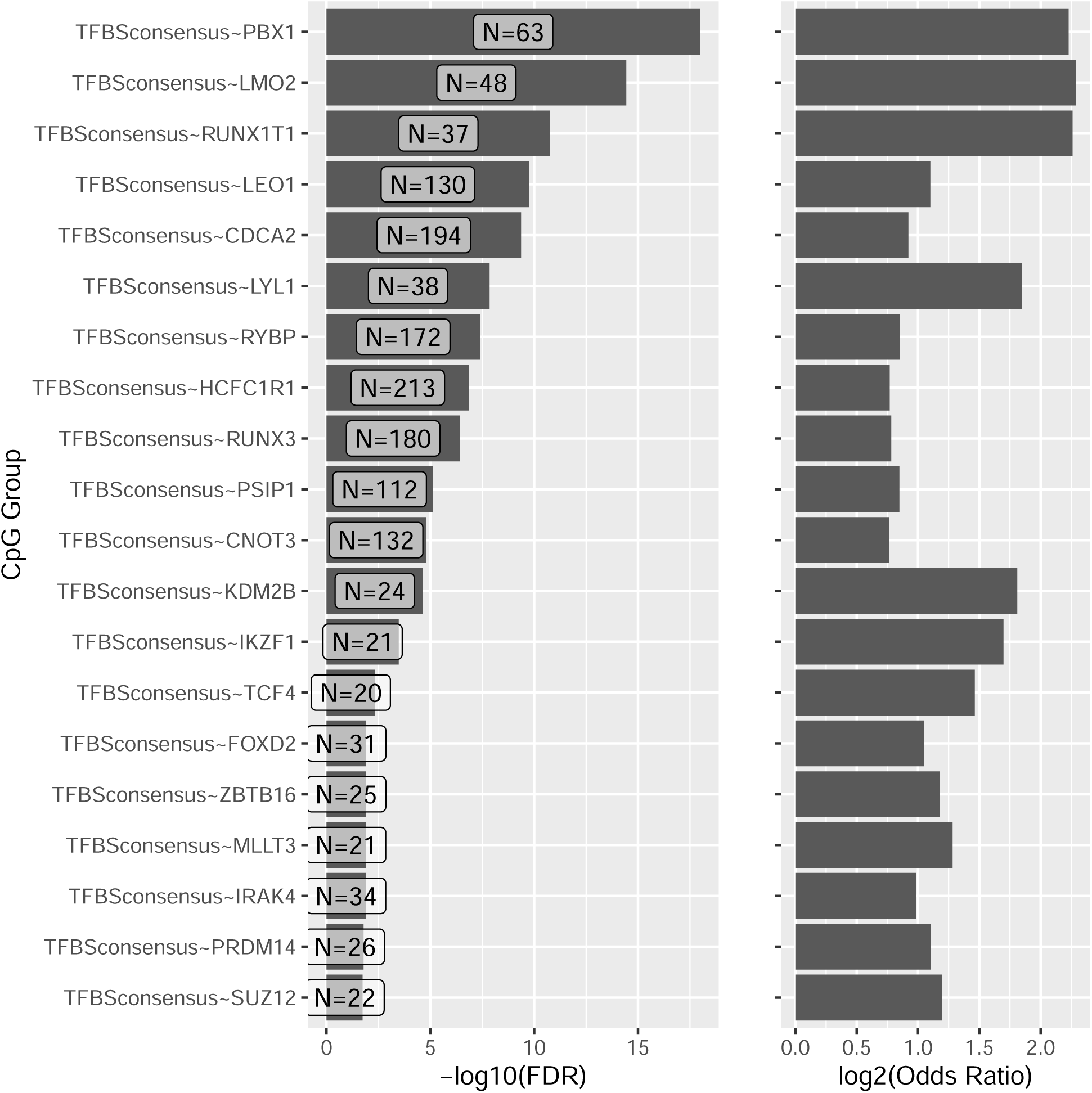
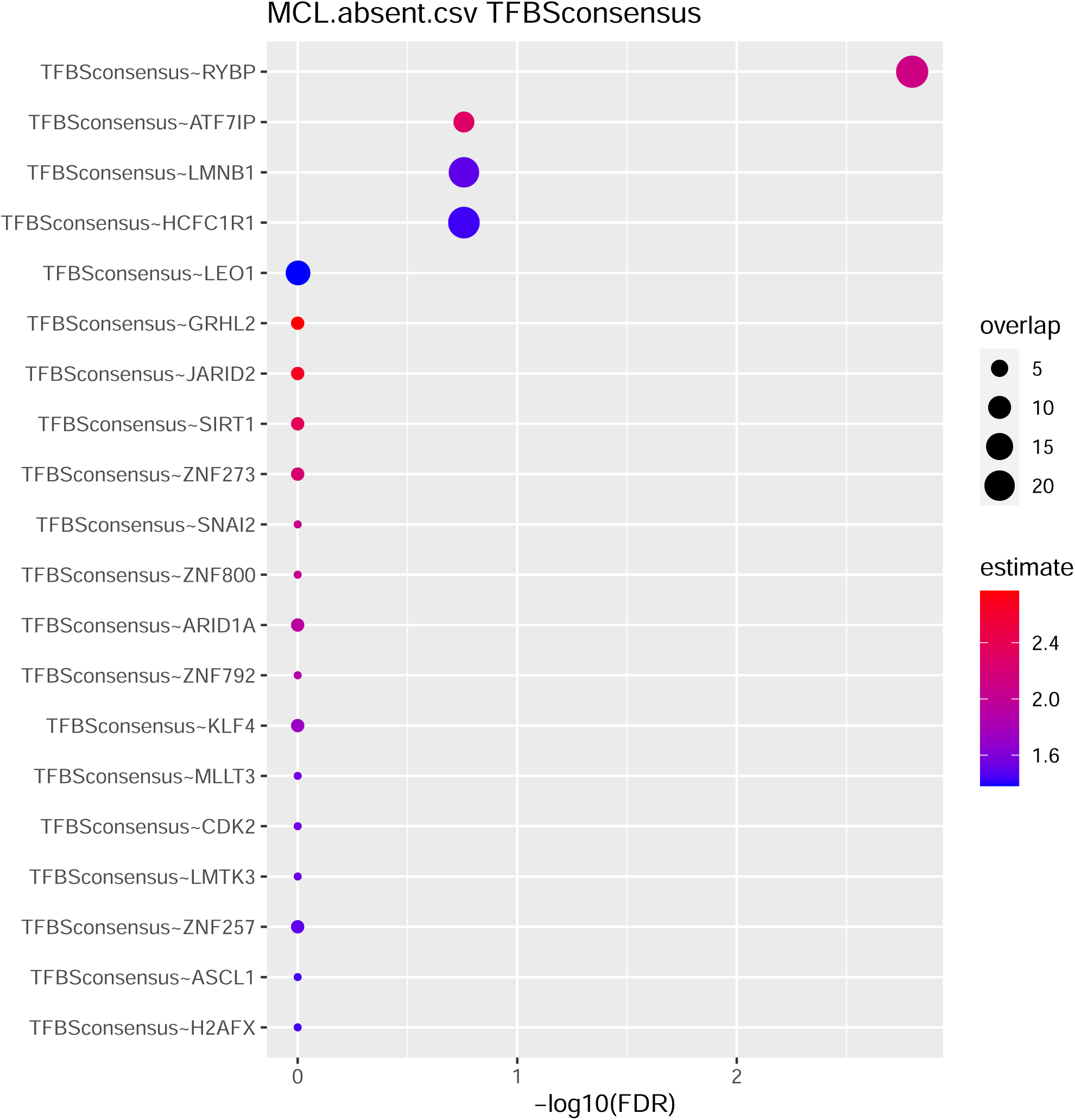
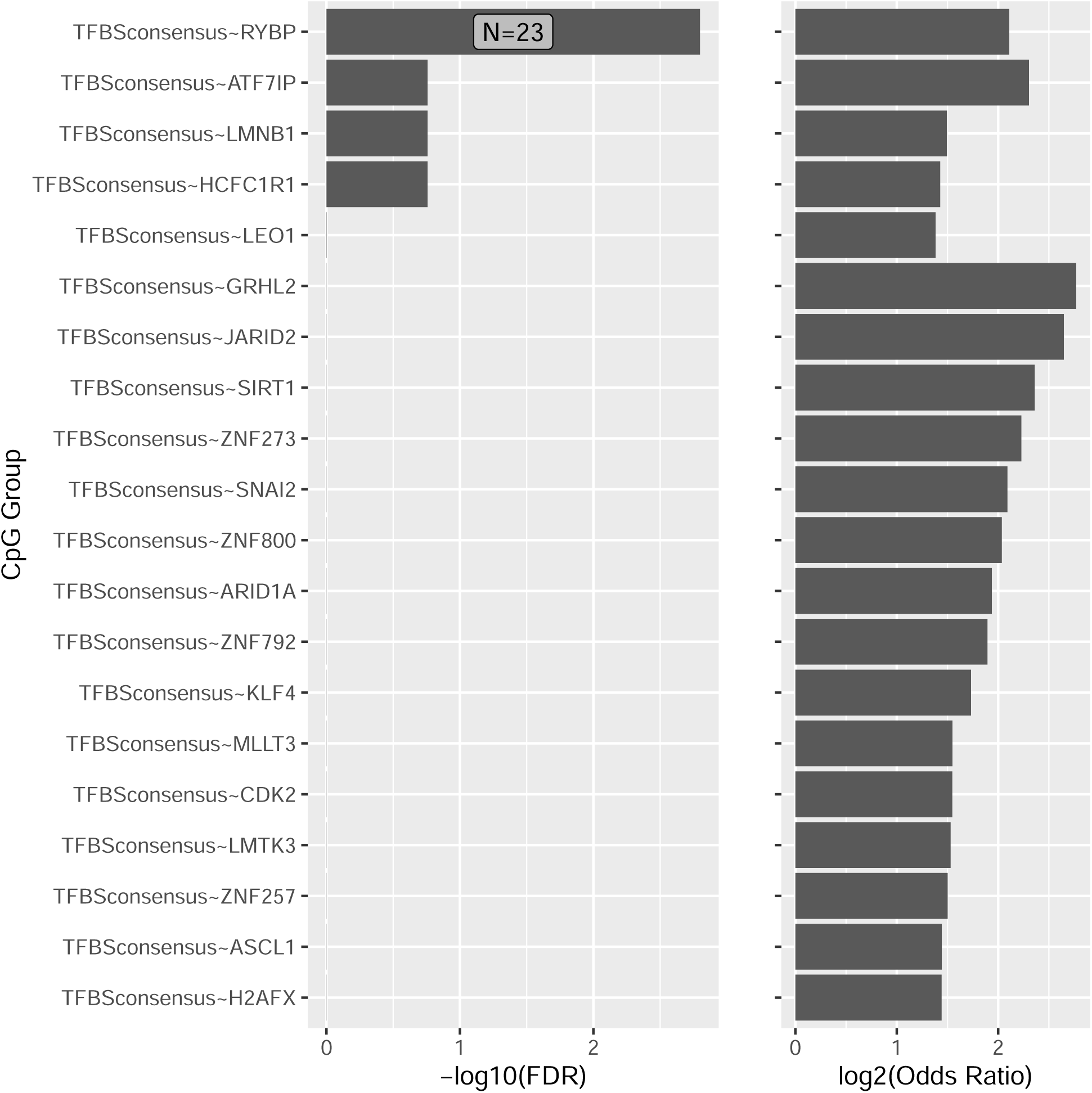
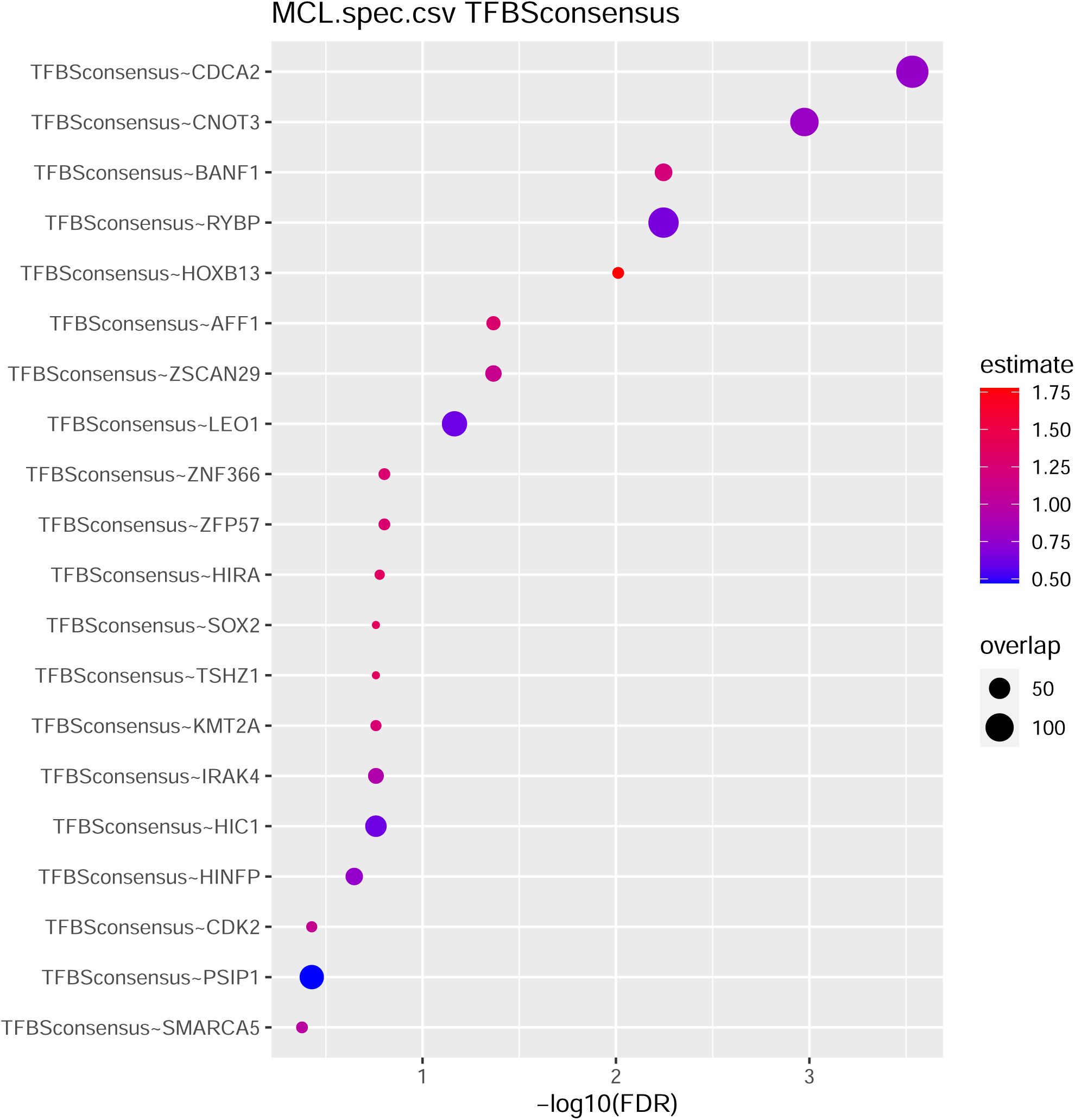
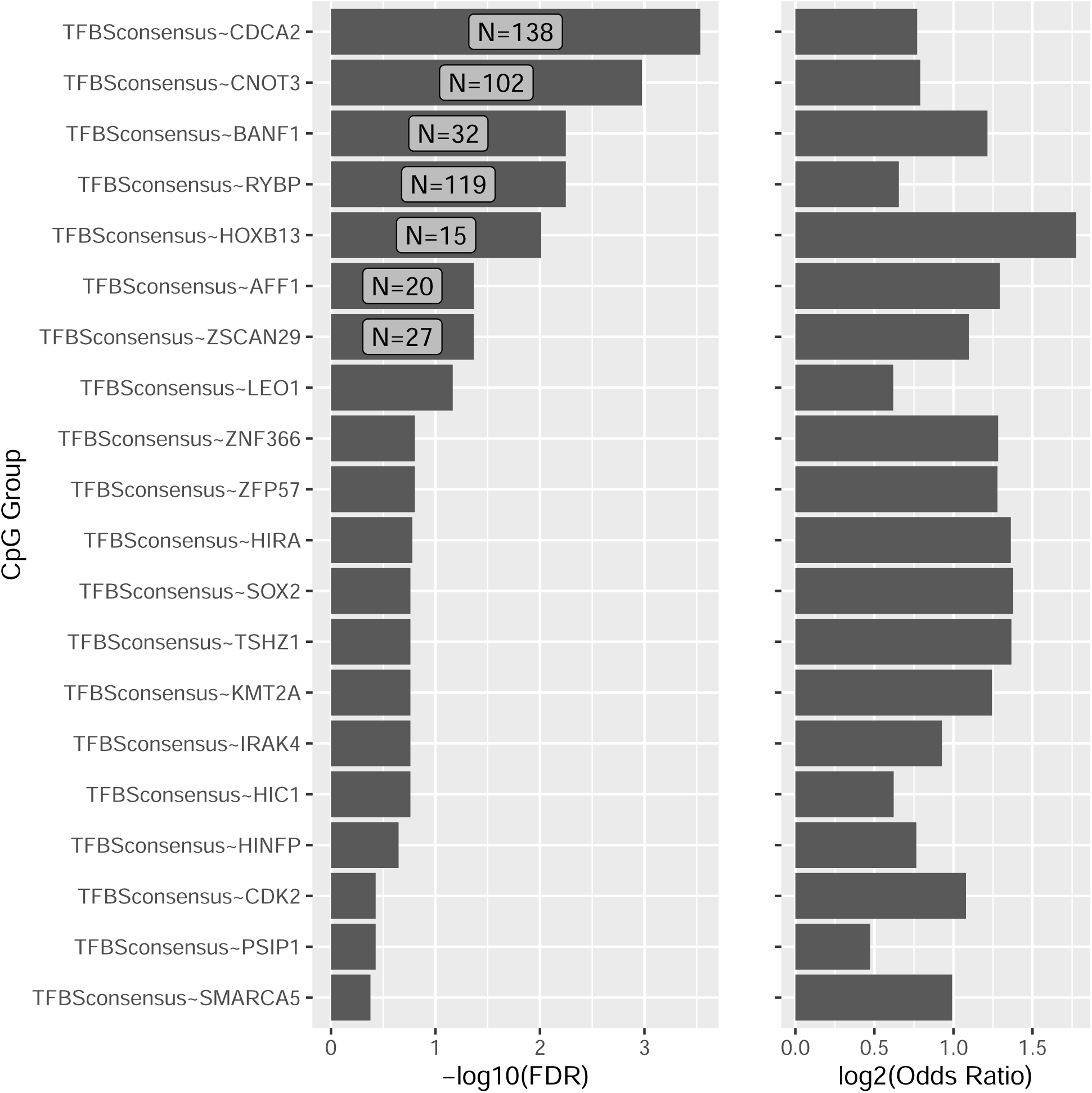
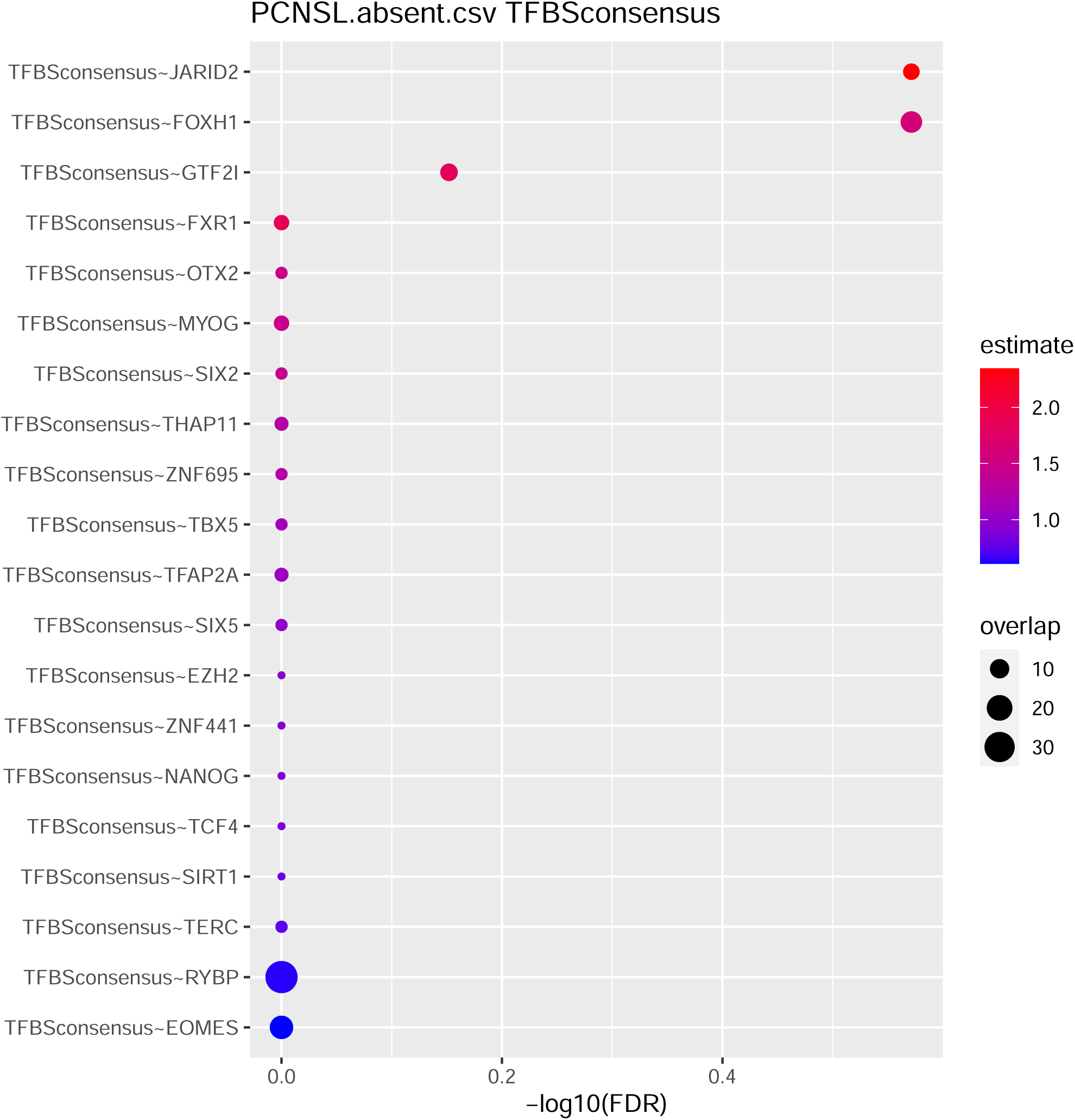
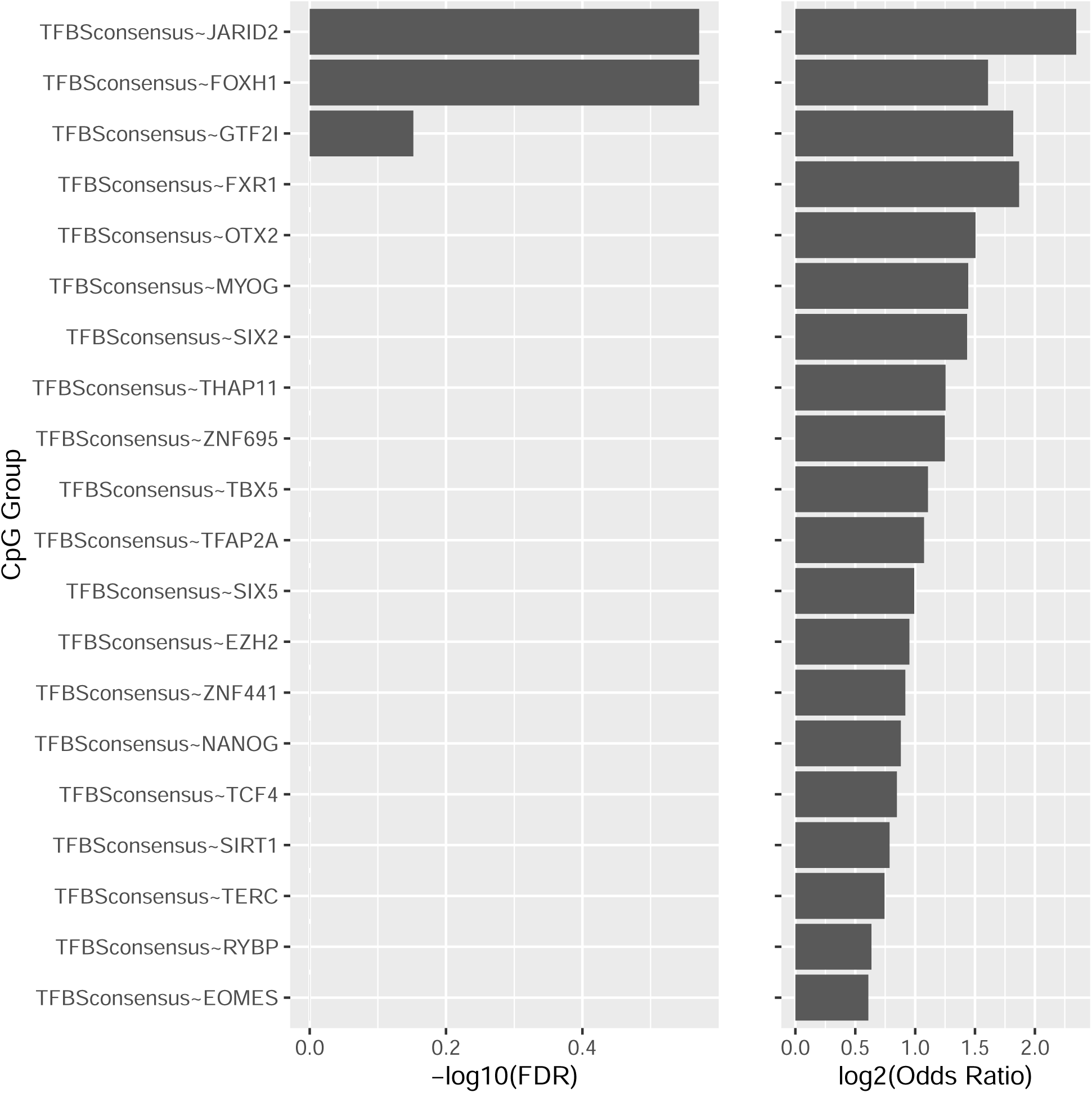
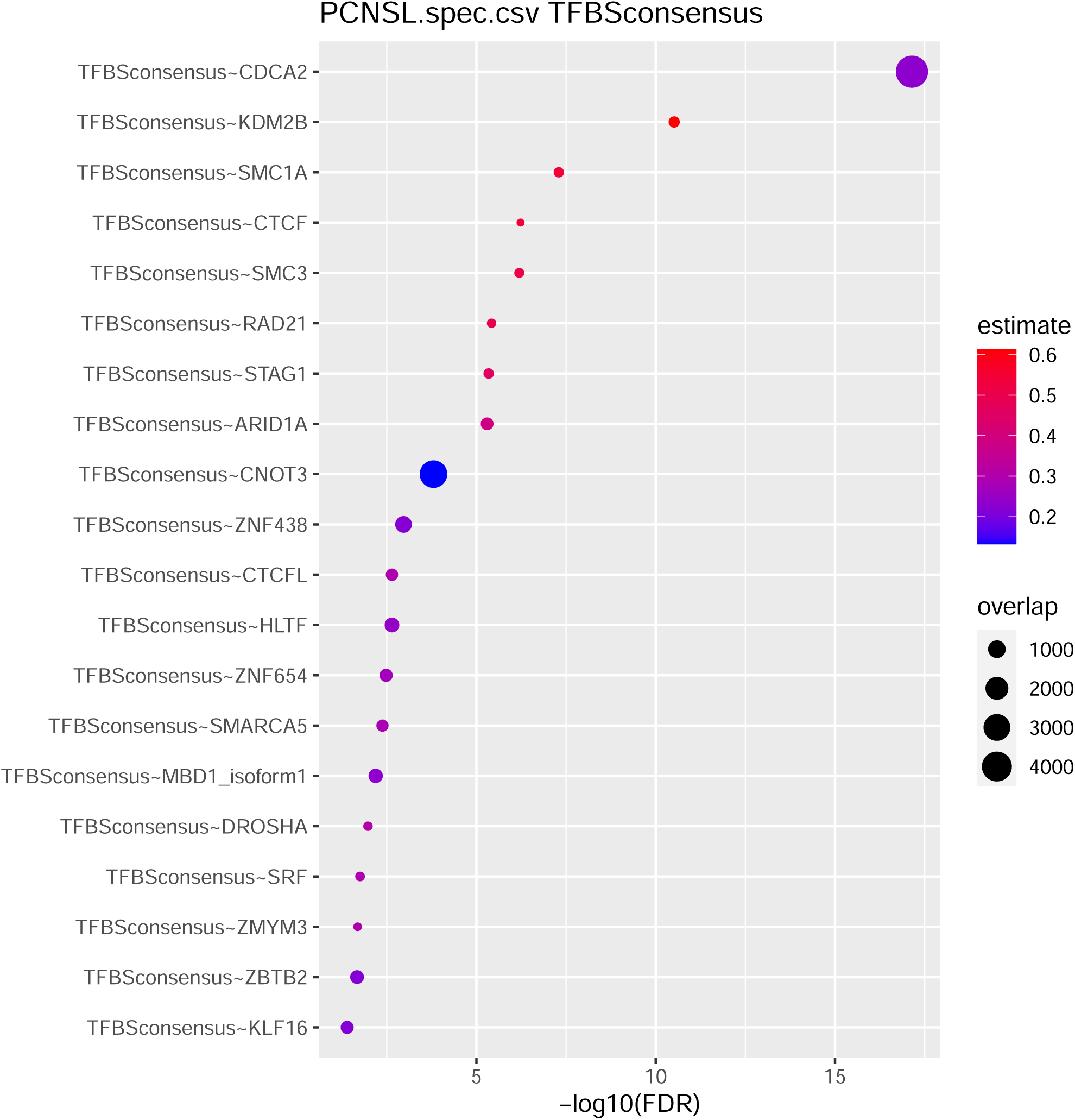
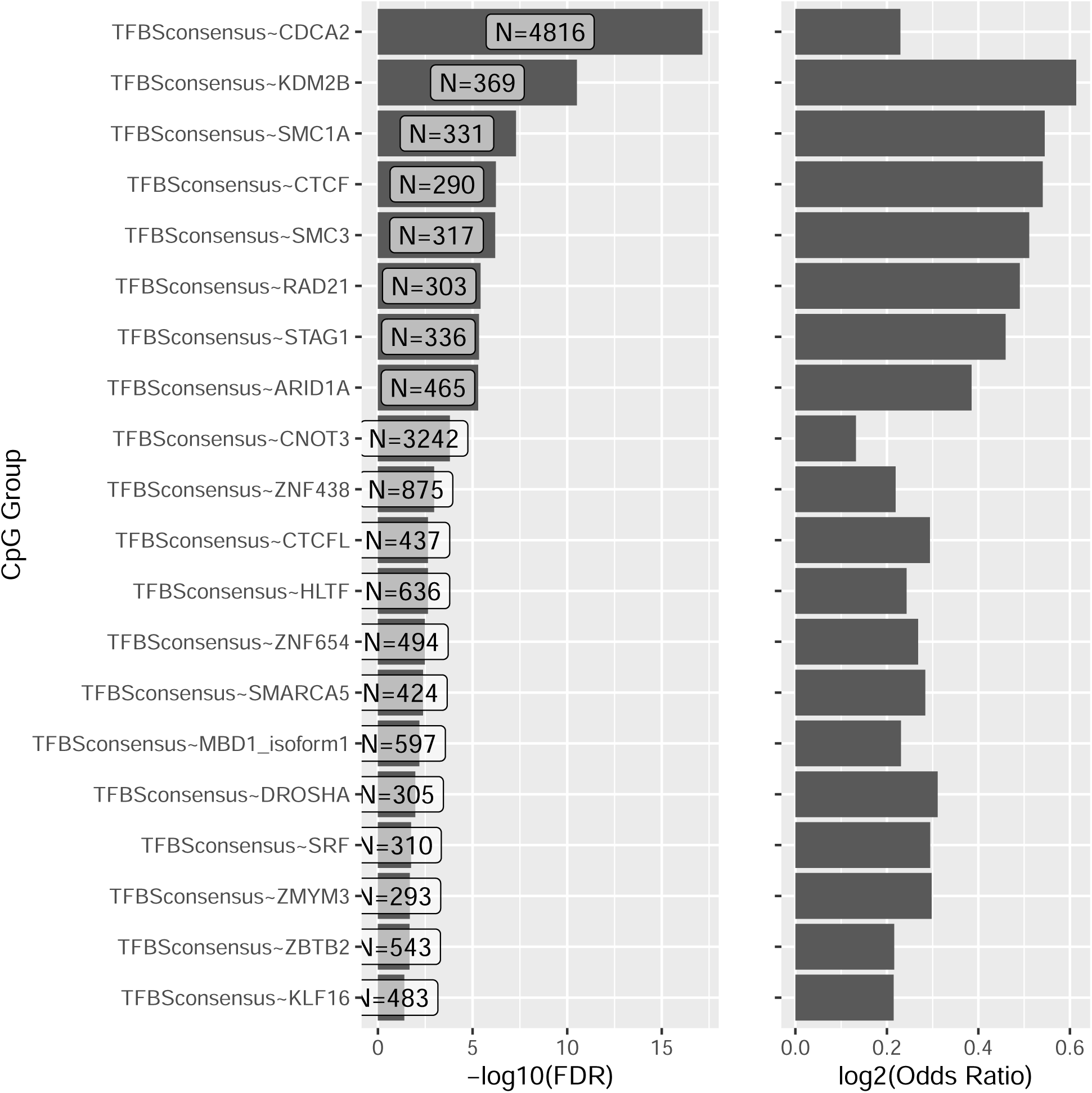
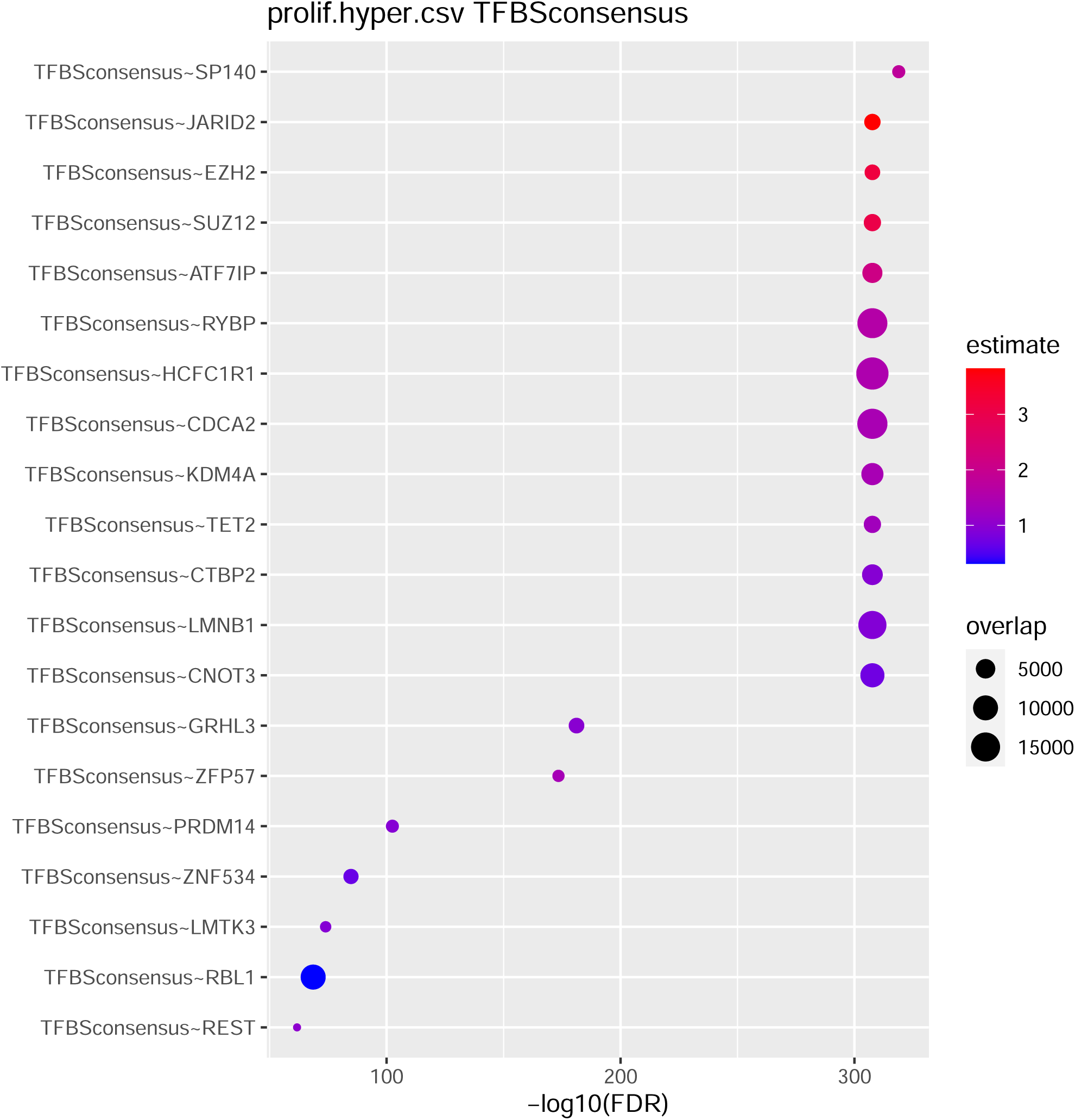
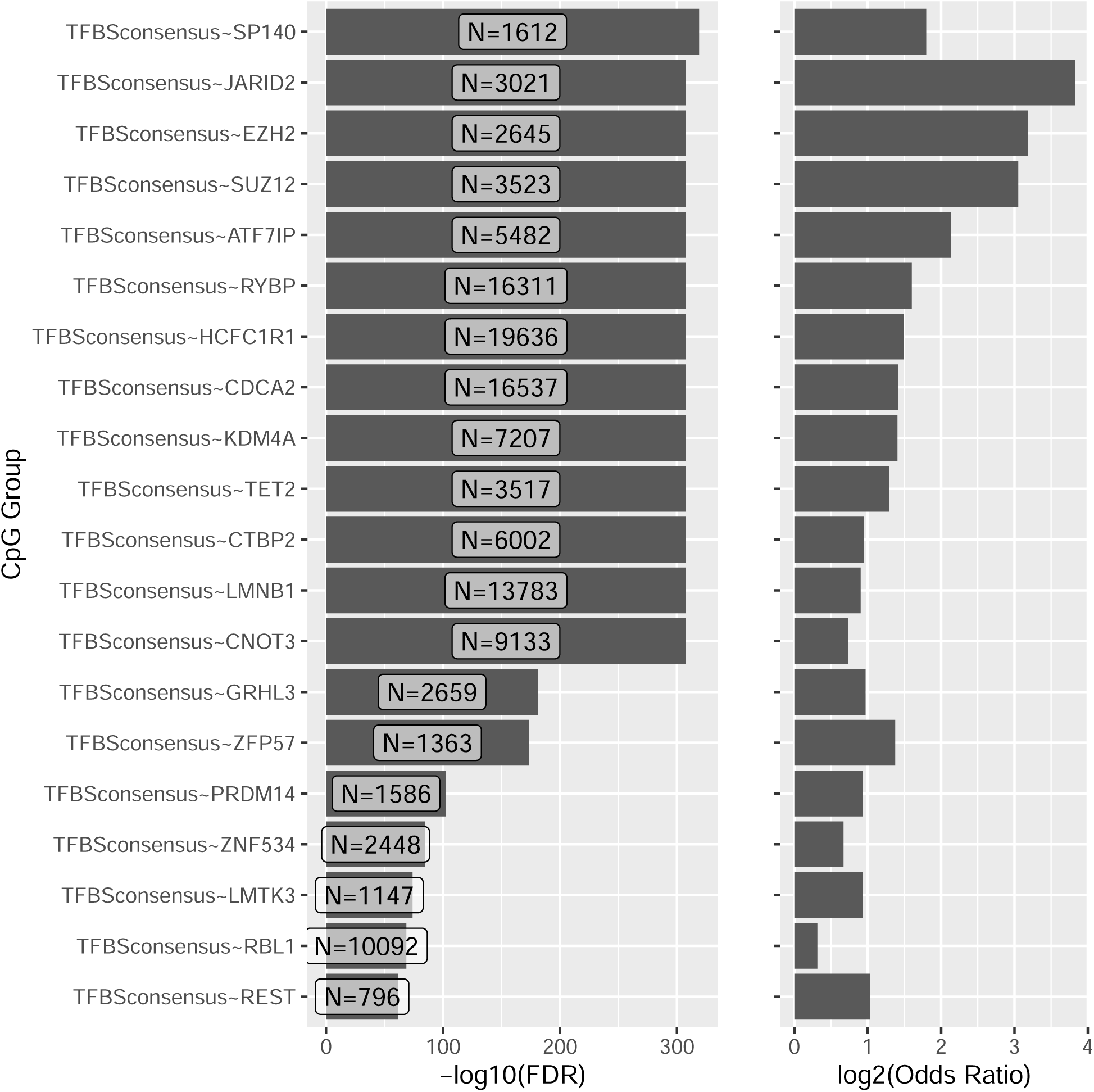

**Supplementary Figure 6.**
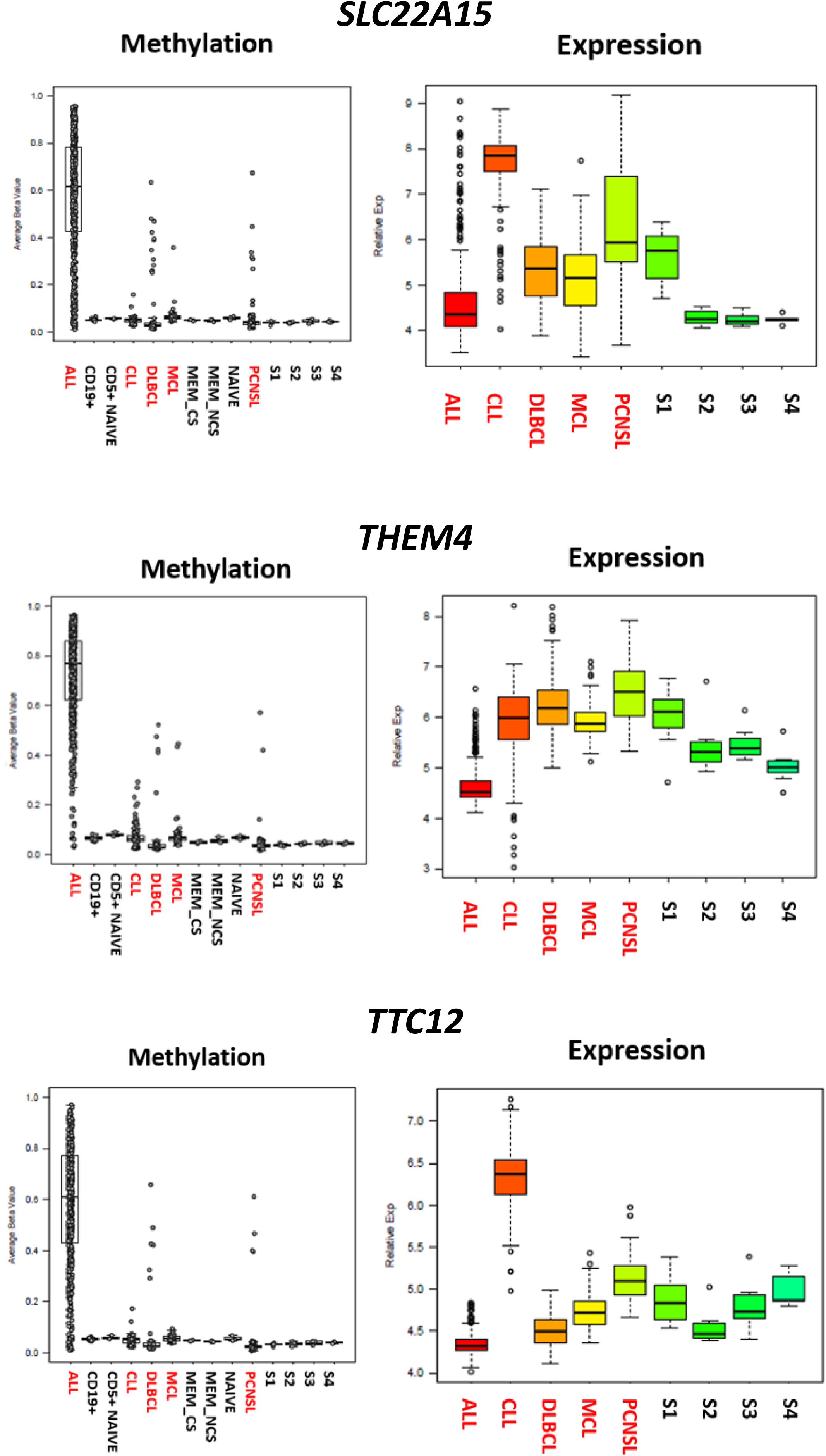
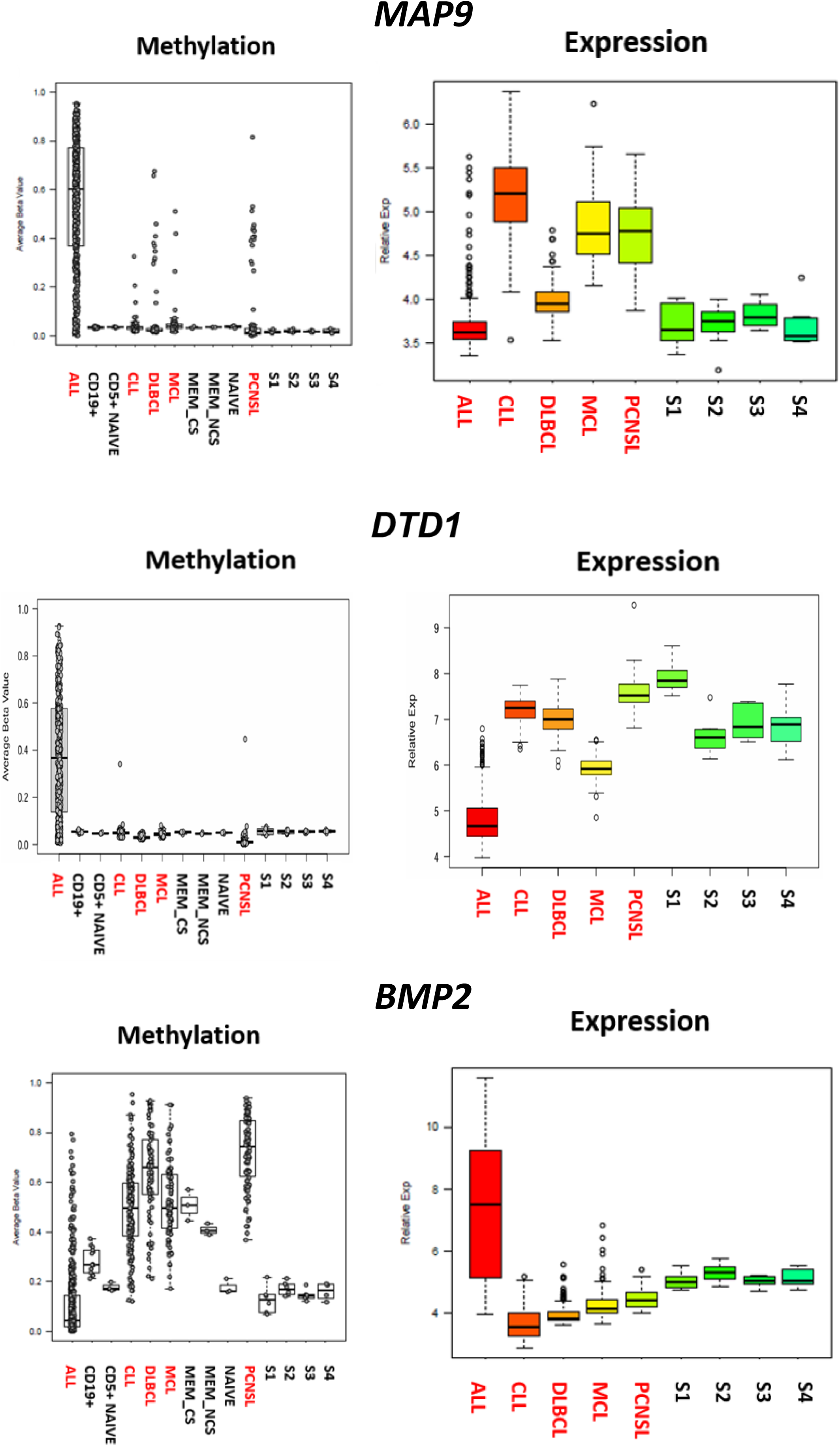
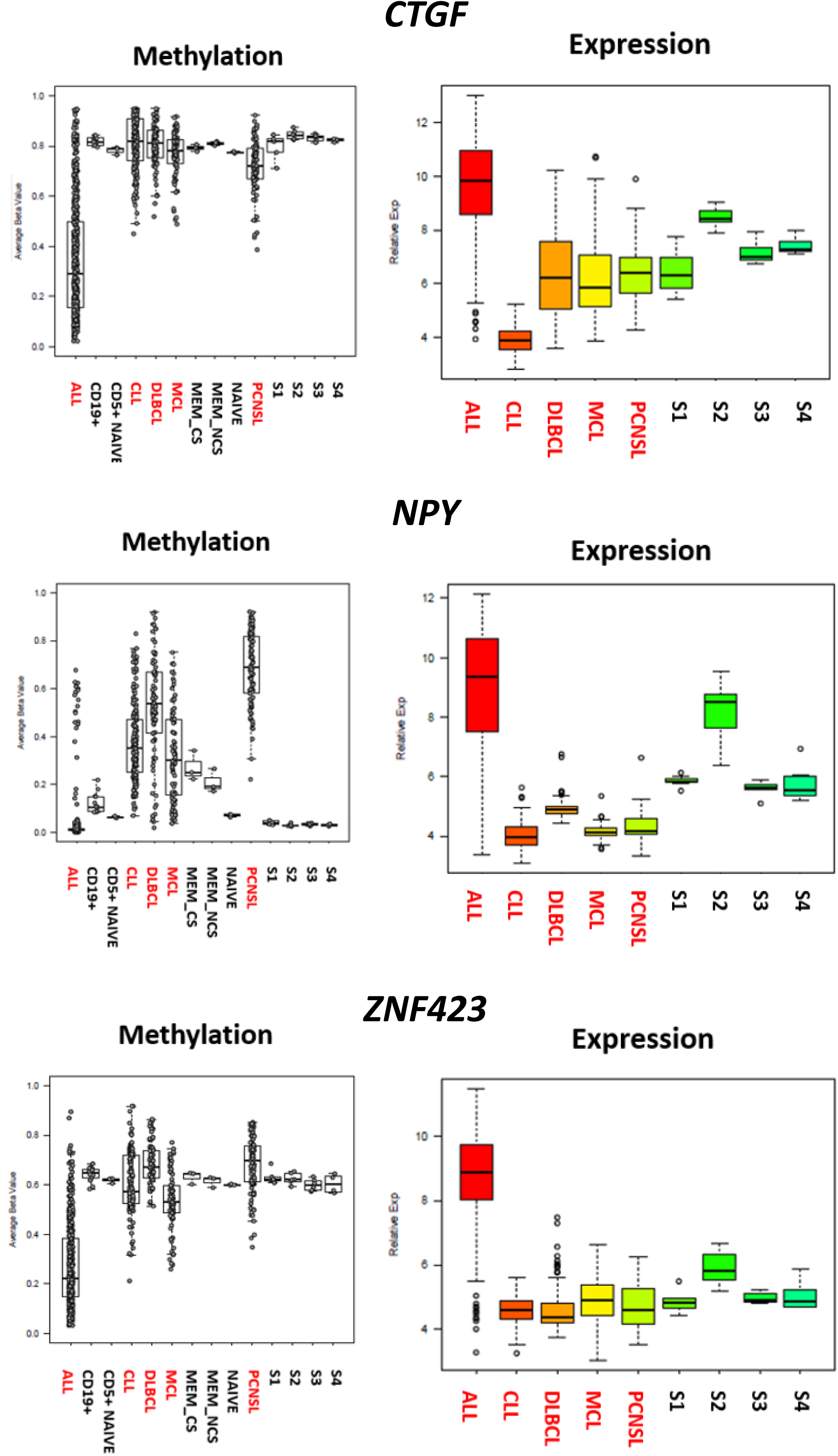
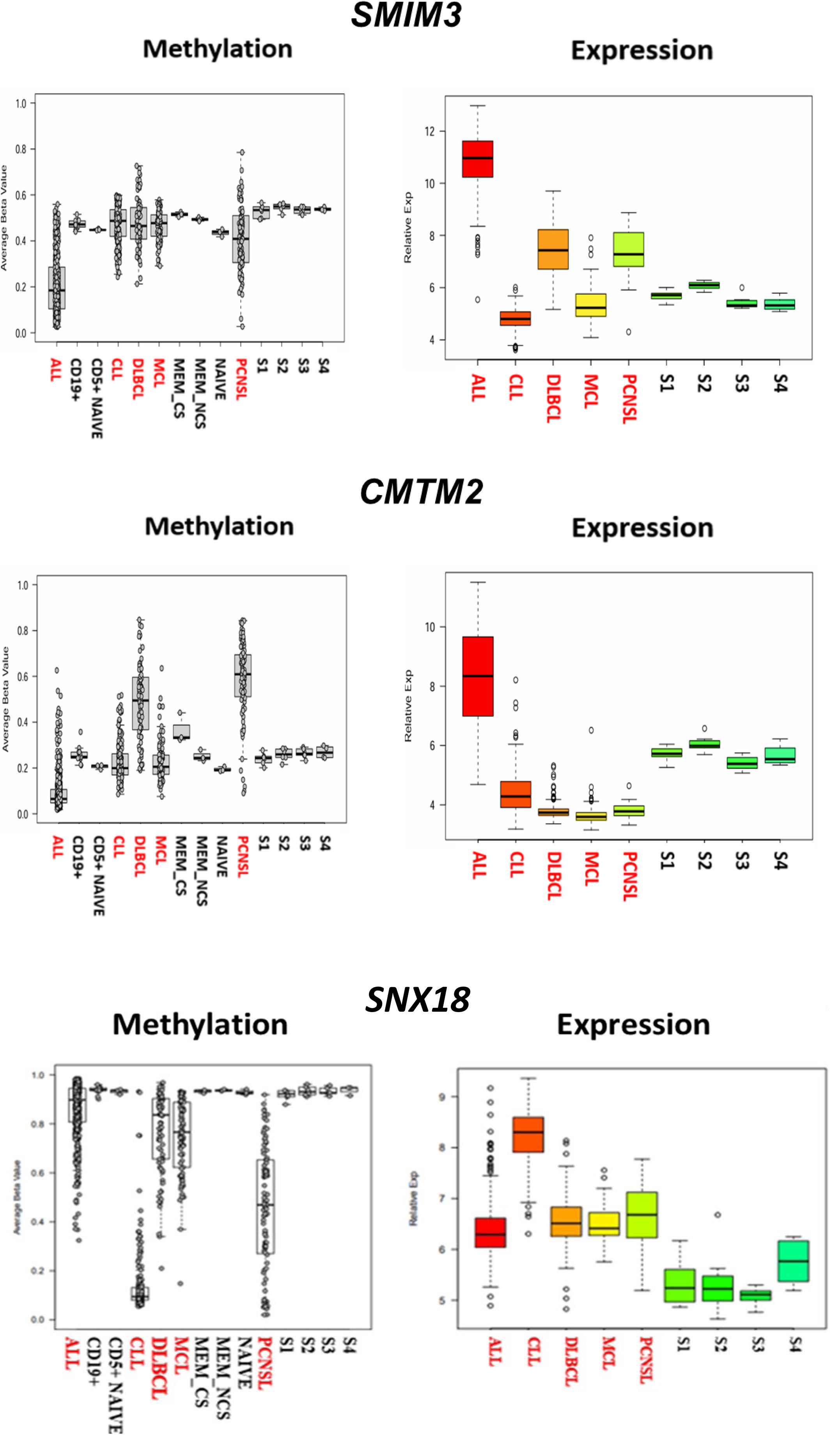
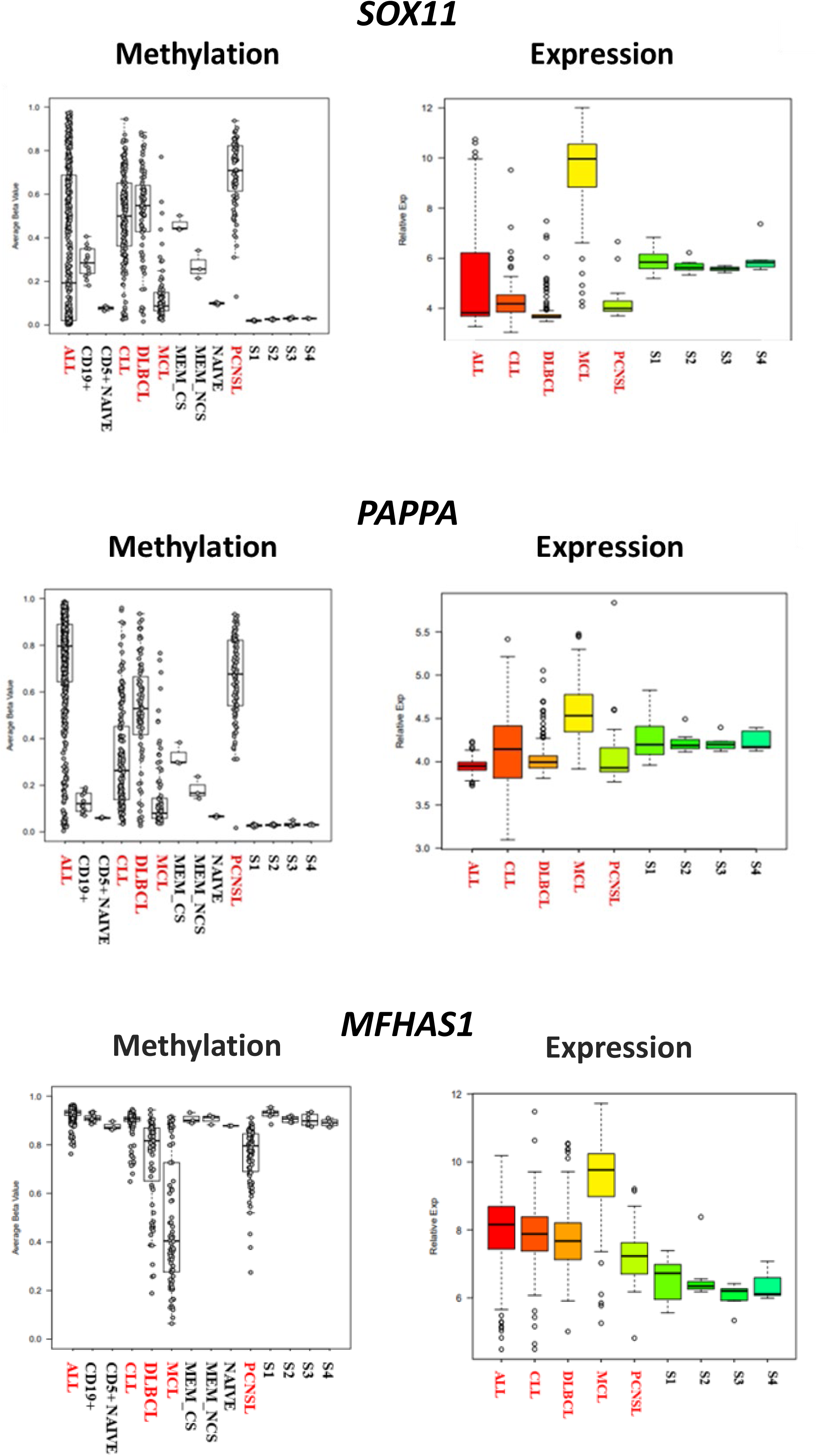
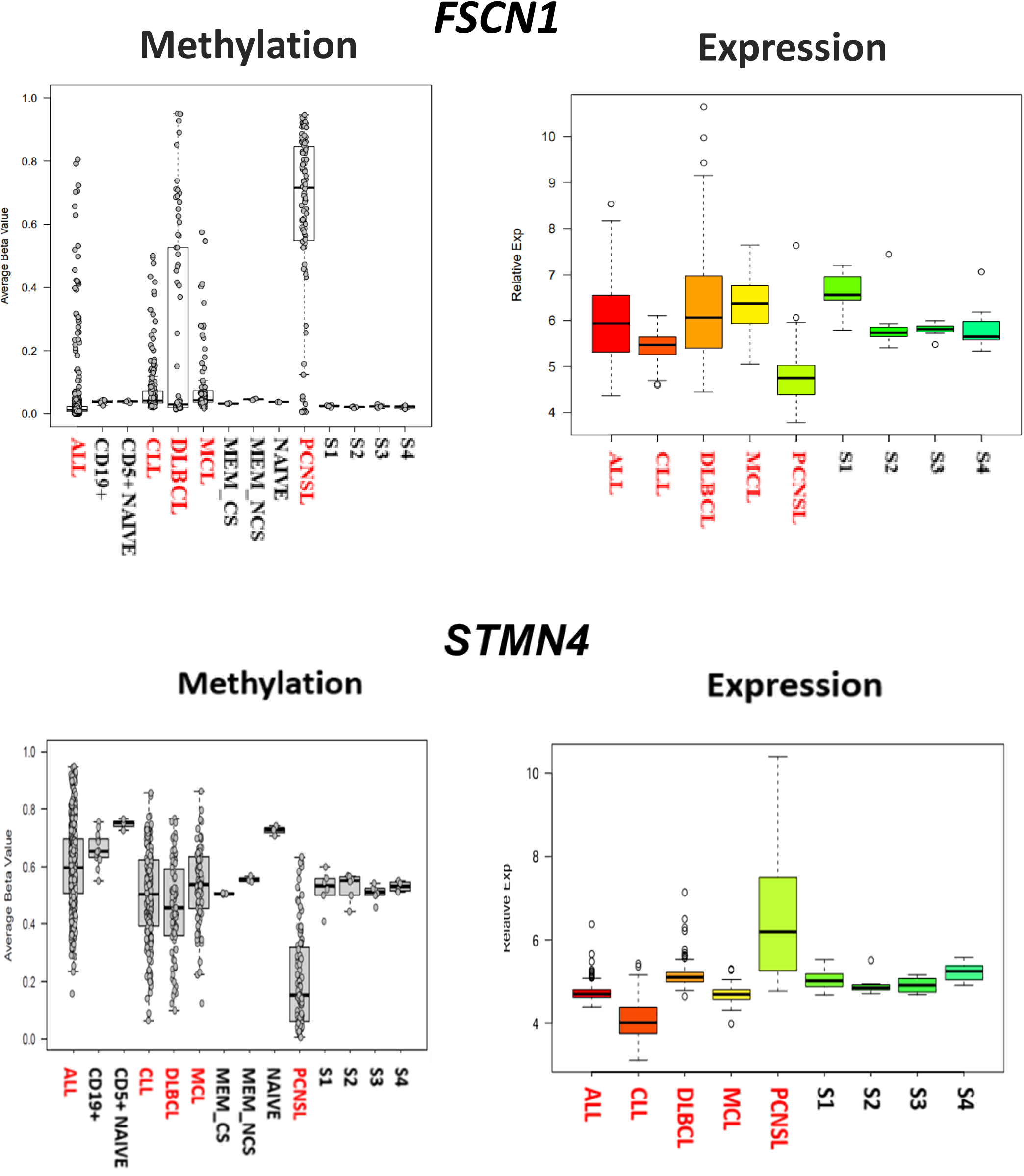

**Supplementary figure 7.**
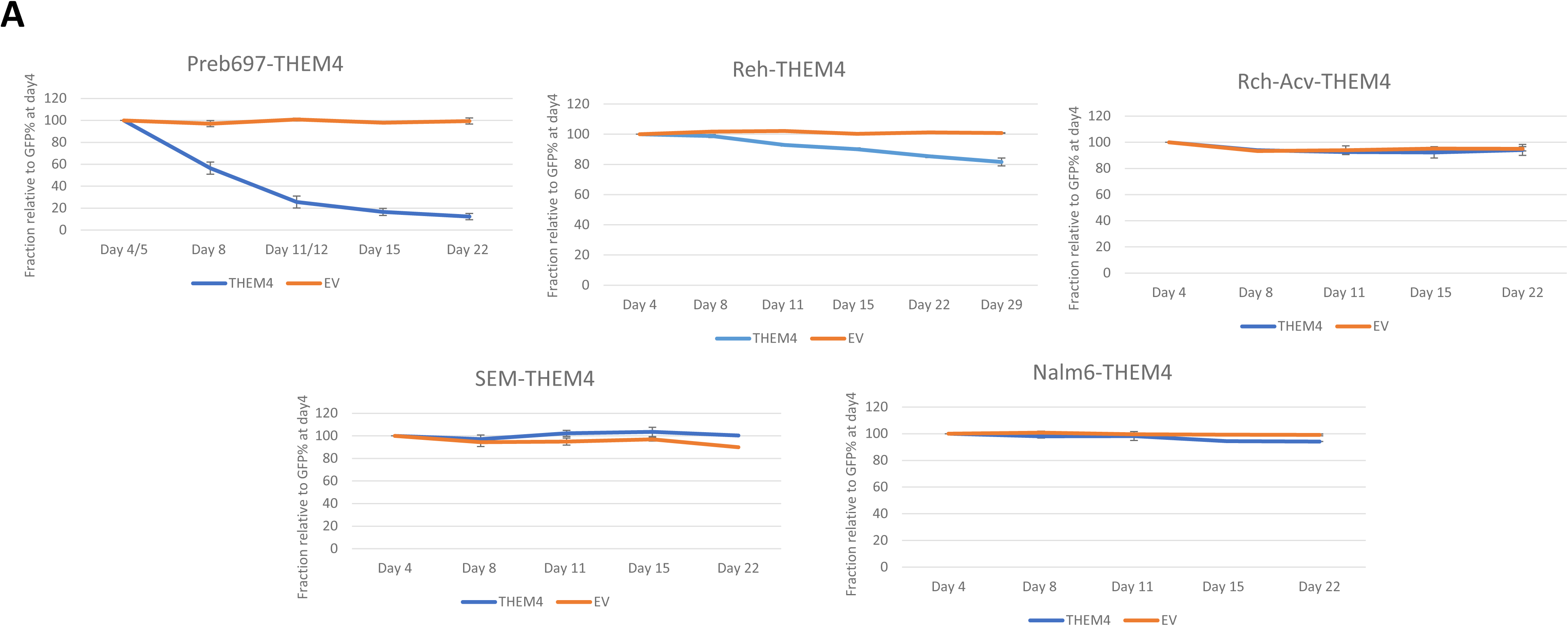
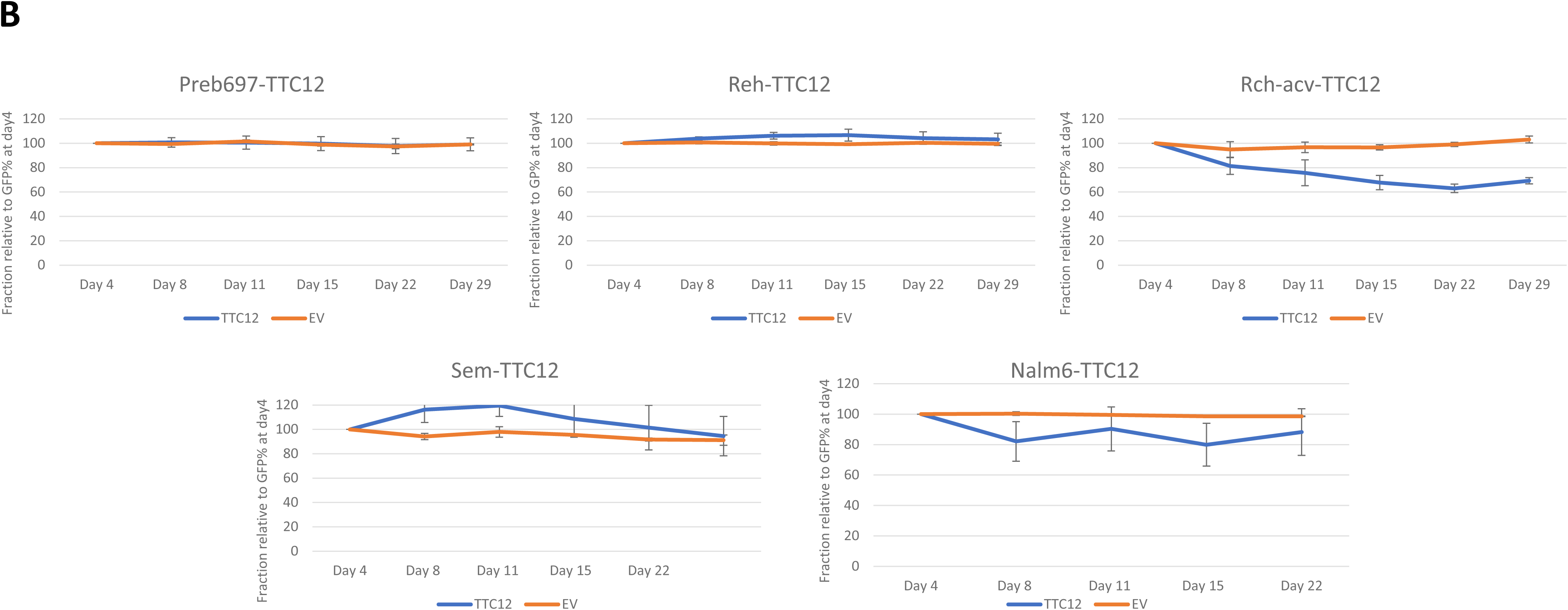
Transduction of ALL cell lines with lentiviral vectors expressing tumour suppressor candidates. A) Five ALL cell lines were transduced with THEM4 expressing lentiviral construct (or an empty vector control). While PreB697 cells exhibited a rapid decrease in THEM4 expressing cells, suggesting a strong negative selection, rapid loss of THEM4 expressing cells was not seen in any of the other four cell lines. B) Five ALL cell lines were transduced with TTC12 expressing lentiviral construct (or an empty vector control). Rch-acv and NALM-6 exhibit a slight fall in the fraction of TTC12 expressing cells. However, loss occurs at a low rate and the other three cell lines do not exhibit loss of TTC12 expressing cells.

